## Supplementary Information 1 for "Analysis of coagulation factor IX in bioreactor cell culture medium predicts yield and quality of the purified product"

**Python script to recalculate peptide and protein abundance with peptide level 1% FDR cut-off.**
This script is a modification of our previously published script [1] which uses the output from the SWATH Acquisition MicroApp in Peakview v2.2 (SCIEX).

```
import pandas as pd
import numpy as np
#------------INPUTS------------#

# Files need to be tab delimited .txt

openPEPTIDESfile = input("What Peptides file are you opening (written like:
PEPTIDES.txt): ")

openFDRfile = input("What FDR file are you opening (written like: FDR.txt): ")

filename = input("What would you like to name the output file (written like:
NEW_PROTEINS.csv): ")

#------------reading in PEPTIDES txt file------------#

MSdata0 = pd.read_csv(openPEPTIDESfile, sep="\t")

#------------reading in FDR txt file------------#

FDRdata1=pd.read_csv(openFDRfile, sep="\t",index_col=[0,1,4])

FDRdata1=FDRdata1[FDRdata1["Decoy"]==False]

#-----------checking FDR file for Peptides with appropriate FDRs --------#

MSdata1=[]

for i,r in MSdata0.iterrows():
    fdr_row = FDRdata1.loc[(r["Protein"], r["Peptide"], r["Precursor Charge"])]
    for c in range(4, len(FDRdata1.columns)):
        if fdr_row[FDRdata1.columns[c]] > 0.01:
            r[MSdata0.columns[c+1]] = np.nan
    MSdata1.append(r)


MSdata1 = pd.DataFrame(MSdata1)
MSdata1.to_csv(filename+"_peptide.txt", sep="\t", index=False)

#------------calculate new PROTEINS abundances----------#

Proteins = []
for i,d in MSdata1.groupby("Protein"):
    Proteins.append([i]+[d[d.columns[i]].sum() for i in range(5, len(d.columns))])
Proteins = pd.DataFrame(Proteins, columns=["Protein"]+[d.columns[i] for i in range(5,
len(d.columns))])

#------------create new txt file for new PROTEINS abundances------------#

Proteins.to_csv(filename, sep="\t", index=False)
```


1. Kerr, E.D., et al., *The intrinsic and regulated proteomes of barley seeds in response to fungal infection.* Anal Biochem, 2019. **580**: p. 30-35.
