## Supplementary Information 2 for "Analysis of coagulation factor IX in bioreactor cell culture medium predicts yield and quality of the purified product"

20190701_Co4Ra2NeuGcOglyLacNAc_20190509_BenSchulz_Luci_H1a_Ch-P.wiff_Byonic_D359_D364 1
20190701_Co4Ra2NeuGcOglyLacNAc_20190509_BenSchulz_Luci_H1a_Ch-P.wiff_Byonic_D359[+16] 2
20190701_Co4Ra2NeuGcOglyLacNAc_20190509_BenSchulz_Luci_H1a_Ch-P.wiff_Byonic_D364[+16] 3
20190701_Co4Ra2NeuGcOglyLacNAc_20190509_BenSchulz_Luci_H1a_Ch.wiff_Byonic_E33_E36_E40 4
20190701_Co4Ra2NeuGcOglyLacNAc_20190509_BenSchulz_Luci_H1a_Ch.wiff_Byonic_E33_E36_E40[+44] 5

20190701_Co4Ra2NeuGcOglyLacNAc_20190509_BenSchulz_Luci_H1a_Ch.wiff_Byonic_E33_E36_E40[+44]x2 6
20190701_Co4Ra2NeuGcOglyLacNAc_20190509_BenSchulz_Luci_H1a_Ch.wiff_Byonic_T179 7
20190701_Co4Ra2NeuGcOglyLacNAc_20190509_BenSchulz_Luci_H1a_Ch.wiff_Byonic_T179[+947] 8
20190701_Co4Ra2NeuGcOglyLacNAc_20190509_BenSchulz_Luci_H1a_Ch.wiff_Byonic_T179[+963] 9
Common4rare2NeuGc50_20181106_Schulz_Luci_H1aG.wiff_20181116_Byonic(1)__S53[+426]_S61[+802]_D64[+16]_S68[+146] 10

Common4rare2NeuGc50_20181106_Schulz_Luci_H1aG.wiff_20181116_Byonic(1)_D49[+16] 11
Common4rare2NeuGc50_20181106_Schulz_Luci_H1aG.wiff_20181116_Byonic(1)_D292[+16] 12
Common4rare2NeuGc50_20181106_Schulz_Luci_H1aG.wiff_20181116_Byonic(1)_E7_E9_E15_[+44]x2 13
Common4rare2NeuGc50_20181106_Schulz_Luci_H1aG.wiff_20181116_Byonic(1)_N258 14
Common4rare2NeuGc50_20181106_Schulz_Luci_H1aG.wiff_20181116_Byonic(1)_N258[+1] 15

Common4rare2NeuGc50_20181106_Schulz_Luci_H1aG.wiff_20181116_Byonic(1)_S53[+426]_S61[+802]_D64[+16]_D65[+16]_S 16
Common4rare2NeuGc50_20181106_Schulz_Luci_H1aGP_wiff_20181116_Byonic_D292[+16] 17
Common4rare2NeuGc50_20181106_Schulz_Luci_H1aGP.wiff_20181116_Byonic_N157[+1] 18
Common4rare2NeuGc50_20181106_Schulz_Luci_H1aGP.wiff_20181116_Byonic_N167[+1} 19
Common4rare2NeuGc50_20181106_Schulz_Luci_H1aGP.wiff_20181116_Byonic_S141[+656] 20

Common4rare2NeuGc50_20181106_Schulz_Luci_H1aGP.wiff_20181116_Byonic_S158[+947] 21
Common4rare2NeuGc50_20181106_Schulz_Luci_H1aGP.wiff_20181116_Byonic_Y155[+80] 22
Common4rare2NeuGc50_20181106_Schulz_Luci_H1aTP.wiff_20181119_Byonic_D64 23
Common4rare2NeuGc50_20181106_Schulz_Luci_H1aTP.wiff_20181119_Byonic_D64[+16] 24
Common4rare2NeuGc50_20181106_Schulz_Luci_H1aTP.wiff_20181119_Byonic_D85[+16] 25

Common4rare2NeuGc50_20181106_Schulz_Luci_H1aTP.wiff_20181119_Byonic_D186 26
Common4rare2NeuGc50_20181106_Schulz_Luci_H1aTP.wiff_20181119_Byonic_D186[+16] 27
Common4rare2NeuGc50_20181106_Schulz_Luci_H1aTP.wiff_20181119_Byonic_D203 28
Common4rare2NeuGc50_20181106_Schulz_Luci_H1aTP.wiff_20181119_Byonic_D203[+16] 29
Common4rare2NeuGc50_20181106_Schulz_Luci_H1aTP.wiff_20181119_Byonic_E7_E9_E15 30

Common4rare2NeuGc50_20181106_Schulz_Luci_H1aTP.wiff_20181119_Byonic_E7_E9_E15{+44] 31
Common4rare2NeuGc50_20181106_Schulz_Luci_H1aTP.wiff_20181119_Byonic_E7_E9_E15{+44]x2 32
Common4rare2NeuGc50_20181106_Schulz_Luci_H1aTP.wiff_20181119_Byonic_E7_E9_E15{+44]x3 33
Common4rare2NeuGc50_20181106_Schulz_Luci_H1aTP.wiff_20181119_Byonic_S141[+947] 34
Common4rare2NeuGc50_20181106_Schulz_Luci_H1aTP.wiff_20181119_Byonic_T38[+656] 35

Common4rare2NeuGc50_20181106_Schulz_Luci_H1aTP.wiff_20181119_Byonic_T38[+947] 36
Common4rare2NeuGc50_20181106_Schulz_Luci_H1aTP.wiff_20181119_Byonic_T38[+963] 37
Common4rare2NeuGc50_20181106_Schulz_Luci_H1bG.wiff_20181116_Byonic D47[+16] 38
Common4rare2NeuGc50_20181106_Schulz_Luci_H1bG.wiff_20181116_Byonic_D85 39
Common4rare2NeuGc50_20181106_Schulz_Luci_H1bG.wiff_20181116_Byonic_S53[+426]_S61[+802]_D64[+16] 40

Common4rare2NeuGc50_20181106_Schulz_Luci_H1bG.wiff_20181116_Byonic_S53[+426]_S61[+802] 41
Common4rare2NeuGc50_20181106_Schulz_Luci_H1bG.wiff_20181116_Byonic_S53[+426]_S61[+818]_D64[+16] 42
Common4rare2NeuGc50_20181106_Schulz_Luci_H1bG.wiff_20181116_Byonic_S110[+426]_T112[+802] 43
Common4rare2NeuGc50_20181106_Schulz_Luci_H1bG.wiff_20181116_Byonic_s123_s141_potential 44
Common4rare2NeuGc50_20181106_Schulz_Luci_H1bG.wiff_20181116_Byonic_S141 45

Common4rare2NeuGc50_20181106_Schulz_Luci_H1bG.wiff_20181116_Byonic_Y45_D47_D49 46
Common4rare2NeuGc50_20181106_Schulz_Luci_H1bG.wiff_20181116_Byonic_Y45[+80} 47
Common4rare2NeuGc50_20181106_Schulz_Luci_H1bG.wiff_20181116_Byonic_Y45[+80}sulfo_immonium 48
Common4rare2NeuGc50_20181106_Schulz_Luci_H1bGP.wiff_20181116_Byonic_D104 49
Common4rare2NeuGc50_20181106_Schulz_Luci_H1bGP.wiff_20181116_Byonic_D104{+16] 50

Common4rare2NeuGc50_20181106_Schulz_Luci_H1bGP.wiff_20181116_Byonic_D276-D292 51
Common4rare2NeuGc50_20181106_Schulz_Luci_H1bGP.wiff_20181116_Byonic_D276[+16] 52
Common4rare2NeuGc50_20181106_Schulz_Luci_H1bGP.wiff_20181116_Byonic_s365_672 T371 426 potential 53
Common4rare2NeuGc50_20181106_Schulz_Luci_H1bGP.wiff_20181116_Byonic_Y155[+80] 54
Common4rare2NeuGc50_20181106_Schulz_Luci_H1bT.wiff_20181119_Byonic_D203 55

Common4rare2NeuGc50_20181106_Schulz_Luci_H1bT.wiff_20181119_Byonic_D203[+16] 56
Common4rare2NeuGc50_20181106_Schulz_Luci_H1bT.wiff_20181119_Byonic_S141[+656] 57
ProteinPilot_E40 58
ProteinPilot_E40[+44] 59
ProteinPilot_S158[+656] 60

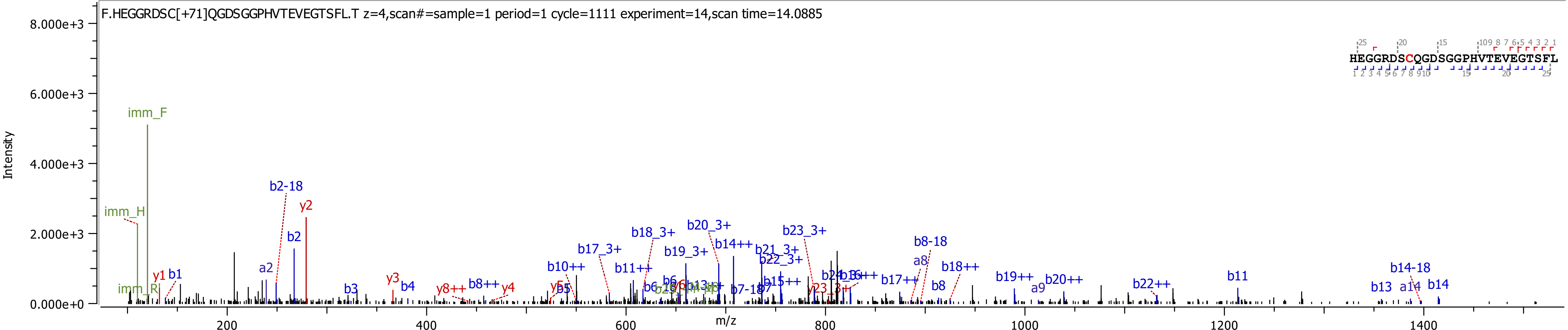
F.HEGGRDSC[+71]QGDSGGPHVTEVEGTSFL.T z=4,scan#=sample=1 period=1 cycle=1111 experiment=14,scan time=14.0885
HEGGRDSCQGDSGGPHVTEVEGTSFL
Intensity
m/z
imm_F
imm_H
imm_R
y1
b1
a2
b2-18
b2
y2
b3
y3
b4
y8++
b8++
y4
b10++
b17_3+
b18_3+
b11++
b19_3+
b20_3+
b14++
b21_3+
b22_3+
b23_3+
b15++
b7-18
b16++
b17++
b8-18
a8
b8
b18++
b19++
a9
b20++
b22++
b11
b14-18
b13
a14
b14

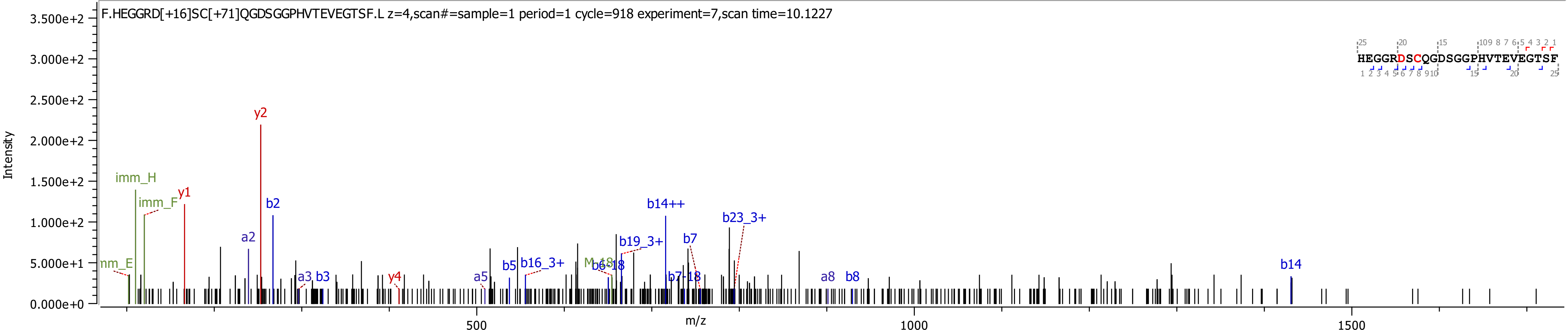

F.HEGGRD[+16]SC[+71]QGDSGGPHVTEVEGTSF.L z=4,scan#=sample=1 period=1 cycle=918 experiment=7,scan time=10.1227
Intensity
m/z
HEGGRDSCQGDSGGPHVTEVEGTSF
imm_H
imm_F
y1
y2
b2
a2
a3
b3
y4
a5
b5
b16_3+
M-18
b6-18
b19_3+
b14++
b7-18
b7
b23_3+
a8
b8
b14

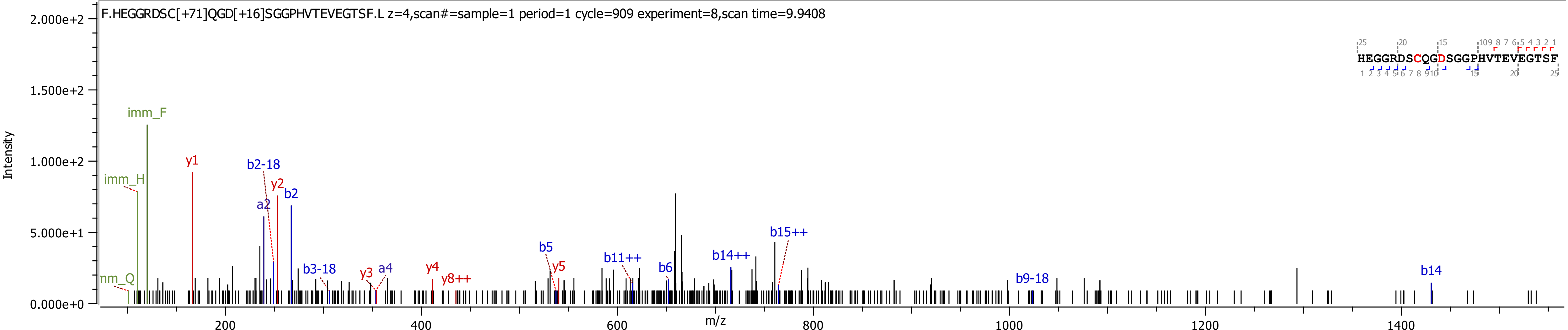
F.HEGGRDSC[+71]QGD[+16]SGGPHVTEVEGTSF.L z=4,scan#=sample=1 period=1 cycle=909 experiment=8,scan time=9.9408
HEGGRDSCQGDSGGPHVTEVEGTSF
Intensity
m/z
imm_F
imm_H
nm_Q
y1
b2-18
y2
b2
a2
b3-18
y3
a4
y4
y8++
b5
y5
b11++
b6
b14++
b15++
b9-18
b14
2.000e+2
1.500e+2
1.000e+2
5.000e+1
0.000e+0
200
400
600
800
1000
1200
1400

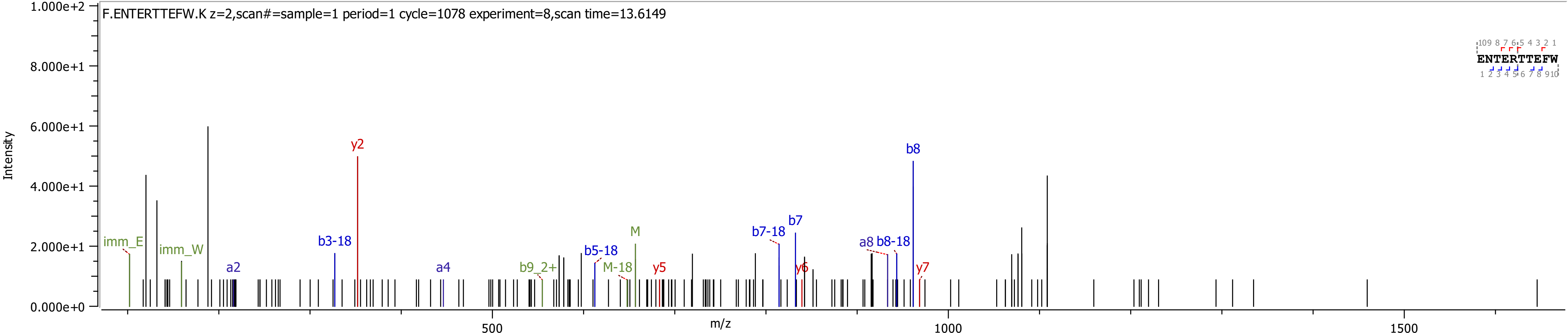
F.ENTERTTEFW.K z=2,scan#=sample=1 period=1 cycle=1078 experiment=8,scan time=13.6149
Intensity
m/z
imm_E
imm_W
a2
b3-18
y2
a4
b9_2+
b5-18
M-18
M
y5
b7-18
b7
y6
a8
b8-18
b8
y7
ENTERTTEFW
0.000e+0
2.000e+1
4.000e+1
6.000e+1
8.000e+1
1.000e+2
500
1000
1500

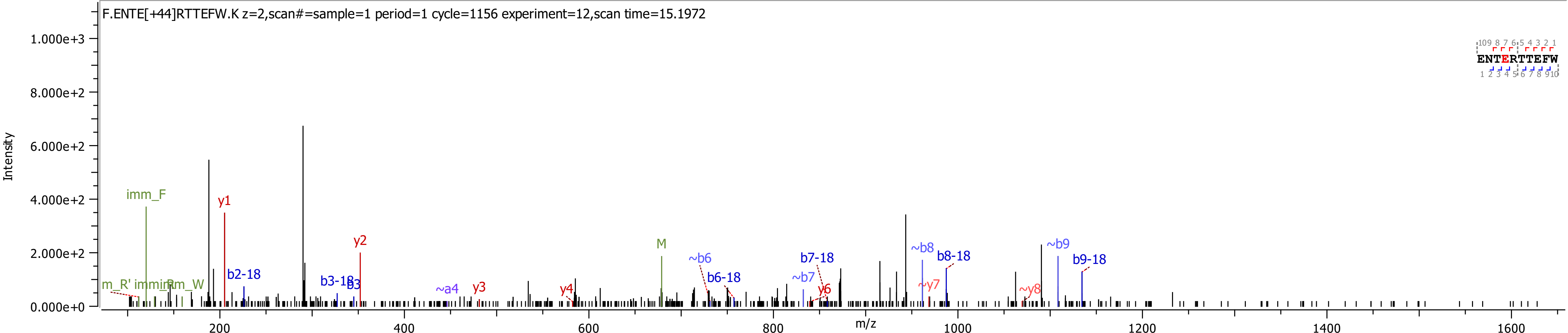
F.ENTE[+44]RTTEFW.K z=2,scan#=sample=1 period=1 cycle=1156 experiment=12,scan time=15.1972
Intensity
m/z
imm_F
y1
y2
y3
y4
y6
~y7
~y8
m_R'
imm_R
imm_W
b2-18
b3-18
b3
~a4
M
~b6
b6-18
~b7
b7-18
~b8
b8-18
~b9
b9-18
ENTERTTEFW

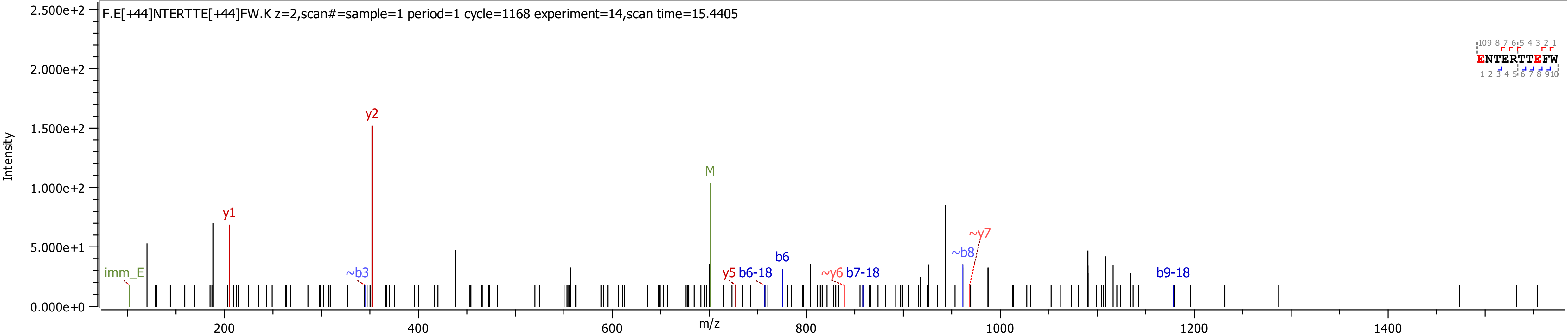
F.E[+44]NTERTTE[+44]FW.K z=2,scan#=sample=1 period=1 cycle=1168 experiment=14,scan time=15.4405
Intensity
m/z
2.500e+2
2.000e+2
1.500e+2
1.000e+2
5.000e+1
0.000e+0
200
400
600
800
1000
1200
1400
imm_E
y1
~b3
y2
M
y5
b6-18
b6
~y6
b7-18
~b8
~y7
b9-18
ENTERTTEFW

F.NDFTRVVGGEDAKPGQFPW.Q z=3,scan#=sample=1 period=1 cycle=1389 experiment=4,scan time=20.0154

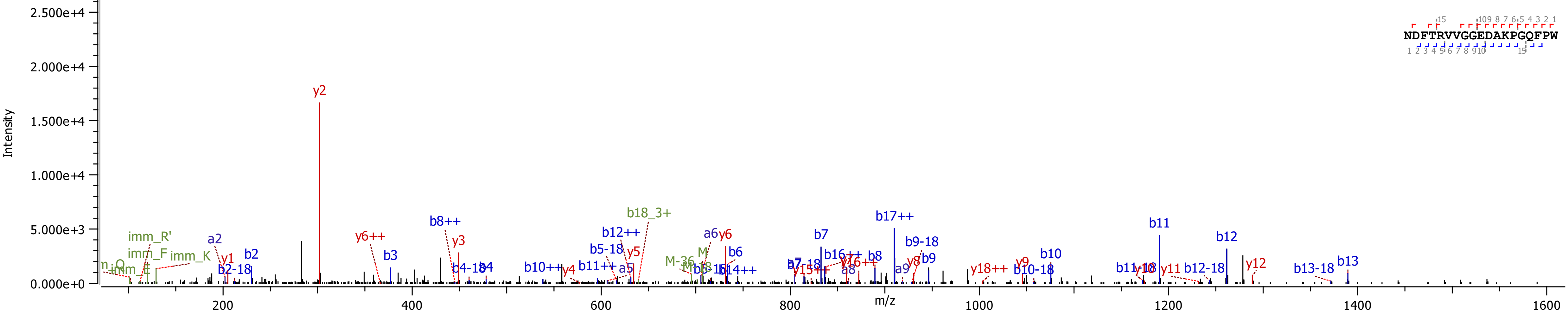

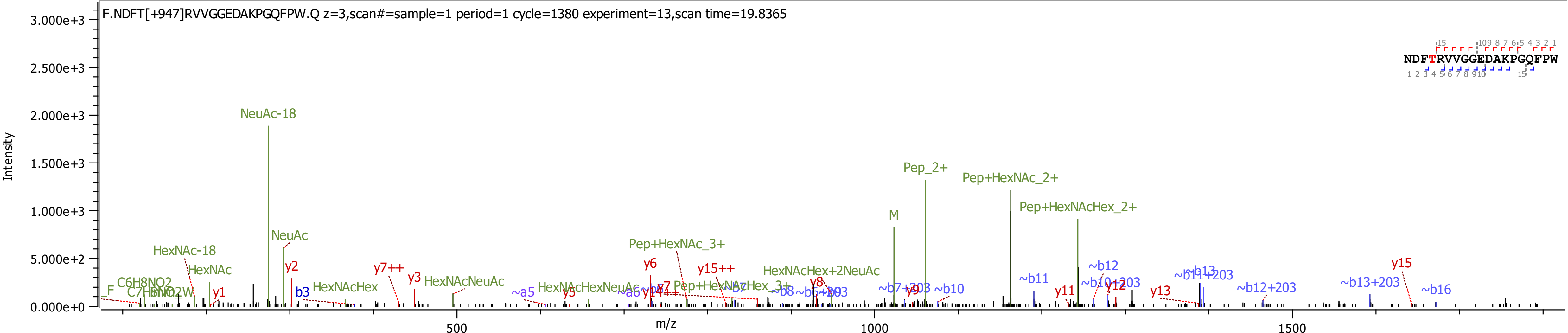

F.NDFT[+947]RVVGGEDAKPGQFPW.Q z=3,scan#=sample=1 period=1 cycle=1380 experiment=13,scan time=19.8365
NDFTRVVGGEDAKPGQFPW
Intensity
3.000e+3
2.500e+3
2.000e+3
1.500e+3
1.000e+3
5.000e+2
0.000e+0
m/z
500
1000
1500
NeuAc-18
NeuAc
HexNAc-18
HexNAc
C6H8NO2
C7H8NO2W
F
y1
y2
b3
HexNAcHex
y7++
y3
HexNAcNeuAc
~a5
y5
HexNAcHexNeuAc
y6
Pep+HexNAc_3+
y15++
Pep+HexNAcHex_3+
HexNAcHex+2NeuAc
y8
M
Pep_2+
Pep+HexNAc_2+
Pep+HexNAcHex_2+
~b7+203
y9
~b10
~b11
~b12
y11
~b10+203
y12
y13
~b11+203
~b12+203
~b13+203
y15
~b16

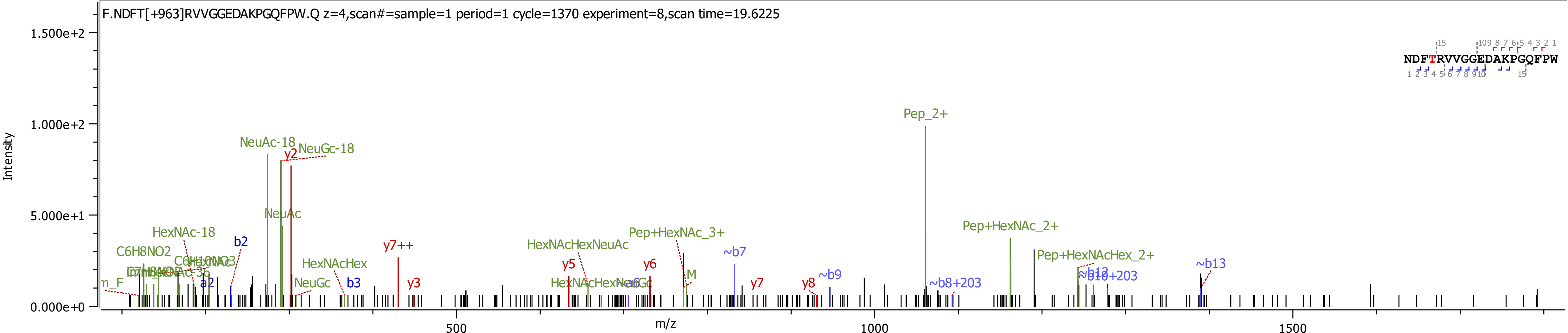
F.NDFT[+963]RVVGGEDAKPGQFPW.Q z=4,scan#=sample=1 period=1 cycle=1370 experiment=8,scan time=19.6225
NDFTRVVGGEDAKPGQFPW
Intensity
m/z
1.500e+2
1.000e+2
5.000e+1
0.000e+0
500
1000
1500
Pep_2+
NeuAc-18
NeuGc-18
y2
NeuAc
HexNAc-18
C6H8NO2
b2
a2
b3
y3
y7++
HexNAcHex
NeuGc
y5
y6
y7
y8
HexNAcHexNeuAc
Pep+HexNAc_3+
~b7
~b9
~b8+203
Pep+HexNAc_2+
Pep+HexNAcHex_2+
~b10+203
~b13
M
m_F

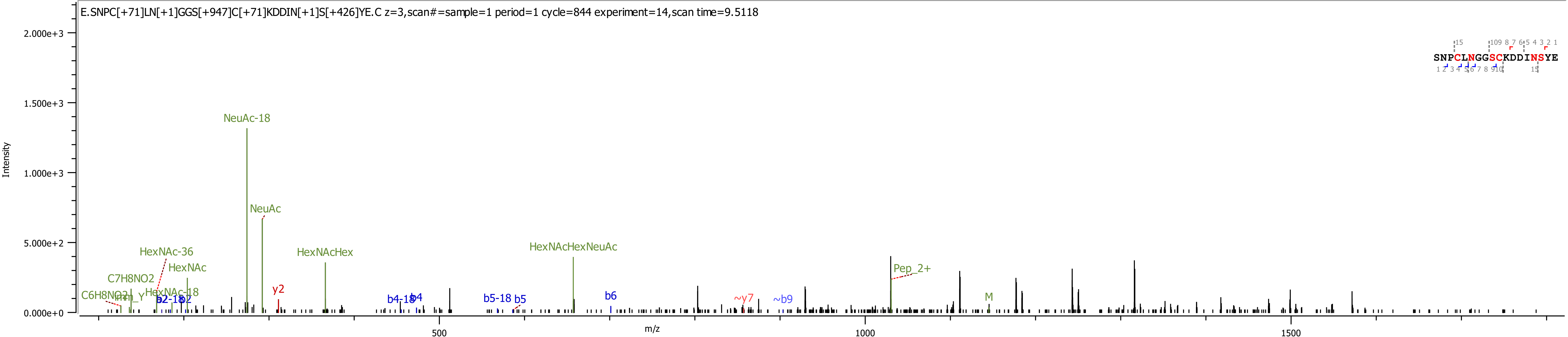
E.SNPC[+71]LN[+1]GGS[+947]C[+71]KDDIN[+1]S[+426]YE.C z=3,scan#=sample=1 period=1 cycle=844 experiment=14,scan time=9.5118
SNPCLNGGSCKDDINSYE
Intensity
m/z
2.000e+3
1.500e+3
1.000e+3
5.000e+2
0.000e+0
500
1000
1500
NeuAc-18
NeuAc
HexNAcHex
HexNAcHexNeuAc
HexNAc-36
HexNAc
C7H8NO2
HexNAc-18
C6H8NO2
y2
b4-18
b4
b5-18
b5
b6
~y7
~b9
Pep_2+
M

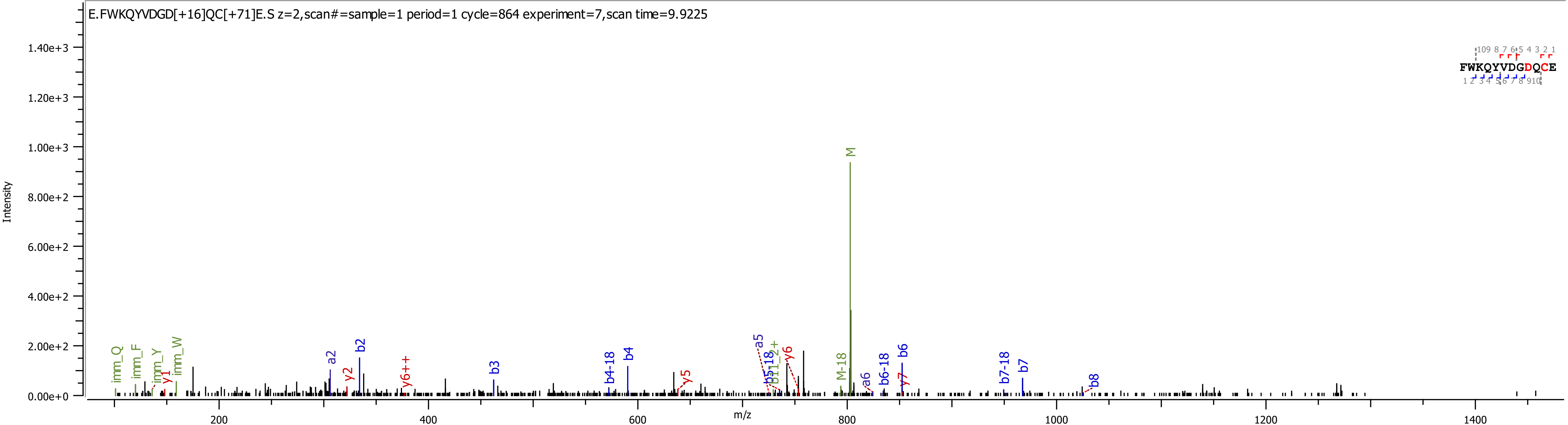

E.FWKQYVDGD[+16]QC[+71]E.S z=2,scan#=sample=1 period=1 cycle=864 experiment=7,scan time=9.9225
Intensity
m/z
FWKQYVDGDQCE
M
M-18
imm_Q
imm_F
imm_Y
imm_W
y1
y2
y5
y6
y7
y6++
a2
a5
a6
b2
b3
b4
b4-18
b6
b6-18
b7
b7-18
b8

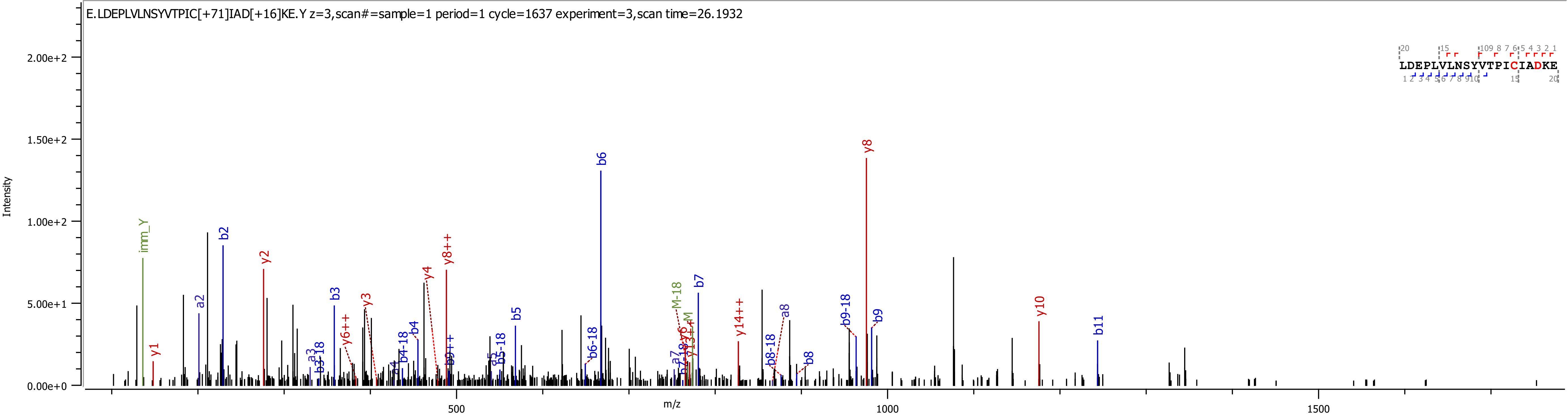
E.LDEPLVLNSYVTPIC[+71]IAD[+16]KE.Y z=3,scan#=sample=1 period=1 cycle=1637 experiment=3,scan time=26.1932
Intensity
m/z
LDEPLVLNSYVTPICIADKE
imm_Y
y1
a2
b2
y2
b3
y3
y6++
y4
y8++
b6
b5
b4
b4-18
b5-18
b6-18
M-18
b7
y14++
a8
b8-18
b8
b9-18
b9
y8
y10
b11
0.00e+0
5.00e+1
1.00e+2
1.50e+2
2.00e+2
500
1000
1500

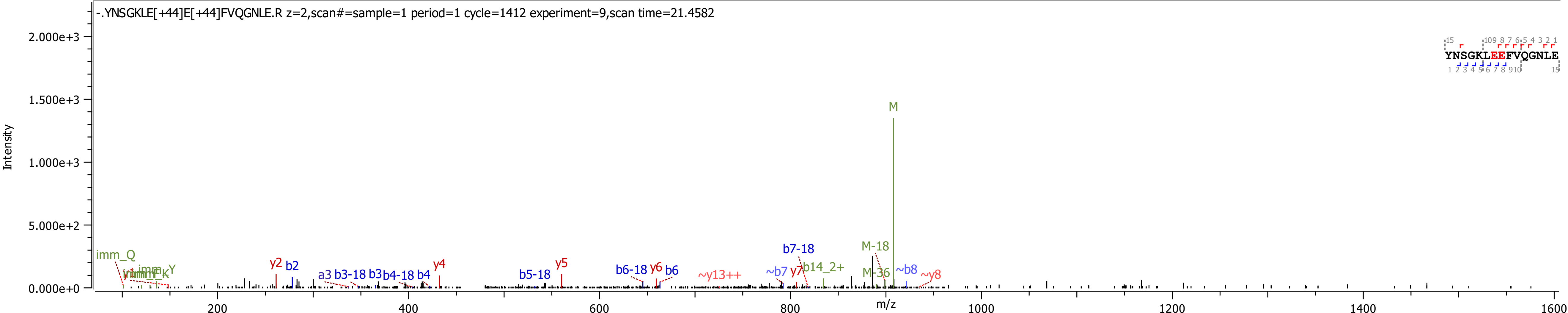
-.YNSGKLE[+44]E[+44]FVQGNLE.R z=2,scan#=sample=1 period=1 cycle=1412 experiment=9,scan time=21.4582
Intensity
2.000e+3
1.500e+3
1.000e+3
5.000e+2
0.000e+0
m/z
200
400
600
800
1000
1200
1400
1600
YNSGKLEEFVQGNLE
M
M-18
M-36
imm_Q
imm_Y
imm_K
y2
b2
a3
b3-18
b3
b4-18
b4
y4
b5-18
y5
b6-18
y6
b6
~y13++
~b7
y7
b7-18
b14_2+
~b8
~y8

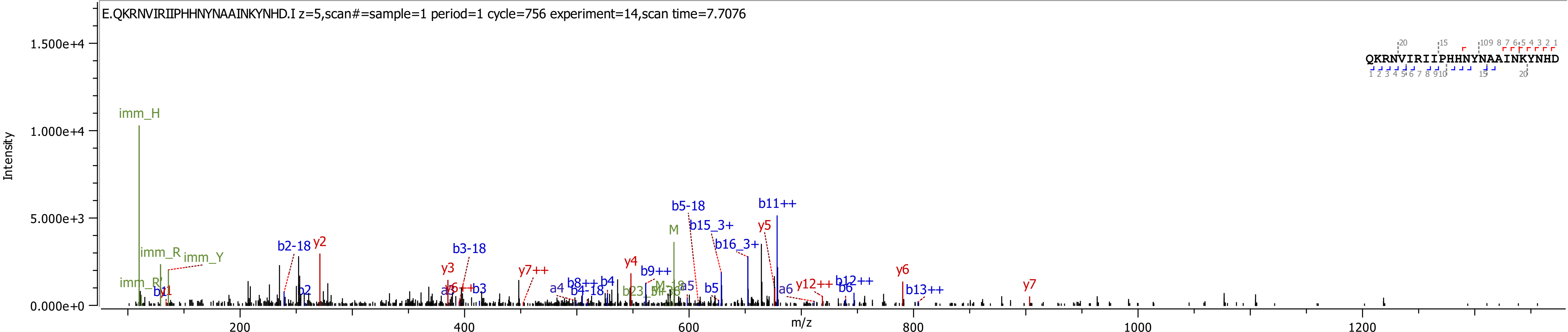
E.QKRNVIRIIPHHNYNAAINKYNHD.I z=5,scan#=sample=1 period=1 cycle=756 experiment=14,scan time=7.7076
Intensity
m/z
1.500e+4
1.000e+4
5.000e+3
0.000e+0
200
400
600
800
1000
1200
QKRNVIRIIPHHNYNAAINKYNHD
imm_H
imm_R
imm_Y
b2-18
y2
b3-18
y3
y7++
y4
b9++
M
b5-18
b15_3+
b16_3+
y5
b11++
y12++
b12++
y6
b13++
y7
a4
b8++
b4
b4-18
a5
b5
a6
b6
b3
b2

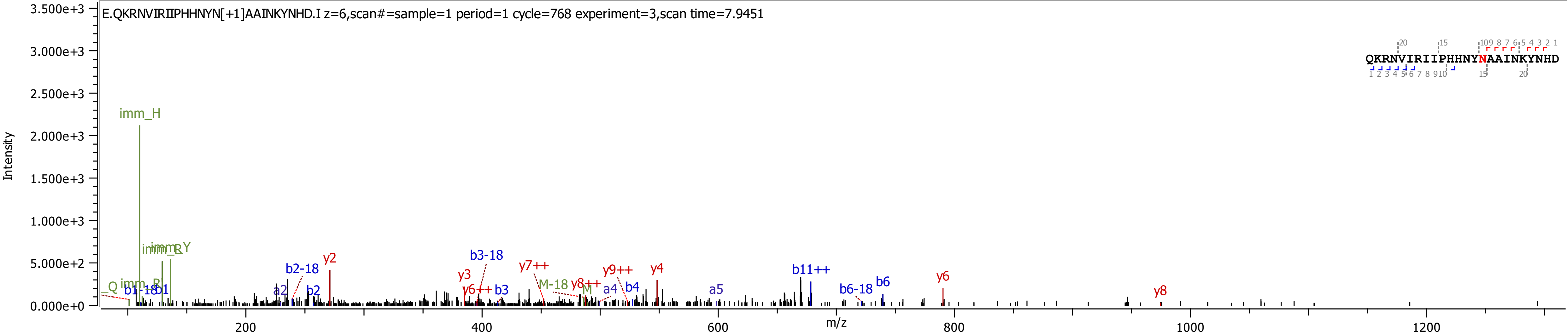

E.QKRNVIRIIPHHNYN[+1]AAINKYNHD.I z=6,scan#=sample=1 period=1 cycle=768 experiment=3,scan time=7.9451
QKRNVIRIIPHHNYNAAINKYNHD
Intensity
m/z
3.500e+3
3.000e+3
2.500e+3
2.000e+3
1.500e+3
1.000e+3
5.000e+2
0.000e+0
200
400
600
800
1000
1200
imm_H
imm_Y
imm_R
y2
b2-18
b3-18
y7++
y9++
y4
b11++
y6
y3
y6++
b3
M-18
y8++
M
a4
b4
a5
b6-18
b6
y8
Q
b1-18
b1
a2
b2

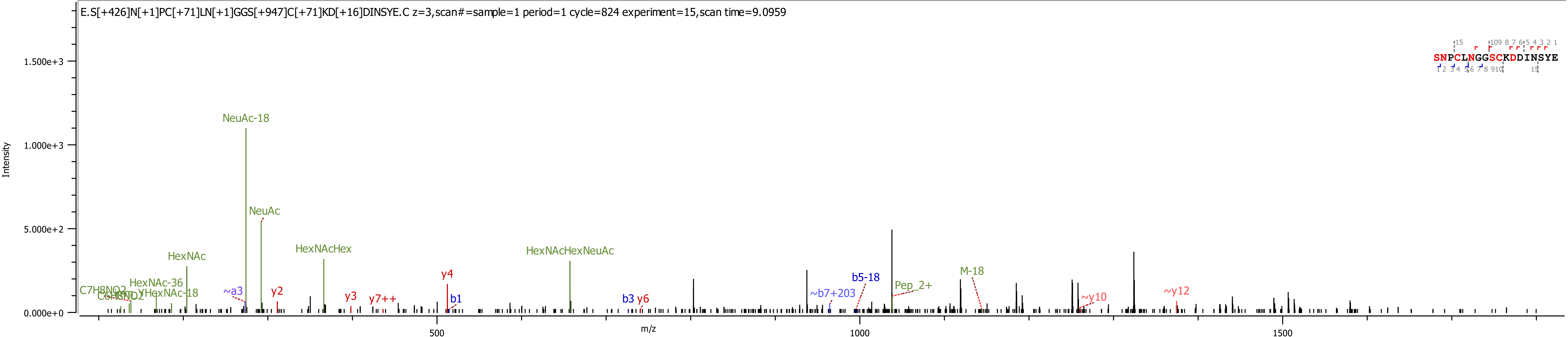
E.S[+426]N[+1]PC[+71]LN[+1]GGS[+947]C[+71]KD[+16]DINSYE.C z=3,scan#=sample=1 period=1 cycle=824 experiment=15,scan time=9.0959
Intensity
1.500e+3
1.000e+3
5.000e+2
0.000e+0
m/z
500
1000
1500
SNPCLNGGSCKDDINSYE
NeuAc-18
NeuAc
HexNAc
HexNAcHex
HexNAcHexNeuAc
HexNAc-36
HexNAc-18
C7H8NO2
~a3
y2
y3
y7++
y4
b1
b3
y6
~b7+203
b5-18
Pep_2+
M-18
~y10
~y12

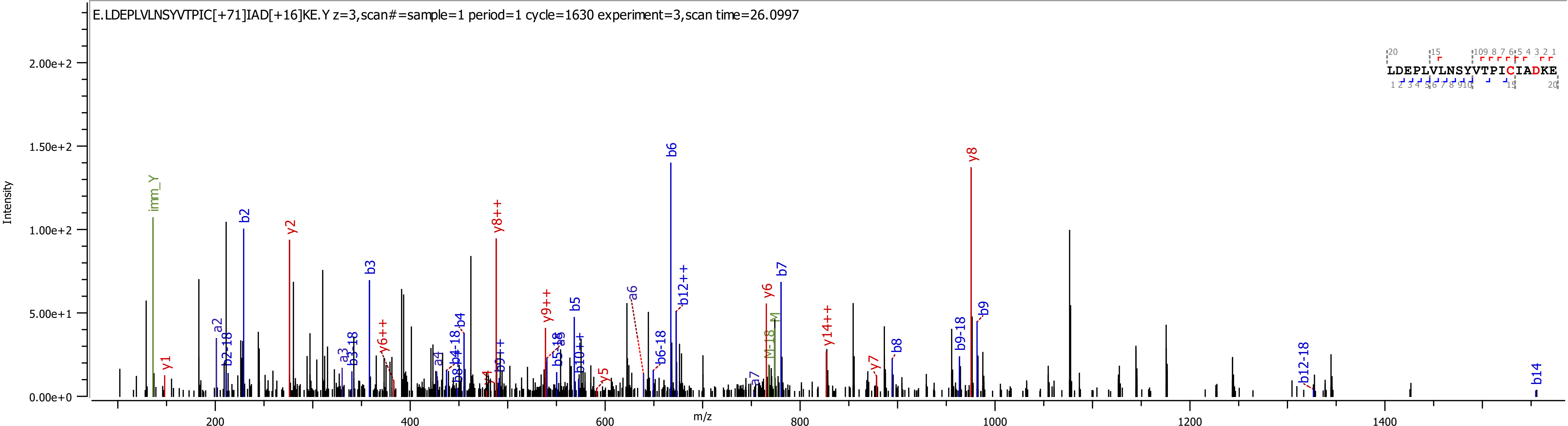

E.LDEPLVLNSYVTPIC[+71]IAD[+16]KE.Y z=3,scan#=sample=1 period=1 cycle=1630 experiment=3,scan time=26.0997
LDEPLVLNSYVTPICIADKE
Intensity
m/z

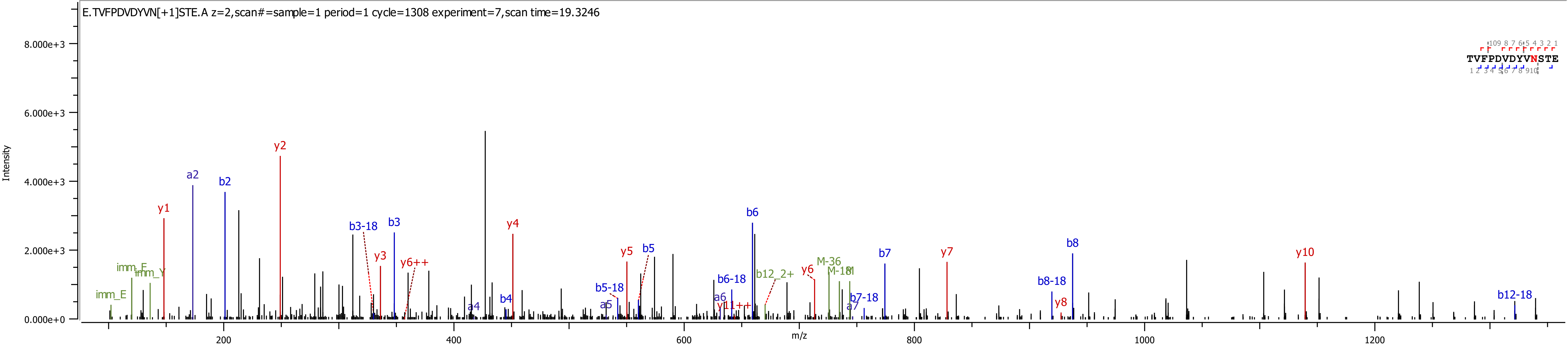
E.TVFPDVDYVN[+1]STE.A z=2,scan#=sample=1 period=1 cycle=1308 experiment=7,scan time=19.3246
TVFPDVDYVNSTE
Intensity
8.000e+3
6.000e+3
4.000e+3
2.000e+3
0.000e+0
m/z
200
400
600
800
1000
1200
imm_E
imm_E
imm_Y
y1
a2
b2
y2
b3-18
b3
y3
y6++
a4
b4
y4
b5-18
a5
y5
b5
a6
b6-18
y11++
b6
b12_2+
y6
M-36
M-18
M
b7-18
a7
b7
y7
b8-18
y8
b8
y10
b12-18

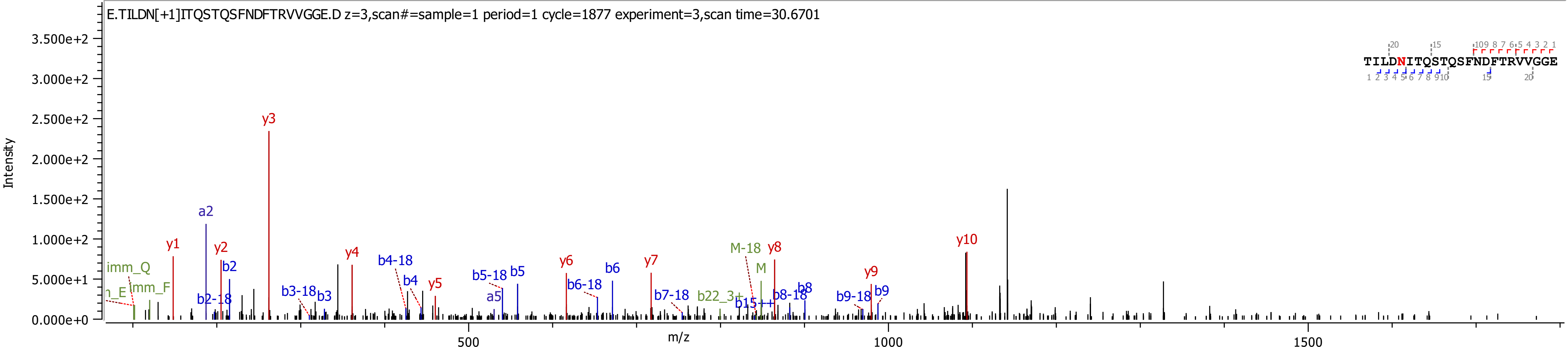

E.TILDN[+1]ITQSTQSFNDFTRVVGGE.D z=3,scan#=sample=1 period=1 cycle=1877 experiment=3,scan time=30.6701
TILDNITQSTQSFNDFTRVVGGE
Intensity
m/z
3.500e+2
3.000e+2
2.500e+2
2.000e+2
1.500e+2
1.000e+2
5.000e+1
0.000e+0
500
1000
1500
y1
y2
y3
y4
y5
y6
y7
y8
y9
y10
a2
a5
b2
b2-18
b3
b3-18
b4
b4-18
b5
b5-18
b6
b6-18
b7-18
b8
b8-18
b9
b9-18
b15++
b22_3+
M
M-18
imm_Q
imm_F

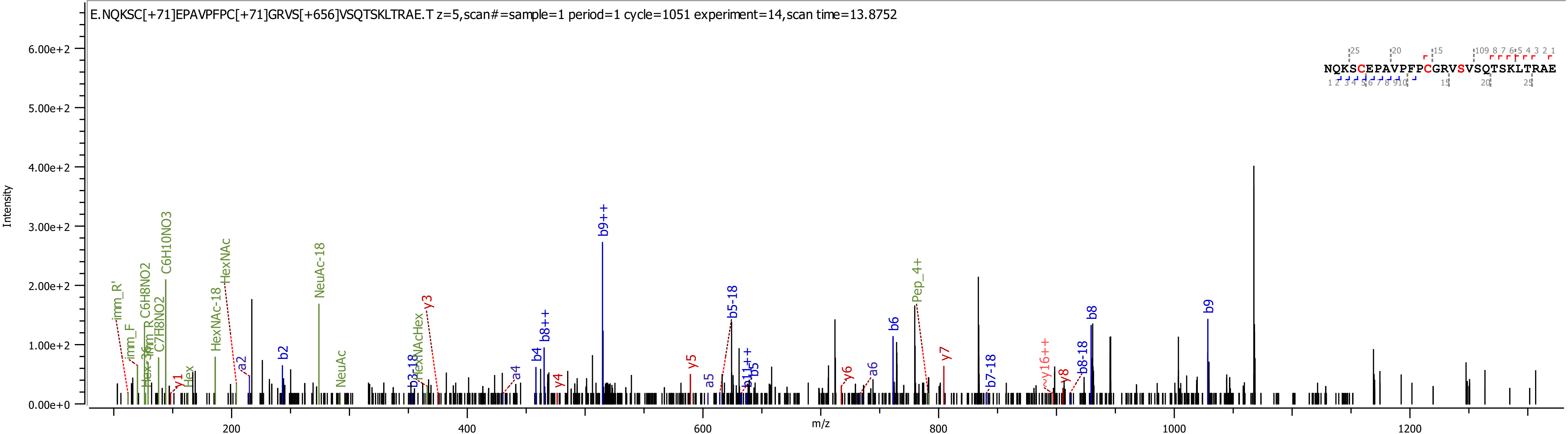
E.NQKSC[+71]EPAVPFPC[+71]GRVS[+656]VSQTSKLTRAE.T z=5,scan#=sample=1 period=1 cycle=1051 experiment=14,scan time=13.8752
NQKSCEPAVPFPCGRVSVSQTSKLTRAE
Intensity
m/z
6.00e+2
5.00e+2
4.00e+2
3.00e+2
2.00e+2
1.00e+2
0.00e+0
200
400
600
800
1000
1200
imm_R'
imm_F
C6H8NO2
C7H8NO2
C6H10NO3
HexNAc-18
HexNAc
NeuAc-18
NeuAc
Hex
y1
a2
b2
b3-18
HexNAcHex
y3
a4
b4
b8++
y4
b9++
y5
a5
b5-18
b11++
b5
y6
a6
b6
Pep_4+
y7
b7-18
y16++
y8
b8-18
b8
b9

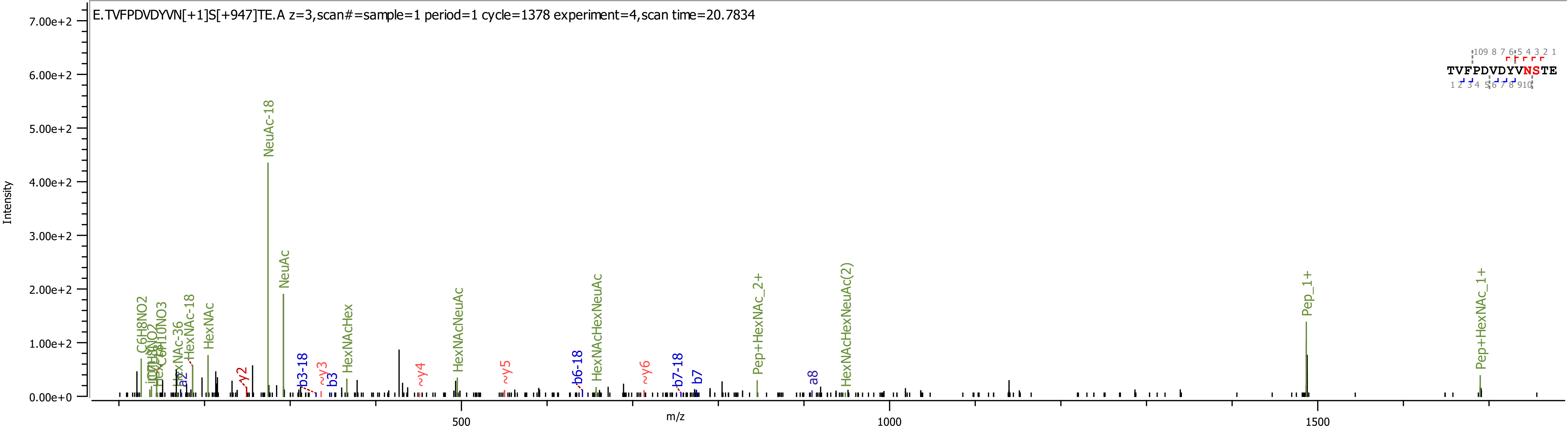
E.TVFPDVDYVN[+1]S[+947]TE.A z=3,scan#=sample=1 period=1 cycle=1378 experiment=4,scan time=20.7834
TVFPDVDYVNSTE
Intensity
7.00e+2
6.00e+2
5.00e+2
4.00e+2
3.00e+2
2.00e+2
1.00e+2
0.00e+0
m/z
500
1000
1500
NeuAc-18
NeuAc
C6H8NO2
HexNAc-36
HexNAc-18
HexNAc
y2
b3-18
~y3
b3
HexNAcHex
~y4
HexNAcNeuAc
~y5
b6-18
HexNAcHexNeuAc
~y6
b7-18
b7
Pep+HexNAc_2+
a8
HexNAcHexNeuAc(2)
Pep_1+
Pep+HexNAc_1+

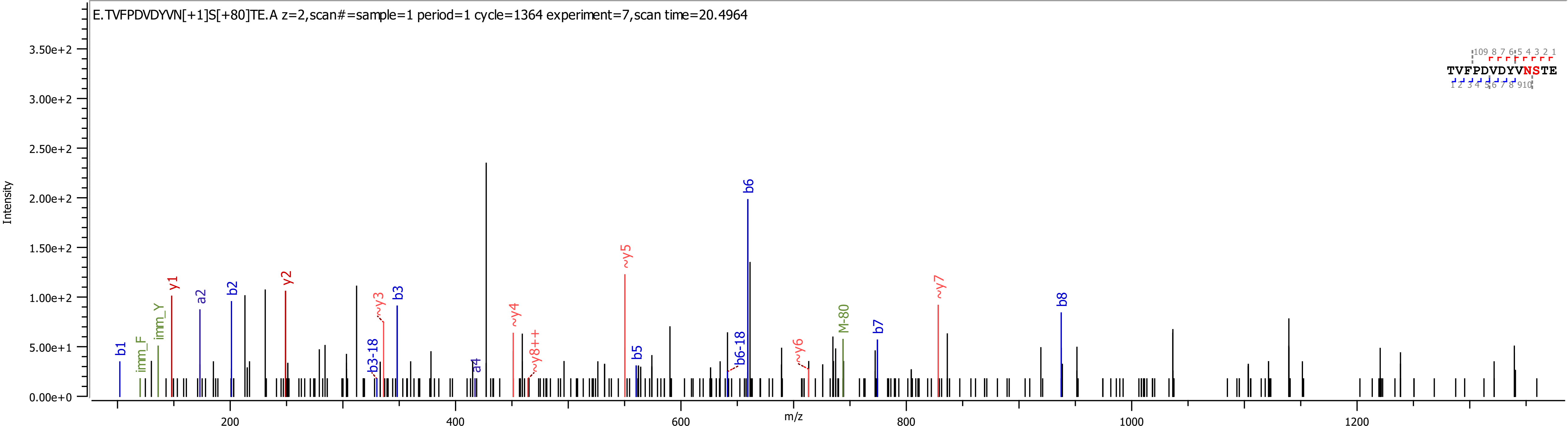
E.TVFPDVDYVN[+1]S[+80]TE.A z=2,scan#=sample=1 period=1 cycle=1364 experiment=7,scan time=20.4964
TVFPDVDYVNSTE
Intensity
m/z
b1
imm_F
imm_Y
y1
a2
b2
y2
b3-18
~y3
b3
a4
~y4
~y8++
~y5
b5
b6-18
b6
~y6
M-80
b7
~y7
b8

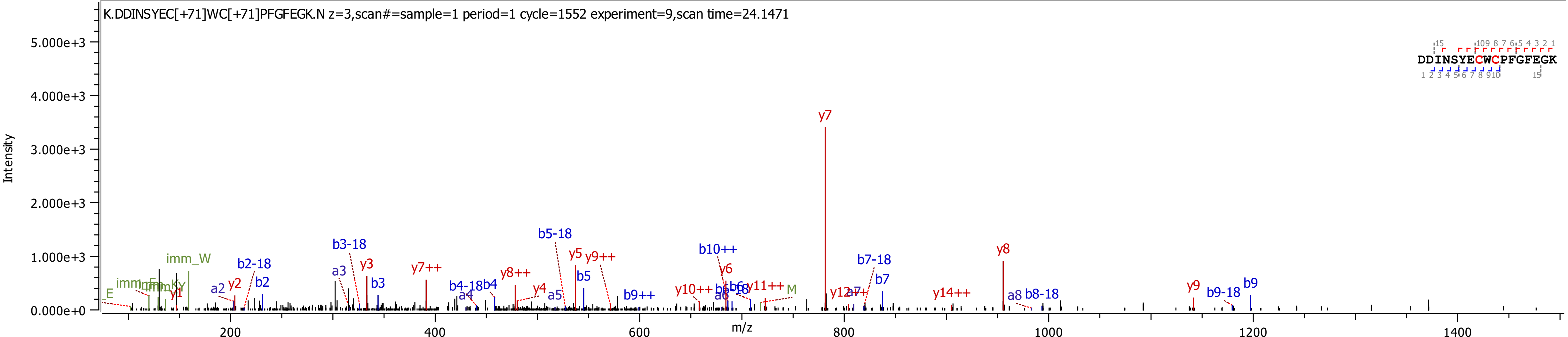
K.DDINSYEC[+71]WC[+71]PFGFEGK.N z=3,scan#=sample=1 period=1 cycle=1552 experiment=9,scan time=24.1471
Intensity
m/z
DDINSYECWCPFGFEGK
y7
y8
y5
y9++
b5
b5-18
b10++
y6
b7-18
b7
y3
b3-18
a3
b3
y7++
y8++
y4
b4-18
b4
b2-18
b2
imm_W
y2
a2
y1
E
y9
b9-18
b9
b9++
y10++
y11++
M
y12
y14++
a8
b8-18
a4
a5
0.000e+0
1.000e+3
2.000e+3
3.000e+3
4.000e+3
5.000e+3
200
400
600
800
1000
1200
1400

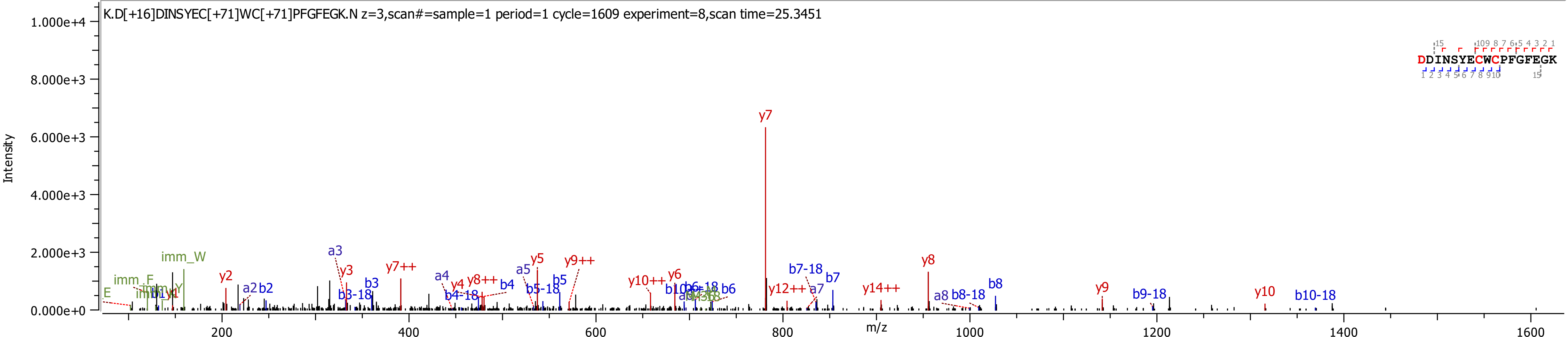
K.D[+16]DINSYEC[+71]WC[+71]PFGFEGK.N z=3,scan#=sample=1 period=1 cycle=1609 experiment=8,scan time=25.3451
DDINSYECWCPFGFEGK
Intensity
m/z
y7
y8
y5
y9++
y7++
y2
y3
a3
imm_W
imm_F
b7-18
b7
b8
y9
y10
b9-18
b10-18
y14++
y12++
a7
a8
b8-18
y10++
y6
b6
b5
b5-18
a5
b4
y8++
y4
a4
b4-18
b3
b3-18
a2
b2
E

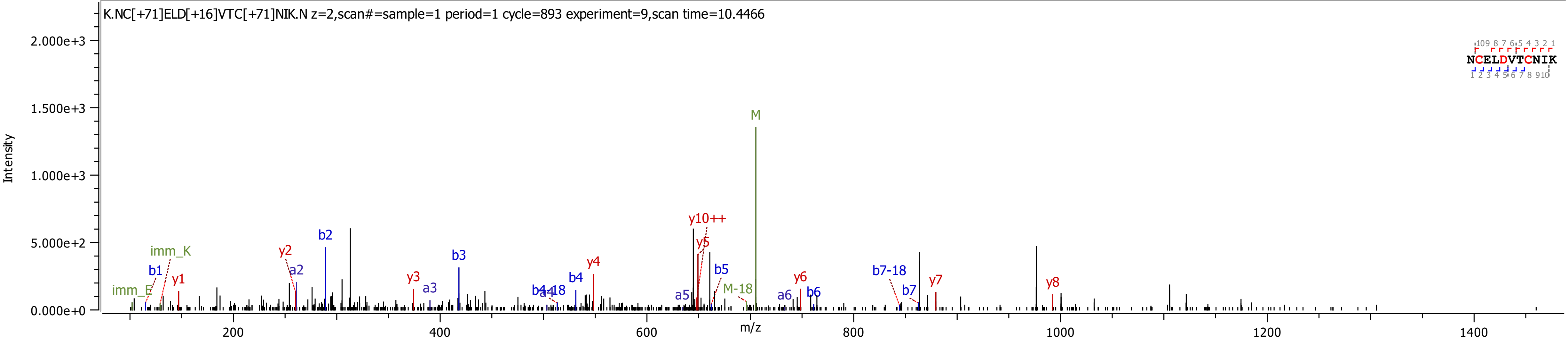
K.NC[+71]ELD[+16]VTC[+71]NIK.N z=2,scan#=sample=1 period=1 cycle=893 experiment=9,scan time=10.4466
Intensity
m/z
2.000e+3
1.500e+3
1.000e+3
5.000e+2
0.000e+0
200
400
600
800
1000
1200
1400
NCELDVTCNIK
M
y10++
y5
b2
a2
y2
imm_K
b1
y1
imm_E
b3
y3
a3
b4-18
a4
b4
y4
a5
b5
M-18
a6
b6
y6
b7-18
b7
y7
y8

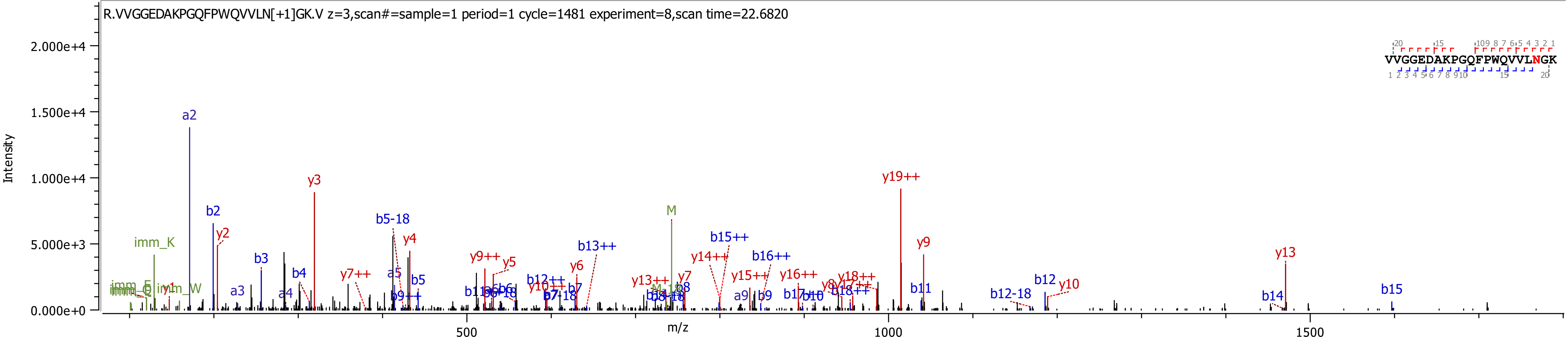
R.VVGGEDAKPGQFPWQVVLN[+1]GK.V z=3,scan#=sample=1 period=1 cycle=1481 experiment=8,scan time=22.6820
VVGGEDAKPGQFPWQVVLNGK
Intensity
m/z
2.000e+4
1.500e+4
1.000e+4
5.000e+3
0.000e+0
500
1000
1500
a2
b2
y2
y3
imm_K
b3
b4
a3
a4
b5-18
y4
a5
b5
y7++
y9++
y5
y6
b12++
b13++
M
y14++
b15++
b16++
y15++
y16++
y18++
y19++
y9
b11
y13
b12
y10
b12-18
b14
b15
y7
b8
a9
b9
y13++
b7

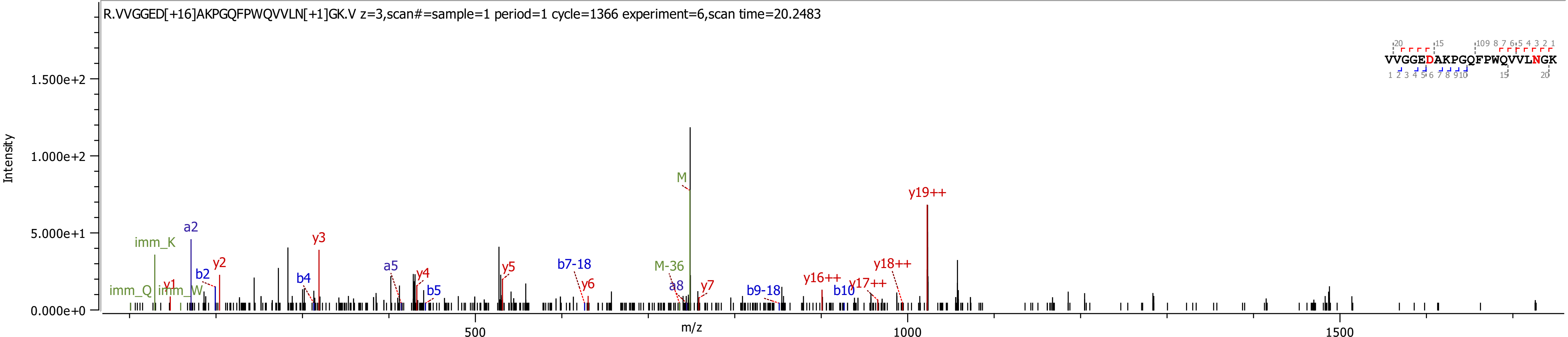
R.VVGGED[+16]AKPGQFPWQVVLN[+1]GK.V z=3,scan#=sample=1 period=1 cycle=1366 experiment=6,scan time=20.2483
VVGGEDAKPGQFPWQVVLNGK
Intensity
1.500e+2
1.000e+2
5.000e+1
0.000e+0
m/z
500
1000
1500
imm_K
imm_Q
imm_W
y1
a2
b2
y2
b4
y3
a5
y4
b5
y5
b7-18
y6
M
M-36
a8
y7
b9-18
y16++
b10
y17++
y18++
y19++

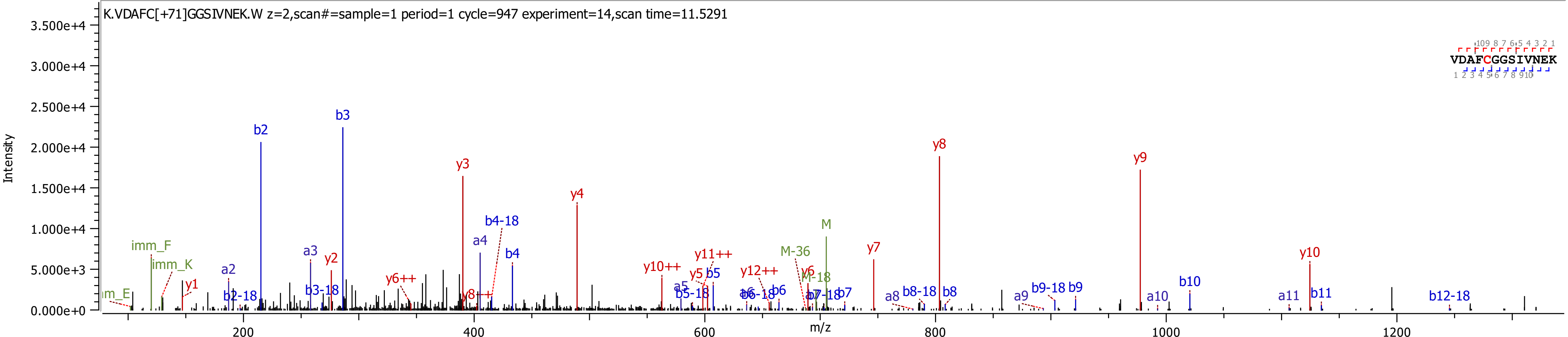

K.VDAFC[+71]GGSIVNEK.W z=2,scan#=sample=1 period=1 cycle=947 experiment=14,scan time=11.5291
VDAFCGGSIVNEK
Intensity
m/z
b2
b3
y3
y4
y8
y9
y7
y10
M
M-36
M-18
b4-18
a4
b4
a3
a2
y2
y1
imm_F
imm_K
b2-18
b3-18
y6++
y8++
y10++
y11++
y12++
y5
b5
a5
b5-18
b6-18
b6
y6
b7-18
b7
a8
b8-18
b8
a9
b9-18
b9
a10
b10
a11
b11
b12-18

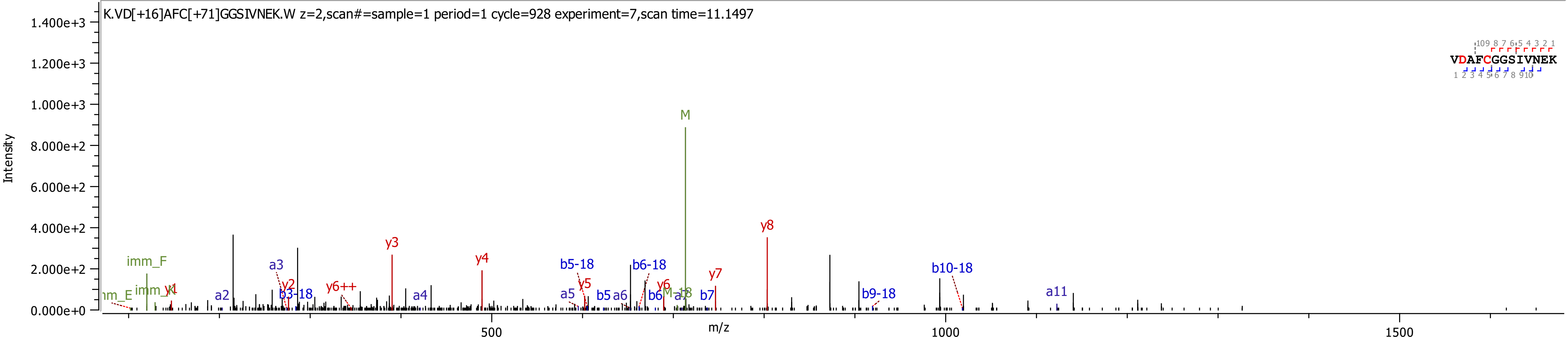

K.VD[+16]AFC[+71]GGSIVNEK.W z=2,scan#=sample=1 period=1 cycle=928 experiment=7,scan time=11.1497
Intensity
m/z
0.000e+0
2.000e+2
4.000e+2
6.000e+2
8.000e+2
1.000e+3
1.200e+3
1.400e+3
500
1000
1500
M
y8
y3
y4
imm_F
a3
y2
b3-18
y6++
a2
a4
b5-18
y5
a5
b5
a6
b6-18
y6
b6
a7
M-18
b7
y7
b9-18
b10-18
a11
VDAFCGGSIVNEK

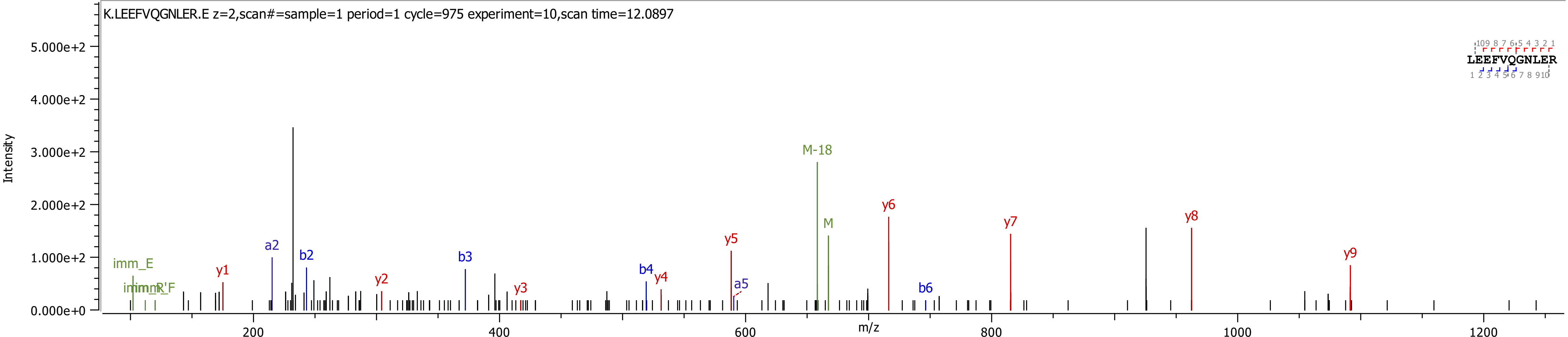
K.LEEFVQGNLER.E z=2,scan#=sample=1 period=1 cycle=975 experiment=10,scan time=12.0897
LEEFVQGNLER
Intensity
m/z
5.000e+2
4.000e+2
3.000e+2
2.000e+2
1.000e+2
0.000e+0
200
400
600
800
1000
1200
imm_E
imm_R'F
y1
a2
b2
y2
b3
y3
b4
y4
y5
a5
M-18
M
y6
b6
y7
y8
y9

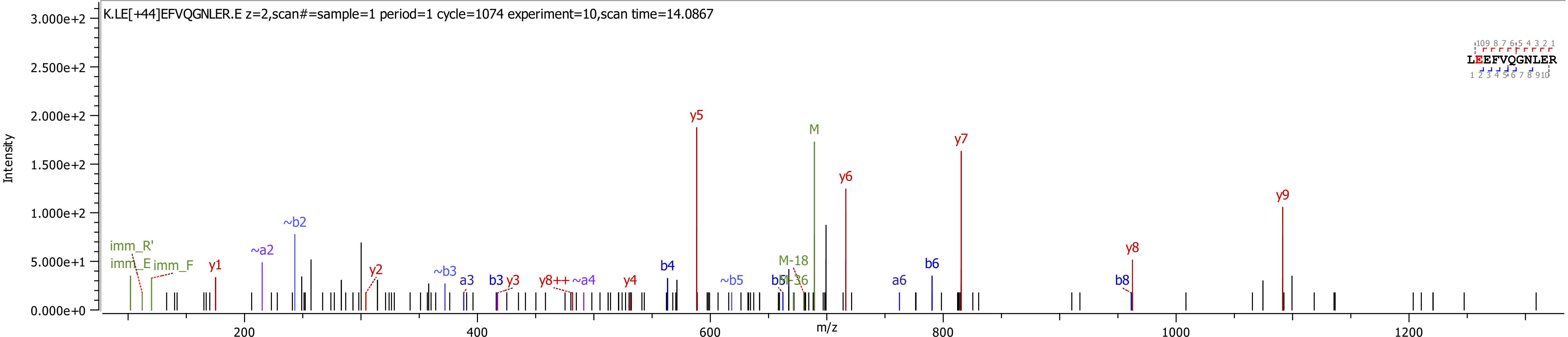
K.LE[+44]EFVQGNLER.E z=2,scan#=sample=1 period=1 cycle=1074 experiment=10,scan time=14.0867
Intensity
3.000e+2
2.500e+2
2.000e+2
1.500e+2
1.000e+2
5.000e+1
0.000e+0
m/z
200
400
600
800
1000
1200
LEEFVQGNLER
imm_R'
imm_E
imm_F
y1
~a2
~b2
y2
~b3
a3
b3
y3
y8++
~a4
y4
b4
y5
~b5
b5
M-36
M-18
M
y6
a6
b6
y7
b8
y8
y9

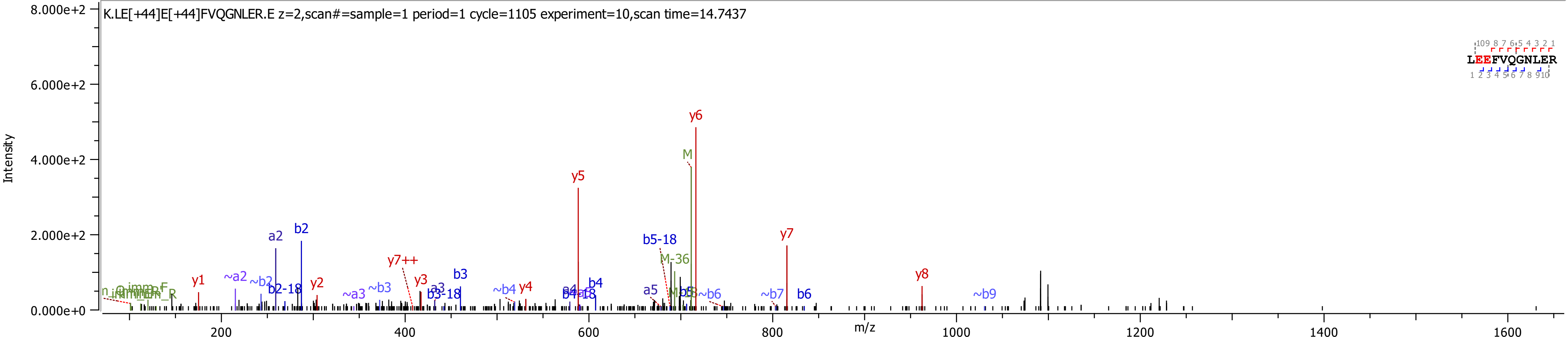
K.LE[+44]E[+44]FVQGNLER.E z=2,scan#=sample=1 period=1 cycle=1105 experiment=10,scan time=14.7437
Intensity
m/z
8.000e+2
6.000e+2
4.000e+2
2.000e+2
0.000e+0
200
400
600
800
1000
1200
1400
1600
LEEFVQGNLER
y1
~a2
~b2
b2-18
a2
b2
y2
~a3
~b3
y7++
y3
a3
b3-18
b3
~b4
y4
a4
b4-18
b4
y5
a5
b5-18
M-36
b5
~b6
M
y6
~b7
y7
b6
y8
~b9

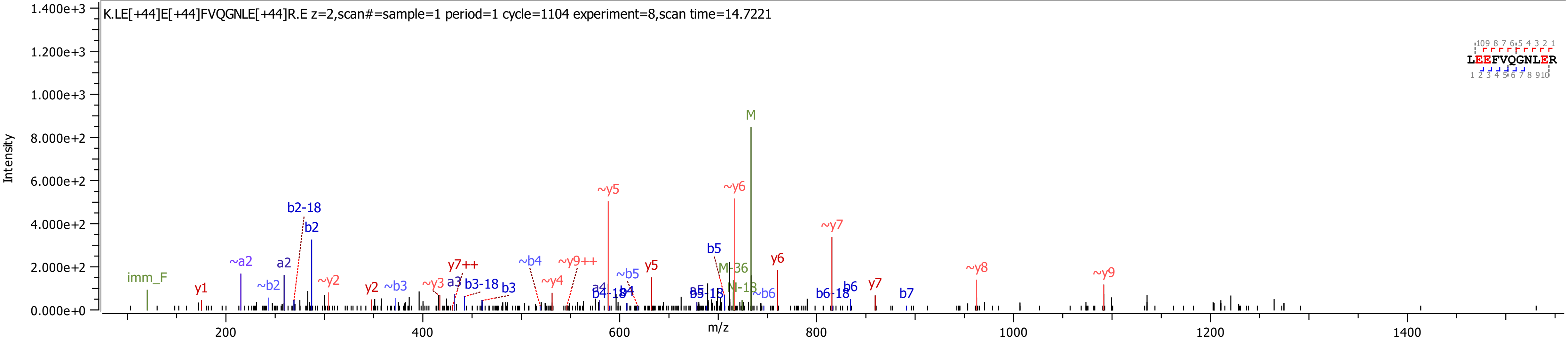
K.LE[+44]E[+44]FVQGNLE[+44]R.E z=2,scan#=sample=1 period=1 cycle=1104 experiment=8,scan time=14.7221
Intensity
m/z
LEEFVQGNLER
M
~y5
~y6
~y7
~y8
~y9
b2-18
b2
a2
~a2
~b2
imm_F
y1
~y2
y2
~b3
~y3
a3
y7++
b3-18
b3
~b4
~y4
~y9++
a4
b4-18
b4
~b5
y5
b5
b5-18
M-36
M-18
~b6
y6
b6-18
b6
y7
b7

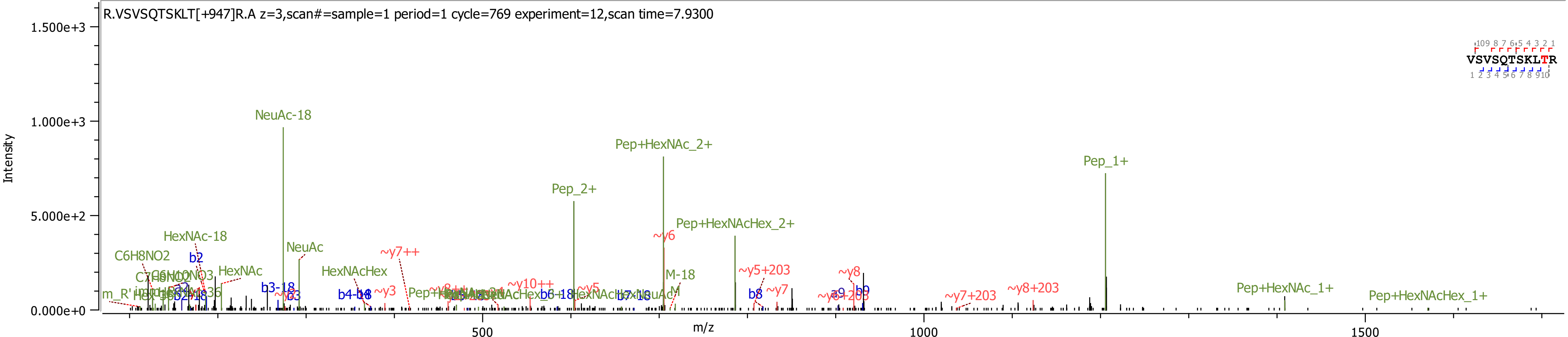
R.VSVSQTSKLT[+947]R.A z=3,scan#=sample=1 period=1 cycle=769 experiment=12,scan time=7.9300
Intensity
1.500e+3
1.000e+3
5.000e+2
0.000e+0
m/z
500
1000
1500
VSVSQTSKLTR
NeuAc-18
Pep+HexNAc_2+
Pep_2+
Pep_1+
Pep+HexNAcHex_2+
~y6
HexNAc-18
NeuAc
C6H8NO2
b2
HexNAc
HexNAcHex
~y7++
M-18
~y5+203
~y8
~y10++
~y5
~y3
b3-18
b4-18
b6-18
b7-18
b8
~y7
a9
b9
~y7+203
~y8+203
Pep+HexNAc_1+
Pep+HexNAcHex_1+

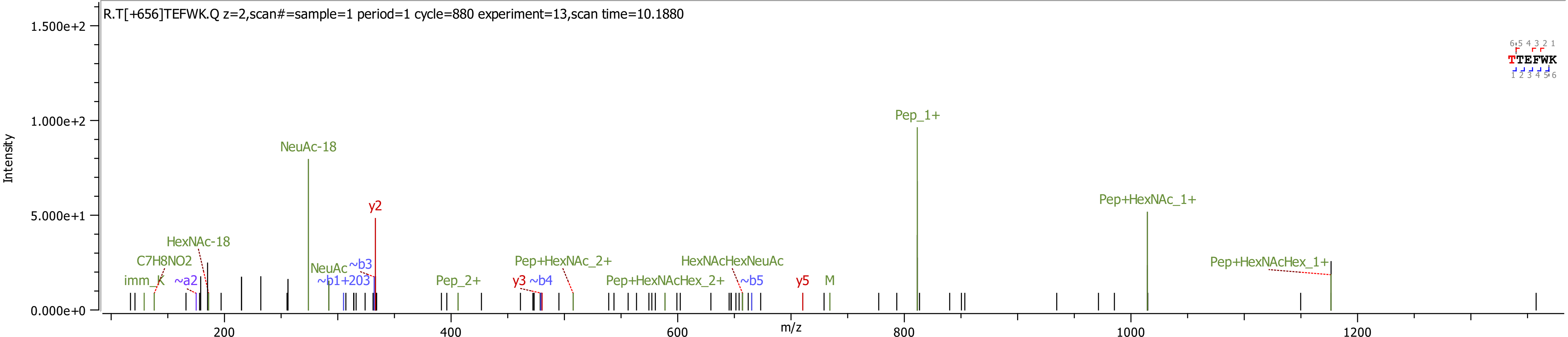

R.T[+656]TEFWK.Q z=2,scan#=sample=1 period=1 cycle=880 experiment=13,scan time=10.1880
Intensity
1.500e+2
1.000e+2
5.000e+1
0.000e+0
m/z
200
400
600
800
1000
1200
TTEFWK
NeuAc-18
y2
Pep_1+
Pep+HexNAc_1+
HexNAc-18
C7H8NO2
imm_K
~a2
NeuAc
~b3
~b1+203
Pep_2+
y3
~b4
Pep+HexNAc_2+
Pep+HexNAcHex_2+
HexNAcHexNeuAc
~b5
y5
M
Pep+HexNAcHex_1+

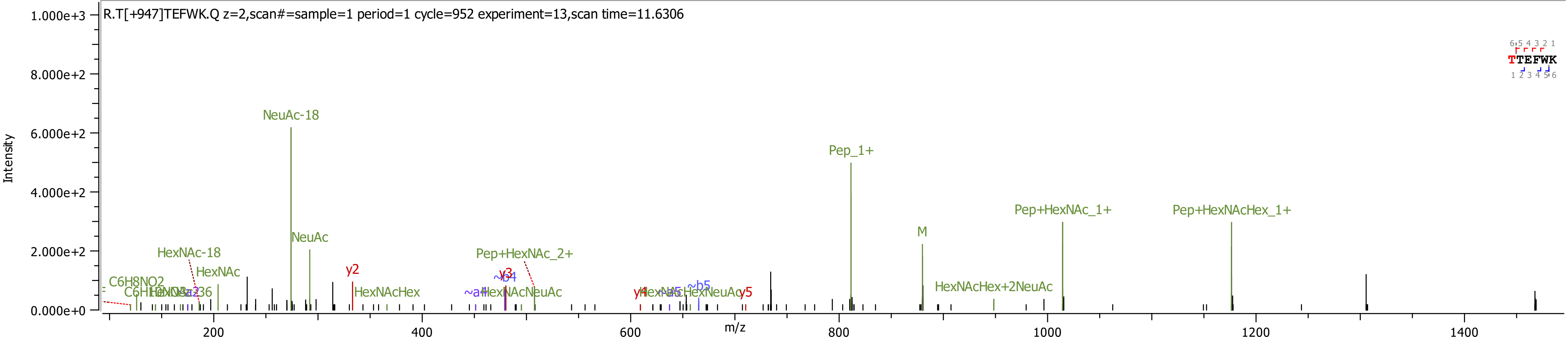
R.T[+947]TEFWK.Q z=2,scan#=sample=1 period=1 cycle=952 experiment=13,scan time=11.6306
Intensity
m/z
NeuAc-18
Pep_1+
Pep+HexNAc_1+
Pep+HexNAcHex_1+
M
NeuAc
HexNAc-18
HexNAc
Pep+HexNAc_2+
y2
y3
~b4
~a4
HexNAcHex
HexNAcNeuAc
HexNAcHex+2NeuAc
C6H8NO2
y4
~a5
~b5
HexNAcHexNeuAc
y5
TTEFWK

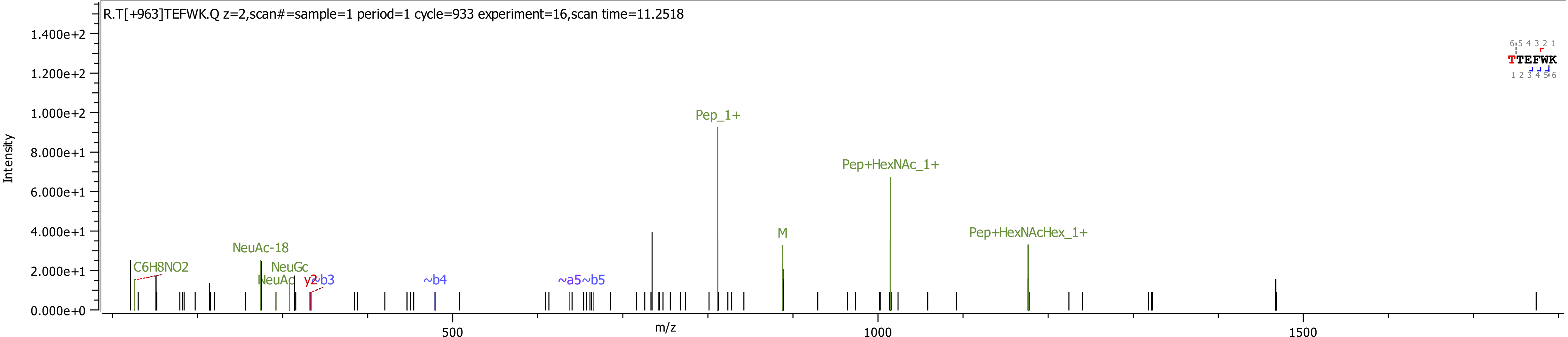
R.T[+963]TEFWK.Q z=2,scan#=sample=1 period=1 cycle=933 experiment=16,scan time=11.2518
Intensity
m/z
TTEFWK
C6H8NO2
NeuAc-18
NeuGc
NeuAc
y2
~b3
~b4
~a5
~b5
Pep_1+
M
Pep+HexNAc_1+
Pep+HexNAcHex_1+

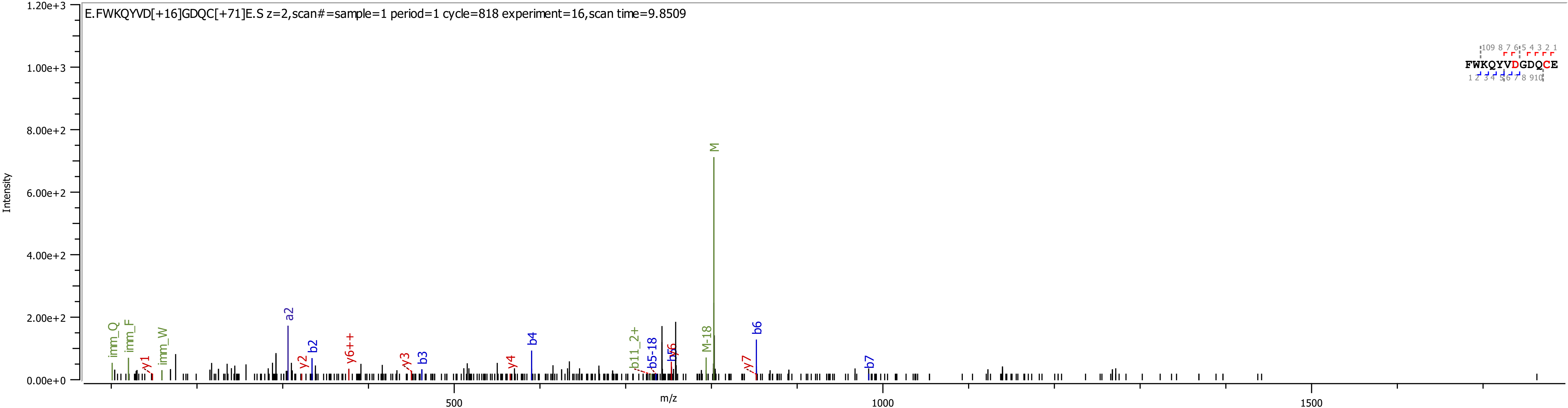

E.FWKQYVD[+16]GDQC[+71]E.S z=2,scan#=sample=1 period=1 cycle=818 experiment=16,scan time=9.8509
Intensity
m/z
0.00e+0
2.00e+2
4.00e+2
6.00e+2
8.00e+2
1.00e+3
1.20e+3
500
1000
1500
FWKQYVDGDQCE
M
M-18
imm_Q
imm_F
imm_W
y1
a2
y2
b2
y6++
y3
b3
y4
b4
b11_2+
b5-18
b5
y5
y7
b6
b7

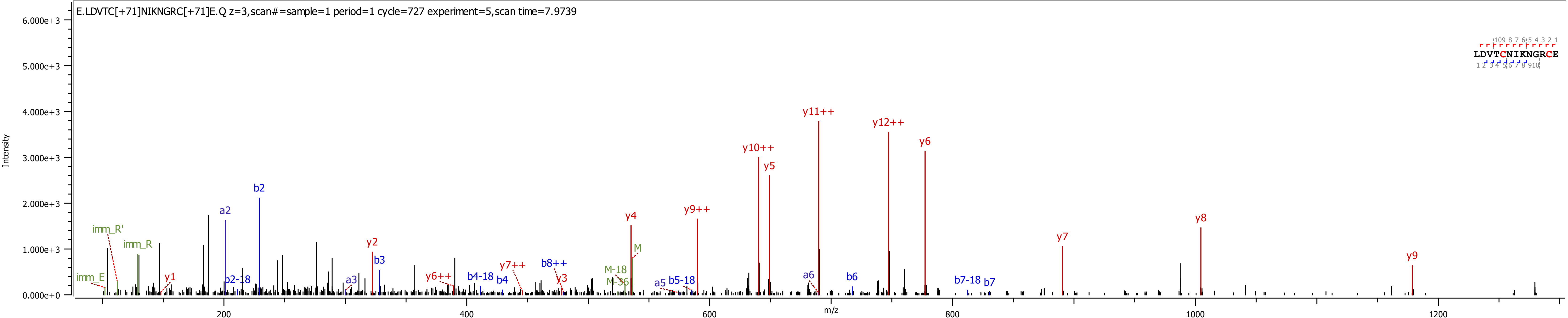
E.LDVTC[+71]NIKNGRC[+71]E.Q z=3,scan#=sample=1 period=1 cycle=727 experiment=5,scan time=7.9739
Intensity
m/z
LDVTCNIKNGRCE
y11++
y12++
y10++
y6
y5
y9++
y4
y8
y7
y9
y2
y1
y6++
y7++
y3
b2
a2
b3
a3
b2-18
b4-18
b4
b8++
a5
b5-18
a6
b6
b7-18
b7
imm_R'
imm_R
imm_E
M
M-18
M-36

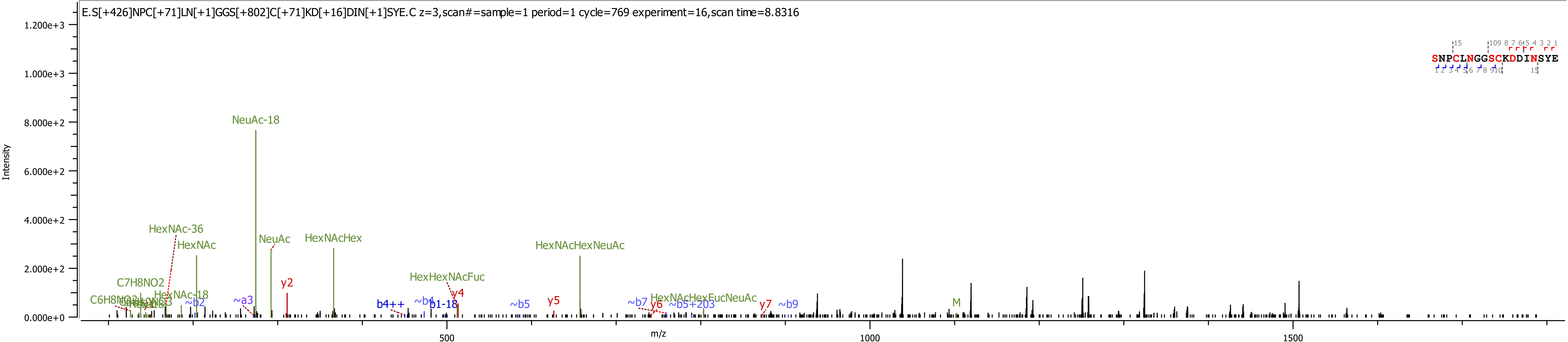
E.S[+426]NPC[+71]LN[+1]GGS[+802]C[+71]KD[+16]DIN[+1]SYE.C z=3,scan#=sample=1 period=1 cycle=769 experiment=16,scan time=8.8316
SNPCLNGGSCKDDINSYE
Intensity
m/z
NeuAc-18
HexNAc-36
HexNAc
NeuAc
HexNAcHex
HexNAcHexNeuAc
HexHexNAcFuc
C7H8NO2
C6H8NO2
HexNAc-18
~b2
~a3
y2
b4++
~b4
b1-18
y4
~b5
y5
~b7
y6
HexNAcHexFucNeuAc
~b5+203
y7
~b9
M
500
1000
1500

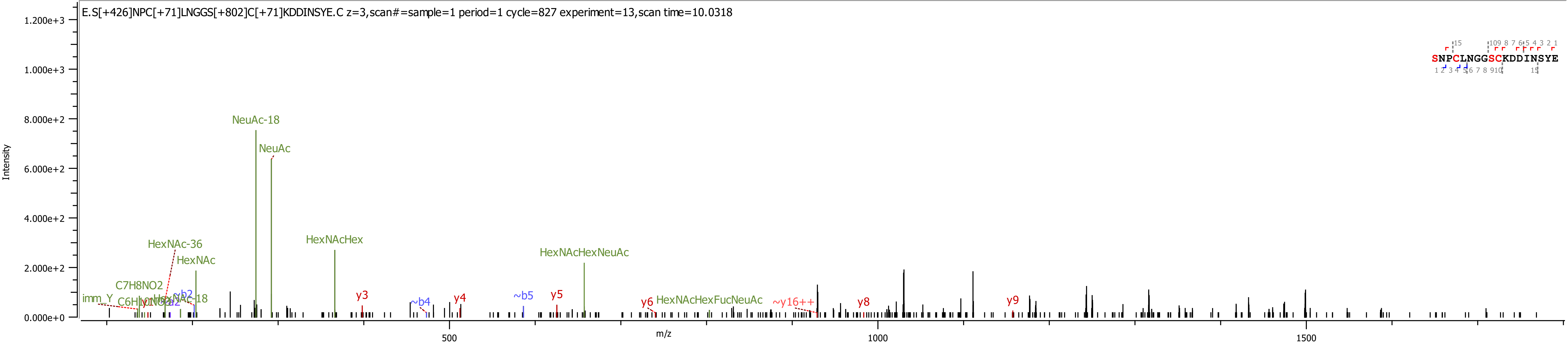
E.S[+426]NPC[+71]LNGGS[+802]C[+71]KDDINSYE.C z=3,scan#=sample=1 period=1 cycle=827 experiment=13,scan time=10.0318
SNPCLNGGSCKDDINSYE
Intensity
1.200e+3
1.000e+3
8.000e+2
6.000e+2
4.000e+2
2.000e+2
0.000e+0
m/z
500
1000
1500
NeuAc-18
NeuAc
HexNAc-36
HexNAc
HexNAcHex
HexNAcHexNeuAc
C7H8NO2
imm_Y
~b2
~b4
~b5
y3
y4
y5
y6
y8
y9
HexNAcHexFucNeuAc
~y16++

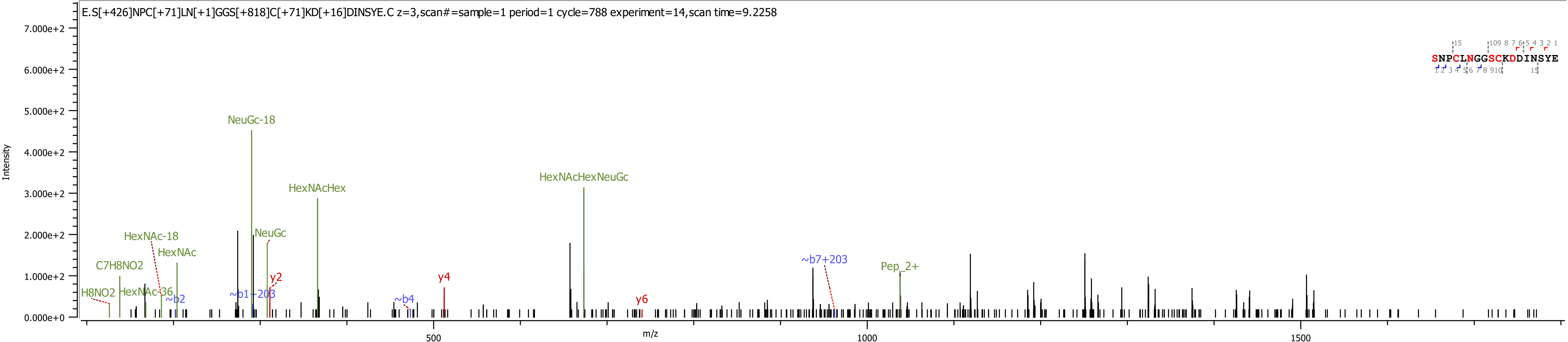
E.S[+426]NPC[+71]LN[+1]GGS[+818]C[+71]KD[+16]DINSYE.C z=3,scan#=sample=1 period=1 cycle=788 experiment=14,scan time=9.2258
Intensity
m/z
NeuGc-18
HexNAcHexNeuGc
HexNAcHex
HexNAc-18
NeuGc
HexNAc
C7H8NO2
y2
y4
~b7+203
Pep_2+
H8NO2
HexNAc-36
~b2
~b1+203
~b4
y6
SNPCLNGGSCKDDINSYE

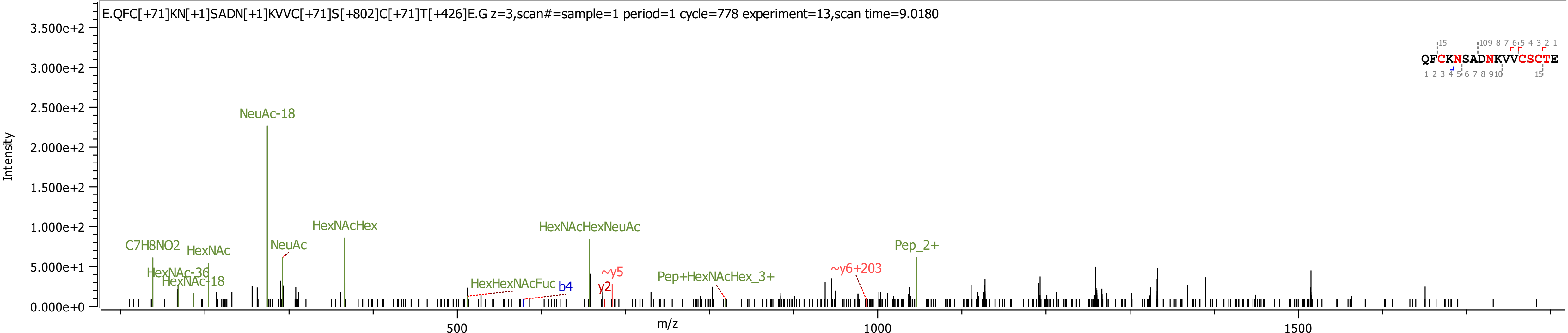
E.QFC[+71]KN[+1]SADN[+1]KVVC[+71]S[+802]C[+71]T[+426]E.G z=3,scan#=sample=1 period=1 cycle=778 experiment=13,scan time=9.0180
QFCKNSADNKVVCSCTE
Intensity
m/z
3.500e+2
3.000e+2
2.500e+2
2.000e+2
1.500e+2
1.000e+2
5.000e+1
0.000e+0
500
1000
1500
NeuAc-18
HexNAcHex
HexNAcHexNeuAc
C7H8NO2
HexNAc
NeuAc
Pep_2+
HexNAc-36
HexNAc-18
HexHexNAcFuc
b4
~y5
y2
Pep+HexNAcHex_3+
~y6+203

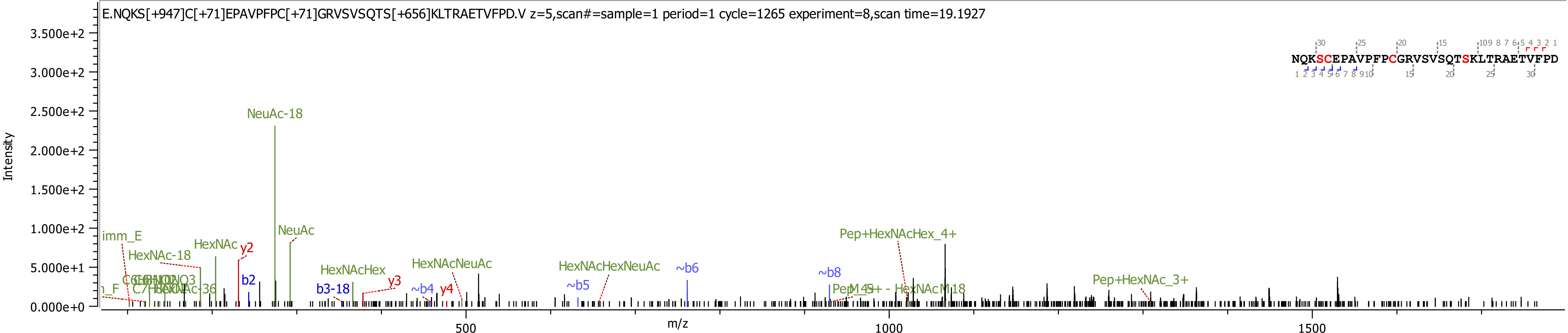

E.NQKS[+947]C[+71]EPAVPFPC[+71]GRVSVSQTS[+656]KLTRAETVFPD.V z=5,scan#=sample=1 period=1 cycle=1265 experiment=8,scan time=19.1927
NQKSCEPAVPFPCGRVSVSQTSKLTRAETVFPD
Intensity
3.500e+2
3.000e+2
2.500e+2
2.000e+2
1.500e+2
1.000e+2
5.000e+1
0.000e+0
m/z
500
1000
1500
NeuAc-18
NeuAc
imm_E
HexNAc-18
HexNAc
y2
b2
b3-18
y3
HexNAcHex
HexNAcNeuAc
~b4
y4
HexNAcHexNeuAc
~b5
~b6
~b8
Pep+HexNAcHex_4+
Pep+HexNAc_3+

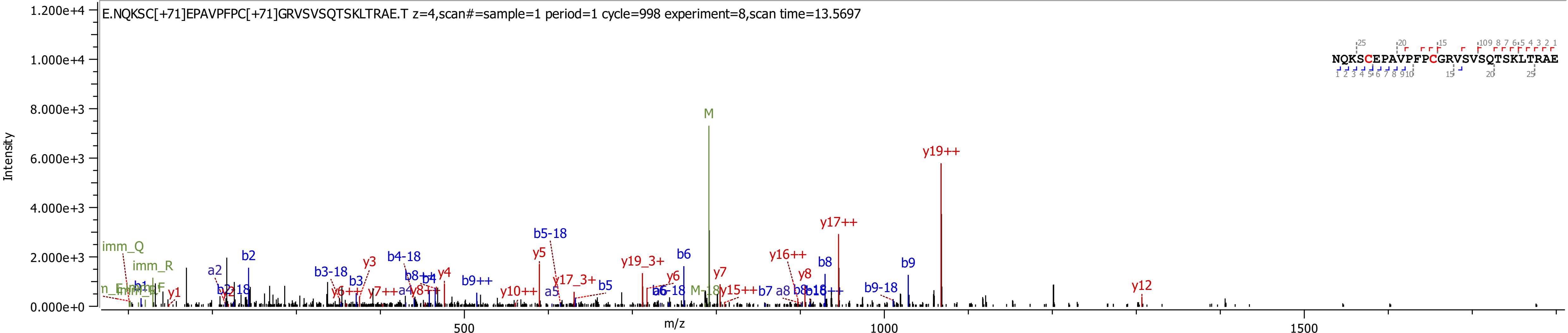
E.NQKSC[+71]EPAVPFPC[+71]GRVSVSQTSKLTRAE.T z=4,scan#=sample=1 period=1 cycle=998 experiment=8,scan time=13.5697
NQKSCEPAVPFPCGRVSVSQTSKLTRAE
Intensity
m/z
1.200e+4
1.000e+4
8.000e+3
6.000e+3
4.000e+3
2.000e+3
0.000e+0
500
1000
1500
M
y19++
y17++
b5-18
imm_Q
imm_R
a2
b2
b3-18
y3
b4-18
y4
b9++
y5
y19_3+
b6
y6
y7
y16++
y8
b8
b9
y17_3+
b5
y10++
a5
b7
a8
b9-18
y12
M-18
y15++
y1
b3
y7++
a4

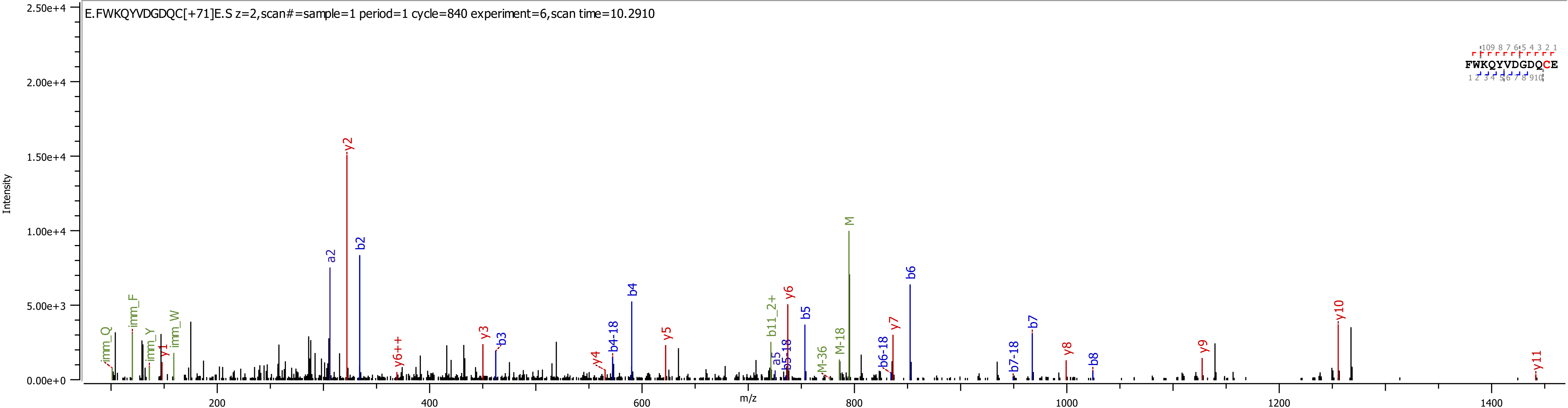
E.FWKQYVDGDQC[+71]E.S z=2,scan#=sample=1 period=1 cycle=840 experiment=6,scan time=10.2910
FWKQYVDGDQCE
Intensity
m/z
2.50e+4
2.00e+4
1.50e+4
1.00e+4
5.00e+3
0.00e+0
200
400
600
800
1000
1200
1400
imm_Q
imm_F
imm_Y
y1
imm_W
a2
y2
b2
y6++
y3
b3
y4
b4-18
b4
y5
b11_2+
a5
b5-18
y6
b5
M-36
M-18
M
b6-18
y7
b6
b7-18
b7
y8
b8
y9
y10
y11

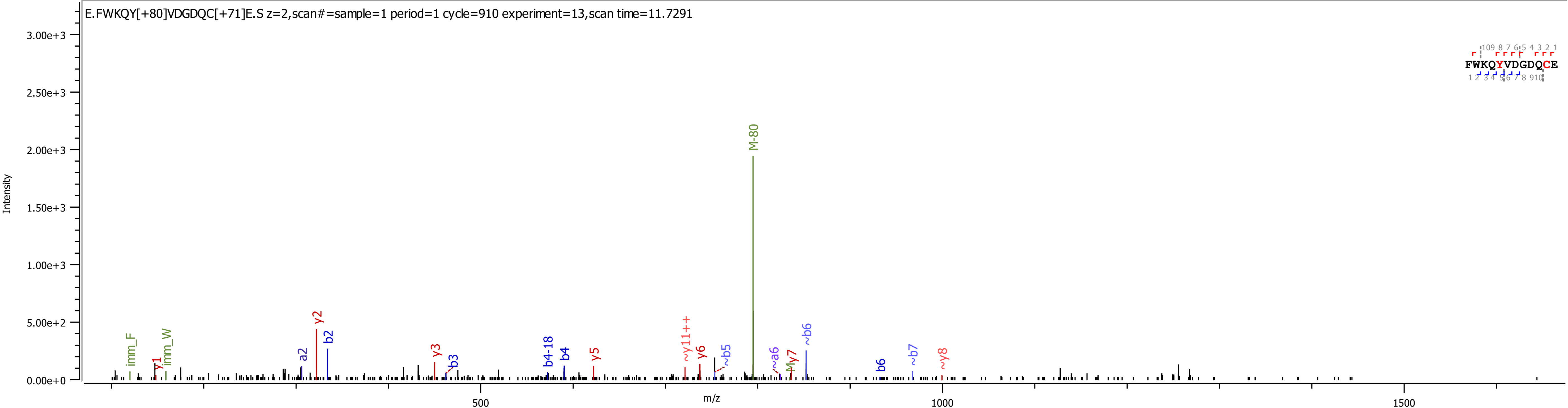

E.FWKQY[+80]VDGDQC[+71]E.S z=2,scan#=sample=1 period=1 cycle=910 experiment=13,scan time=11.7291
Intensity
0.00e+0
5.00e+2
1.00e+3
1.50e+3
2.00e+3
2.50e+3
3.00e+3
m/z
500
1000
1500
M-80
imm_F
y1
imm_W
a2
y2
b2
y3
~b3
b4-18
b4
y5
~y11++
y6
~b5
~a6
~M
y7
~b6
b6
~b7
~y8
FWKQYVDGDQCE

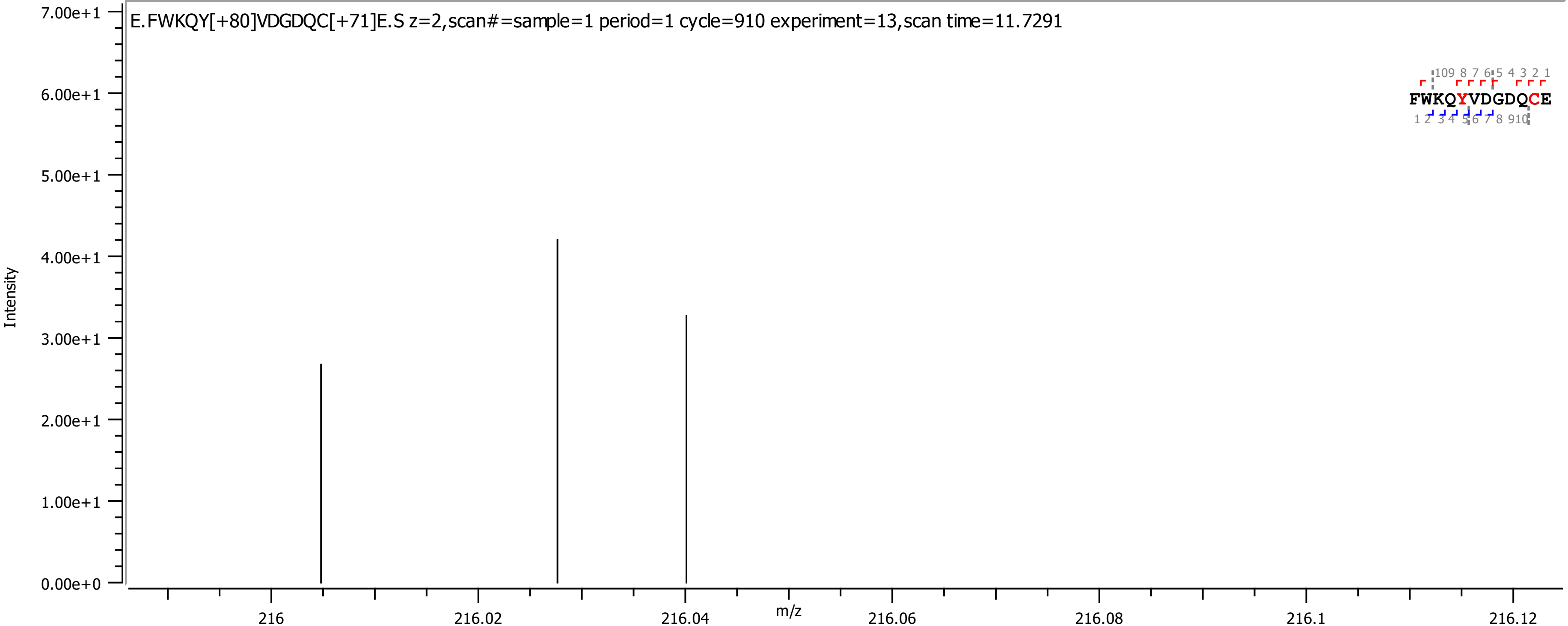
E.FWKQY[+80]VDGDQC[+71]E.S z=2,scan#=sample=1 period=1 cycle=910 experiment=13,scan time=11.7291
FWKQYVDGDQCE
Intensity
m/z
7.00e+1
6.00e+1
5.00e+1
4.00e+1
3.00e+1
2.00e+1
1.00e+1
0.00e+0
216
216.02
216.04
216.06
216.08
216.1
216.12

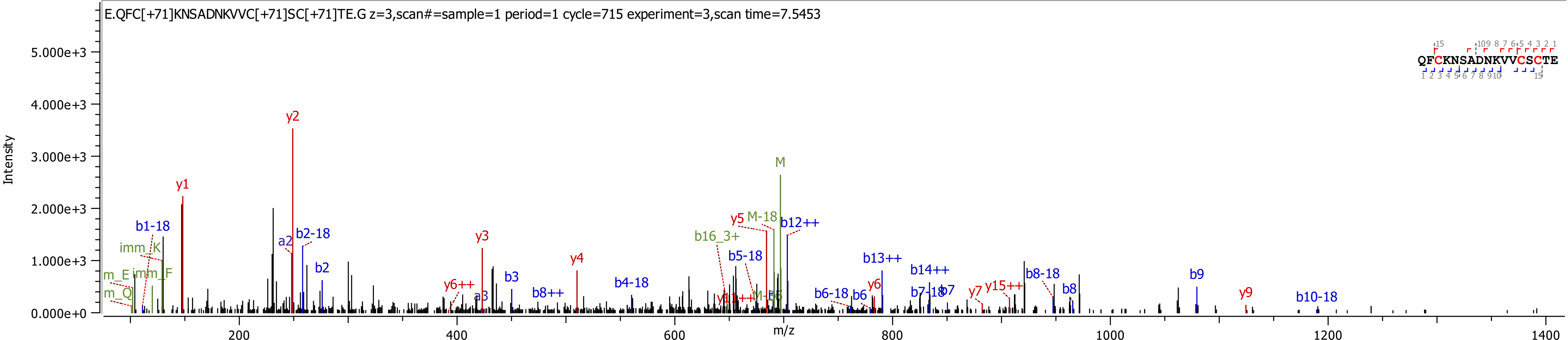
E.QFC[+71]KNSADNKVVC[+71]SC[+71]TE.G z=3,scan#=sample=1 period=1 cycle=715 experiment=3,scan time=7.5453
Intensity
m/z
QFCKNSADNKVVCSCTE
y2
M
y1
b1-18
imm_K
m_E
imm_F
m_Q
a2
b2-18
b2
y3
y4
y6++
a3
b3
b8++
b4-18
y5
M-18
b12++
b16_3+
b5-18
b13++
b14++
b6-18
b6
y6
b7-18
b7
y7
y15++
b8-18
b8
b9
y9
b10-18

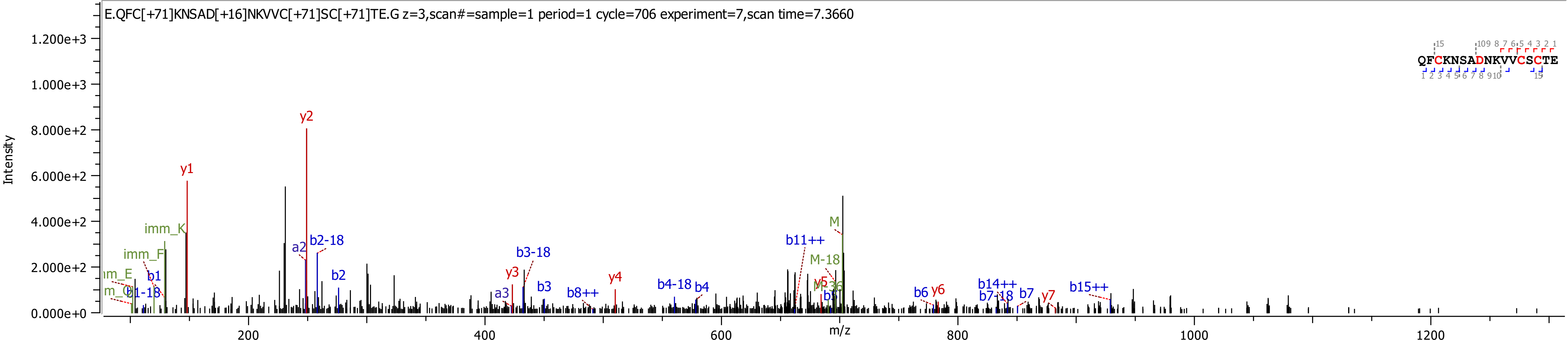

E.QFC[+71]KNSAD[+16]NKVVC[+71]SC[+71]TE.G z=3,scan#=sample=1 period=1 cycle=706 experiment=7,scan time=7.3660
Intensity
m/z
0.000e+0
2.000e+2
4.000e+2
6.000e+2
8.000e+2
1.000e+3
1.200e+3
200
400
600
800
1000
1200
QFCKNSADNKVVCSCTE
y1
y2
y3
y4
y5
y6
y7
a2
a3
b1
b1-18
b2
b2-18
b3
b3-18
b4
b4-18
b6
b7
b7-18
b8++
b11++
b14++
b15++
imm_K
imm_F
M
M-18

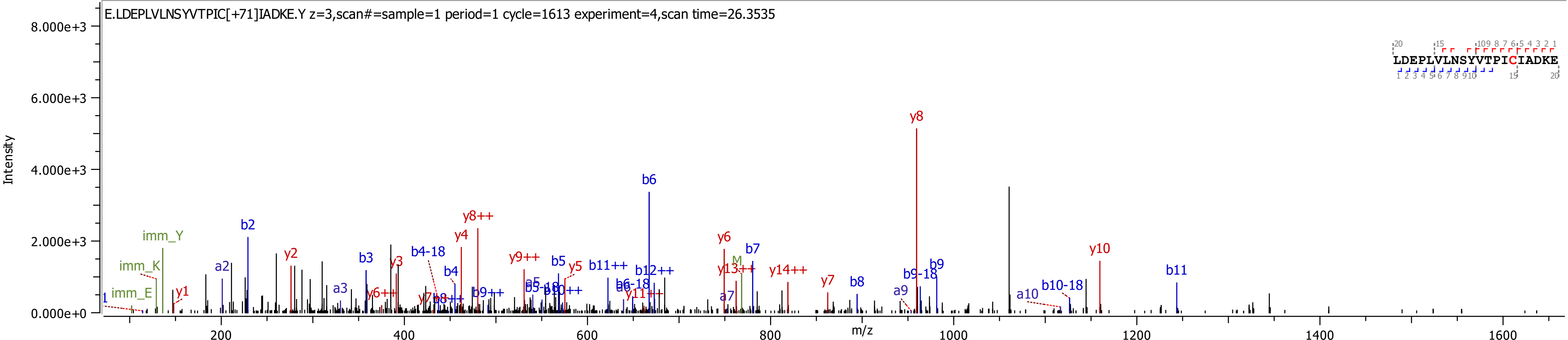

E.LDEPLVLNSYVTPIC[+71]IADKE.Y z=3,scan#=sample=1 period=1 cycle=1613 experiment=4,scan time=26.3535
LDEPLVLNSYVTPICIADKE
Intensity
m/z
8.000e+3
6.000e+3
4.000e+3
2.000e+3
0.000e+0
200
400
600
800
1000
1200
1400
1600

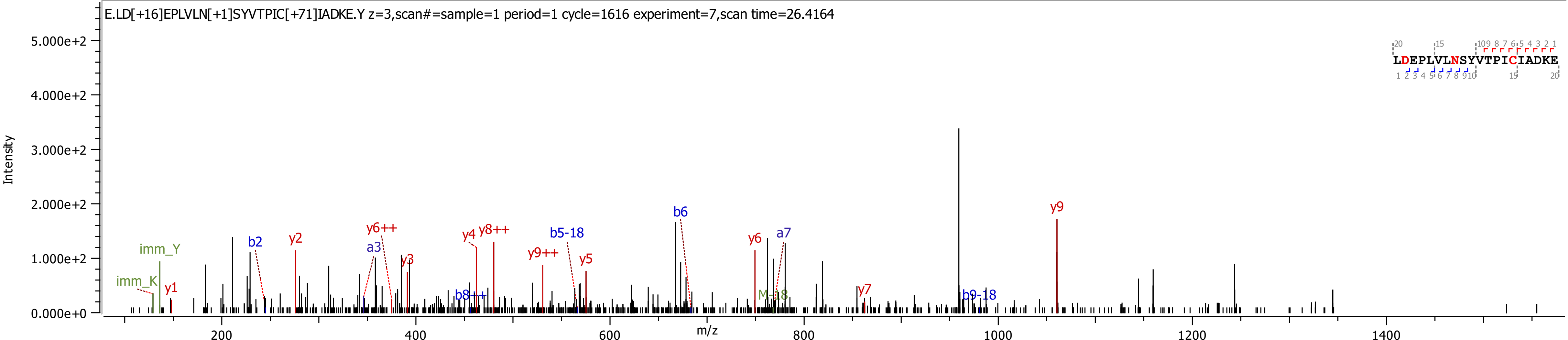
E.LD[+16]EPLVLN[+1]SYVTPIC[+71]IADKE.Y z=3,scan#=sample=1 period=1 cycle=1616 experiment=7,scan time=26.4164
Intensity
m/z
LDEPLVLNSYVTPICIADKE
imm_Y
imm_K
y1
b2
y2
a3
y6++
y3
b8++
y4
y8++
y9++
b5-18
y5
b6
y6
a7
M-18
y7
b9-18
y9

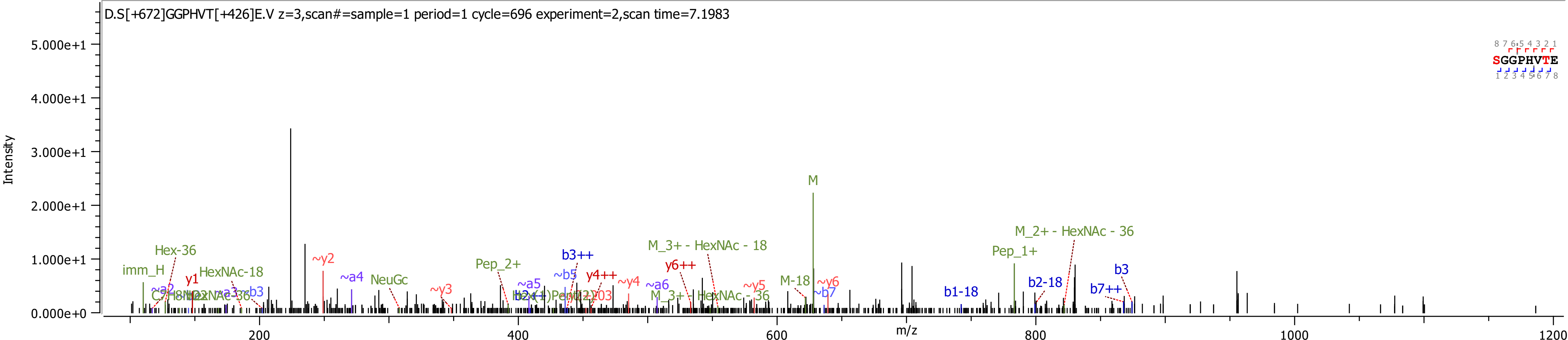
D.S[+672]GGPHVT[+426]E.V z=3,scan#=sample=1 period=1 cycle=696 experiment=2,scan time=7.1983
Intensity
5.000e+1
4.000e+1
3.000e+1
2.000e+1
1.000e+1
0.000e+0
m/z
200
400
600
800
1000
1200
SGGPHVTE
M
M_2+ - HexNAc - 36
Pep_1+
M_3+ - HexNAc - 18
M_3+ - HexNAc - 36
Hex-36
imm_H
HexNAc-18
y1
~y2
~a4
NeuGc
Pep_2+
b3++
~b5
~a5
y4++
~y4
~a6
y6++
~y3
~y5
~y6
M-18
~b7
b1-18
b2-18
b3
b7++

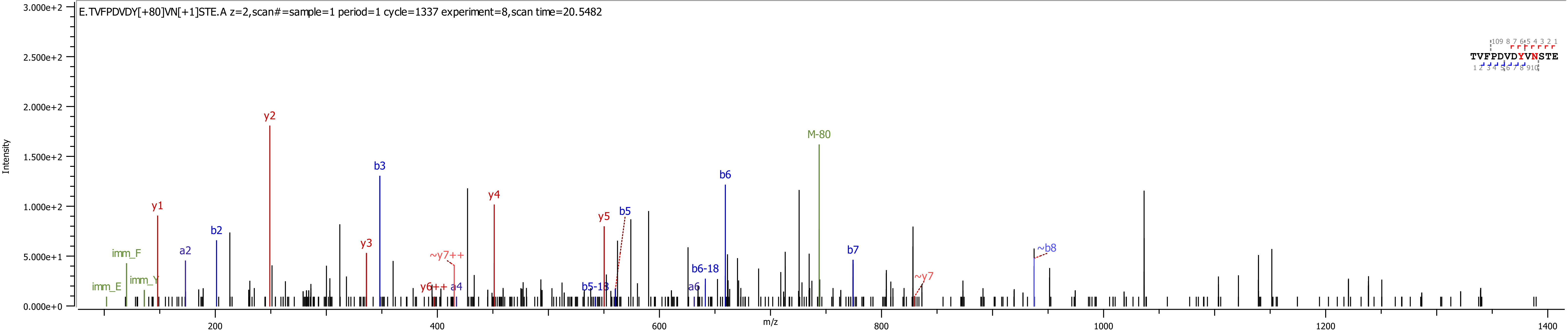
E.TVFPDVDY[+80]VN[+1]STE.A z=2,scan#=sample=1 period=1 cycle=1337 experiment=8,scan time=20.5482
Intensity
m/z
TVFPDVDYVNSTE
y1
y2
y3
y4
y5
b2
b3
b5
b6
b7
a2
a4
a6
imm_F
imm_E
imm_Y
M-80
~y7++
y6++
~y7
~b8
b5-18
b6-18
3.000e+2
2.500e+2
2.000e+2
1.500e+2
1.000e+2
5.000e+1
0.000e+0
200
400
600
800
1000
1200
1400

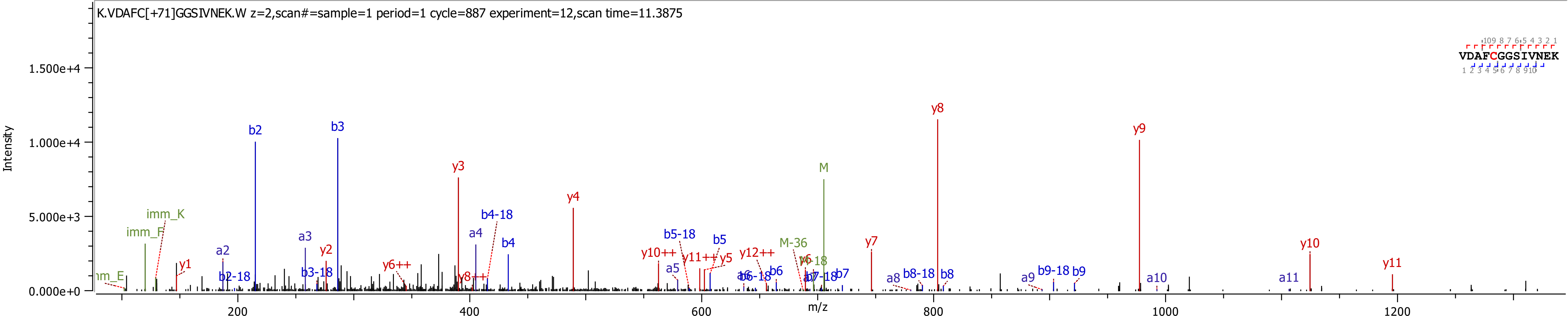
K.VDAFC[+71]GGSIVNEK.W z=2,scan#=sample=1 period=1 cycle=887 experiment=12,scan time=11.3875
VDAFCGGSIVNEK
Intensity
1.500e+4
1.000e+4
5.000e+3
0.000e+0
m/z
200
400
600
800
1000
1200
imm_K
imm_F
y1
a2
b2
b2-18
a3
b3
b3-18
y2
y3
y6++
y8++
a4
b4
b4-18
y4
y10++
a5
b5-18
y11++
y5
b5
y12++
b6-18
b6
M-36
y6
M
b7-18
b7
y7
a8
b8-18
b8
y8
a9
b9-18
b9
y9
a10
a11
y10
y11

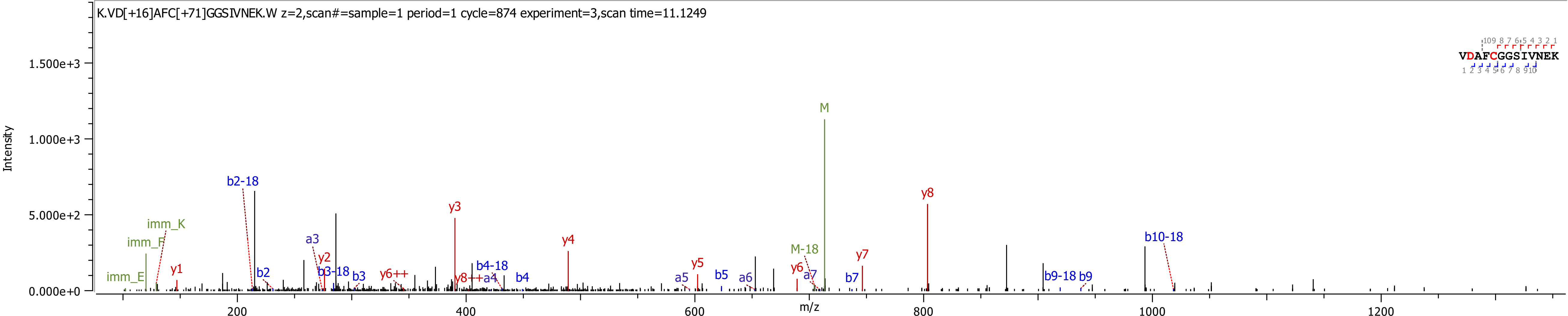
K.VD[+16]AFC[+71]GGSIVNEK.W z=2,scan#=sample=1 period=1 cycle=874 experiment=3,scan time=11.1249
Intensity
m/z
VDAFCGGSIVNEK
imm_E
imm_F
imm_K
y1
b2-18
b2
a3
y2
b3-18
b3
y6++
y3
y8++
a4
b4-18
b4
y4
a5
y5
b5
a6
y6
M-18
a7
M
b7
y7
y8
b9-18
b9
b10-18

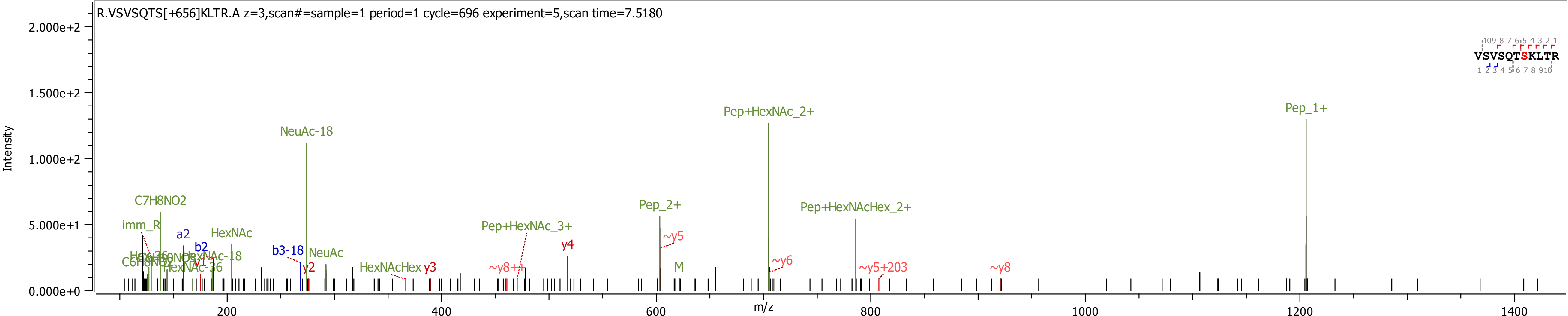
R.VSVSQTS[+656]KLTR.A z=3,scan#=sample=1 period=1 cycle=696 experiment=5,scan time=7.5180
Intensity
m/z
NeuAc-18
Pep+HexNAc_2+
Pep_1+
C7H8NO2
imm_R
a2
HexNAc
b2
b3-18
NeuAc
y2
HexNAcHex
y3
~y8++
Pep+HexNAc_3+
y4
Pep_2+
~y5
M
~y6
Pep+HexNAcHex_2+
~y5+203
~y8
VSVSQTSKLTR

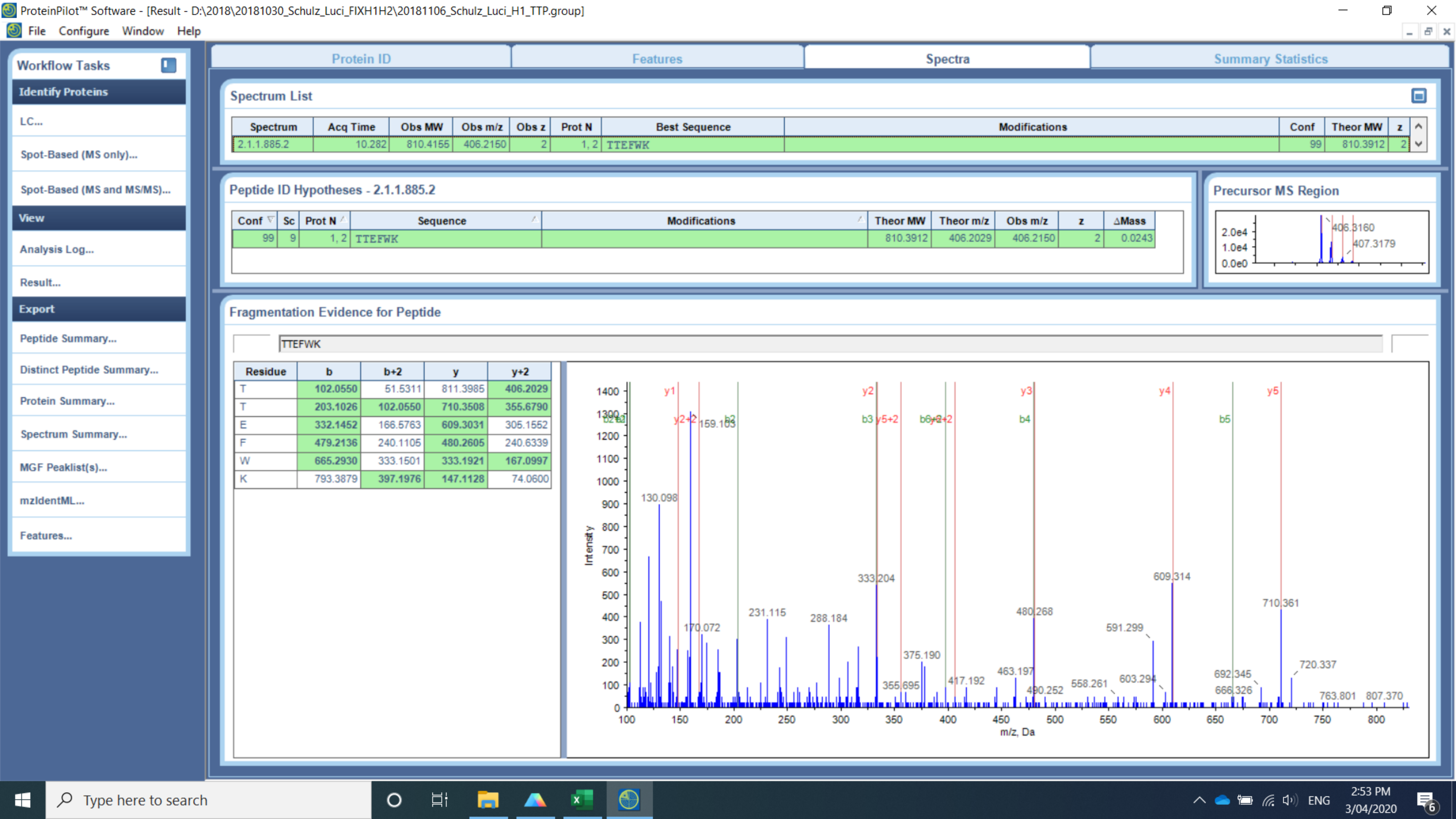
ProteinPilot™ Software - [Result - D:\2018\20181030_Schulz_Luci_FIXH1H2\20181106_Schulz_Luci_H1_TTP.group]
File Configure Window Help
Workflow Tasks
Identify Proteins
LC...
Spot-Based (MS only)...
Spot-Based (MS and MS/MS)...
View
Analysis Log...
Result...
Export
Peptide Summary...
Distinct Peptide Summary...
Protein Summary...
Spectrum Summary...
MGF Peaklist(s)...
mzIdentML...
Features...
Protein ID
Features
Spectra
Summary Statistics
Spectrum List
Spectrum | Acq Time | Obs MW | Obs m/z | Obs z | Prot N | Best Sequence | Modifications | Conf | Theor MW | z
2.1.1.885.2 | 10.282 | 810.4155 | 406.2150 | 2 | 1, 2 | TTEFWK | | 99 | 810.3912 | 2
Peptide ID Hypotheses - 2.1.1.885.2
Conf | Sc | Prot N | Sequence | Modifications | Theor MW | Theor m/z | Obs m/z | z | ΔMass
99 | 9 | 1, 2 | TTEFWK | | 810.3912 | 406.2029 | 406.2150 | 2 | 0.0243
Precursor MS Region
2.0e4
1.0e4
0.0e0
406.3160
407.3179
Fragmentation Evidence for Peptide
TTEFWK
Residue | b | b+2 | y | y+2
T | 102.0550 | 51.5311 | 811.3985 | 406.2029
T | 203.1026 | 102.0550 | 710.3508 | 355.6790
E | 332.1452 | 166.5763 | 609.3031 | 305.1552
F | 479.2136 | 240.1105 | 480.2605 | 240.6339
W | 665.2930 | 333.1501 | 333.1921 | 167.0997
K | 793.3879 | 397.1976 | 147.1128 | 74.0600
y1
y2
y3
y4
y5
y2+2
b2
b3
y5+2
b4
b5
159.103
130.098
170.072
231.115
288.184
333.204
355.695
375.190
417.192
463.197
480.268
490.252
558.261
591.299
603.294
609.314
666.326
692.345
710.361
720.337
763.801
807.370
Intensity
m/z, Da
Type here to search
ENG
2:53 PM
3/04/2020

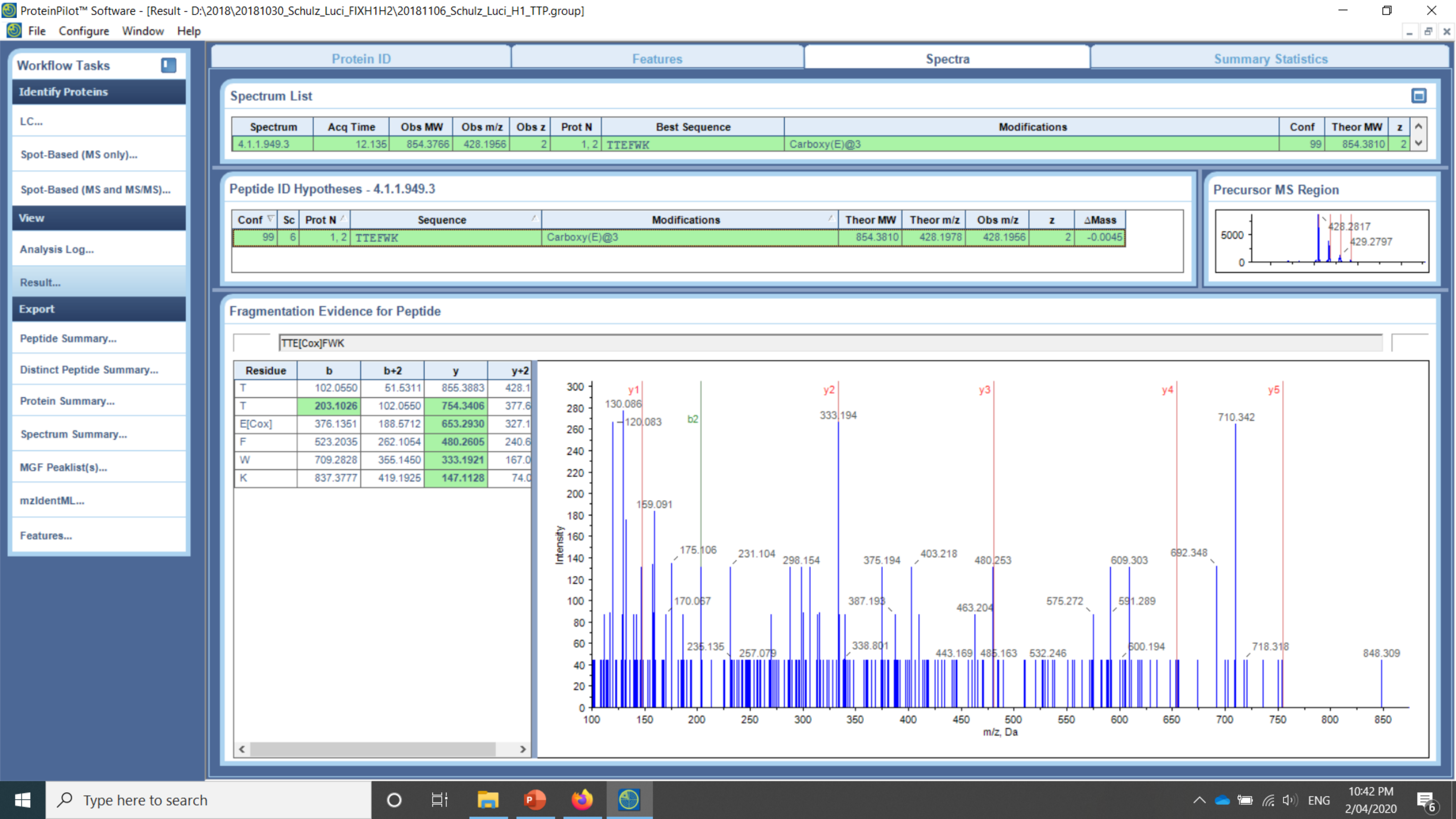
ProteinPilot™ Software - [Result - D:\2018\20181030_Schulz_Luci_FIXH1H2\20181106_Schulz_Luci_H1_TTP.group]
File Configure Window Help
Workflow Tasks
Identify Proteins
LC...
Spot-Based (MS only)...
Spot-Based (MS and MS/MS)...
View
Analysis Log...
Result...
Export
Peptide Summary...
Distinct Peptide Summary...
Protein Summary...
Spectrum Summary...
MGF Peaklist(s)...
mzIdentML...
Features...
Protein ID
Features
Spectra
Summary Statistics
Spectrum List
Spectrum | Acq Time | Obs MW | Obs m/z | Obs z | Prot N | Best Sequence | Modifications | Conf | Theor MW | z
4.1.1.949.3 | 12.135 | 854.3766 | 428.1956 | 2 | 1, 2 | TTEFWK | Carboxy(E)@3 | 99 | 854.3810 | 2
Peptide ID Hypotheses - 4.1.1.949.3
Conf | Sc | Prot N | Sequence | Modifications | Theor MW | Theor m/z | Obs m/z | z | ΔMass
99 | 6 | 1, 2 | TTEFWK | Carboxy(E)@3 | 854.3810 | 428.1978 | 428.1956 | 2 | -0.0045
Precursor MS Region
5000
0
428.2817
429.2797
Fragmentation Evidence for Peptide
TTE[Cox]FWK
Residue | b | b+2 | y | y+2
T | 102.0550 | 51.5311 | 855.3883 | 428.1
T | 203.1026 | 102.0550 | 754.3406 | 377.6
E[Cox] | 376.1351 | 188.5712 | 653.2930 | 327.1
F | 523.2035 | 262.1054 | 480.2605 | 240.6
W | 709.2828 | 355.1450 | 333.1921 | 167.0
K | 837.3777 | 419.1925 | 147.1128 | 74.0
y1
y2
y3
y4
y5
b2
130.086
120.083
159.091
175.106
170.067
231.104
236.135
257.079
298.154
333.194
338.801
375.194
387.193
403.218
443.169
463.204
480.253
486.163
532.246
575.272
591.289
600.194
609.303
692.348
710.342
718.318
848.309
Intensity
m/z, Da
Type here to search
ENG
10:42 PM
2/04/2020

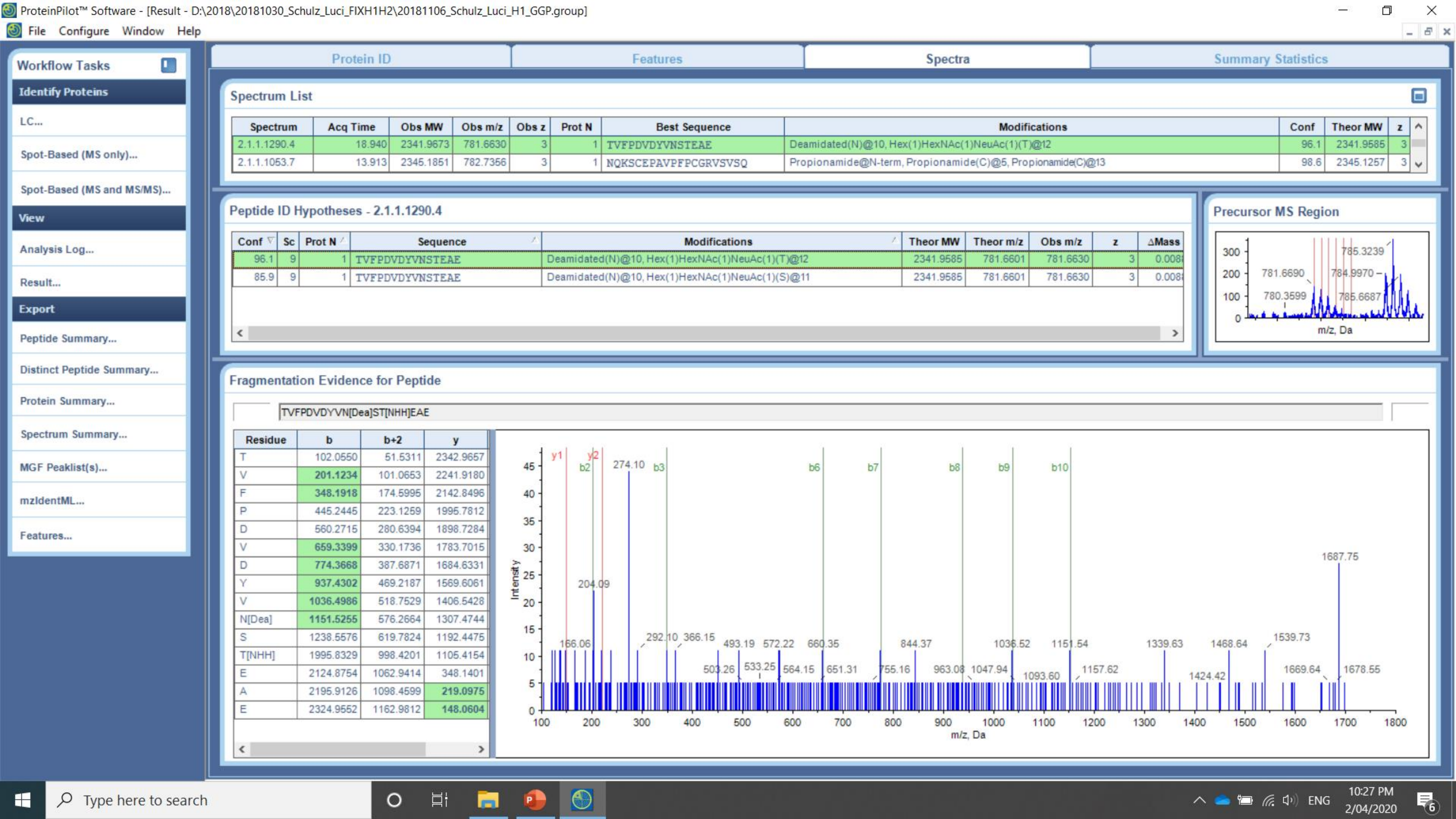
ProteinPilot™ Software - [Result - D:\2018\20181030_Schulz_Luci_FIXH1H2\20181106_Schulz_Luci_H1_GGP.group]
File Configure Window Help
Workflow Tasks
Identify Proteins
LC...
Spot-Based (MS only)...
Spot-Based (MS and MS/MS)...
View
Analysis Log...
Result...
Export
Peptide Summary...
Distinct Peptide Summary...
Protein Summary...
Spectrum Summary...
MGF Peaklist(s)...
mzIdentML...
Features...
Protein ID
Features
Spectra
Summary Statistics
Spectrum List
Spectrum Acq Time Obs MW Obs m/z Obs z Prot N Best Sequence Modifications Conf Theor MW z
2.1.1.1290.4 18.940 2341.9673 781.6630 3 1 TVFPDVDYVNSTEAE Deamidated(N)@10, Hex(1)HexNAc(1)NeuAc(1)(T)@12 96.1 2341.9585 3
2.1.1.1053.7 13.913 2345.1851 782.7356 3 1 NQKSCEPAVPFPCGRVSVSQ Propionamide@N-term, Propionamide(C)@5, Propionamide(C)@13 98.6 2345.1257 3
Peptide ID Hypotheses - 2.1.1.1290.4
Conf Sc Prot N Sequence Modifications Theor MW Theor m/z Obs m/z z ΔMass
96.1 9 1 TVFPDVDYVNSTEAE Deamidated(N)@10, Hex(1)HexNAc(1)NeuAc(1)(T)@12 2341.9585 781.6601 781.6630 3 0.008
85.9 9 1 TVFPDVDYVNSTEAE Deamidated(N)@10, Hex(1)HexNAc(1)NeuAc(1)(S)@11 2341.9585 781.6601 781.6630 3 0.008
Precursor MS Region
785.3239
781.6690
784.9970
780.3599
785.6687
m/z, Da
Fragmentation Evidence for Peptide
TVFPDVDYVN[Dea]ST[NHH]EAE
Residue b b+2 y
T 102.0550 51.5311 2342.9657
V 201.1234 101.0653 2241.9180
F 348.1918 174.5995 2142.8496
P 445.2445 223.1259 1995.7812
D 560.2715 280.6394 1898.7284
V 659.3399 330.1736 1783.7015
D 774.3668 387.6871 1684.6331
Y 937.4302 469.2187 1569.6061
V 1036.4986 518.7529 1406.5428
N[Dea] 1151.5255 576.2664 1307.4744
S 1238.5576 619.7824 1192.4475
T[NHH] 1995.8329 998.4201 1105.4154
E 2124.8754 1062.9414 348.1401
A 2195.9126 1098.4599 219.0975
E 2324.9552 1162.9812 148.0604
y1 y2 b2 b3 b6 b7 b8 b9 b10
274.10
204.09
166.06
292.10
366.15
493.19
572.22
660.35
844.37
1036.52
1151.54
1339.63
1468.64
1539.73
1687.75
503.26
533.25
564.15
651.31
755.16
963.08
1047.94
1093.60
1157.62
1424.42
1669.64
1678.55
Intensity
m/z, Da
Type here to search
ENG
10:27 PM
2/04/2020
