## Supplementary Information 3 for "Analysis of coagulation factor IX in bioreactor cell culture medium predicts yield and quality of the purified product"

20190702_Co4Ra2NeuGcOglyLacNAc_20190509_BenSchulz_Luci_H2a_Ch.wiff_Byonic_D359_D364 1
20190702_Co4Ra2NeuGcOglyLacNAc_20190509_BenSchulz_Luci_H2a_Ch.wiff_Byonic_E15_E17_E20_E21 2
20190702_Co4Ra2NeuGcOglyLacNAc_20190509_BenSchulz_Luci_H2a_Ch.wiff_Byonic_E20_E21_E26_E27_E30[+44] 3
20190702_Co4Ra2NeuGcOglyLacNAc_20190509_BenSchulz_Luci_H2a_Ch.wiff_Byonic_E20_E21_E30_E33_E36 4
20190702_Co4Ra2NeuGcOglyLacNAc_20190509_BenSchulz_Luci_H2a_Ch.wiff_Byonic_E33_E36_E40 5

20190702_Co4Ra2NeuGcOglyLacNAc_20190509_BenSchulz_Luci_H2a_Ch.wiff_Byonic_E33_E36_E40[+44] 6
20190702_Co4Ra2NeuGcOglyLacNAc_20190509_BenSchulz_Luci_H2a_Ch.wiff_Byonic_E33_E36_E40[+44]x2 7
20190702_Co4Ra2NeuGcOglyLacNAc_20190509_BenSchulz_Luci_H2a_Ch.wiff_Byonic_S53[+426] 8
20190702_Co4Ra2NeuGcOglyLacNAc_20190509_BenSchulz_Luci_H2a_Ch.wiff_Byonic_T179[+656] 9
20190702_Co4Ra2NeuGcOglyLacNAc_20190509_BenSchulz_Luci_H2a_Ch.wiff_Byonic_T179[+947] 10

Common4rare2NeuGc50_20181106_Schulz_Luci_H2aG.wiff_20181116_Byonic_D47[+16] 11
Common4rare2NeuGc50_20181106_Schulz_Luci_H2aG.wiff_20181116_Byonic_D49[+16] 12
Common4rare2NeuGc50_20181106_Schulz_Luci_H2aG.wiff_20181116_Byonic_D104 13
Common4rare2NeuGc50_20181106_Schulz_Luci_H2aG.wiff_20181116_Byonic_D104[+16] 14
Common4rare2NeuGc50_20181106_Schulz_Luci_H2aG.wiff_20181116_Byonic_D276_D292 15

Common4rare2NeuGc50_20181106_Schulz_Luci_H2aG.wiff_20181116_Byonic_D292[+16] 16
Common4rare2NeuGc50_20181106_Schulz_Luci_H2aG.wiff_20181116_Byonic_N157[+1] 17
Common4rare2NeuGc50_20181106_Schulz_Luci_H2aG.wiff_20181116_Byonic_N258 18
Common4rare2NeuGc50_20181106_Schulz_Luci_H2aG.wiff_20181116_Byonic_N258[+1] 19
Common4rare2NeuGc50_20181106_Schulz_Luci_H2aG.wiff_20181116_Byonic_S53[+426]_S61[+802]_D64[+16]_D65[+16]_S68[+ 20

Common4rare2NeuGc50_20181106_Schulz_Luci_H2aG.wiff_20181116_Byonic_S110[+426]_T112[+802] 21
Common4rare2NeuGc50_20181106_Schulz_Luci_H2aG.wiff_20181116_Byonic_Y45_D47_D49 22
Common4rare2NeuGc50_20181106_Schulz_Luci_H2aG.wiff_20181116_Byonic_Y45[+80] 23
Common4rare2NeuGc50_20181106_Schulz_Luci_H2aGP.wiff_20181116_Byonic_N167[+1] 24
Common4rare2NeuGc50_20181106_Schulz_Luci_H2aGP.wiff_20181116_Byonic_S158[+947] 25

Common4rare2NeuGc50_20181106_Schulz_Luci_H2aT.wiff_20181119_Byonic_D64 26
Common4rare2NeuGc50_20181106_Schulz_Luci_H2aT.wiff_20181119_Byonic_D64[+16] 27
Common4rare2NeuGc50_20181106_Schulz_Luci_H2aT.wiff_20181119_Byonic_D85 28
Common4rare2NeuGc50_20181106_Schulz_Luci_H2aT.wiff_20181119_Byonic_D85[+16] 29
Common4rare2NeuGc50_20181106_Schulz_Luci_H2aT.wiff_20181119_Byonic_D203 30

Common4rare2NeuGc50_20181106_Schulz_Luci_H2aT.wiff_20181119_Byonic_D203[+16] 31
Common4rare2NeuGc50_20181106_Schulz_Luci_H2aT.wiff_20181119_Byonic_E7_E9_E15 32
Common4rare2NeuGc50_20181106_Schulz_Luci_H2aT.wiff_20181119_Byonic_E7_E9_E15[+44] 33
Common4rare2NeuGc50_20181106_Schulz_Luci_H2aT.wiff_20181119_Byonic_E7_E9_E15[+44]x2 34
Common4rare2NeuGc50_20181106_Schulz_Luci_H2aT.wiff_20181119_Byonic_E7_E9_E15[+44]x3 35

Common4rare2NeuGc50_20181106_Schulz_Luci_H2aT.wiff_20181119_Byonic_S141[+656] 36
Common4rare2NeuGc50_20181106_Schulz_Luci_H2aT.wiff_20181119_Byonic_S141[+947] 37
Common4rare2NeuGc50_20181106_Schulz_Luci_H2aT.wiff_20181119_Byonic_T38[+656] 38
Common4rare2NeuGc50_20181106_Schulz_Luci_H2aT.wiff_20181119_Byonic_T38[+947] 39
Common4rare2NeuGc50_20181106_Schulz_Luci_H2aT.wiff_20181119_Byonic_T38[+963] 40

Common4rare2NeuGc50_20181106_Schulz_Luci_H2bG.wiff_20181116_Byonic_Y155[+80] 41
Common4rare2NeuGc50_20181106_Schulz_Luci_H2bGP.wiff_20181116_Byonic_D276{+16] 42
Common4rare2NeuGc50_20181106_Schulz_Luci_H2bGP.wiff_20181116_Byonic_T169orT172[+947] 43
Common4rare2NeuGc50_20181106_Schulz_Luci_H2bT.wiff_20181119_Byonic_E26_E27_E30_E33_E36[+44]x3 44
Common4rare2NeuGc50_20181106_Schulz_Luci_H2bTP.wiff_20181119_Byonic_S53[+426]_S61 45

ProteinPilot_E40[+44] ______________________________________________ 46

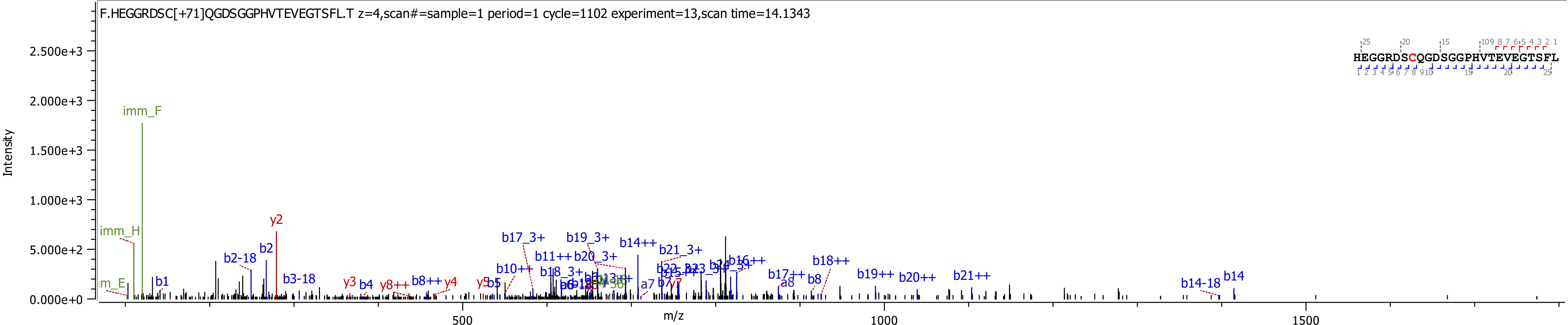

F.HEGGRDSC[+71]QGDSGGPHVTEVEGTSFL.T z=4,scan#=sample=1 period=1 cycle=1102 experiment=13,scan time=14.1343
HEGGRDSCQGDSGGPHVTEVEGTSFL
Intensity
2.500e+3
2.000e+3
1.500e+3
1.000e+3
5.000e+2
0.000e+0
m/z
500
1000
1500
imm_F
imm_H
y2
b2-18
b2
b3-18
b1
y3
b4
y8++
b8++
y4
y5
b5
b10++
b17_3+
b19_3+
b11++
b20_3+
b18_3+
b14++
b21_3+
b16++
b17++
a8
b8
b18++
b19++
b20++
b21++
b14-18
b14
a7
b7
y7

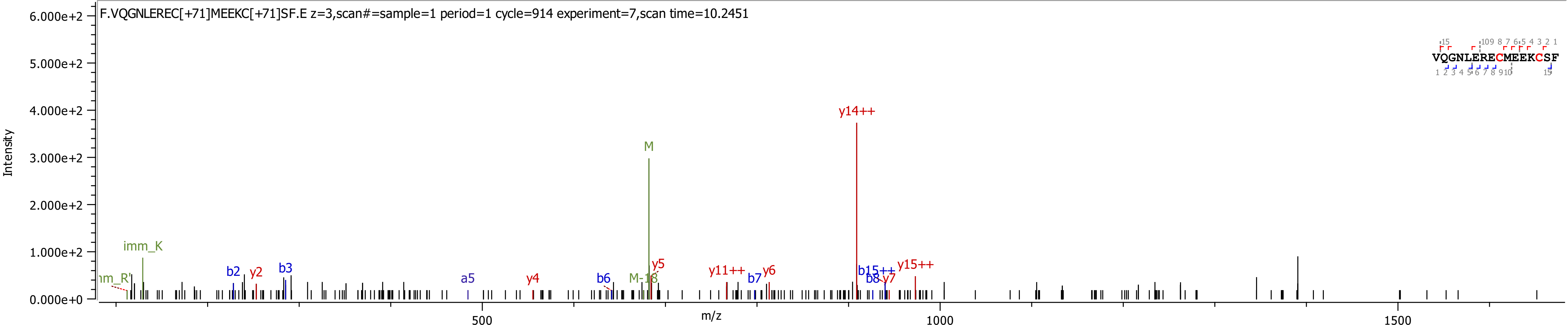

F.VQGNLEREC[+71]MEEKC[+71]SF.E z=3,scan#=sample=1 period=1 cycle=914 experiment=7,scan time=10.2451
Intensity
m/z
VQGNLERECMEEKCSF
y14++
M
imm_K
b2
y2
b3
a5
y4
b6
M-18
y5
y11++
b7
y6
b15++
b8
y7
y15++

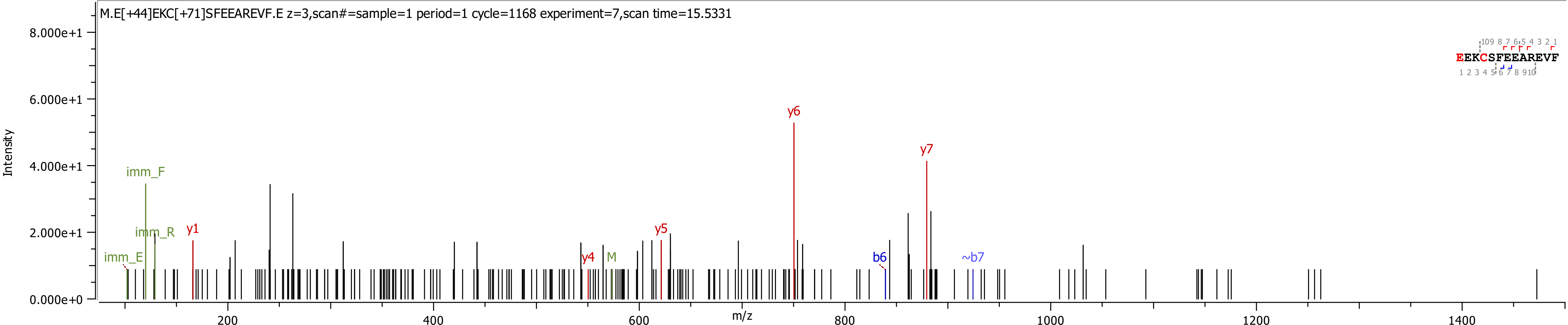

M.E[+44]EKC[+71]SFEEAREVF.E z=3,scan#=sample=1 period=1 cycle=1168 experiment=7,scan time=15.5331
Intensity
8.000e+1
6.000e+1
4.000e+1
2.000e+1
0.000e+0
imm_F
imm_R
imm_E
y1
y4
M
y5
y6
y7
b6
~b7
EEKCSFEEAREVF
200
400
600
800
1000
1200
1400
m/z

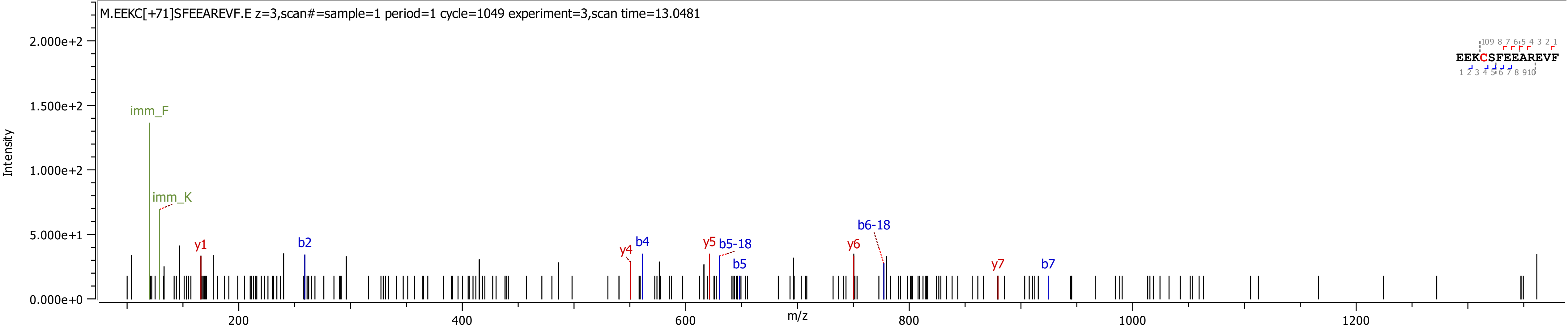
M.EEKC[+71]SFEEAREVF.E z=3,scan#=sample=1 period=1 cycle=1049 experiment=3,scan time=13.0481
EEKCSFEEAREVF
Intensity
m/z
imm_F
imm_K
y1
b2
y4
b4
y5
b5-18
b5
y6
b6-18
y7
b7
2.000e+2
1.500e+2
1.000e+2
5.000e+1
0.000e+0
200
400
600
800
1000
1200

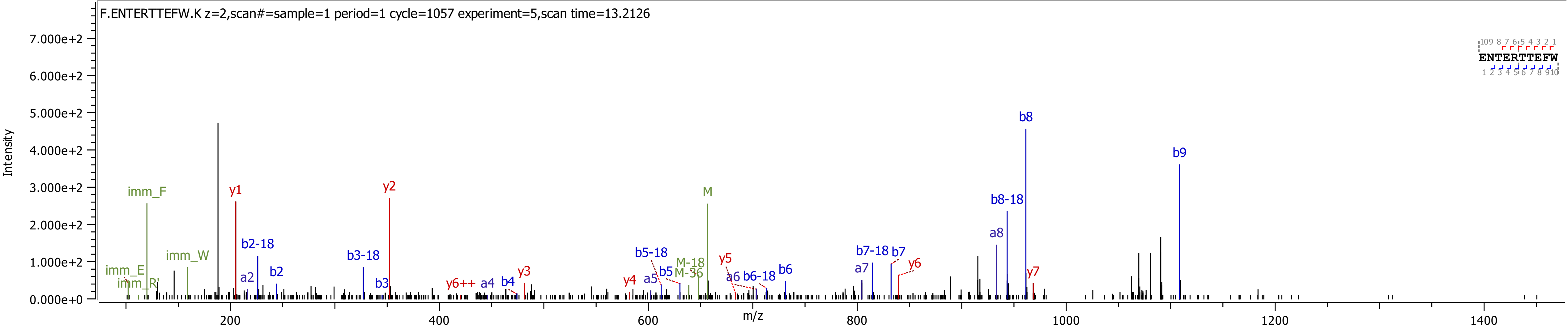

F.ENTERTTEFW.K z=2,scan#=sample=1 period=1 cycle=1057 experiment=5,scan time=13.2126
Intensity
7.000e+2
6.000e+2
5.000e+2
4.000e+2
3.000e+2
2.000e+2
1.000e+2
0.000e+0
m/z
200
400
600
800
1000
1200
1400
ENTERTTEFW
imm_F
imm_E
imm_R
imm_W
y1
y2
y3
y4
y5
y6
y6++
y7
a2
a4
a5
a6
a7
a8
b2
b2-18
b3
b3-18
b4
b5
b5-18
b6
b6-18
b7
b7-18
b8
b8-18
b9
M
M-18
M-36

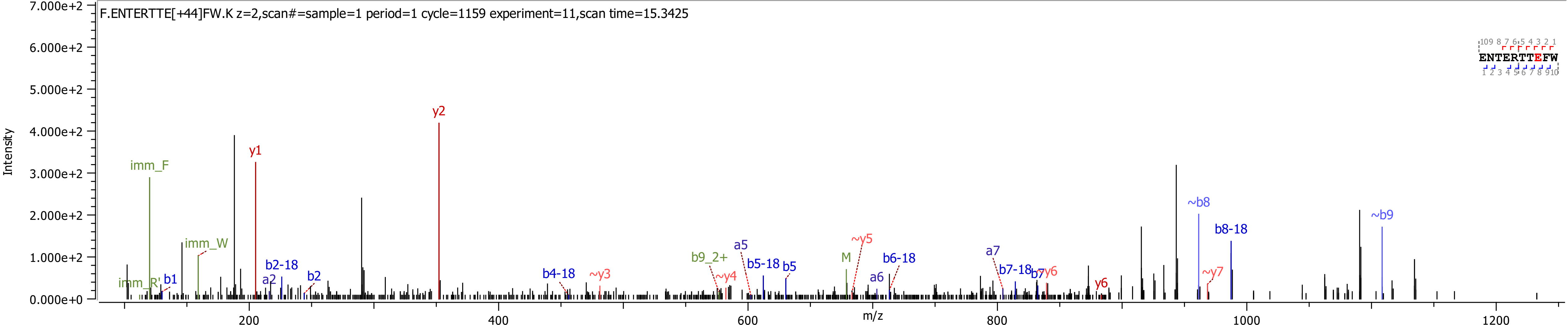
F.ENTERTTE[+44]FW.K z=2,scan#=sample=1 period=1 cycle=1159 experiment=11,scan time=15.3425
Intensity
m/z
7.000e+2
6.000e+2
5.000e+2
4.000e+2
3.000e+2
2.000e+2
1.000e+2
0.000e+0
200
400
600
800
1000
1200
imm_F
imm_W
imm_R'
b1
a2
b2-18
b2
y1
y2
b4-18
~y3
b9_2+
~y4
a5
b5-18
b5
M
~y5
a6
b6-18
a7
b7-18
b7
~y6
y6
~b8
~y7
b8-18
~b9
ENTERTTEFW

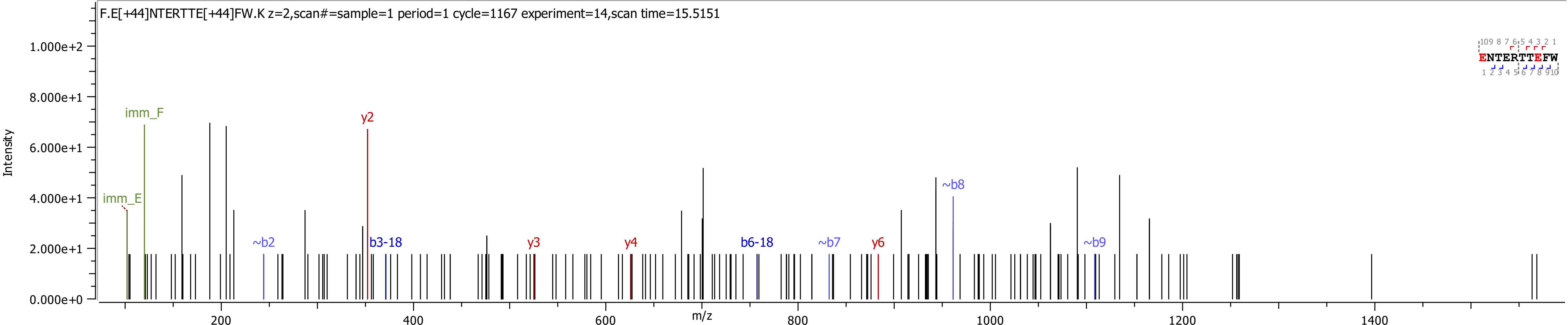
F.E[+44]NTERTTE[+44]FW.K z=2,scan#=sample=1 period=1 cycle=1167 experiment=14,scan time=15.5151
Intensity
m/z
imm_F
imm_E
y2
~b2
b3-18
y3
y4
b6-18
~b7
y6
~b8
~b9
ENTERTTEFW
0.000e+0
2.000e+1
4.000e+1
6.000e+1
8.000e+1
1.000e+2
200
400
600
800
1000
1200
1400

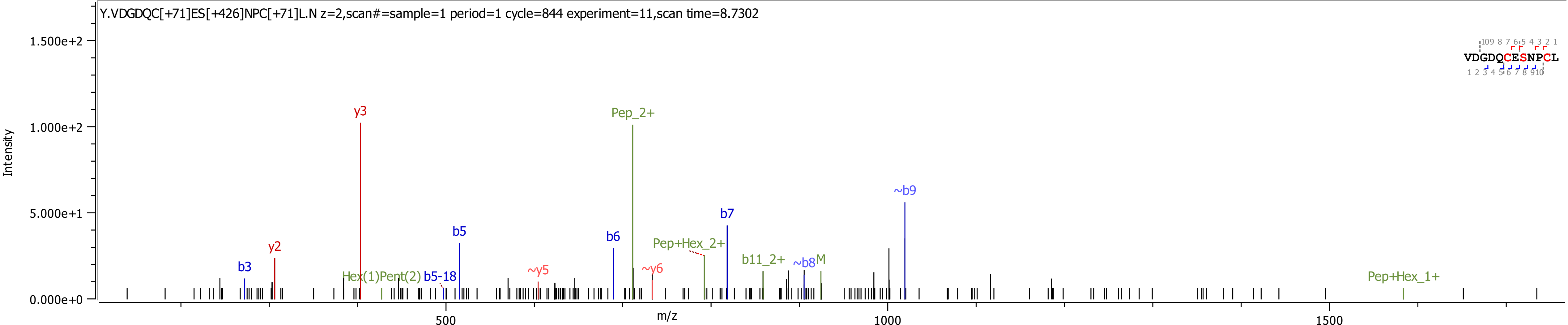

Y.VDGDQC[+71]ES[+426]NPC[+71]L.N z=2,scan#=sample=1 period=1 cycle=844 experiment=11,scan time=8.7302
Intensity
1.500e+2
1.000e+2
5.000e+1
0.000e+0
m/z
500
1000
1500
y3
Pep_2+
~b9
b7
b5
b6
y2
b3
Pep+Hex_2+
b11_2+
~b8
M
~y5
~y6
Hex(1)Pent(2)
b5-18
Pep+Hex_1+
VDGDQCESNPCL

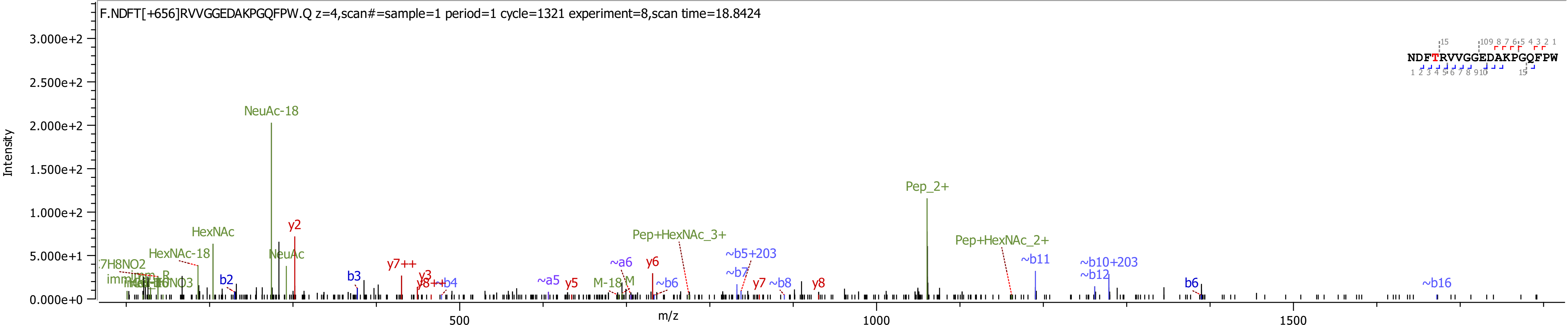

F.NDFT[+656]RVVGGEDAKPGQFPW.Q z=4,scan#=sample=1 period=1 cycle=1321 experiment=8,scan time=18.8424
NDFTRVVGGEDAKPGQFPW
Intensity
m/z
NeuAc-18
Pep_2+
HexNAc
HexNAc-18
NeuAc
y2
Pep+HexNAc_3+
Pep+HexNAc_2+
~b5+203
~b11
~b10+203
~b12
y7++
~a6
y6
~b7
b2
b3
y3
y8++
~b4
~a5
y5
M-18
M
~b6
y7
~b8
y8
b6
~b16

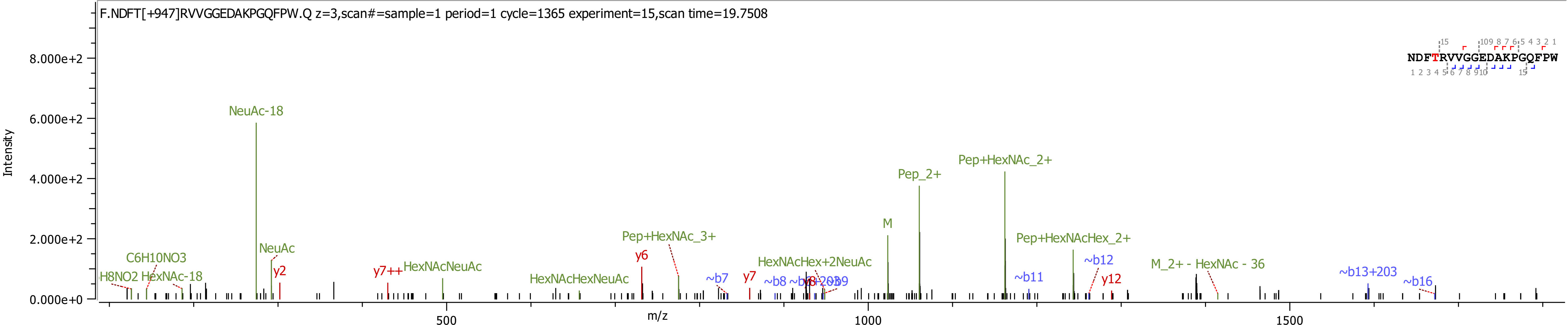

F.NDFT[+947]RVVGGEDAKPGQFPW.Q z=3,scan#=sample=1 period=1 cycle=1365 experiment=15,scan time=19.7508
NDFTRVVGGEDAKPGQFPW
Intensity
m/z
8.000e+2
6.000e+2
4.000e+2
2.000e+2
0.000e+0
500
1000
1500
NeuAc-18
NeuAc
C6H10NO3
H8NO2
HexNAc-18
y2
y7++
HexNAcNeuAc
HexNAcHexNeuAc
Pep+HexNAc_3+
y6
~b7
y7
~b8
HexNAcHex+2NeuAc
M
Pep_2+
Pep+HexNAc_2+
Pep+HexNAcHex_2+
~b11
~b12
y12
M_2+ - HexNAc - 36
~b13+203
~b16

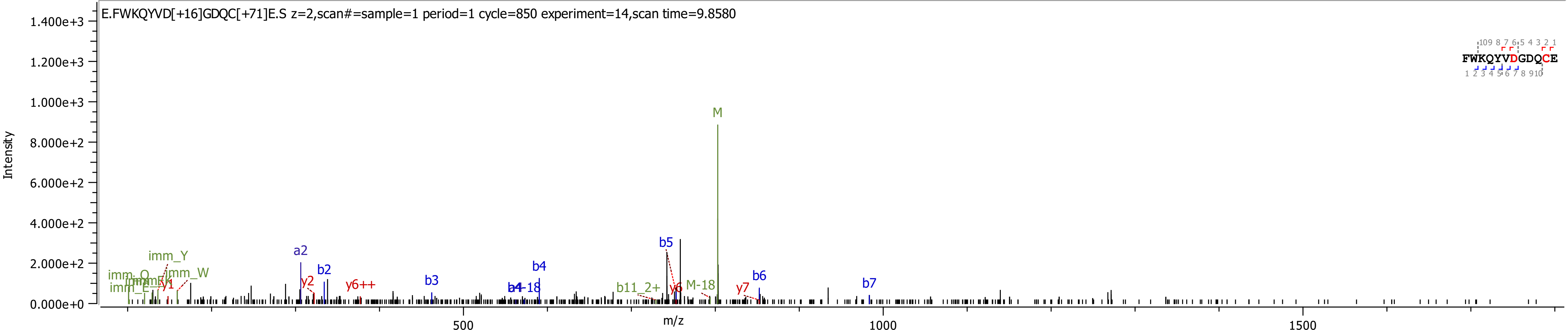
E.FWKQYVD[+16]GDQC[+71]E.S z=2,scan#=sample=1 period=1 cycle=850 experiment=14,scan time=9.8580
Intensity
m/z
FWKQYVDGDQCE
M
b5
b4
a2
b2
b3
b6
b7
y2
y6++
y7
M-18
b11_2+
imm_Y
imm_W

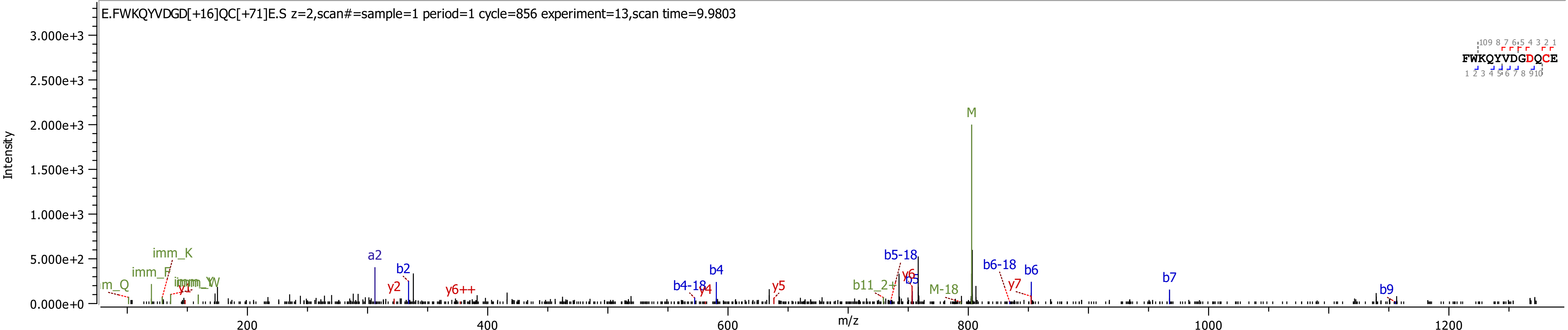

E.FWKQYVDGD[+16]QC[+71]E.S z=2,scan#=sample=1 period=1 cycle=856 experiment=13,scan time=9.9803
Intensity
3.000e+3
2.500e+3
2.000e+3
1.500e+3
1.000e+3
5.000e+2
0.000e+0
m/z
200
400
600
800
1000
1200
FWKQYVDGDQCE
M
imm_K
imm_F
m_Q
a2
b2
y2
y6++
b4
b4-18
y4
y5
b5-18
b5
y6
b11_2+
M-18
b6-18
b6
y7
b7
b9

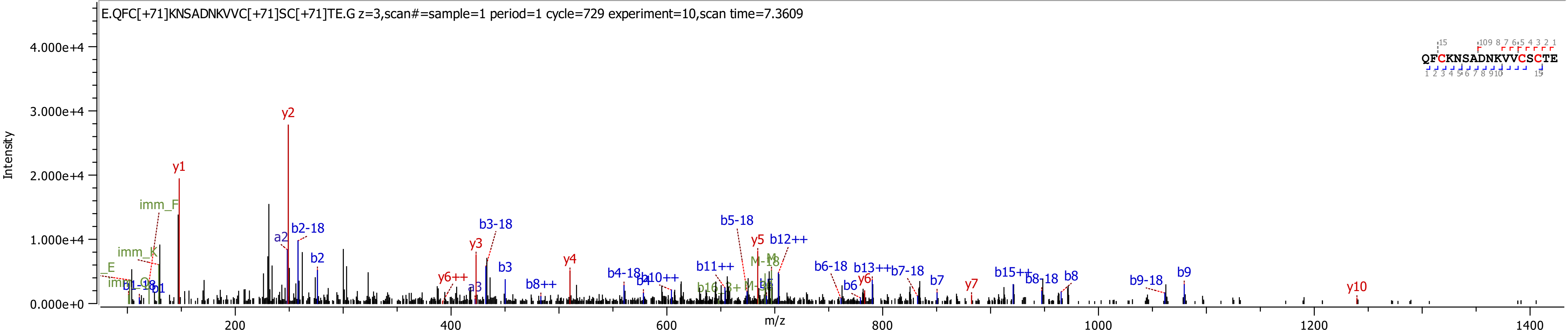
E.QFC[+71]KNSADNKVVC[+71]SC[+71]TE.G z=3,scan#=sample=1 period=1 cycle=729 experiment=10,scan time=7.3609
QFCKNSADNKVVCSCTE
Intensity
m/z
4.000e+4
3.000e+4
2.000e+4
1.000e+4
0.000e+0
200
400
600
800
1000
1200
1400
y1
y2
imm_F
imm_K
a2
b2-18
b2
y3
b3-18
b3
y4
y6++
a3
b8++
b4-18
b4
b10++
b11++
b5-18
y5
b12++
M-18
b6-18
b6
y6
b13++
b7-18
b7
y7
b15++
b8-18
b8
b9-18
b9
y10
b1
b1-18

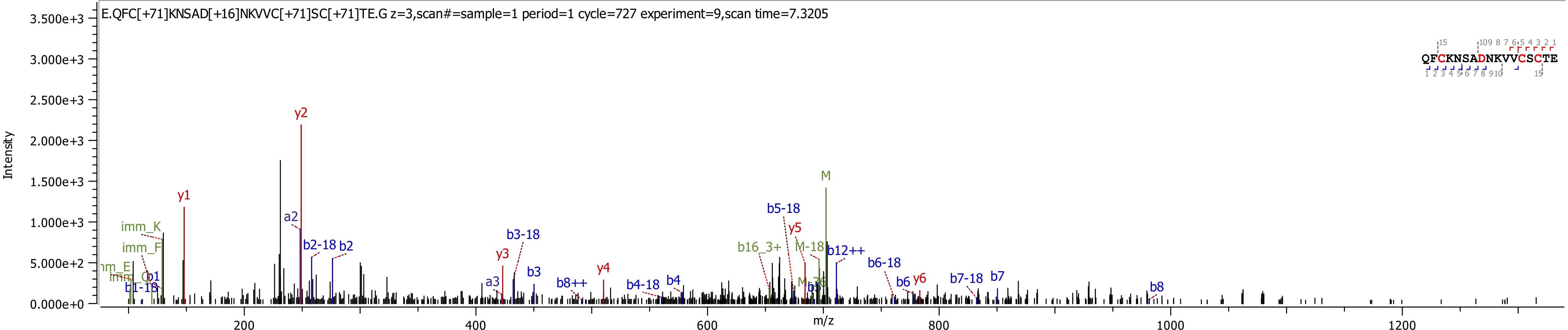
E.QFC[+71]KNSAD[+16]NKVVC[+71]SC[+71]TE.G z=3,scan#=sample=1 period=1 cycle=727 experiment=9,scan time=7.3205
QFCKNSADNKVVCSCTE
Intensity
m/z
y1
y2
a2
b2-18
b2
imm_K
imm_F
b1
b1-18
y3
a3
b3-18
b3
b8++
y4
b4-18
b4
b16_3+
b5-18
y5
M-18
M
b12++
b6-18
b6
y6
b7-18
b7
b8

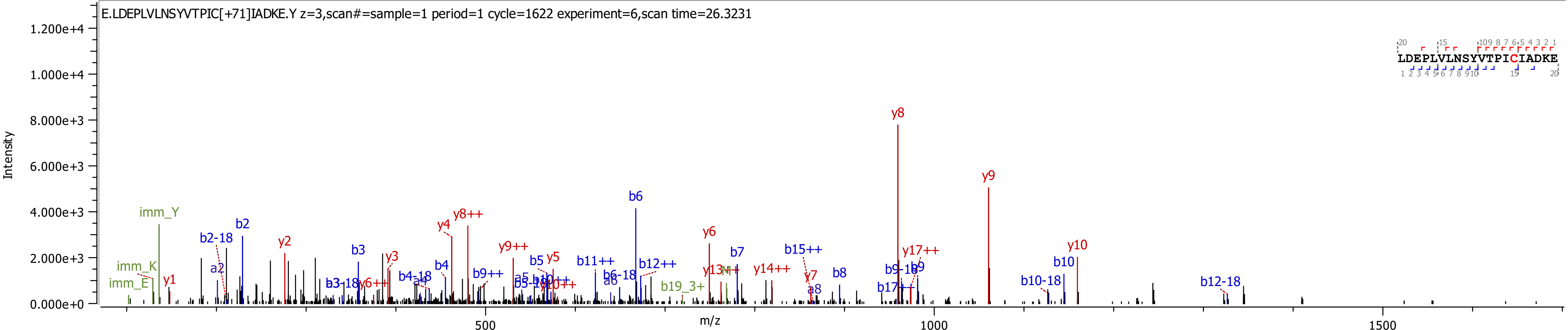
E.LDEPLVLNSYVTPIC[+71]IADKE.Y z=3,scan#=sample=1 period=1 cycle=1622 experiment=6,scan time=26.3231
LDEPLVLNSYVTPICIADKE
Intensity
m/z
y8
y9
b6
imm_Y
y8++
y4
b2
y2
b2-18
y6
y10
b10
b12-18
b10-18
y17++
b15++
b7
b8
b3
y3
a2
imm_K
imm_E
y1
b11++
b12++
y14++
y13++
b9++
y9++
b5
y5
b4
b19_3+
y7
a8
b9-18
b9
b17++
b6-18
a6
y6++
b3-18
b4-18
a4
a5
b5-18
b10++
y10++

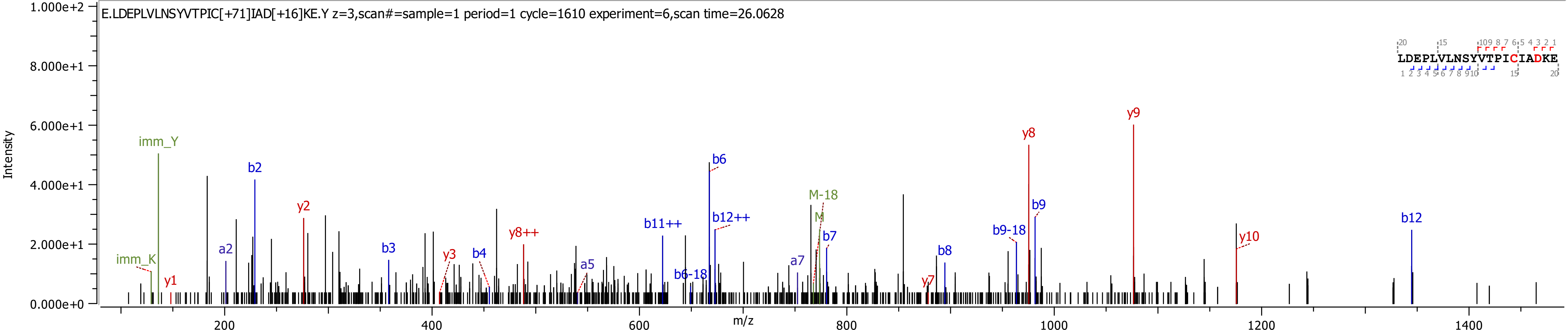
E.LDEPLVLNSYVTPIC[+71]IAD[+16]KE.Y z=3,scan#=sample=1 period=1 cycle=1610 experiment=6,scan time=26.0628
LDEPLVLNSYVTPICIADKE
Intensity
m/z
imm_Y
imm_K
y1
a2
b2
y2
b3
y3
b4
y8++
a5
b11++
b6-18
b6
b12++
a7
M-18
M
b7
y7
b8
b9-18
y8
b9
y9
y10
b12

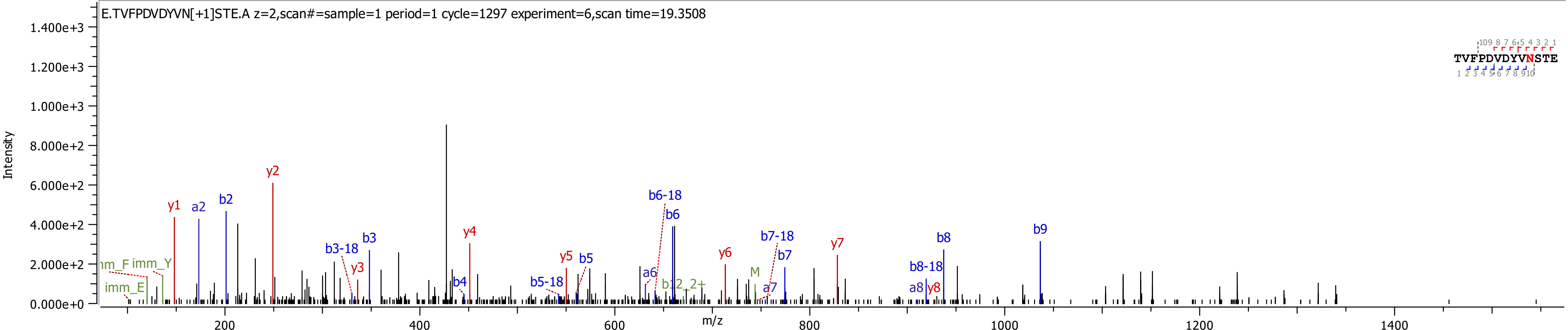
E.TVFPDVDYVN[+1]STE.A z=2,scan#=sample=1 period=1 cycle=1297 experiment=6,scan time=19.3508
Intensity
m/z
TVFPDVDYVNSTE
y1
a2
b2
y2
b3-18
b3
y3
b4
y4
b5-18
y5
b5
a6
b6-18
b6
b12_2+
y6
M
a7
b7-18
b7
y7
b8-18
a8
y8
b8
b9
imm_F
imm_Y
imm_E

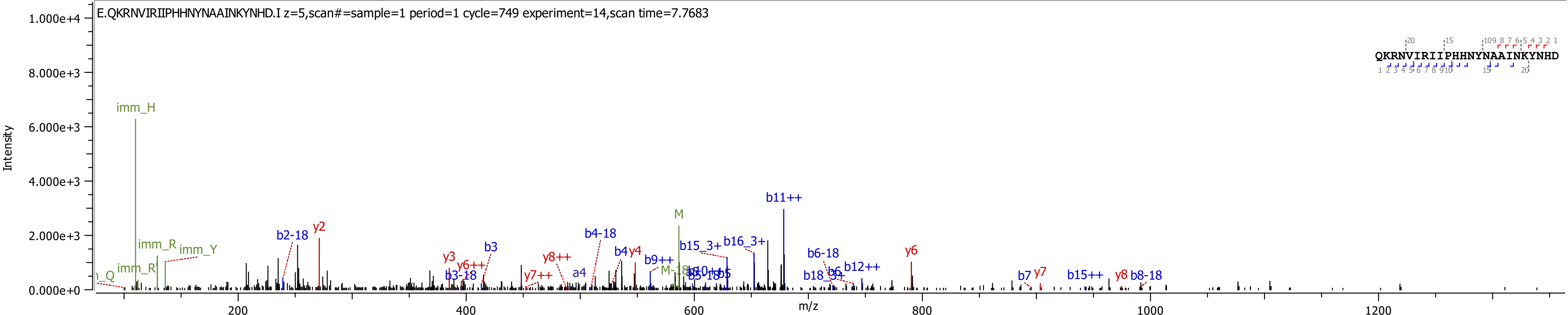

E.QKRNVIRIIPHHNYNAAINKYNHD.I z=5,scan#=sample=1 period=1 cycle=749 experiment=14,scan time=7.7683
QKRNVIRIIPHHNYNAAINKYNHD
Intensity
m/z
1.000e+4
8.000e+3
6.000e+3
4.000e+3
2.000e+3
0.000e+0
200
400
600
800
1000
1200
imm_H
imm_R
imm_Y
imm_R
b2-18
y2
y3
b3
y6++
b3-18
y8++
y7++
a4
b4-18
b4
y4
b9++
M
M-18
b15_3+
b16_3+
b11++
b5-18
b5
b6-18
b6
b12++
b18_3+
y6
b7
y7
b15++
y8
b8-18

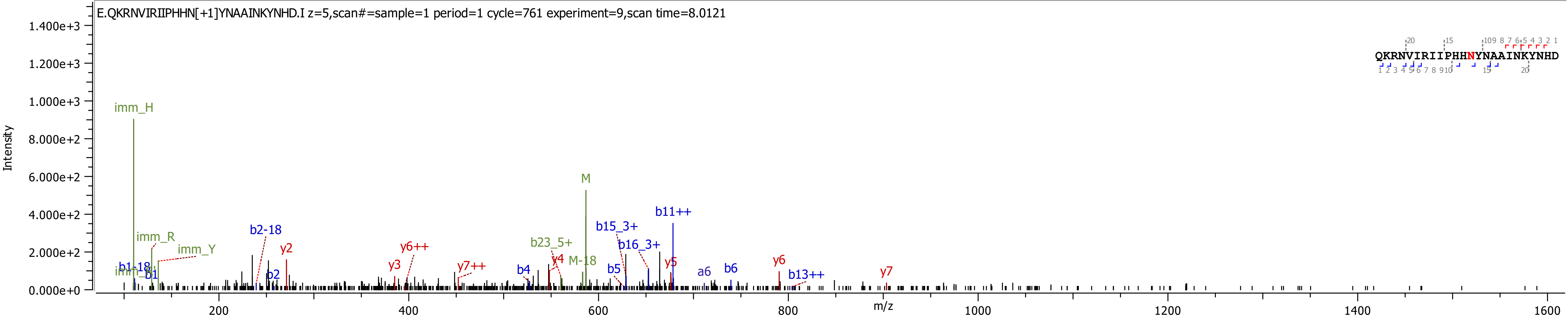

E.QKRNVIRIIPHHN[+1]YNAAINKYNHD.I z=5,scan#=sample=1 period=1 cycle=761 experiment=9,scan time=8.0121
QKRNVIRIIPHHNYNAAINKYNHD
Intensity
m/z
imm_H
imm_R
imm_Y
b1-18
b1
b2-18
b2
y2
y3
y6++
y7++
b4
y4
b23_5+
M-18
M
b15_3+
b16_3+
b5
b11++
y5
a6
b6
y6
b13++
y7

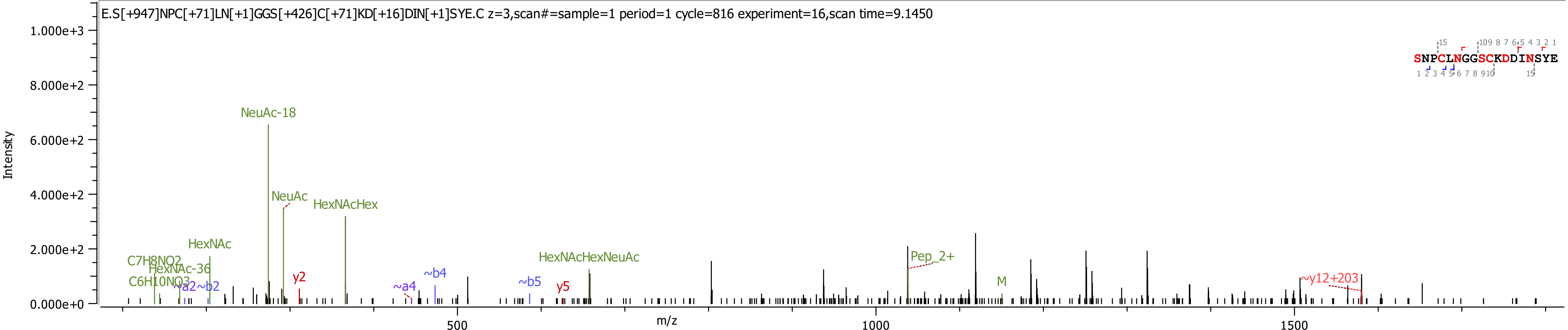

E.S[+947]NPC[+71]LN[+1]GGS[+426]C[+71]KD[+16]DIN[+1]SYE.C z=3,scan#=sample=1 period=1 cycle=816 experiment=16,scan time=9.1450
SNPCLNGGSCKDDINSYE
Intensity
1.000e+3
8.000e+2
6.000e+2
4.000e+2
2.000e+2
0.000e+0
m/z
500
1000
1500
NeuAc-18
NeuAc
HexNAcHex
HexNAc
C7H8NO2
HexNAc-36
C6H10NO3
~a2
~b2
y2
~a4
~b4
~b5
y5
HexNAcHexNeuAc
Pep_2+
M
~y12+203

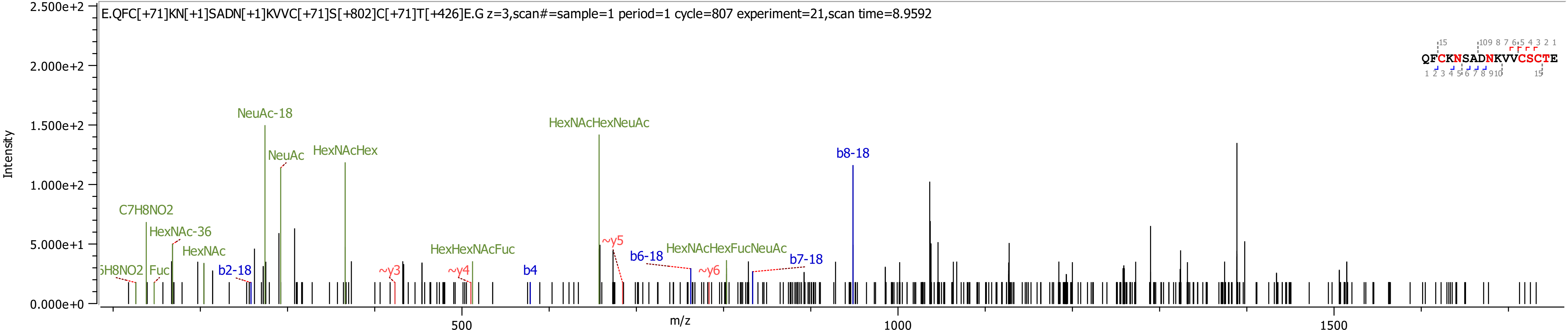

E.QFC[+71]KN[+1]SADN[+1]KVVC[+71]S[+802]C[+71]T[+426]E.G z=3,scan#=sample=1 period=1 cycle=807 experiment=21,scan time=8.9592
QFCKNSADNKVVCSCTE
Intensity
m/z
2.500e+2
2.000e+2
1.500e+2
1.000e+2
5.000e+1
0.000e+0
500
1000
1500
NeuAc-18
HexNAcHexNeuAc
NeuAc
HexNAcHex
b8-18
C7H8NO2
HexNAc-36
HexNAc
~y5
HexHexNAcFuc
b6-18
HexNAcHexFucNeuAc
b7-18
Fuc
b2-18
~y3
~y4
b4
~y6

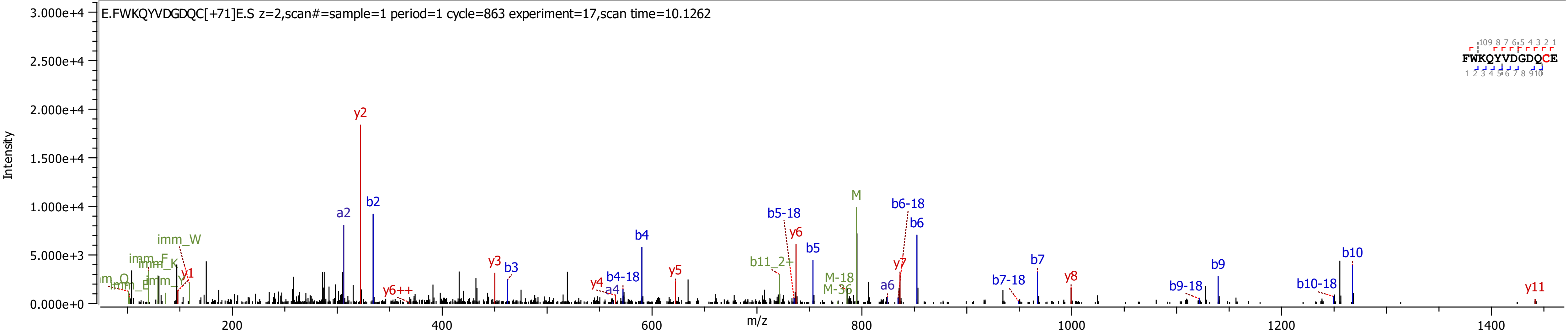
E.FWKQYVDGDQC[+71]E.S z=2,scan#=sample=1 period=1 cycle=863 experiment=17,scan time=10.1262
Intensity
m/z
FWKQYVDGDQCE
y2
a2
b2
b4
b5-18
y6
b5
M
b6-18
b6
b10
b7
b9
y8
y11
imm_W
imm_F
imm_K
y1
y3
b3
y4
a4
b4-18
y5
b11_2+
M-18
M-36
y7
a6
b7-18
b9-18
b10-18
y6++

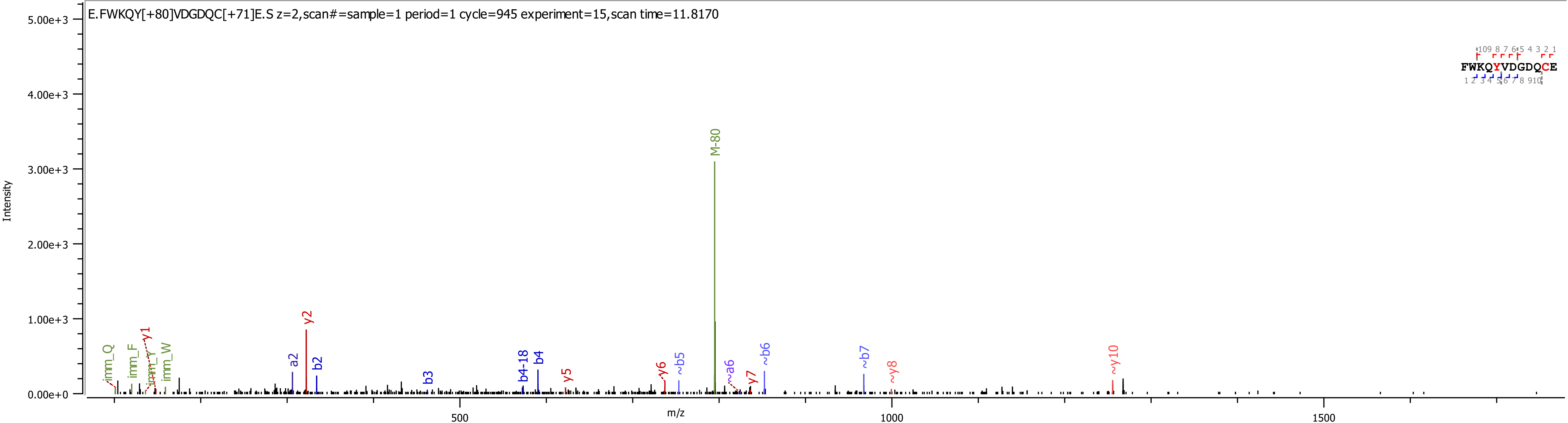

E.FWKQY[+80]VDGDQC[+71]E.S z=2,scan#=sample=1 period=1 cycle=945 experiment=15,scan time=11.8170
FWKQYVDGDQCE
Intensity
5.00e+3
4.00e+3
3.00e+3
2.00e+3
1.00e+3
0.00e+0
m/z
500
1000
1500
M-80
imm_Q
imm_F
imm_Y
imm_W
y1
a2
y2
b2
b3
b4-18
b4
y5
y6
~b5
~a6
y7
~b6
~b7
~y8
~y10

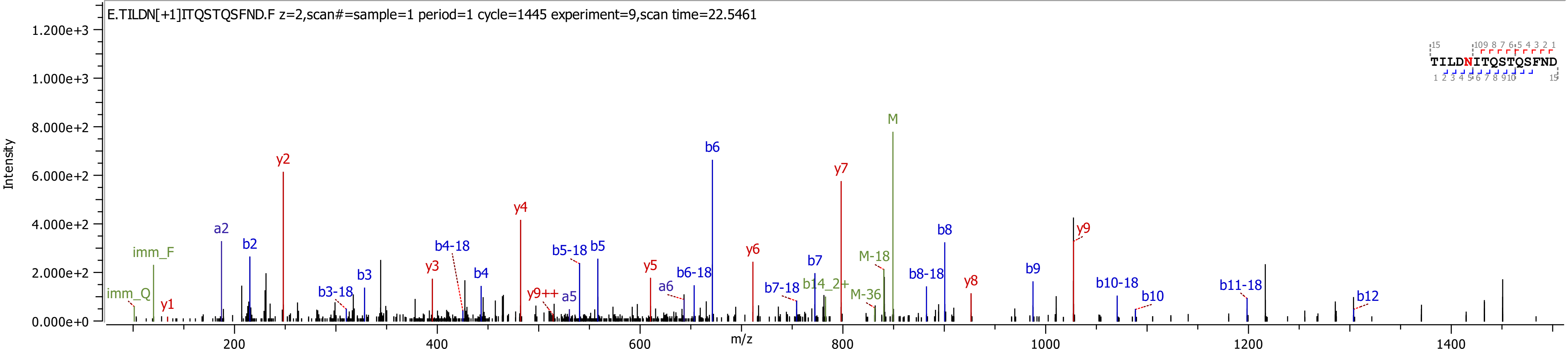
E.TILDN[+1]ITQSTQSFND.F z=2,scan#=sample=1 period=1 cycle=1445 experiment=9,scan time=22.5461
TILDNITQSTQSFND
Intensity
m/z
imm_Q
imm_F
y1
a2
b2
y2
b3-18
b3
y3
b4-18
b4
y4
y9++
a5
b5-18
b5
y5
a6
b6-18
b6
y6
b7-18
b7
b14_2+
y7
M-18
M-36
M
b8-18
b8
y8
b9
y9
b10-18
b10
b11-18
b12

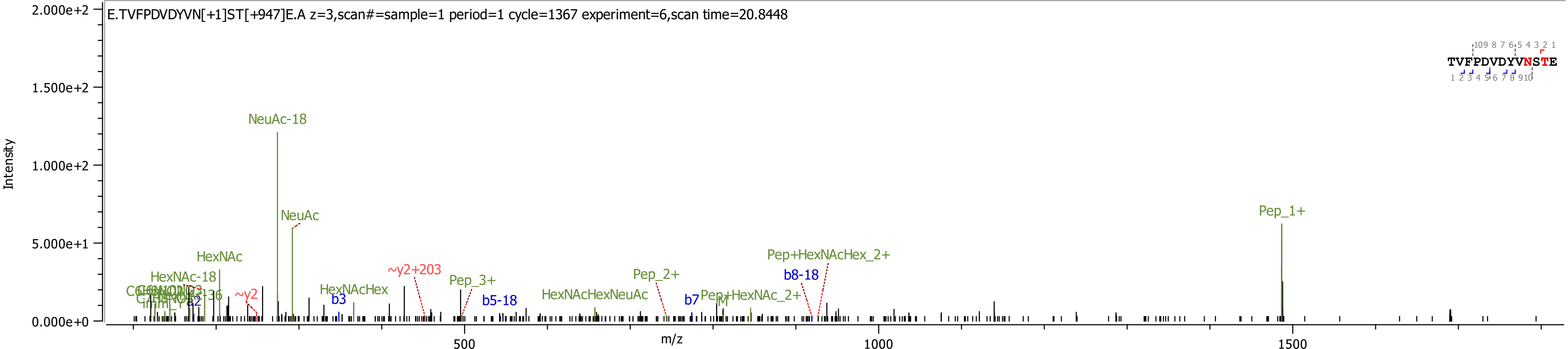
E.TVFPDVDYVN[+1]ST[+947]E.A z=3,scan#=sample=1 period=1 cycle=1367 experiment=6,scan time=20.8448
Intensity
2.000e+2
1.500e+2
1.000e+2
5.000e+1
0.000e+0
m/z
500
1000
1500
TVFPDVDYVNSTE
NeuAc-18
NeuAc
HexNAc
HexNAc-18
~y2
~y2+203
HexNAcHex
b3
Pep_3+
b5-18
HexNAcHexNeuAc
Pep_2+
b7
Pep+HexNAc_2+
b8-18
Pep+HexNAcHex_2+
Pep_1+

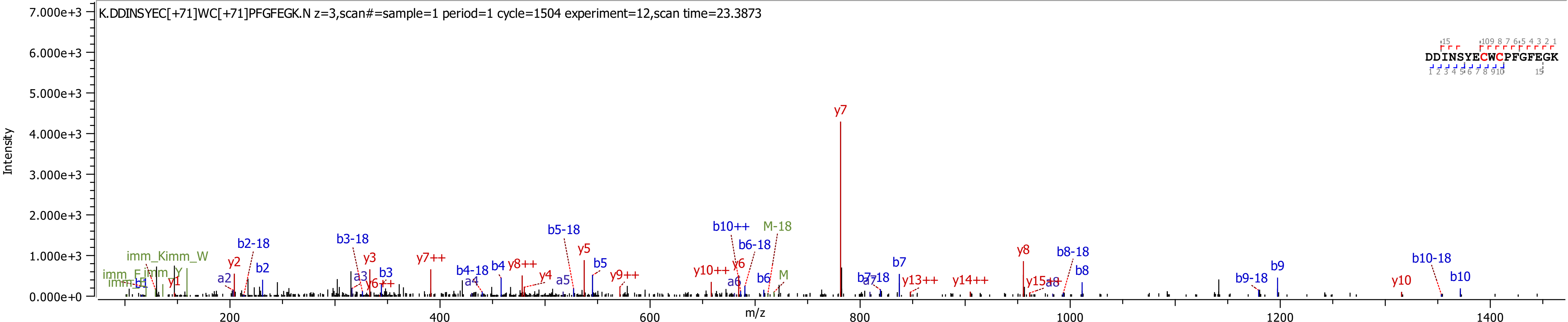
K.DDINSYEC[+71]WC[+71]PFGFEGK.N z=3,scan#=sample=1 period=1 cycle=1504 experiment=12,scan time=23.3873
DDINSYECWCPFGFEGK
Intensity
7.000e+3
6.000e+3
5.000e+3
4.000e+3
3.000e+3
2.000e+3
1.000e+3
0.000e+0
m/z
200
400
600
800
1000
1200
1400
y7
b10++
M-18
b5-18
b6-18
b2-18
b3-18
y5
b8-18
b10-18
y8
imm_Kimm_W
y2
b2
y3
y7++
b4-18
b4
y8++
b5
y4
a5
y9++
y10++
y6
b7
b9
b8
b10
a2
a3
b3
a4
y1
y6++
a6
b6
M
y13++
y14++
y15++
a8
b9-18
y10

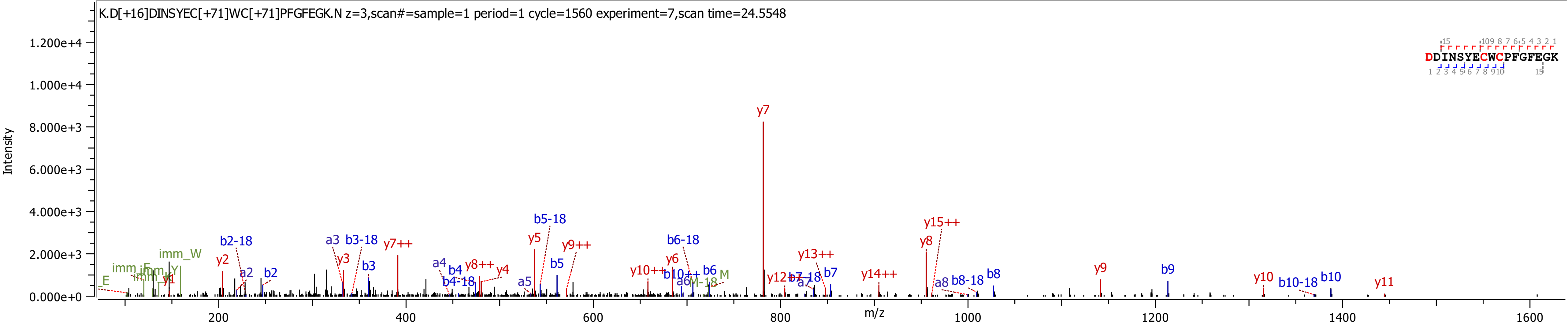
K.D[+16]DINSYEC[+71]WC[+71]PFGFEGK.N z=3,scan#=sample=1 period=1 cycle=1560 experiment=7,scan time=24.5548
DDINSYECWCPFGFEGK
Intensity
m/z
y7
b5-18
y15++
y5
y9++
y8
b2-18
a3
b3-18
y7++
b6-18
y13++
y2
y3
b3
a4
b4
y8++
y4
b5
y6
y10++
b10++
b6
M
a6
M-18
y12++
b7-18
a7
b7
y14++
y9
b9
a8
b8-18
b8
y10
b10-18
b10
y11
imm_W
imm_F
imm_Y
E
a2
b2
b4-18
a5
y1

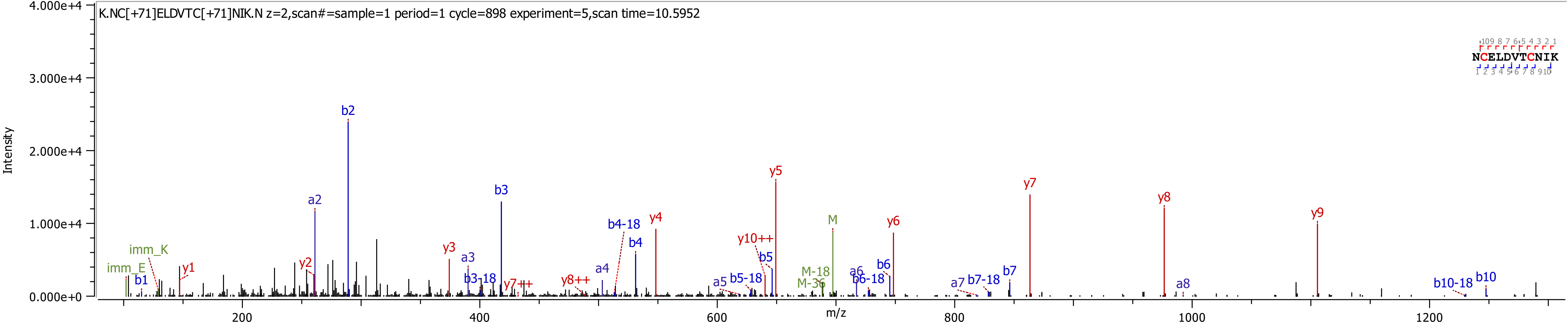
K.NC[+71]ELDVTC[+71]NIK.N z=2,scan#=sample=1 period=1 cycle=898 experiment=5,scan time=10.5952
NCELDVTCNIK
Intensity
m/z
4.000e+4
3.000e+4
2.000e+4
1.000e+4
0.000e+0
200
400
600
800
1000
1200
imm_K
imm_E
b1
y1
a2
y2
b2
y3
a3
b3-18
b3
y7++
y8++
a4
b4-18
b4
y4
a5
b5-18
y10++
b5
y5
M-18
M-36
M
a6
b6-18
b6
y6
a7
b7-18
b7
y7
y8
a8
y9
b10-18
b10

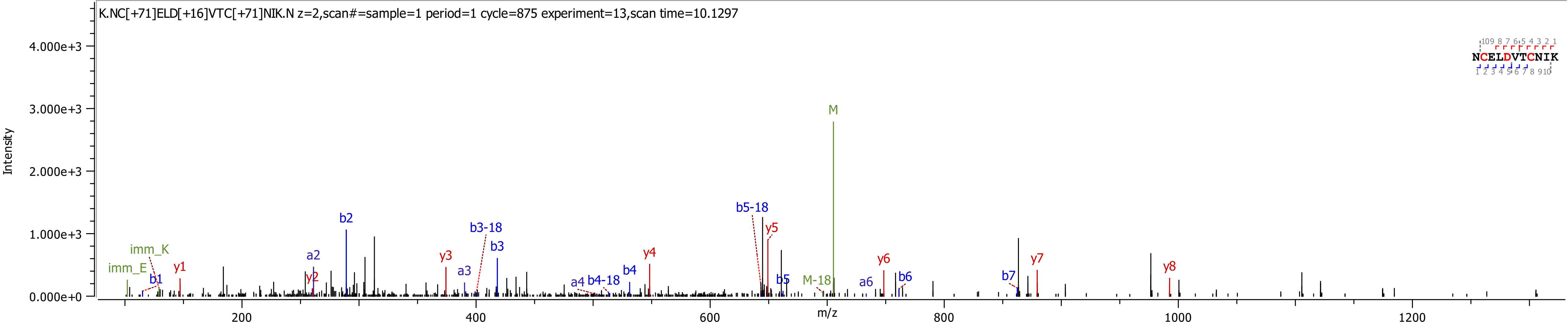

K.NC[+71]ELD[+16]VTC[+71]NIK.N z=2,scan#=sample=1 period=1 cycle=875 experiment=13,scan time=10.1297
Intensity
m/z
NCELDVTCNIK
M
b5-18
y5
b2
b3-18
b3
imm_K
imm_E
y1
b1
a2
y2
y3
a3
a4
b4-18
b4
y4
b5
M-18
a6
y6
b6
b7
y7
y8

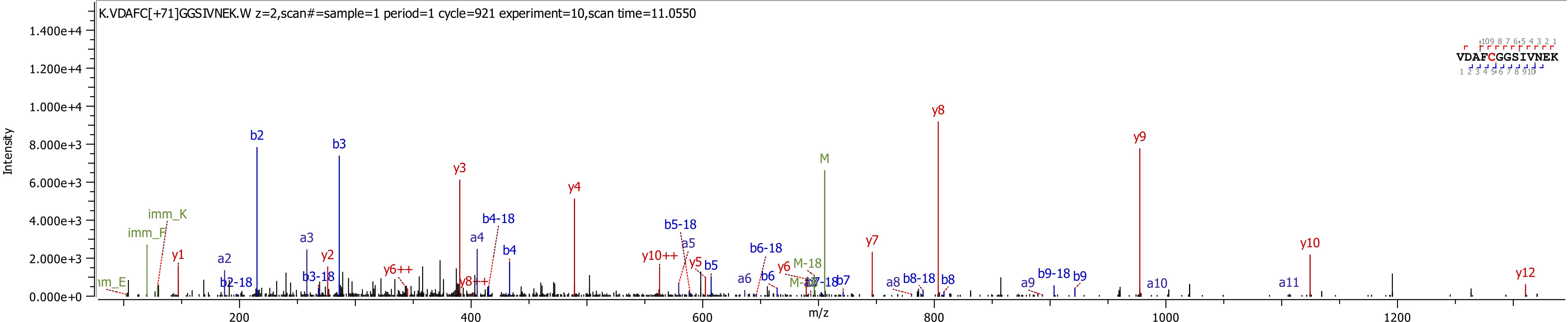
K.VDAFC[+71]GGSIVNEK.W z=2,scan#=sample=1 period=1 cycle=921 experiment=10,scan time=11.0550
VDAFCGGSIVNEK
Intensity
m/z
1.400e+4
1.200e+4
1.000e+4
8.000e+3
6.000e+3
4.000e+3
2.000e+3
0.000e+0
200
400
600
800
1000
1200
imm_E
imm_F
imm_K
y1
a2
b2-18
b2
a3
b3-18
y2
b3
y6++
y3
y8++
a4
b4-18
b4
y4
y10++
b5-18
a5
y5
b5
a6
b6-18
b6
y6
M-18
M-17
b7-18
b7
M
y7
a8
b8-18
b8
y8
a9
b9-18
b9
y9
a10
a11
y10
y12

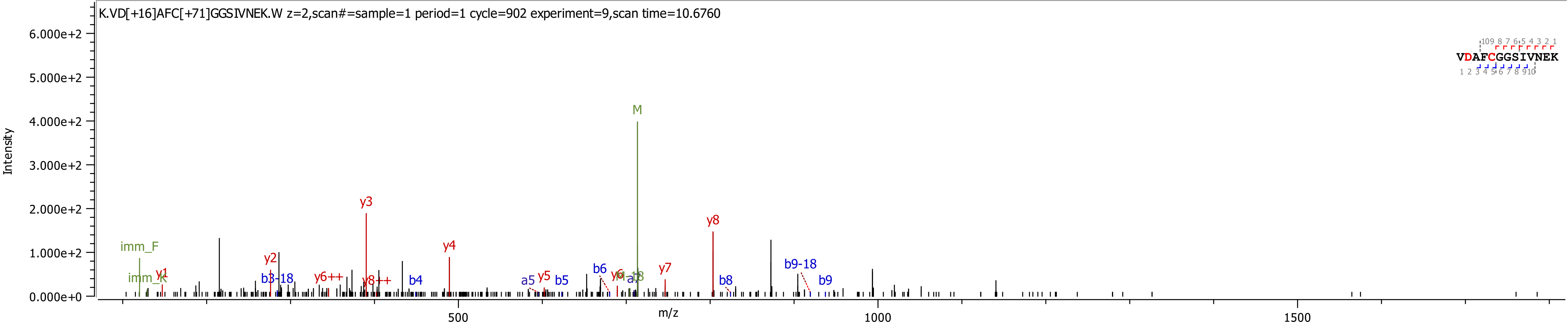
K.VD[+16]AFC[+71]GGSIVNEK.W z=2,scan#=sample=1 period=1 cycle=902 experiment=9,scan time=10.6760
VDAFCGGSIVNEK
Intensity
m/z
6.000e+2
5.000e+2
4.000e+2
3.000e+2
2.000e+2
1.000e+2
0.000e+0
500
1000
1500
M
y3
y8
y4
imm_F
imm_K
y1
y2
b3-18
y6++
y8++
b4
a5
y5
b5
b6
y7
b8
b9-18
b9

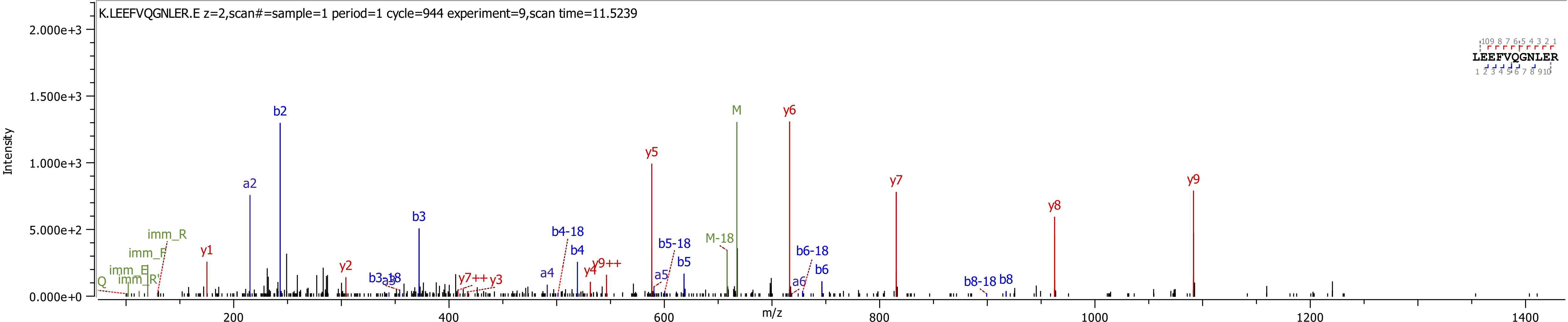

K.LEEFVQGNLER.E z=2,scan#=sample=1 period=1 cycle=944 experiment=9,scan time=11.5239
Intensity
m/z
LEEFVQGNLER
y6
M
b2
y5
y7
y9
y8
a2
b3
y1
b4-18
b4
b5-18
b5
b6-18
b6
M-18
y4
y9++
y2
y7++
y3
a4
a5
a6
b3-18
a3
b8-18
b8
imm_R
imm_F
imm_E
imm_R'
Q

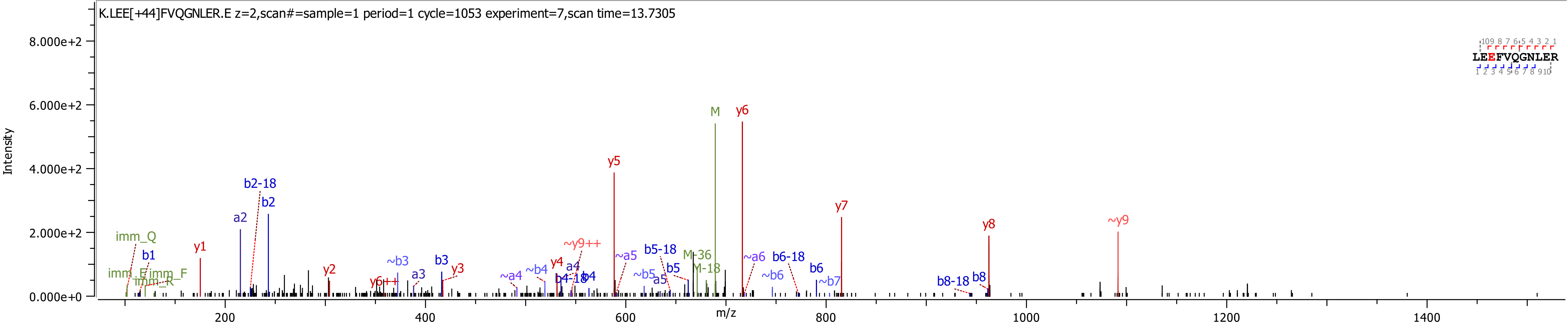
K.LEE[+44]FVQGNLER.E z=2,scan#=sample=1 period=1 cycle=1053 experiment=7,scan time=13.7305
Intensity
m/z
LEEFVQGNLER
M
y6
y5
y7
y8
~y9
b2-18
b2
a2
imm_Q
b1
y1
y2
b3
y3
~b3
a3
~y9++
y4
a4
b4
b4-18
~a4
~b4
~a5
~b5
a5
b5
b5-18
M-36
M-18
~a6
~b6
b6-18
b6
~b7
b8-18
b8
y6++
imm_E
imm_F
imm_R

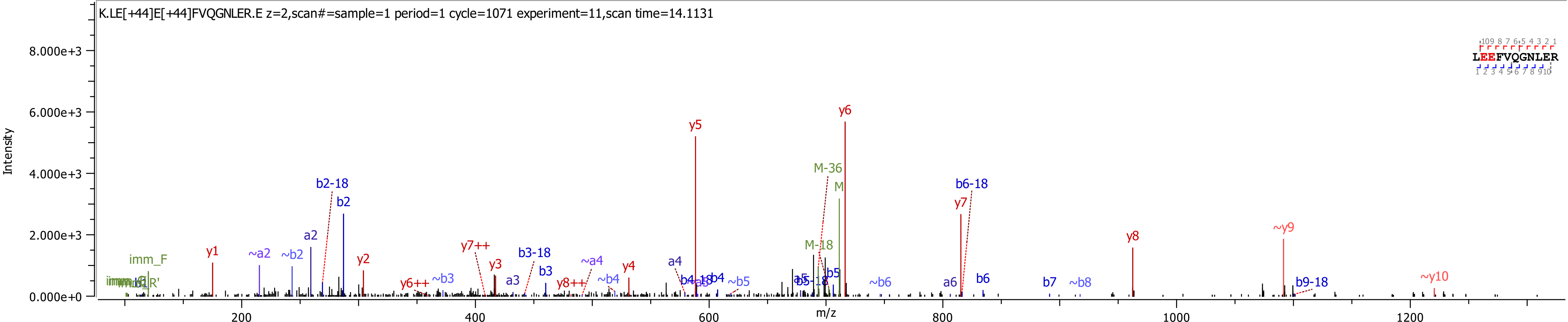

K.LE[+44]E[+44]FVQGNLER.E z=2,scan#=sample=1 period=1 cycle=1071 experiment=11,scan time=14.1131
LEEFVQGNLER
Intensity
m/z
8.000e+3
6.000e+3
4.000e+3
2.000e+3
0.000e+0
200
400
600
800
1000
1200
y1
~a2
~b2
a2
b2-18
b2
y2
y6++
~b3
y7++
y3
a3
b3-18
b3
y8++
~a4
~b4
y4
a4
y5
b4
~b5
b5
M-18
M-36
M
y6
~b6
a6
y7
b6-18
b6
b7
~b8
y8
~y9
b9-18
~y10
imm_F

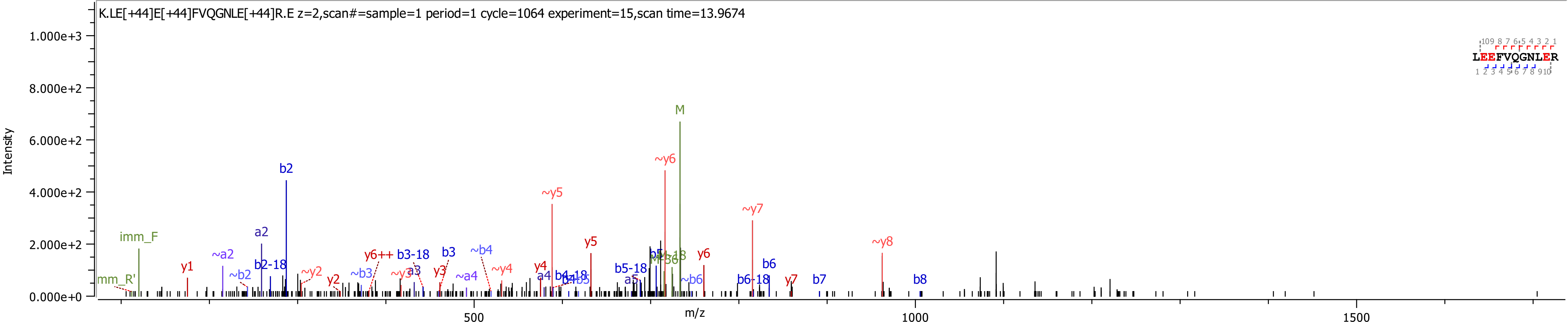
K.LE[+44]E[+44]FVQGNLE[+44]R.E z=2,scan#=sample=1 period=1 cycle=1064 experiment=15,scan time=13.9674
Intensity
m/z
1.000e+3
8.000e+2
6.000e+2
4.000e+2
2.000e+2
0.000e+0
500
1000
1500
LEEFVQGNLER
M
~y6
b2
~y5
~y7
a2
imm_F
y5
~y8
y1
~a2
b2-18
~y2
~b2
y2
y6++
b3-18
b3
~b4
~y4
y4
b5-18
y6
b6
b7
b8
y7
b6-18
~b6
~a4
a4
a5
imm_R'

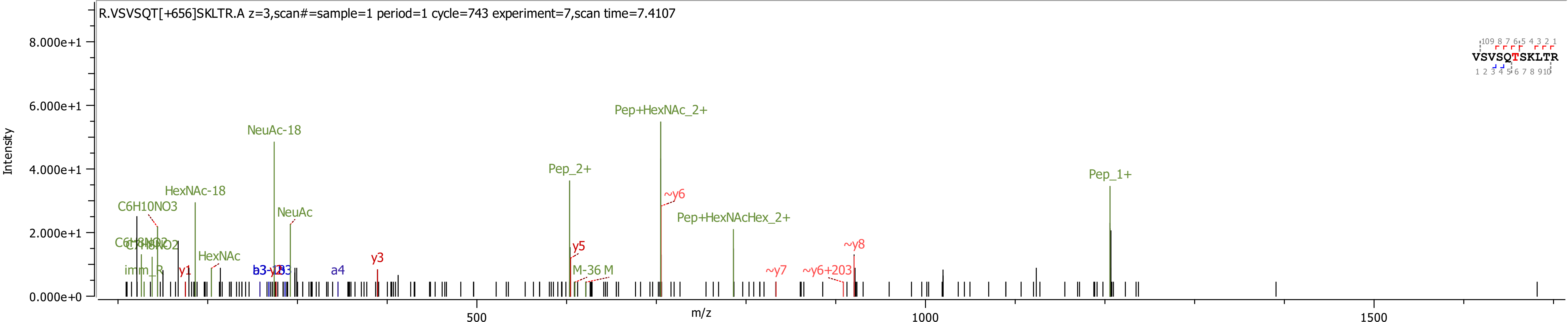
R.VSVSQT[+656]SKLTR.A z=3,scan#=sample=1 period=1 cycle=743 experiment=7,scan time=7.4107
Intensity
8.000e+1
6.000e+1
4.000e+1
2.000e+1
0.000e+0
m/z
500
1000
1500
VSVSQTSKLTR
NeuAc-18
Pep+HexNAc_2+
Pep_2+
Pep_1+
HexNAc-18
C6H10NO3
NeuAc
~y6
Pep+HexNAcHex_2+
HexNAc
y5
~y8
y3
imm_R
y1
a4
M-36 M
~y7
~y6+203

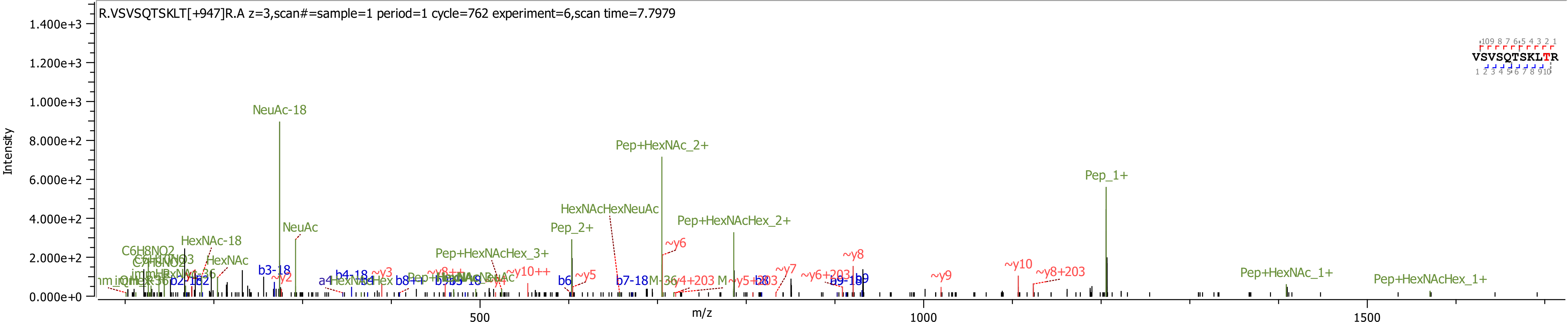
R.VSVSQTSKLT[+947]R.A z=3,scan#=sample=1 period=1 cycle=762 experiment=6,scan time=7.7979
Intensity
m/z
NeuAc-18
Pep+HexNAc_2+
Pep_1+
HexNAcHexNeuAc
Pep+HexNAcHex_2+
NeuAc
Pep_2+
HexNAc-18
HexNAc
Pep+HexNAcHex_3+
Pep+HexNAc_1+
Pep+HexNAcHex_1+
VSVSQTSKLTR

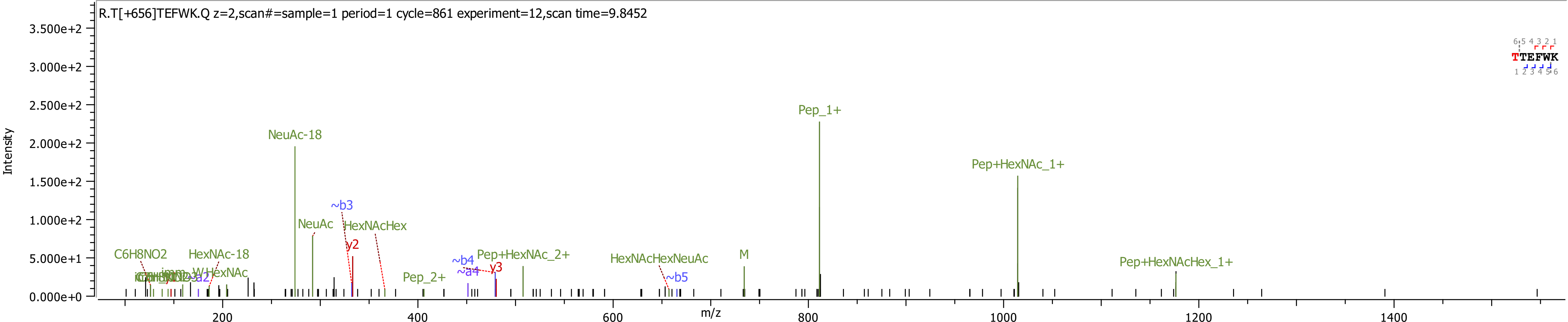

R.T[+656]TEFWK.Q z=2,scan#=sample=1 period=1 cycle=861 experiment=12,scan time=9.8452
Intensity
3.500e+2
3.000e+2
2.500e+2
2.000e+2
1.500e+2
1.000e+2
5.000e+1
0.000e+0
m/z
200
400
600
800
1000
1200
1400
NeuAc-18
NeuAc
HexNAcHex
~b3
y2
C6H8NO2
HexNAc-18
Pep_2+
~b4
~a4
y3
Pep+HexNAc_2+
HexNAcHexNeuAc
~b5
M
Pep_1+
Pep+HexNAc_1+
Pep+HexNAcHex_1+
TTEFWK

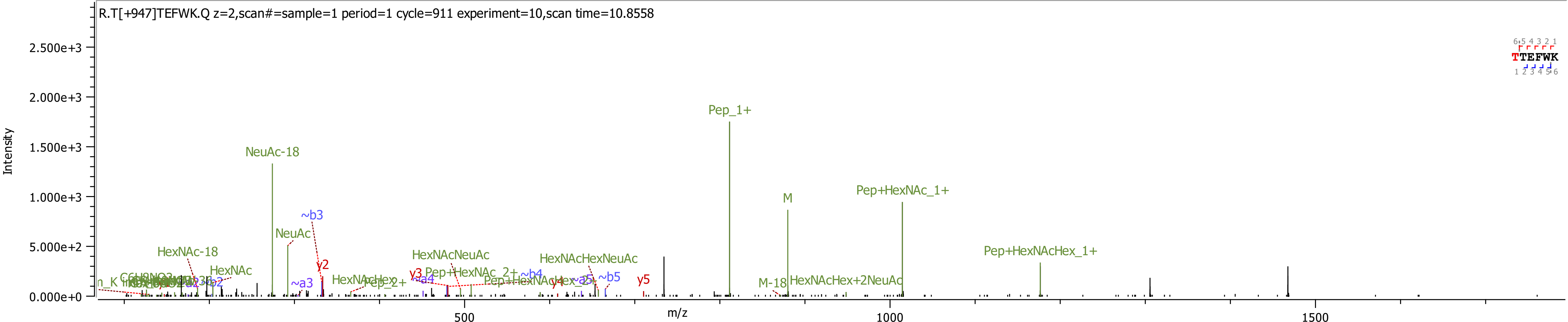

R.T[+947]TEFWK.Q z=2,scan#=sample=1 period=1 cycle=911 experiment=10,scan time=10.8558
Intensity
m/z
TTEFWK
NeuAc-18
NeuAc
HexNAc-18
HexNAc
~b3
y2
~a3
HexNAcHex
Pep_2+
y3
~a4
HexNAcNeuAc
Pep+HexNAc_2+
~b4
HexNAcHexNeuAc
Pep+HexNAcHex_2+
y4
~a5
~b5
y5
Pep_1+
M
M-18
HexNAcHex+2NeuAc
Pep+HexNAc_1+
Pep+HexNAcHex_1+
0.000e+0
5.000e+2
1.000e+3
1.500e+3
2.000e+3
2.500e+3
500
1000
1500

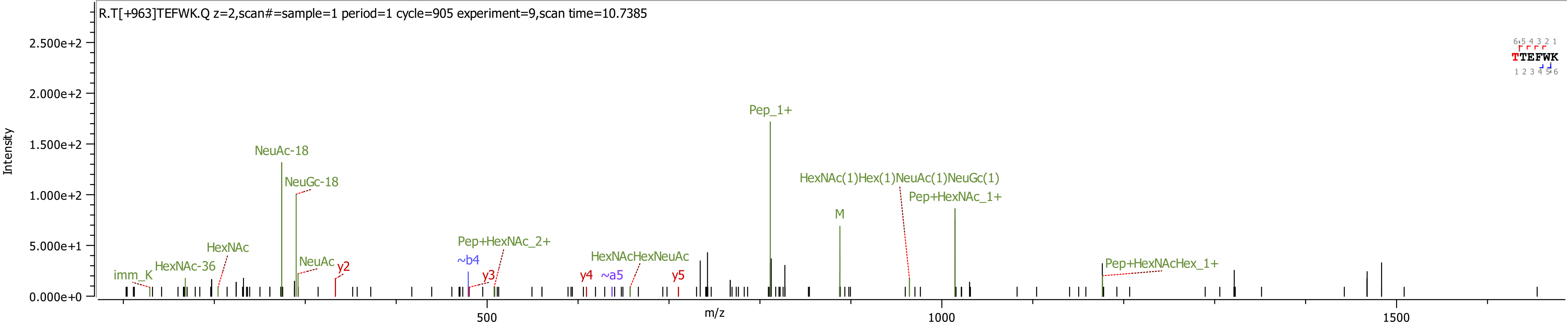

R.T[+963]TEFWK.Q z=2,scan#=sample=1 period=1 cycle=905 experiment=9,scan time=10.7385
Intensity
m/z
TTEFWK
Pep_1+
NeuAc-18
NeuGc-18
HexNAc(1)Hex(1)NeuAc(1)NeuGc(1)
Pep+HexNAc_1+
M
HexNAc
NeuAc
y2
imm_K
HexNAc-36
~b4
y3
Pep+HexNAc_2+
y4
~a5
HexNAcHexNeuAc
y5
Pep+HexNAcHex_1+

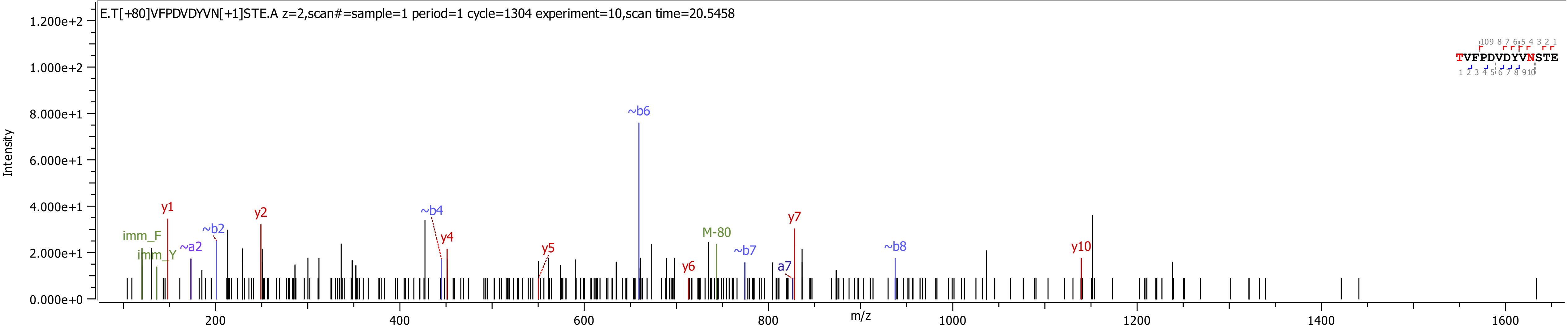

E.T[+80]VFPDVDYVN[+1]STE.A z=2,scan#=sample=1 period=1 cycle=1304 experiment=10,scan time=20.5458
Intensity
m/z
TVFPDVDYVNSTE
imm_F
imm_Y
y1
~a2
~b2
y2
~b4
y4
y5
~b6
y6
M-80
~b7
a7
y7
~b8
y10

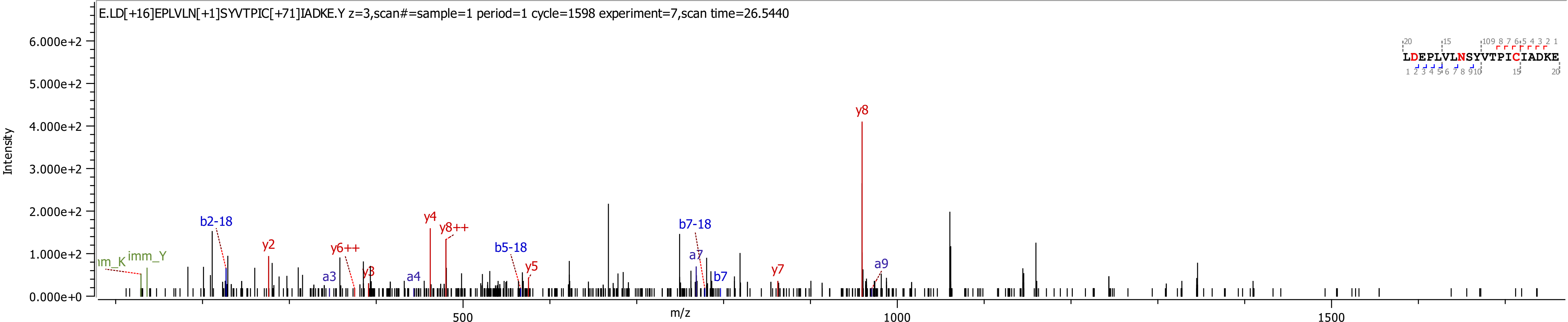
E.LD[+16]EPLVLN[+1]SYVTPIC[+71]IADKE.Y z=3,scan#=sample=1 period=1 cycle=1598 experiment=7,scan time=26.5440
Intensity
m/z
6.000e+2
5.000e+2
4.000e+2
3.000e+2
2.000e+2
1.000e+2
0.000e+0
500
1000
1500
LDEPLVLNSYVTPICIADKE
y8
b2-18
y2
y6++
y3
a3
a4
y4
y8++
b5-18
y5
b7-18
a7
b7
y7
a9
imm_Y
m_K

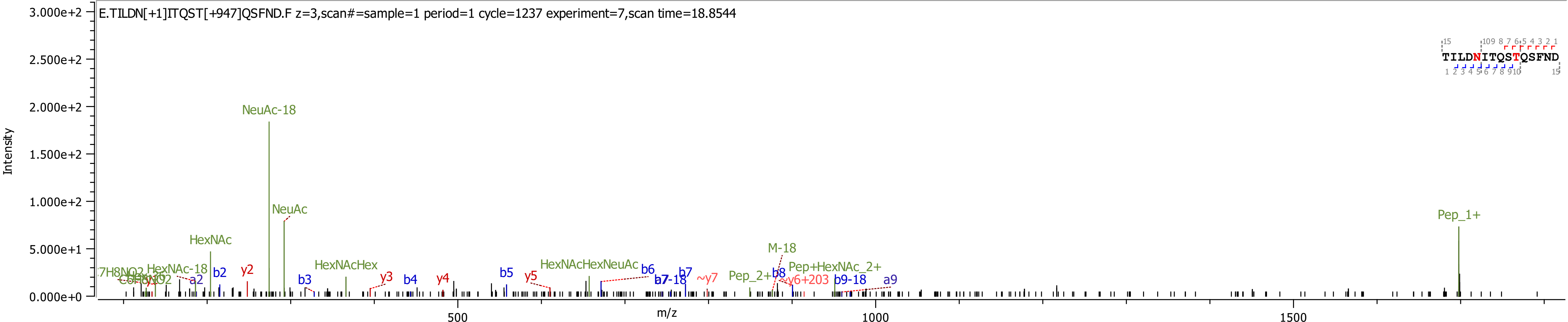

E.TILDN[+1]ITQST[+947]QSFND.F z=3,scan#=sample=1 period=1 cycle=1237 experiment=7,scan time=18.8544
TILDNITQSTQSFND
Intensity
m/z
3.000e+2
2.500e+2
2.000e+2
1.500e+2
1.000e+2
5.000e+1
0.000e+0
500
1000
1500
NeuAc-18
NeuAc
HexNAc
HexNAc-18
HexNAcHex
HexNAcHexNeuAc
Pep_1+
M-18
Pep_2+
Pep+HexNAc_2+
a2
b2
y2
b3
y3
b4
y4
b5
y5
b6
b7
b7-18
~y7
b8
y6+203
b9-18
a9

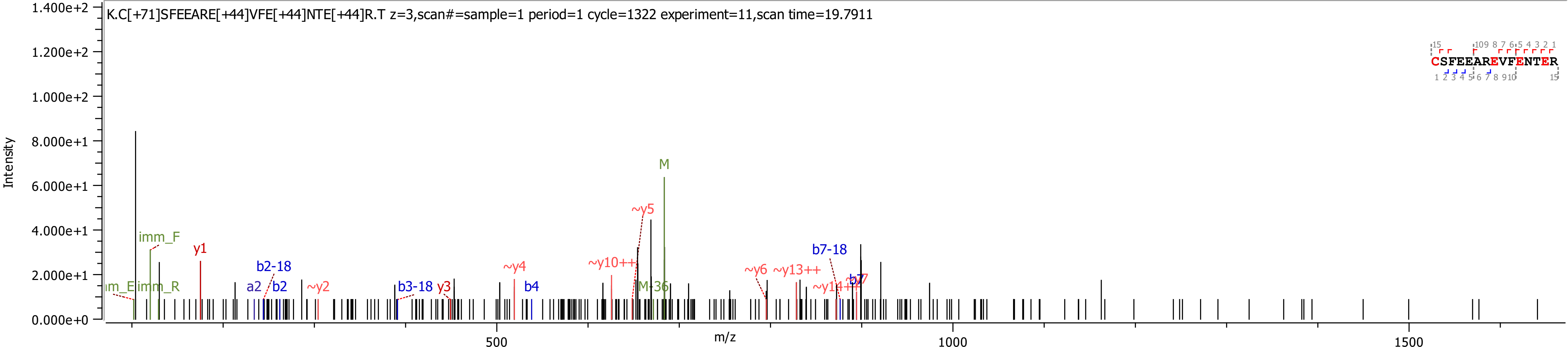
K.C[+71]SFEEARE[+44]VFE[+44]NTE[+44]R.T z=3,scan#=sample=1 period=1 cycle=1322 experiment=11,scan time=19.7911
Intensity
m/z
CSFEEAREVFENTER
M
~y5
imm_F
y1
b2-18
a2
b2
~y2
b3-18
y3
~y4
b4
~y10++
M-36
~y6
~y13++
b7-18
~y14++
b7

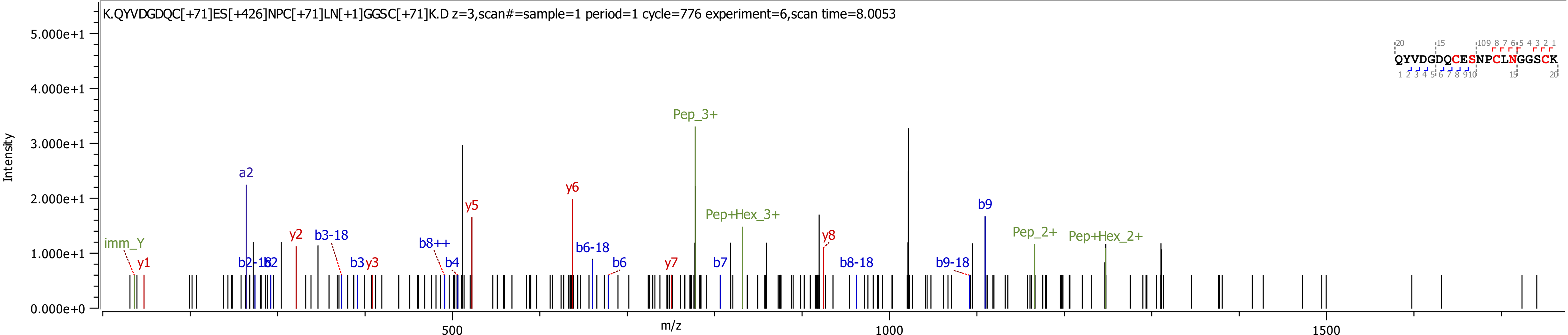

K.QYVDGDQC[+71]ES[+426]NPC[+71]LN[+1]GGSC[+71]K.D z=3,scan#=sample=1 period=1 cycle=776 experiment=6,scan time=8.0053
Intensity
m/z
QYVDGDQCESNPCLNGGSCK
imm_Y
y1
a2
b2-18
b2
y2
b3-18
b3
y3
b8++
b4
y5
y6
b6-18
b6
y7
Pep_3+
b7
Pep+Hex_3+
y8
b8-18
b9-18
b9
Pep_2+
Pep+Hex_2+

ProteinPilot™ Software - [Result - D:\2018\20181030_Schulz_Luci_FIXH1H2\20181106_Schulz_Luci_H2_TTP.group]

File Configure Window Help

**Workflow Tasks**

**Identify Proteins**
- LC...
- Spot-Based (MS only)...
- Spot-Based (MS and MS/MS)...

**View**
- Analysis Log...
- Result...

**Export**
- Peptide Summary...
- Distinct Peptide Summary...
- Protein Summary...
- Spectrum Summary...
- MGF Peaklist(s)...
- mzIdentML...
- Features...

Protein ID | Features | Spectra | Summary Statistics

### Spectrum List

| Spectrum | Acq Time | Obs MW | Obs m/z | Obs z | Prot N | Best Sequence | Modifications | Conf | Theor MW | z |
|---|---|---|---|---|---|---|---|---|---|---|
| 3.1.1.919.3 | 11.303 | 854.3748 | 428.1947 | 2 | 1, 3 | TTEFWK | Carboxy(E)@3 | 60.8 | 854.3810 | 2 |

### Peptide ID Hypotheses - 3.1.1.919.3

| Conf | Sc | Prot N | Sequence | Modifications | Theor MW | Theor m/z | Obs m/z | z | ΔMass |
|---|---|---|---|---|---|---|---|---|---|
| 60.8 | 7 | 1, 3 | TTEFWK | Carboxy(E)@3 | 854.3810 | 428.1978 | 428.1947 | 2 | -0.0063 |

### Precursor MS Region

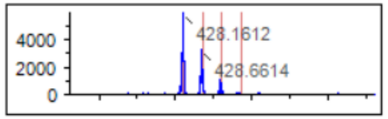


### Fragmentation Evidence for Peptide

TTE[Cox]FWK

| Residue | b | b+2 | y | y+2 |
|---|---|---|---|---|
| T | 102.0550 | 51.5311 | 855.3883 | 428.1 |
| T | 203.1026 | 102.0550 | 754.3406 | 377.6 |
| E[Cox] | 376.1351 | 188.5712 | 653.2930 | 327.1 |
| F | 523.2035 | 262.1054 | 480.2605 | 240.6 |
| W | 709.2828 | 355.1450 | 333.1921 | 167.0 |
| K | 837.3777 | 419.1925 | 147.1128 | 74.0 |

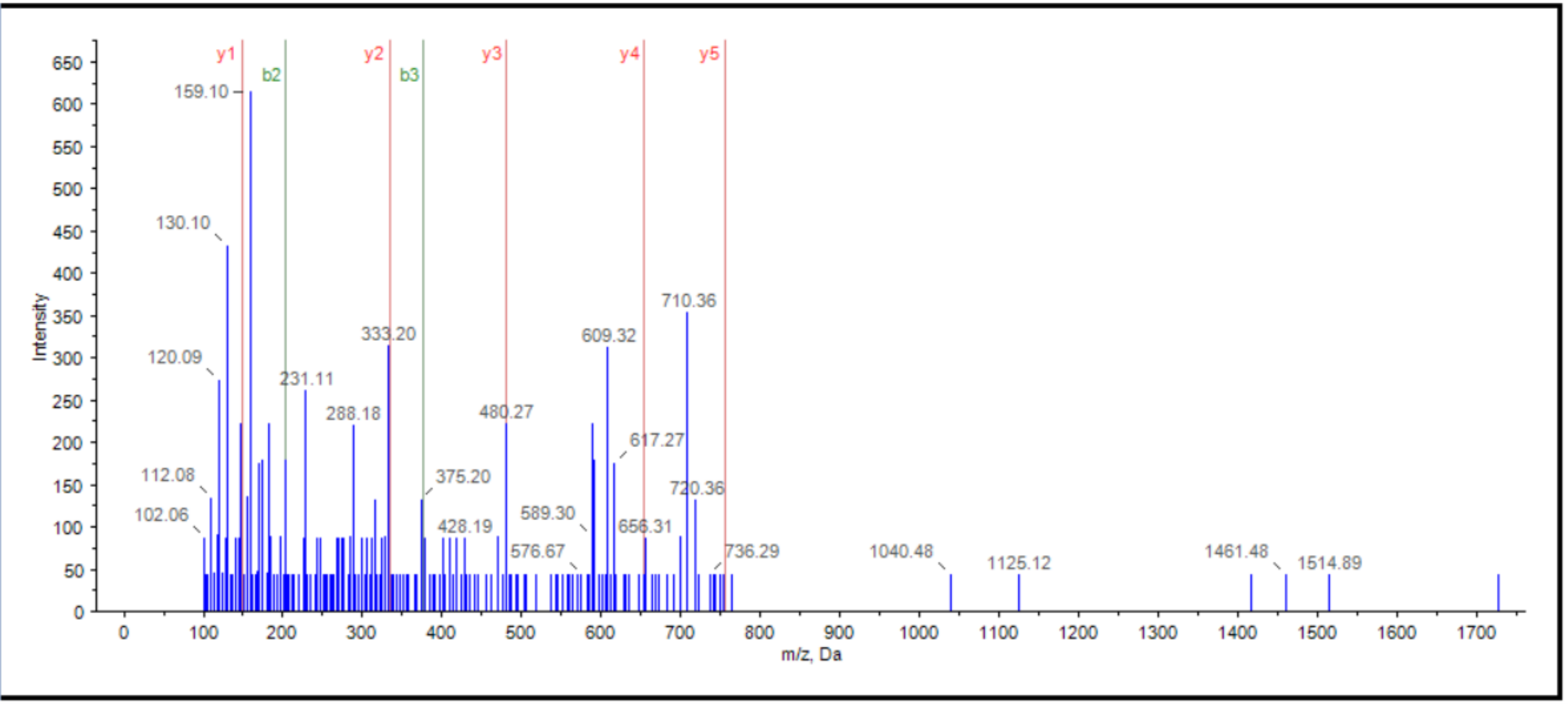


Type here to search

ENG 11:11 PM 2/04/2020
