## Supplementary Information 4 for "Analysis of coagulation factor IX in bioreactor cell culture medium predicts yield and quality of the purified product"

20190702_Co4Ra2NeuGcOglyLacNAc_20190509_BenSchulz_Luci_pdFIXa_Ch.wiff_Byonic_S53[+294]_S61[+802] 1

20190702_Co4Ra2NeuGcOglyLacNAc_20190509_BenSchulz_Luci_pdFIXa_Ch.wiff_Byonic_S53[+426]_S61[+802] 2

Common4rare2NeuGc50_20181106_Schulz_Luci_FIXaG.wiff_20181116_Byonic_D104 3

Common4rare2NeuGc50_20181106_Schulz_Luci_FIXaG.wiff_20181116_Byonic_D104[+16] 4

Common4rare2NeuGc50_20181106_Schulz_Luci_FIXaG.wiff_20181116_Byonic_D276[+16] 5

Common4rare2NeuGc50_20181106_Schulz_Luci_FIXaG.wiff_20181116_Byonic_N258 6
Common4rare2NeuGc50_20181106_Schulz_Luci_FIXaG.wiff_20181116_Byonic_S53[+294]_S61[+802] 7
Common4rare2NeuGc50_20181106_Schulz_Luci_FIXaG.wiff_20181116_Byonic_S141 8
Common4rare2NeuGc50_20181106_Schulz_Luci_FIXaG.wiff_20181116_Byonic_Y45 9
Common4rare2NeuGc50_20181106_Schulz_Luci_FIXaGP.wiff_20181116_Byonic T159[+947] 10

Common4rare2NeuGc50_20181106_Schulz_Luci_FIXaGP.wiff_20181116_Byonic_N157[+1] 11
Common4rare2NeuGc50_20181106_Schulz_Luci_FIXaGP.wiff_20181116_Byonic_N167[+1] 12
Common4rare2NeuGc50_20181106_Schulz_Luci_FIXaGP.wiff_20181116_Byonic_T169orT172[+656] 13
Common4rare2NeuGc50_20181106_Schulz_Luci_FIXaGP.wiff_20181116_Byonic_Y155[+80] 14
Common4rare2NeuGc50_20181106_Schulz_Luci_FIXaT.wiff_20181119_Byonic_D49[+16]_S53[+294]_S61[+802] 15

Common4rare2NeuGc50_20181106_Schulz_Luci_FIXaT.wiff_20181119_Byonic_D64_S68 16
Common4rare2NeuGc50_20181106_Schulz_Luci_FIXaT.wiff_20181119_Byonic_D64[+16] 17
Common4rare2NeuGc50_20181106_Schulz_Luci_FIXaT.wiff_20181119_Byonic_D85 18
Common4rare2NeuGc50_20181106_Schulz_Luci_FIXaT.wiff_20181119_Byonic_D85[+16] 19
Common4rare2NeuGc50_20181106_Schulz_Luci_FIXaT.wiff_20181119_Byonic_D186 20

Common4rare2NeuGc50_20181106_Schulz_Luci_FIXaT.wiff_20181119_Byonic_D186[+16] 21
Common4rare2NeuGc50_20181106_Schulz_Luci_FIXaT.wiff_20181119_Byonic_D203 22
Common4rare2NeuGc50_20181106_Schulz_Luci_FIXaT.wiff_20181119_Byonic_D203[+16] 23
Common4rare2NeuGc50_20181106_Schulz_Luci_FIXaT.wiff_20181119_Byonic_D276_D292 24
Common4rare2NeuGc50_20181106_Schulz_Luci_FIXaT.wiff_20181119_Byonic_D292[+16] 25

Common4rare2NeuGc50_20181106_Schulz_Luci_FIXaT.wiff_20181119_Byonic_D358_D364 26
Common4rare2NeuGc50_20181106_Schulz_Luci_FIXaT.wiff_20181119_Byonic_D358[+16] 27
Common4rare2NeuGc50_20181106_Schulz_Luci_FIXaT.wiff_20181119_Byonic_D364[+16] 28
Common4rare2NeuGc50_20181106_Schulz_Luci_FIXaT.wiff_20181119_Byonic_N258 29
Common4rare2NeuGc50_20181106_Schulz_Luci_FIXaT.wiff_20181119_Byonic_N258[+2204] 30

Common4rare2NeuGc50_20181106_Schulz_Luci_FIXaT.wiff_20181119_Byonic_S53[+294]_S61[+802] 31
Common4rare2NeuGc50_20181106_Schulz_Luci_FIXaT.wiff_20181119_Byonic_S68[+80] 32
Common4rare2NeuGc50_20181106_Schulz_Luci_FIXaT.wiff_20181119_Byonic_S141[+947] 33
Common4rare2NeuGc50_20181106_Schulz_Luci_FIXaTP.wiff_20181119_Byonic_E40[+44] 34

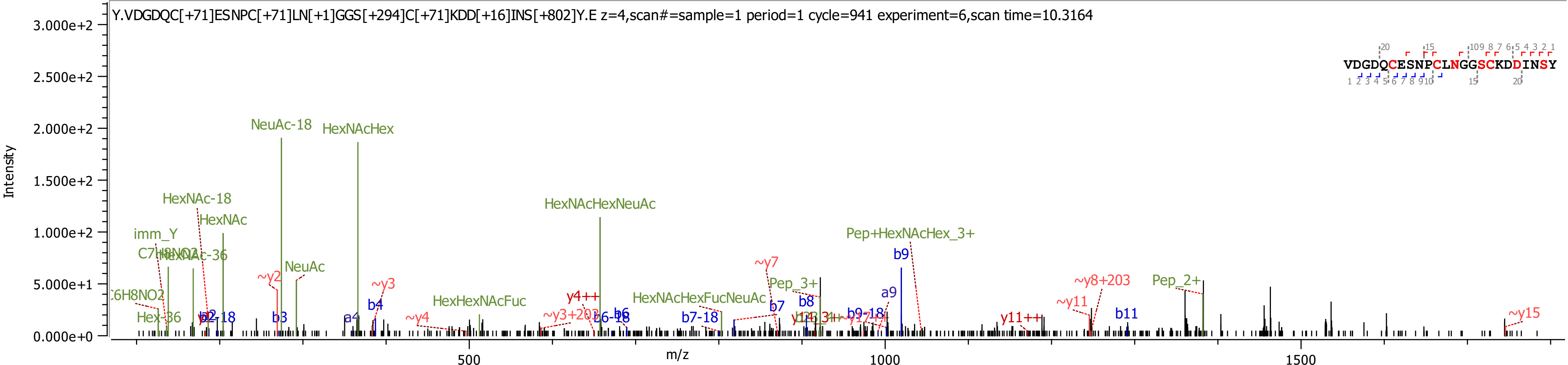

Y.VDGDQC[+71]ESNPC[+71]LN[+1]GGS[+294]C[+71]KDD[+16]INS[+802]Y.E z=4,scan#=sample=1 period=1 cycle=941 experiment=6,scan time=10.3164
VDGDQCESNPCLNGGSCKDDINSY
Intensity
m/z
3.000e+2
2.500e+2
2.000e+2
1.500e+2
1.000e+2
5.000e+1
0.000e+0
500
1000
1500
NeuAc-18
HexNAcHex
HexNAc-18
HexNAc
imm_Y
HexNAcHexNeuAc
Pep+HexNAcHex_3+
NeuAc
~y2
~y3
~y4
~y7
b9
a9
b4
b3
HexHexNAcFuc
HexNAcHexFucNeuAc
Pep_3+
Pep_2+
b8
b7
b7-18
b6
b6-18
y4++
~y3+203
~y8+203
~y11
y11++
b11
~y15
b9-18
Hex-36

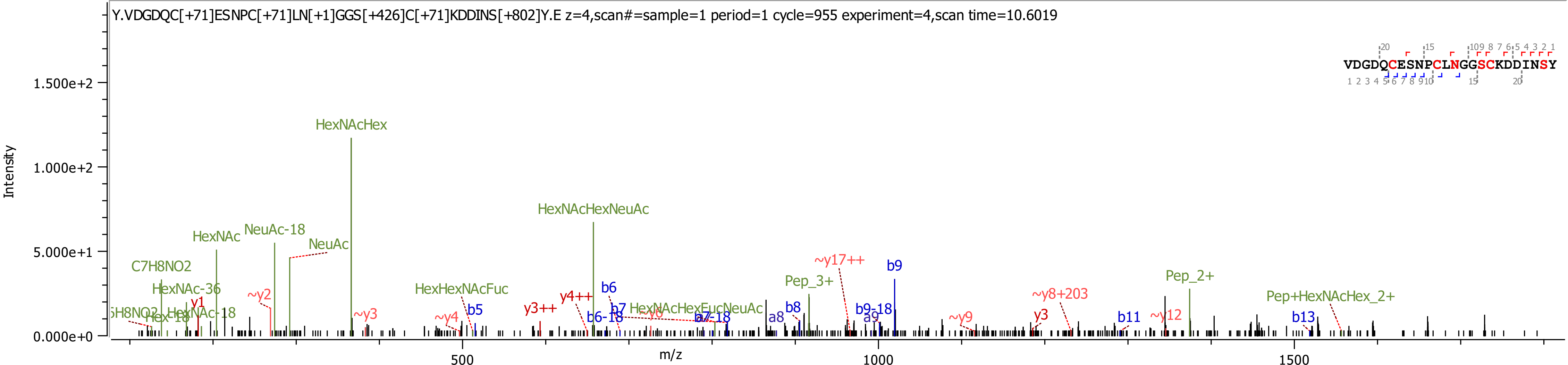
Y.VDGDQC[+71]ESNPC[+71]LN[+1]GGS[+426]C[+71]KDDINS[+802]Y.E z=4,scan#=sample=1 period=1 cycle=955 experiment=4,scan time=10.6019
VDGDQCESNPCLNGGSCKDDINSY
Intensity
1.500e+2
1.000e+2
5.000e+1
0.000e+0
m/z
500
1000
1500
HexNAcHex
HexNAcHexNeuAc
HexNAc
NeuAc-18
NeuAc
C7H8NO2
HexNAc-36
y1
~y2
~y3
~y4
HexHexNAcFuc
b5
y3++
y4++
b6
b6-18
b7
HexNAcHexFucNeuAc
~y6
b7-18
a8
b8
Pep_3+
~y17++
b9
b9-18
a9
~y9
~y8+203
y3
b11
~y12
Pep_2+
Pep+HexNAcHex_2+
b13
Hex-18
HexNAc-18

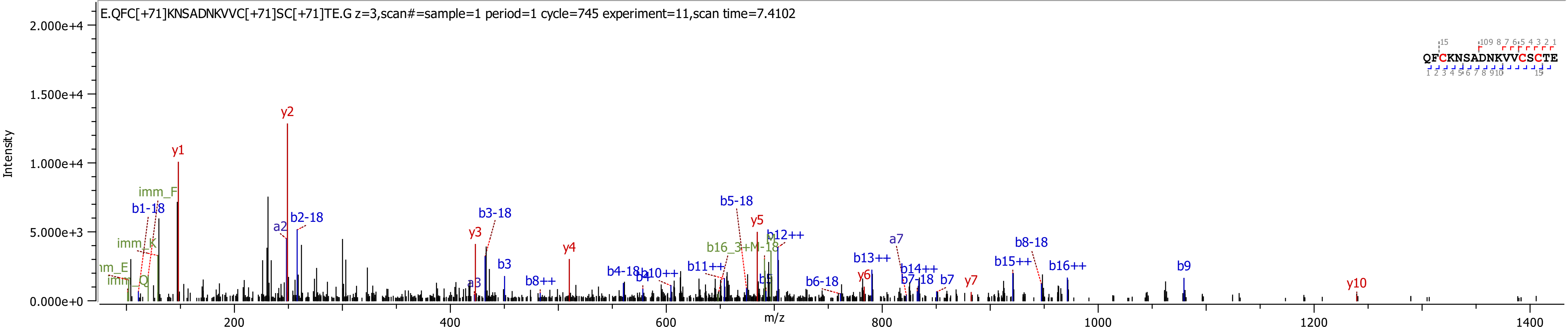

E.QFC[+71]KNSADNKVVC[+71]SC[+71]TE.G z=3,scan#=sample=1 period=1 cycle=745 experiment=11,scan time=7.4102
QFCKNSADNKVVCSCTE
Intensity
m/z
2.000e+4
1.500e+4
1.000e+4
5.000e+3
0.000e+0
200
400
600
800
1000
1200
1400
y1
y2
y3
y4
y5
y6
y7
y10
imm_F
imm_K
imm_Q
b1-18
a2
b2-18
b3-18
b3
a3
b8++
b4-18
b4
b10++
b11++
b5-18
b16_3+M-18
b5
b12++
b6-18
b13++
a7
b14++
b7-18
b7
b15++
b8-18
b16++
b9

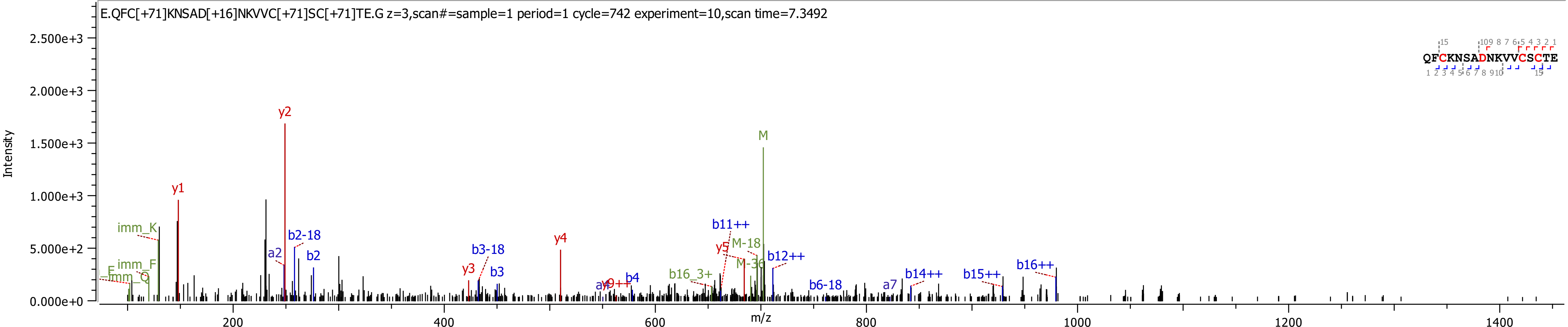
E.QFC[+71]KNSAD[+16]NKVVC[+71]SC[+71]TE.G z=3,scan#=sample=1 period=1 cycle=742 experiment=10,scan time=7.3492
Intensity
m/z
QFCKNSADNKVVCSCTE
y1
y2
y3
y4
y5
M
M-18
M-36
imm_K
imm_F
imm_Q
a2
a7
b2
b2-18
b3
b3-18
b4
b6-18
b11++
b12++
b14++
b15++
b16++
b16_3+

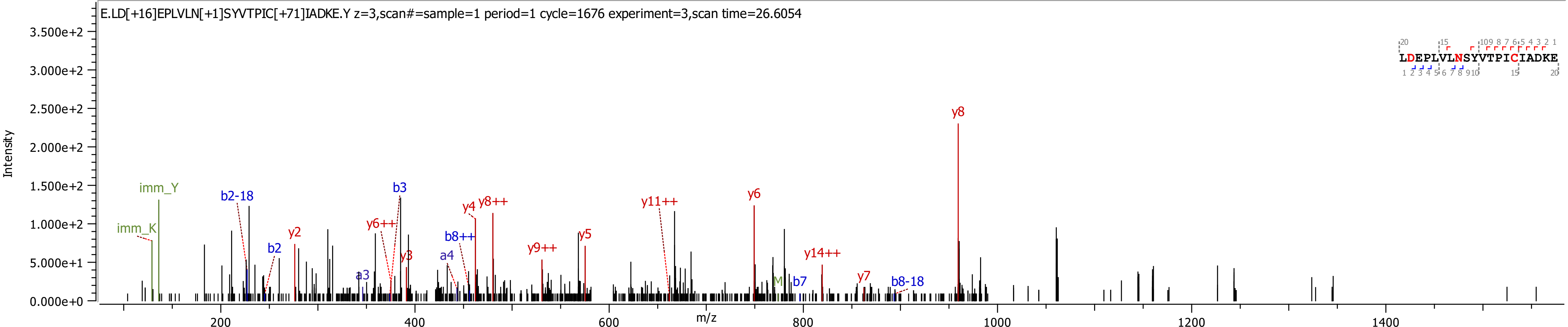
E.LD[+16]EPLVLN[+1]SYVTPIC[+71]IADKE.Y z=3,scan#=sample=1 period=1 cycle=1676 experiment=3,scan time=26.6054
Intensity
m/z
LDEPLVLNSYVTPICIADKE
imm_Y
imm_K
b2-18
b2
y2
b3
y6++
y3
a3
a4
y4
y8++
b8++
y9++
y5
y11++
y6
b7
y14++
y7
b8-18
y8

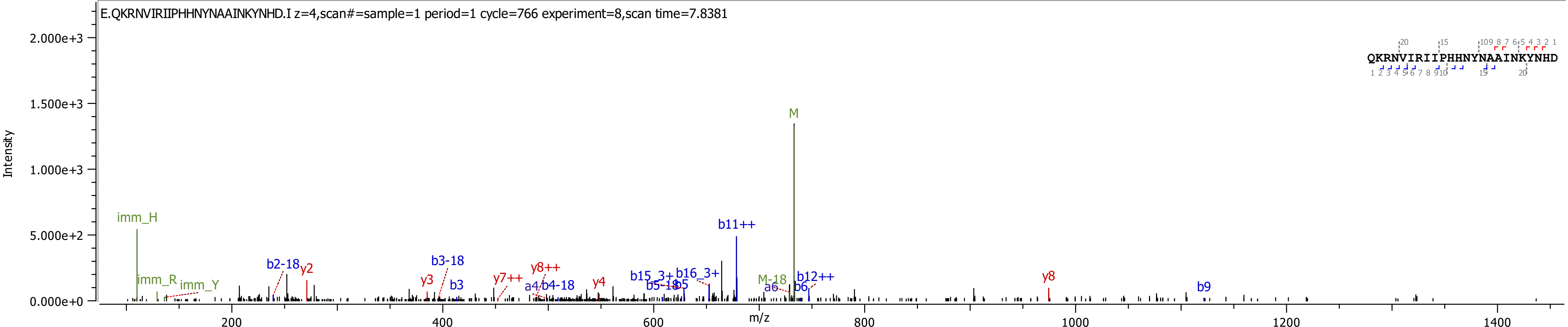

E.QKRNVIRIIPHHNYNAAINKYNHD.I z=4,scan#=sample=1 period=1 cycle=766 experiment=8,scan time=7.8381
QKRNVIRIIPHHNYNAAINKYNHD
Intensity
m/z
2.000e+3
1.500e+3
1.000e+3
5.000e+2
0.000e+0
200
400
600
800
1000
1200
1400
M
imm_H
imm_R
imm_Y
b2-18
y2
y3
b3-18
b3
y7++
y8++
a4
b4-18
y4
b15_3+
b5-18
b5
b16_3+
b11++
M-18
a6
b6
b12++
y8
b9

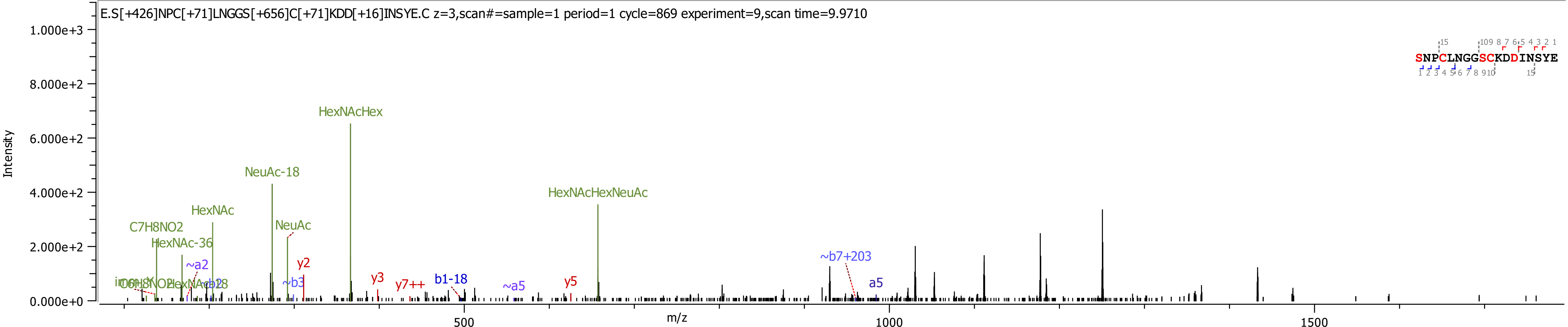
E.S[+426]NPC[+71]LNGGS[+656]C[+71]KDD[+16]INSYE.C z=3,scan#=sample=1 period=1 cycle=869 experiment=9,scan time=9.9710
SNPCLNGGSCKDDINSYE
Intensity
m/z
HexNAcHex
NeuAc-18
HexNAcHexNeuAc
HexNAc
C7H8NO2
HexNAc-36
NeuAc
~a2
y2
~b3
y3
y7++
b1-18
~a5
y5
~b7+203
a5
0.000e+0
2.000e+2
4.000e+2
6.000e+2
8.000e+2
1.000e+3
500
1000
1500

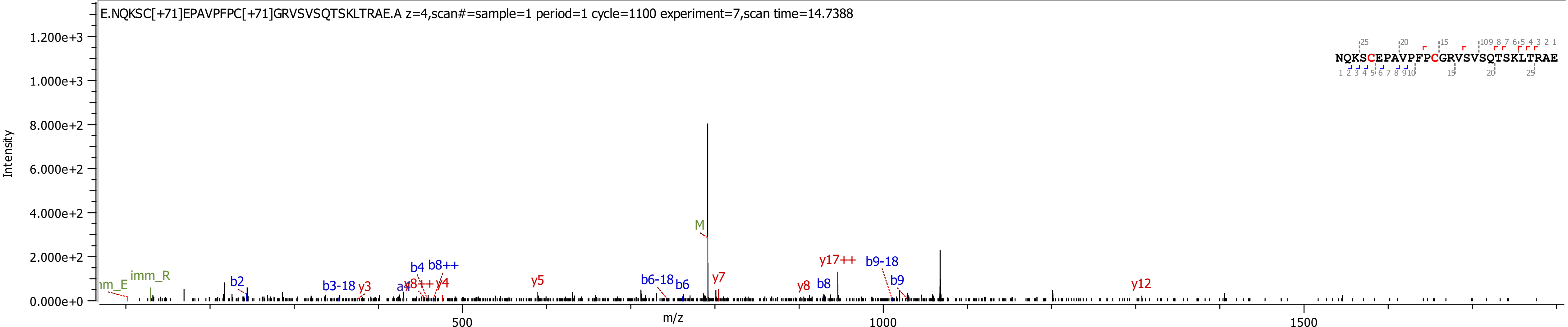

E.NQKSC[+71]EPAVPFPC[+71]GRVSVSQTSKLTRAE.A z=4,scan#=sample=1 period=1 cycle=1100 experiment=7,scan time=14.7388
NQKSCEPAVPFPCGRVSVSQTSKLTRAE
Intensity
m/z
0.000e+0
2.000e+2
4.000e+2
6.000e+2
8.000e+2
1.000e+3
1.200e+3
500
1000
1500
imm_R
b2
b3-18
y3
b4
b8++
y4
y5
b6-18
b6
M
y7
y8
b8
y17++
b9-18
b9
y12

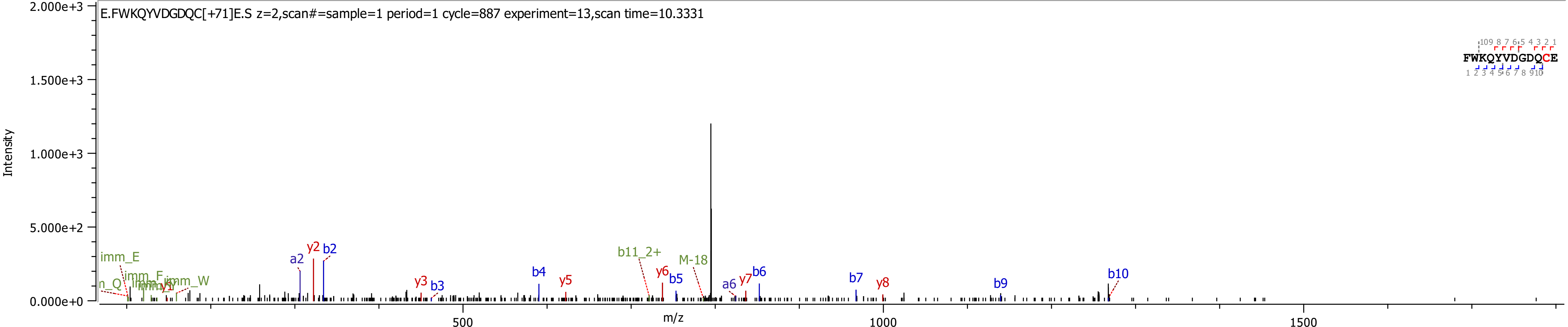
E.FWKQYVDGDQC[+71]E.S z=2,scan#=sample=1 period=1 cycle=887 experiment=13,scan time=10.3331
FWKQYVDGDQCE
Intensity
2.000e+3
1.500e+3
1.000e+3
5.000e+2
0.000e+0
m/z
500
1000
1500
imm_E
n_Q
imm_F
imm_W
a2
y2
b2
y3
b3
b4
y5
b11_2+
y6
b5
M-18
a6
y7
b6
b7
y8
b9
b10

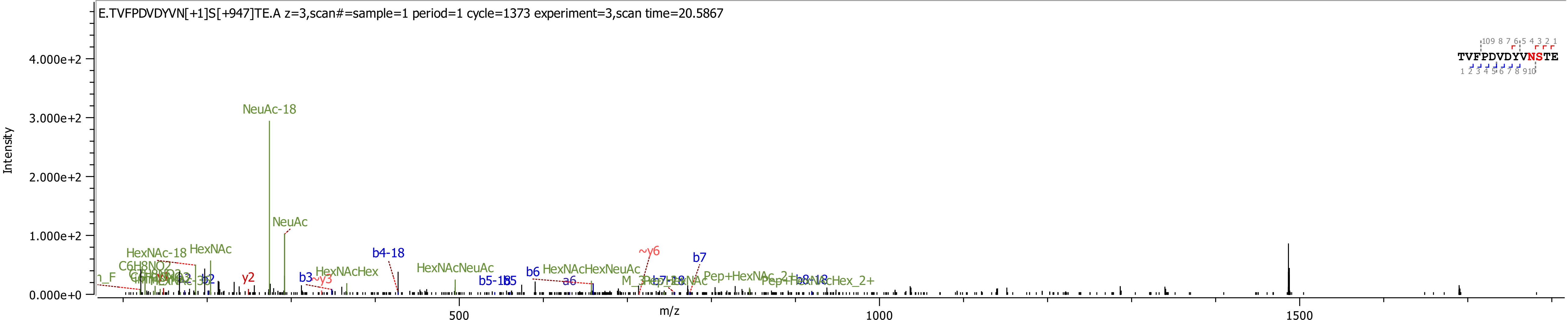

E.TVFPDVDYVN[+1]S[+947]TE.A z=3,scan#=sample=1 period=1 cycle=1373 experiment=3,scan time=20.5867
TVFPDVDYVNSTE
Intensity
4.000e+2
3.000e+2
2.000e+2
1.000e+2
0.000e+0
m/z
500
1000
1500
NeuAc-18
NeuAc
HexNAc-18
HexNAc
C6H8NO2
y2
b3
~y3
HexNAcHex
b4-18
HexNAcNeuAc
b5-18
b5
b6
HexNAcHexNeuAc
a6
~y6
b7
Pep+HexNAc_2+

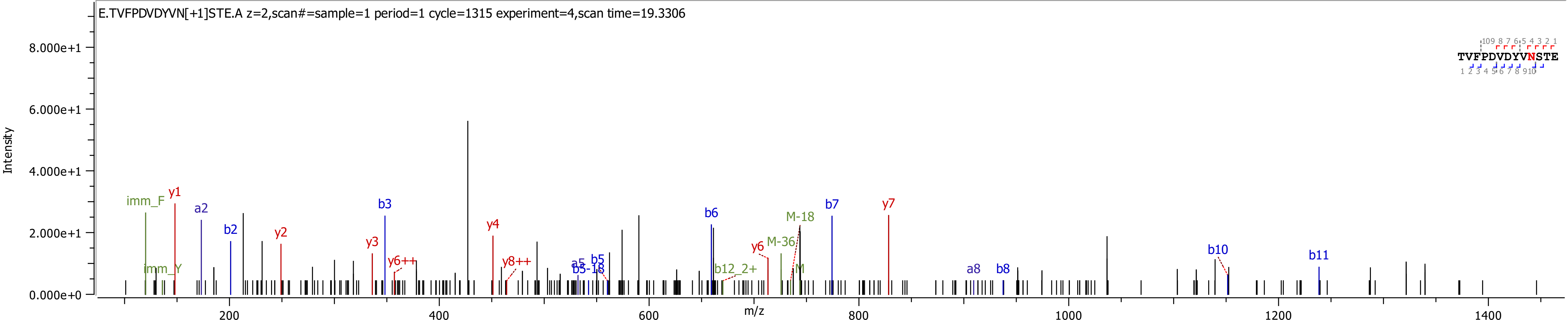

E.TVFPDVDYVN[+1]STE.A z=2,scan#=sample=1 period=1 cycle=1315 experiment=4,scan time=19.3306
Intensity
m/z
8.000e+1
6.000e+1
4.000e+1
2.000e+1
0.000e+0
200
400
600
800
1000
1200
1400
imm_F
imm_Y
y1
a2
b2
y2
y3
b3
y6++
y4
y8++
a5
b5
b5-18
b6
b12_2+
y6
M-36
M-18
M
b7
y7
a8
b8
b10
b11
TVFPDVDYVNSTE

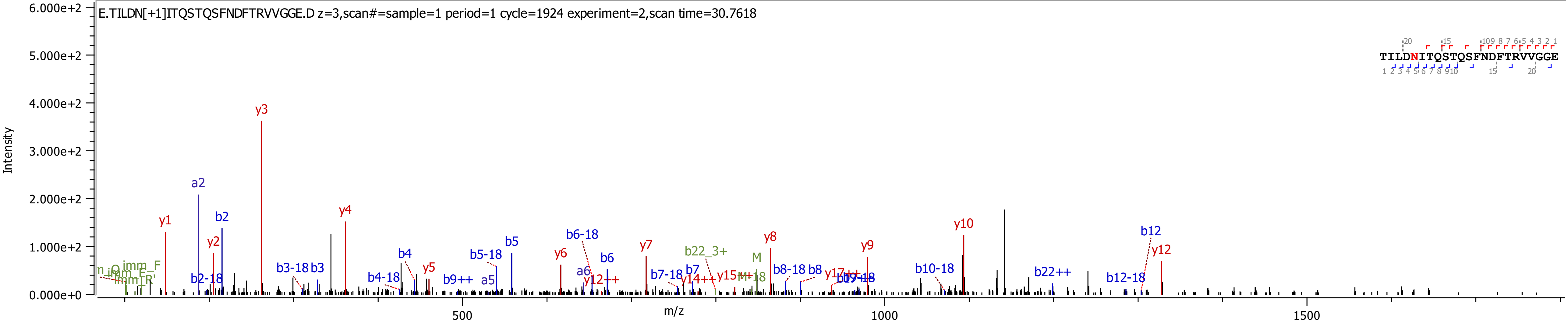
E.TILDN[+1]ITQSTQSFNDFTRVVGGE.D z=3,scan#=sample=1 period=1 cycle=1924 experiment=2,scan time=30.7618
TILDNITQSTQSFNDFTRVVGGE
Intensity
m/z
6.000e+2
5.000e+2
4.000e+2
3.000e+2
2.000e+2
1.000e+2
0.000e+0
500
1000
1500
y1
a2
b2
y2
y3
b2-18
b3-18
b3
y4
b4-18
b4
y5
b9++
a5
b5-18
b5
b6-18
y6
a6
b6
y12++
y7
b7-18
b7
y14++
b22_3+
y15++
M
M-18
y8
b8-18
b8
y17++
y9
b10-18
y10
b22++
b12
b12-18
y12

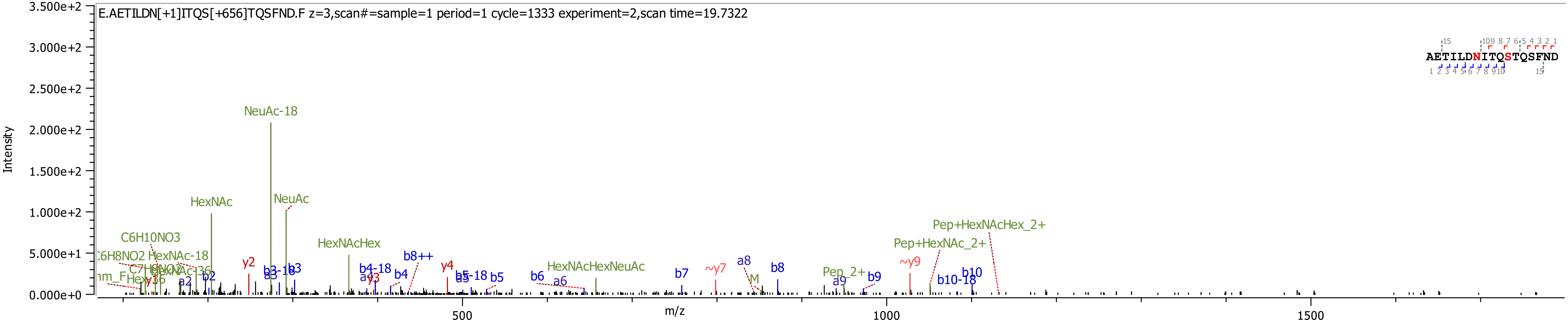

E.AETILDN[+1]ITQS[+656]TQSFND.F z=3,scan#=sample=1 period=1 cycle=1333 experiment=2,scan time=19.7322
AETILDNITQSTQSFND
Intensity
m/z
3.500e+2
3.000e+2
2.500e+2
2.000e+2
1.500e+2
1.000e+2
5.000e+1
0.000e+0
500
1000
1500
NeuAc-18
NeuAc
HexNAc
C6H10NO3
C6H8NO2
HexNAc-18
HexNAc-36
HexNAcHex
HexNAcHexNeuAc
Pep+HexNAcHex_2+
Pep+HexNAc_2+
Pep_2+
M
y2
y4
~y7
~y9
b2
b3
b3-18
b4
b4-18
b5
b5-18
b6
b7
b8
b8++
b9
b10
b10-18
a2
a3
a4
a6
a8
a9

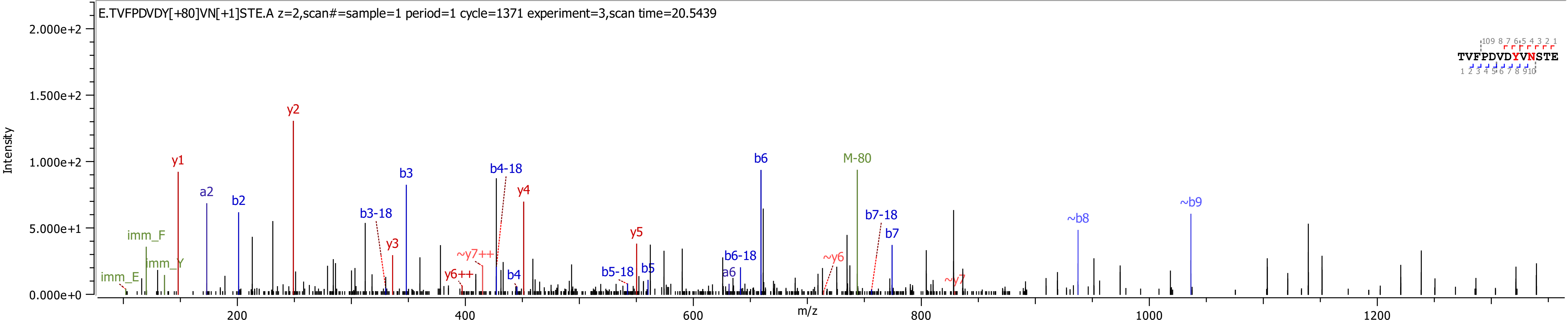
E.TVFPDVDY[+80]VN[+1]STE.A z=2,scan#=sample=1 period=1 cycle=1371 experiment=3,scan time=20.5439
Intensity
2.000e+2
1.500e+2
1.000e+2
5.000e+1
0.000e+0
m/z
200
400
600
800
1000
1200
TVFPDVDYVNSTE
y1
y2
a2
b2
b3
b3-18
y3
b4-18
y4
~y7++
y6++
b4
y5
b5-18
b5
b6
b6-18
a6
~y6
M-80
b7-18
b7
~y7
~b8
~b9
imm_F
imm_Y
imm_E

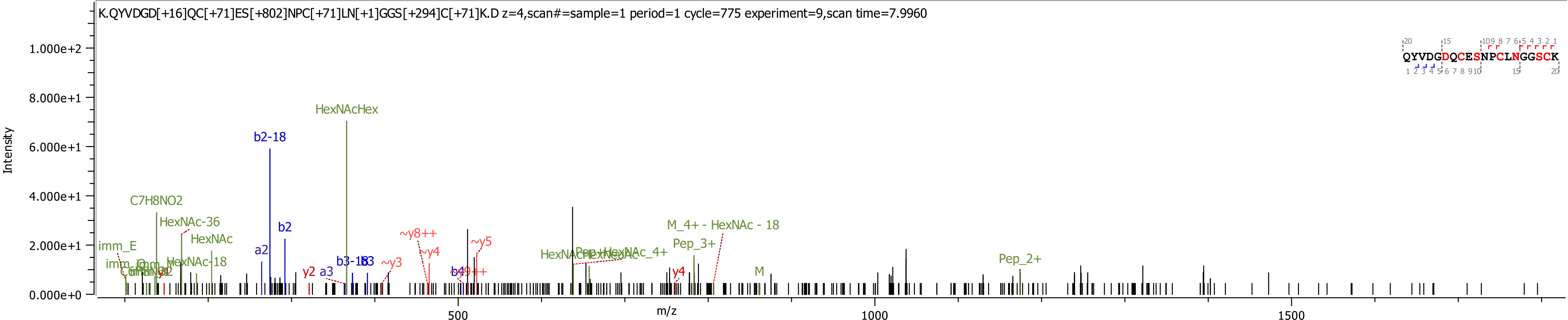
K.QYVDGD[+16]QC[+71]ES[+802]NPC[+71]LN[+1]GGS[+294]C[+71]K.D z=4,scan#=sample=1 period=1 cycle=775 experiment=9,scan time=7.9960
Intensity
m/z
1.000e+2
8.000e+1
6.000e+1
4.000e+1
2.000e+1
0.000e+0
500
1000
1500
QYVDGDQCESNPCLNGGSCK
HexNAcHex
b2-18
C7H8NO2
HexNAc-36
HexNAc
HexNAc-18
imm_E
b2
a2
y2
a3
b3-18
b3
~y3
~y8++
~y4
~y5
b4
b9++
M_4+ - HexNAc - 18
Pep_3+
y4
M
Pep_2+

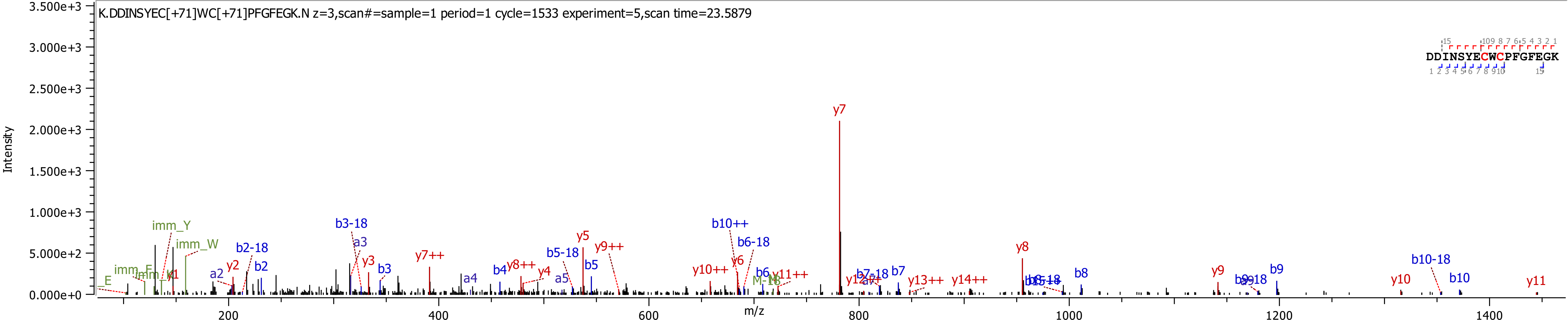
K.DDINSYEC[+71]WC[+71]PFGFEGK.N z=3,scan#=sample=1 period=1 cycle=1533 experiment=5,scan time=23.5879
DDINSYECWCPFGFEGK
Intensity
m/z
y7
y8
y9
y10
y11
b10-18
b10
b9
b8
b10++
b6-18
b3-18
y5
y9++
b5-18
b5
y7++
imm_Y
imm_W
b2-18
b2
a2
y2
a3
y3
b3
y10++
y6
y11++
y13++
y14++

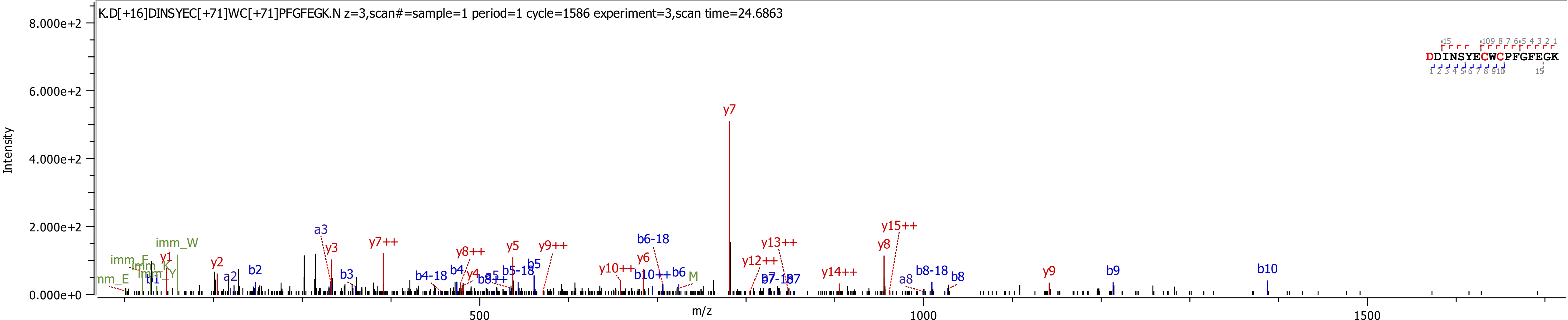
K.D[+16]DINSYEC[+71]WC[+71]PFGFEGK.N z=3,scan#=sample=1 period=1 cycle=1586 experiment=3,scan time=24.6863
DDINSYECWCPFGFEGK
Intensity
m/z
8.000e+2
6.000e+2
4.000e+2
2.000e+2
0.000e+0
500
1000
1500
y7
y15++
y8
y13++
y12++
y14++
y9
b9
b10
b8-18
b8
a8
b6-18
y6
y10++
b10++
b6
y9++
y5
b5
b5-18
y8++
b4
y4
b4-18
y7++
a3
y3
b3
y2
a2
b2
y1
imm_W
imm_F
imm_K
imm_Y
imm_E
M

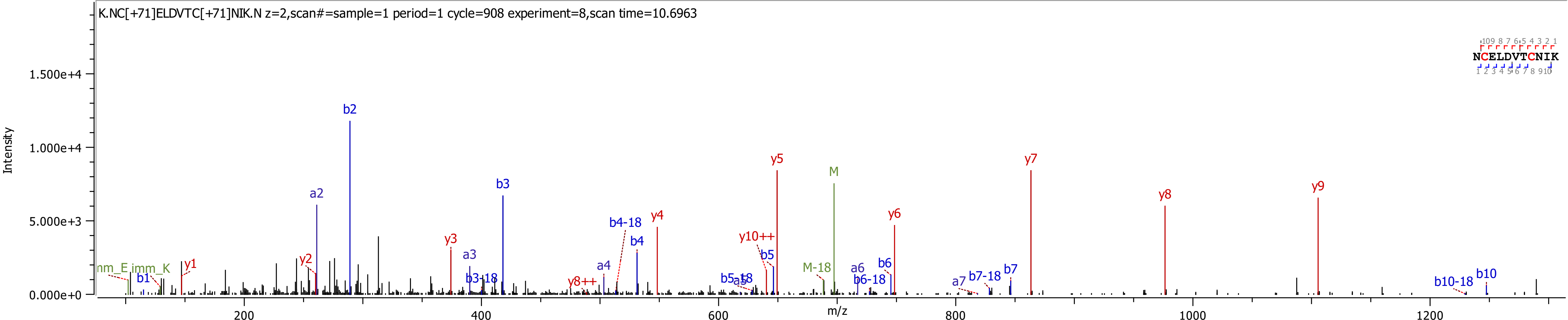

K.NC[+71]ELDVTC[+71]NIK.N z=2,scan#=sample=1 period=1 cycle=908 experiment=8,scan time=10.6963
Intensity
m/z
NCELDVTCNIK
1.500e+4
1.000e+4
5.000e+3
0.000e+0
200
400
600
800
1000
1200
b2
y5
y7
M
a2
b3
y9
y8
y4
b4-18
y6
y3
y10++
b4
a3
b5
a4
b6
a6
b7
M-18
b10
y1
y2
b1
imm_K
nm_E
b3-18
y8++
b5-18
a5
b6-18
a7
b7-18
b10-18

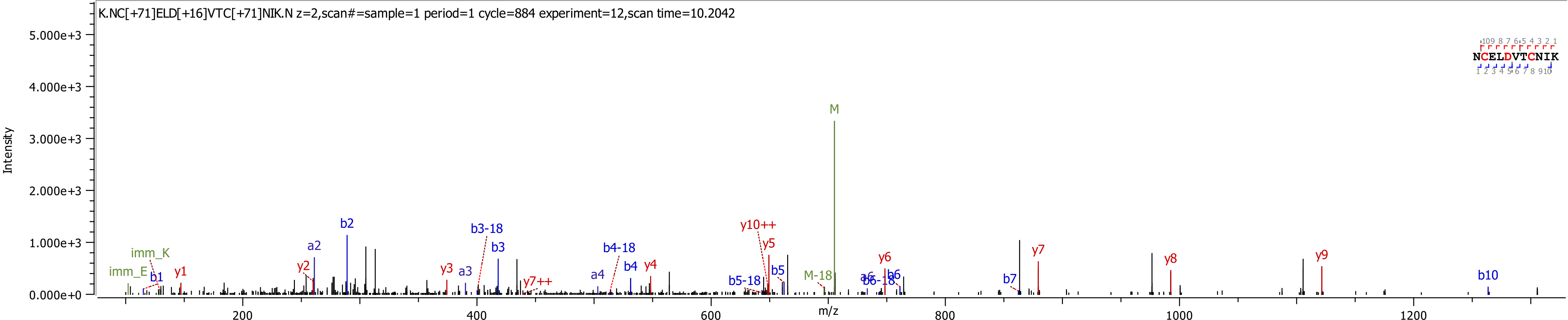
K.NC[+71]ELD[+16]VTC[+71]NIK.N z=2,scan#=sample=1 period=1 cycle=884 experiment=12,scan time=10.2042
Intensity
m/z
NCELDVTCNIK
M
M-18
y10++
y5
b2
b3-18
a2
b3
b4-18
imm_K
y1
y2
y3
a3
a4
b4
y4
imm_E
b1
y7++
b5-18
b5
y6
a6
b6-18
b6
b7
y7
y8
y9
b10
0.000e+0
1.000e+3
2.000e+3
3.000e+3
4.000e+3
5.000e+3
200
400
600
800
1000
1200

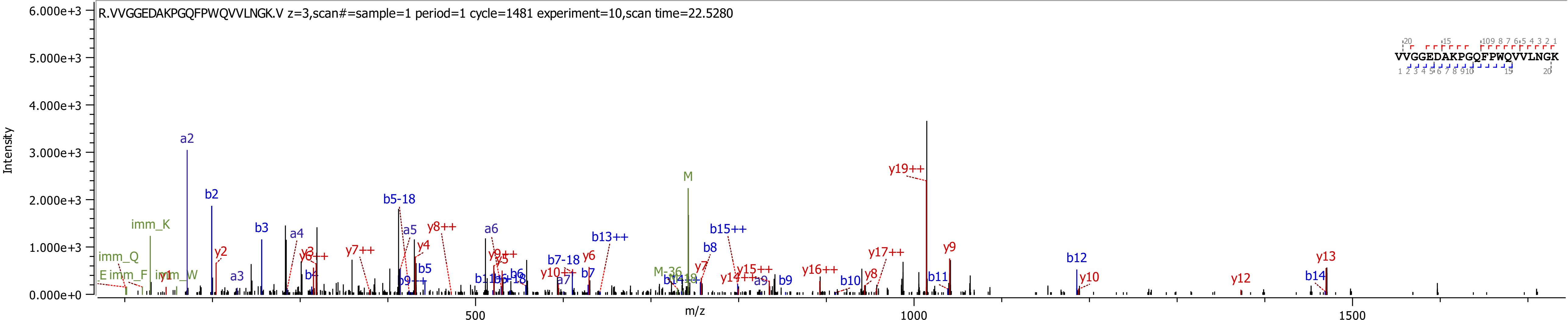
R.VVGGEDAKPGQFPWQVVLNGK.V z=3,scan#=sample=1 period=1 cycle=1481 experiment=10,scan time=22.5280
VVGGEDAKPGQFPWQVVLNGK
Intensity
m/z
6.000e+3
5.000e+3
4.000e+3
3.000e+3
2.000e+3
1.000e+3
0.000e+0
500
1000
1500
a2
b2
b3
a3
a4
b4
b5-18
a5
b5
a6
y19++
M
y9
b12
y13
y10
y12
b14
y8++
y7++
y2
y1
imm_K
imm_Q
imm_F
imm_W
b7-18
b7
y6
b13++
b15++
b8
y7
y15++
y14++
a9
b9
y16++
b10
y8
y17++
b11
M-36
a7
y10++
y4
y5
y9++
b6
b6-18
y3
y6++
b9++

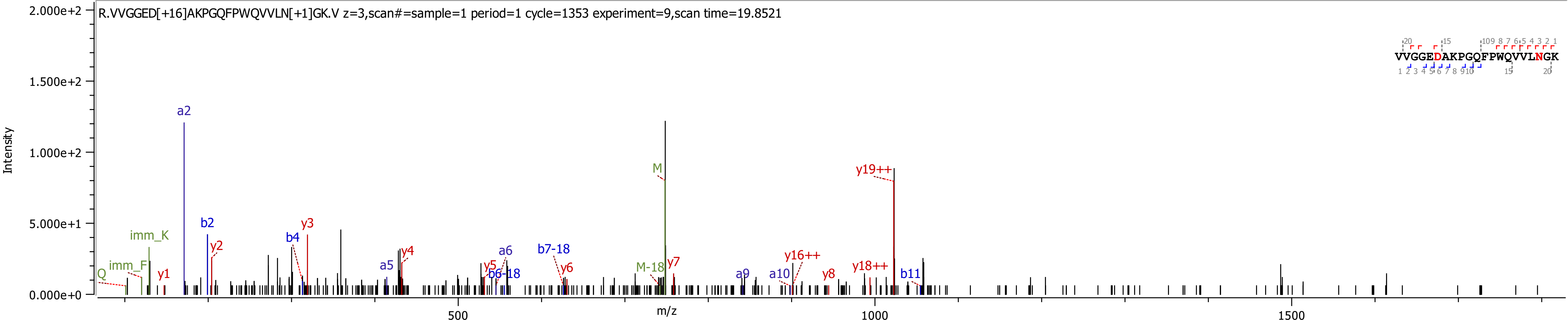
R.VVGGED[+16]AKPGQFPWQVVLN[+1]GK.V z=3,scan#=sample=1 period=1 cycle=1353 experiment=9,scan time=19.8521
VVGGEDAKPGQFPWQVVLNGK
Intensity
m/z
2.000e+2
1.500e+2
1.000e+2
5.000e+1
0.000e+0
500
1000
1500
a2
b2
y2
y3
b4
imm_K
imm_F
Q
y1
a5
y4
a6
b7-18
y5
b6-18
y6
M
M-18
y7
a9
a10
y16++
y8
y18++
y19++
b11

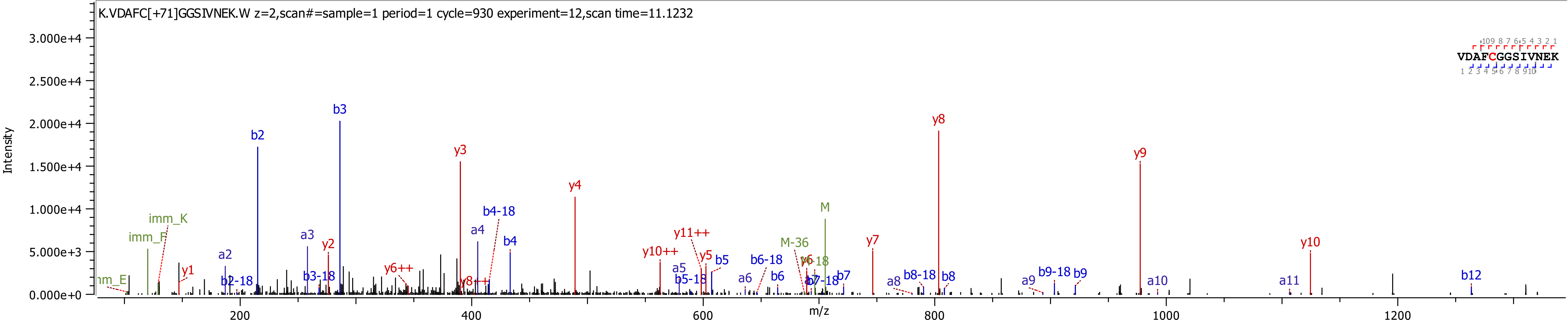
K.VDAFC[+71]GGSIVNEK.W z=2,scan#=sample=1 period=1 cycle=930 experiment=12,scan time=11.1232
VDAFCGGSIVNEK
Intensity
m/z
3.000e+4
2.500e+4
2.000e+4
1.500e+4
1.000e+4
5.000e+3
0.000e+0
200
400
600
800
1000
1200
b3
b2
y8
y9
y3
y4
M
imm_K
imm_F
b4-18
a4
b4
a3
a2
y2
y1
b2-18
b3-18
y6++
y8++
y10++
y11++
y5
b5
a5
b5-18
a6
b6-18
b6
M-36
y6
b7-18
b7
y7
a8
b8-18
b8
a9
b9-18
b9
a10
a11
y10
b12

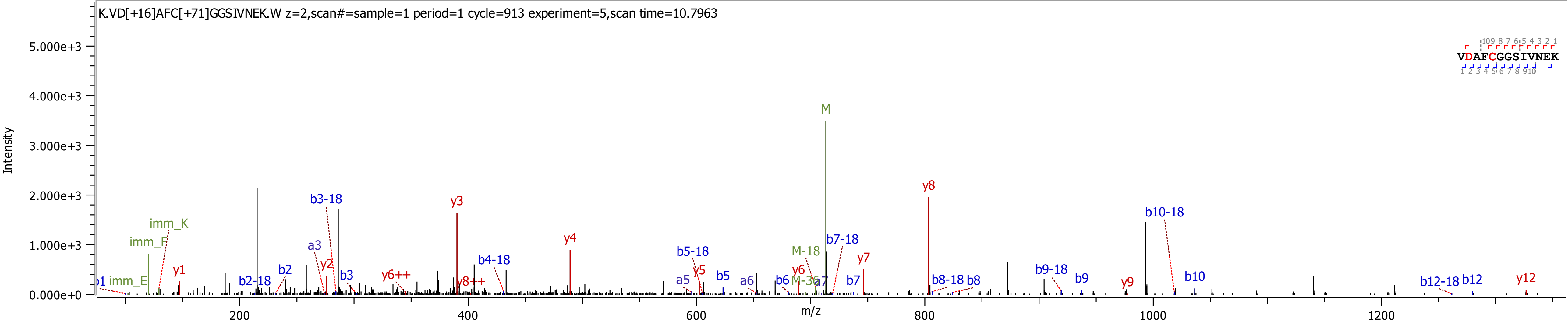
K.VD[+16]AFC[+71]GGSIVNEK.W z=2,scan#=sample=1 period=1 cycle=913 experiment=5,scan time=10.7963
Intensity
m/z
VDAFCGGSIVNEK
M
y8
y3
b3-18
b10-18
imm_K
imm_F
y4
b7-18
b5-18
M-18
y7
b4-18
a3
y2
y6
b2
y5
b3
y6++
b5
a6
b6
M-36
b7
b2-18
y8++
b8-18
b8
b9-18
b9
y9
b10
b12-18
b12
y12
imm_E
y1
a5
a7
5.000e+3
4.000e+3
3.000e+3
2.000e+3
1.000e+3
0.000e+0
200
400
600
800
1000
1200

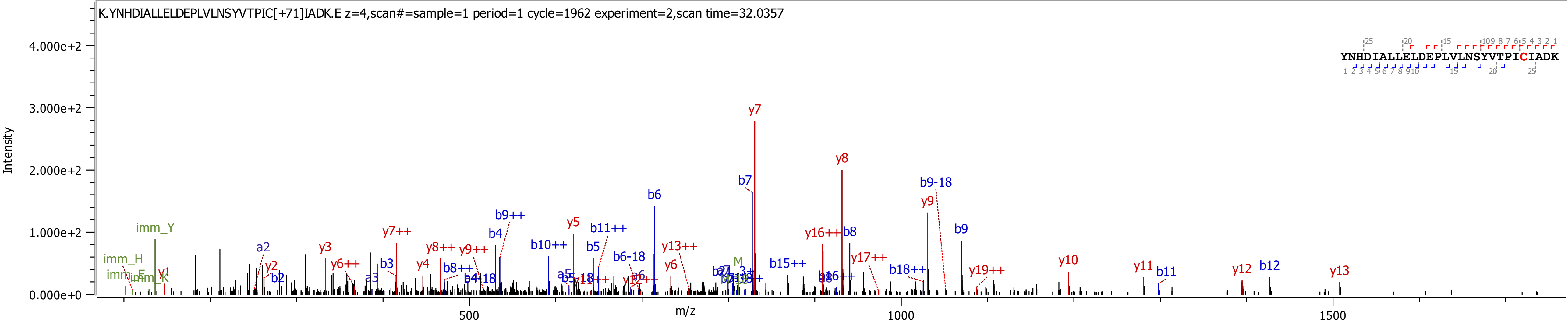
K.YNHDIALLELDEPLVLNSYVTPIC[+71]IADK.E z=4,scan#=sample=1 period=1 cycle=1962 experiment=2,scan time=32.0357
YNHDIALLELDEPLVLNSYVTPICIADK
Intensity
4.000e+2
3.000e+2
2.000e+2
1.000e+2
0.000e+0
m/z
500
1000
1500
imm_Y
imm_H
y1
a2
y2
b2
y3
y6++
b3
a3
y7++
y4
y8++
b8++
y9++
b4
b9++
b10++
y5
b5
b11++
b6-18
b6
y13++
y6
y7
b7
b15++
y16++
b8
y8
y17++
b18++
b9-18
y9
b9
y19++
y10
y11
b11
y12
b12
y13

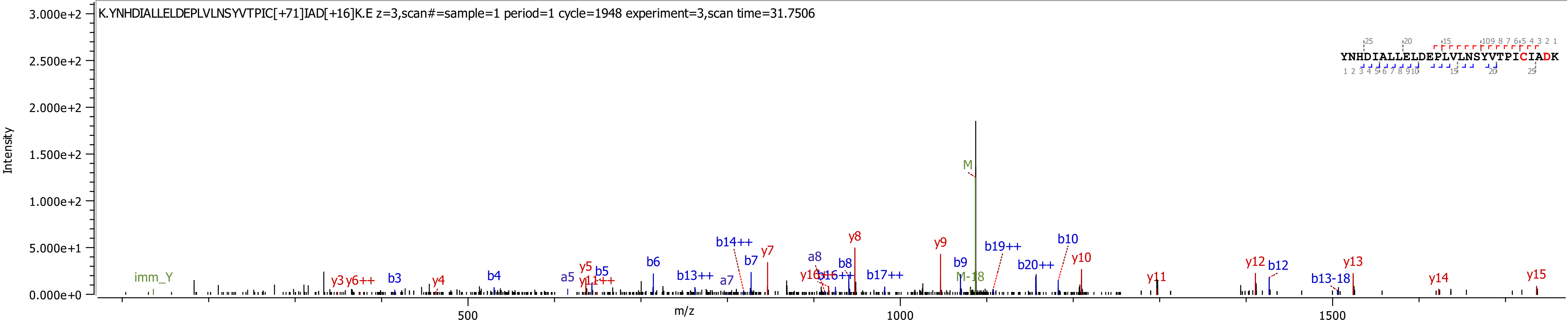

K.YNHDIALLELDEPLVLNSYVTPIC[+71]IAD[+16]K.E z=3,scan#=sample=1 period=1 cycle=1948 experiment=3,scan time=31.7506
YNHDIALLELDEPLVLNSYVTPICIADK
Intensity
m/z
M
M-18
imm_Y
y3 y6++ b3 y4 b4 a5 y5 y11++ b5 b6 b13++ a7 b14++ b7 y7 a8 y16 b16++ b8 y8 b17++ y9 b9 b19++ b20++ b10 y10 y11 y12 b12 b13-18 y13 y14 y15

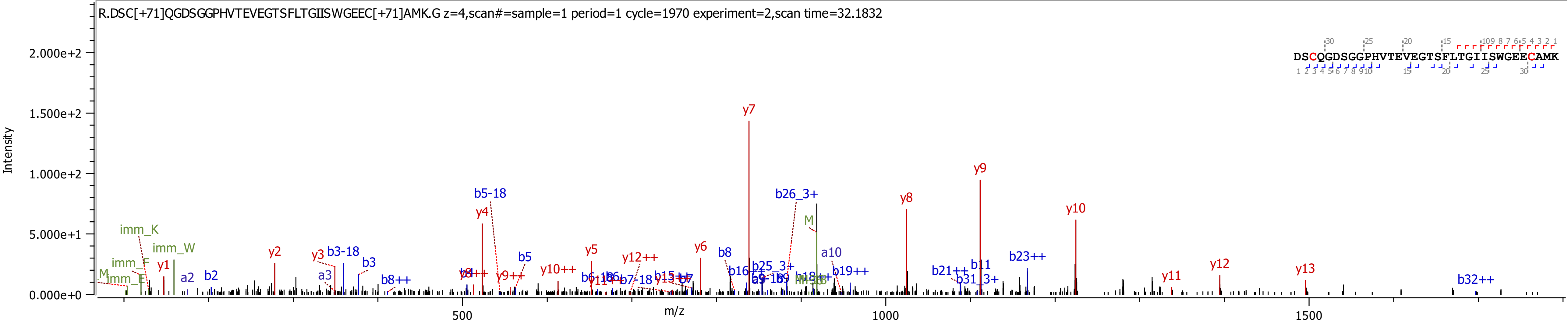

R.DSC[+71]QGDSGGPHVTEVEGTSFLTGIISWGEEC[+71]AMK.G z=4,scan#=sample=1 period=1 cycle=1970 experiment=2,scan time=32.1832
DSCQGDSGGPHVTEVEGTSFLTGIISWGEECAMK
Intensity
m/z
2.000e+2
1.500e+2
1.000e+2
5.000e+1
0.000e+0
500
1000
1500
y7
y9
y8
y10
b26_3+
b5-18
y4
b23++
y13
y12
y11
imm_K
imm_W
y1
a2
b2
y2
y3
b3-18
b3
a3
b8++
b5
y5
y6
b8
y10++
y12++
a10
b19++
b21++
b11
b31_3+
b32++

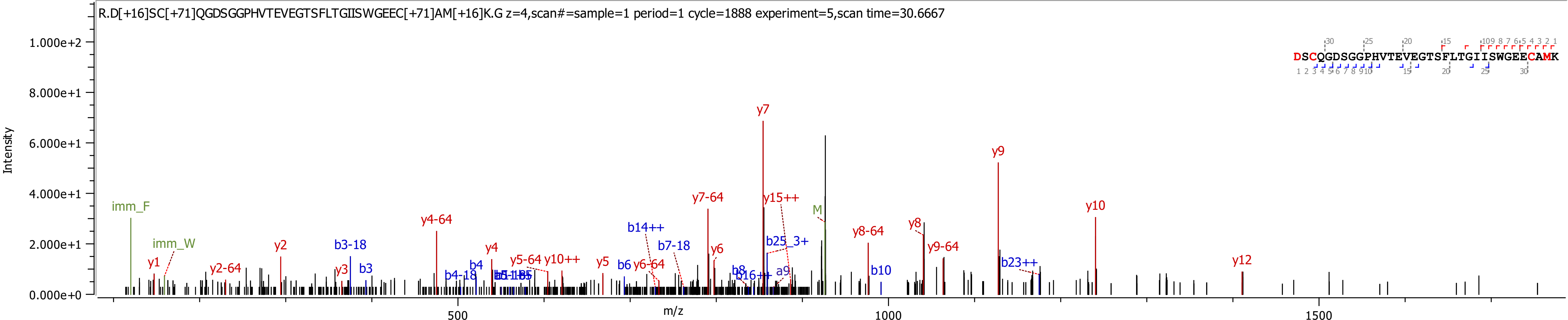
R.D[+16]SC[+71]QGDSGGPHVTEVEGTSFLTGIISWGEEC[+71]AM[+16]K.G z=4,scan#=sample=1 period=1 cycle=1888 experiment=5,scan time=30.6667
DSCQGDSGGPHVTEVEGTSFLTGIISWGEECAMK
Intensity
m/z
1.000e+2
8.000e+1
6.000e+1
4.000e+1
2.000e+1
0.000e+0
500
1000
1500
imm_F
imm_W
y1
y2-64
y2
b3-18
y3
b3
y4-64
b4-18
b4
y4
y5-64
y10++
y5
b6
y6-64
b14++
b7-18
y7-64
y6
y7
y15++
b25_3+
b8
b16++
a9
M
y8-64
b10
y8
y9-64
y9
b23++
y10
y12

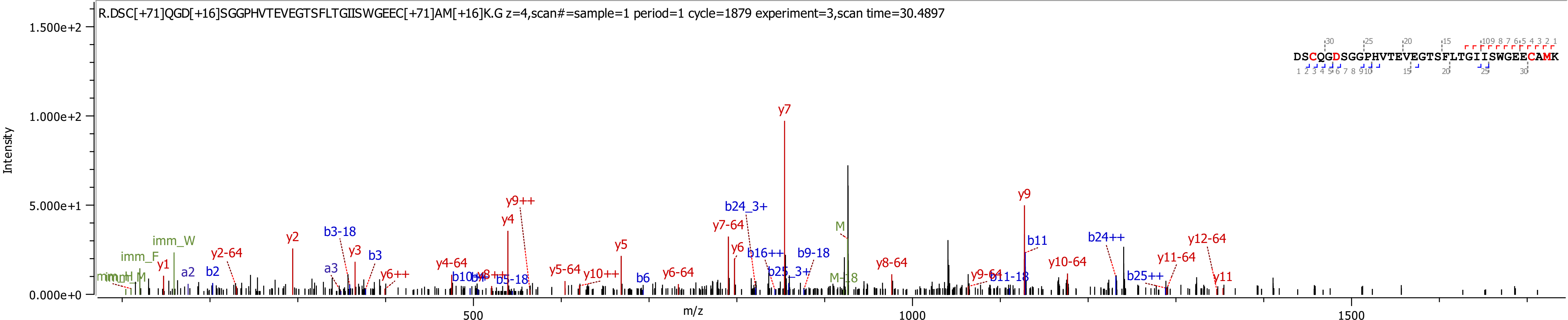
R.DSC[+71]QGD[+16]SGGPHVTEVEGTSFLTGIISWGEEC[+71]AM[+16]K.G z=4,scan#=sample=1 period=1 cycle=1879 experiment=3,scan time=30.4897
DSCQGDSGGPHVTEVEGTSFLTGIISWGEECAMK
Intensity
m/z
1.500e+2
1.000e+2
5.000e+1
0.000e+0
500
1000
1500
y7
y9
y9++
y4
b24_3+
y7-64
M
b11
b24++
y12-64
y2
b3-18
y3
b3
imm_W
imm_F
y1
a2
b2
y2-64
a3
y6++
y4-64
y5
y5-64
y10++
b6
y6-64
y6
b16++
b9-18
b25_3+
M-18
y8-64
y9-64
b11-18
y10-64
b25++
y11-64
y11
b5-18

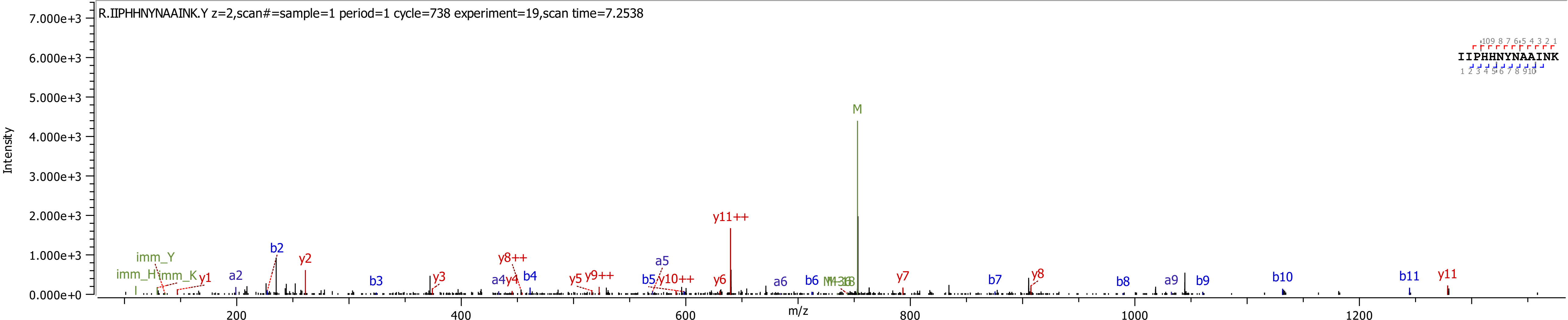

R.IIPHHNYNAAINK.Y z=2,scan#=sample=1 period=1 cycle=738 experiment=19,scan time=7.2538
IIPHHNYNAAINK
Intensity
m/z
M
y11++
b2
y2
imm_Y
imm_H
imm_K
y1
a2
b3
y3
y8++
a4
y4
b4
y5
y9++
a5
b5
y10++
y6
a6
b6
y7
b7
y8
b8
a9
b9
b10
b11
y11

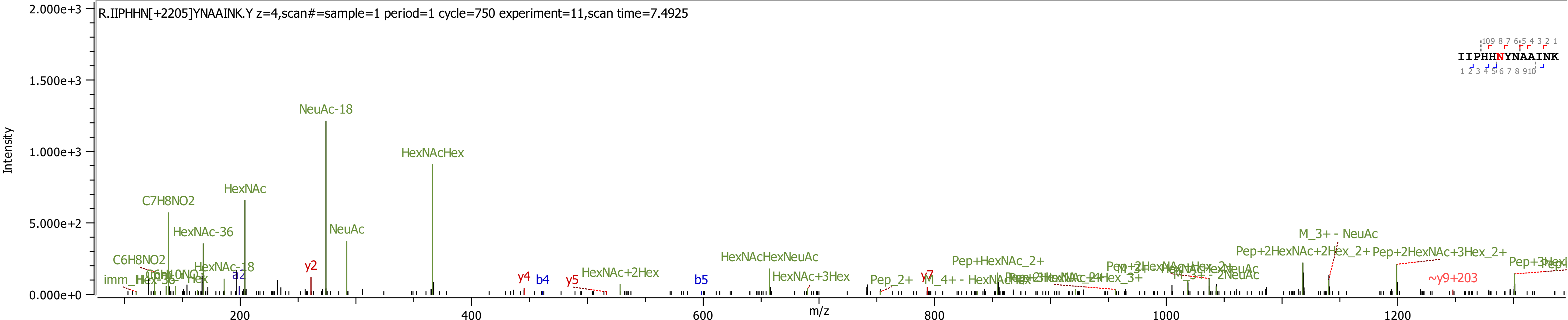
R.IIPHHN[+2205]YNAAINK.Y z=4,scan#=sample=1 period=1 cycle=750 experiment=11,scan time=7.4925
Intensity
m/z
NeuAc-18
HexNAcHex
HexNAc
C7H8NO2
HexNAc-36
NeuAc
C6H8NO2
HexNAc-18
y2
a2
y4
b4
y5
b5
HexNAc+2Hex
HexNAcHexNeuAc
HexNAc+3Hex
Pep_2+
y7
Pep+HexNAc_2+
M_3+ - NeuAc
Pep+2HexNAc+2Hex_2+
Pep+2HexNAc+3Hex_2+
~y9+203
IIPHHNYNAAINK

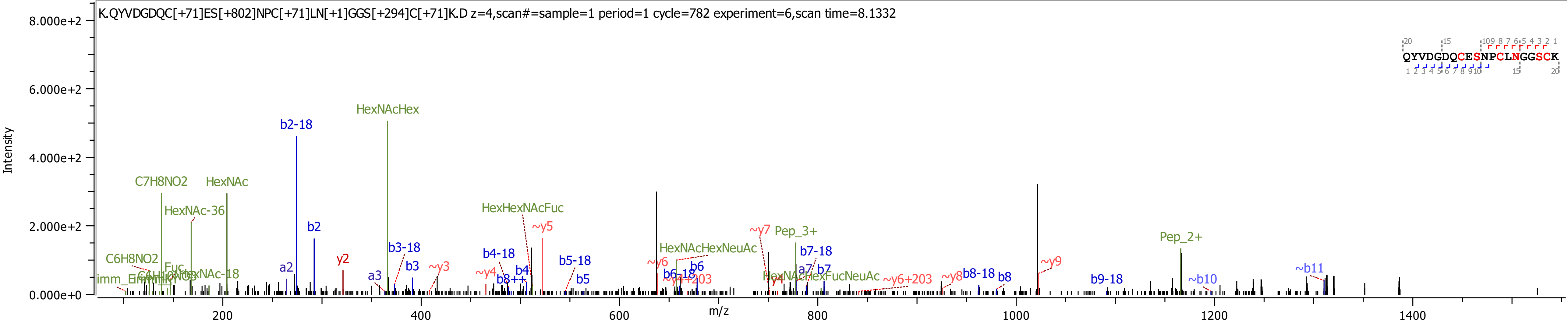
K.QYVDGDQC[+71]ES[+802]NPC[+71]LN[+1]GGS[+294]C[+71]K.D z=4,scan#=sample=1 period=1 cycle=782 experiment=6,scan time=8.1332
QYVDGDQCESNPCLNGGSCK
Intensity
m/z
8.000e+2
6.000e+2
4.000e+2
2.000e+2
0.000e+0
200
400
600
800
1000
1200
1400
HexNAcHex
b2-18
C7H8NO2
HexNAc
HexNAc-36
HexHexNAcFuc
~y5
b2
y2
a2
b3-18
b3
~y3
a3
b4-18
b4
~y4
b8++
b5-18
b5
C6H8NO2
imm_
~y6
b6-18
b6
HexNAcHexNeuAc
~y7
Pep_3+
b7-18
a7
b7
HexNAcHexFucNeuAc
~y6+203
~y8
b8-18
b8
~y9
b9-18
Pep_2+
~b10
~b11

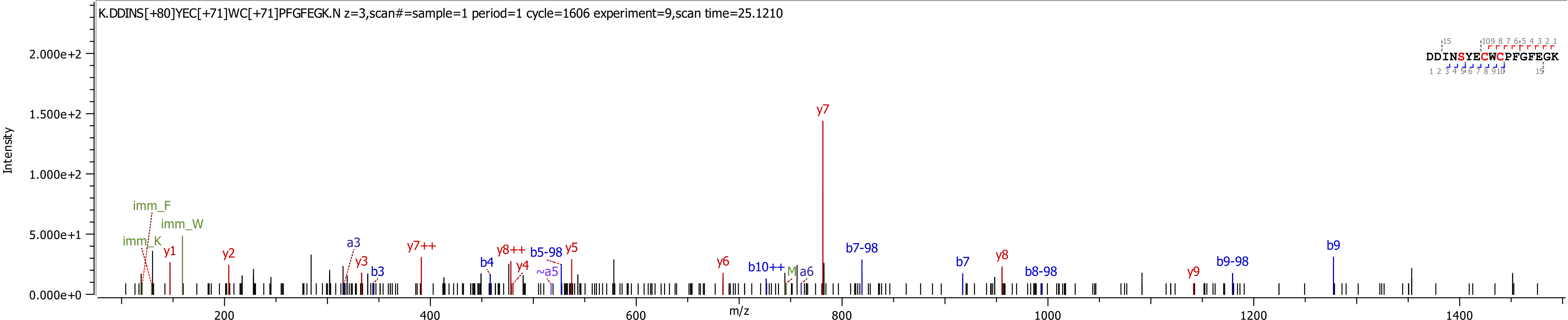

K.DDINS[+80]YEC[+71]WC[+71]PFGFEGK.N z=3,scan#=sample=1 period=1 cycle=1606 experiment=9,scan time=25.1210
DDINSYECWCPFGFEGK
Intensity
m/z
2.000e+2
1.500e+2
1.000e+2
5.000e+1
0.000e+0
200
400
600
800
1000
1200
1400
imm_F
imm_W
imm_K
y1
y2
a3
y3
b3
y7++
b4
y8++
y4
b5-98
~a5
y5
y6
b10++
M
a6
y7
b7-98
b7
y8
b8-98
y9
b9-98
b9

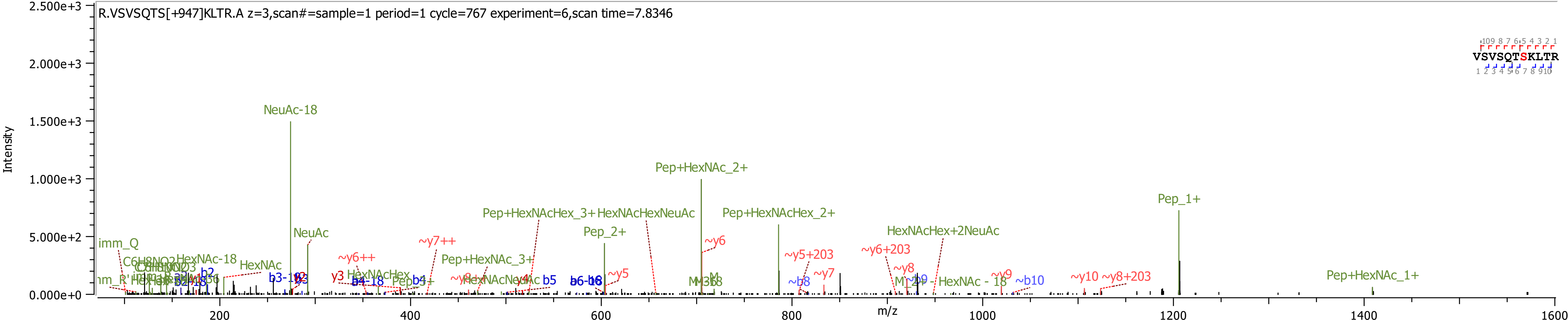

R.VSVSQTS[+947]KLTR.A z=3,scan#=sample=1 period=1 cycle=767 experiment=6,scan time=7.8346
VSVSQTSKLTR
Intensity
m/z
2.500e+3
2.000e+3
1.500e+3
1.000e+3
5.000e+2
0.000e+0
200
400
600
800
1000
1200
1400
1600
NeuAc-18
NeuAc
Pep+HexNAc_2+
Pep+HexNAcHex_3+
HexNAcHexNeuAc
Pep+HexNAcHex_2+
Pep_2+
Pep_1+
HexNAcHex+2NeuAc
Pep+HexNAc_3+
Pep+HexNAc_1+
imm_Q
HexNAc-18
HexNAc
C6H8NO2
HexNAcHex
~y6++
~y7++
~y6
~y5
~y5+203
~y6+203
~y7
~y8
~y9
~y10
~y8+203
~b8
~b9
~b10
b2
b3-18
b4-18
b4
b5
b6-18
HexNAc - 18

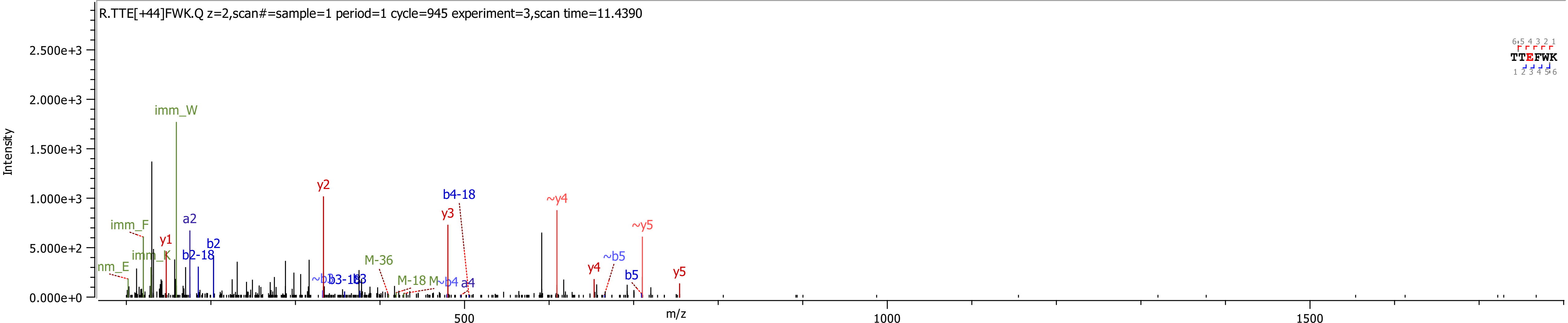
R.TTE[+44]FWK.Q z=2,scan#=sample=1 period=1 cycle=945 experiment=3,scan time=11.4390
Intensity
m/z
TTEFWK
imm_W
imm_F
imm_K
y1
a2
b2
b2-18
y2
b4-18
y3
~y4
~y5
y4
~b5
b5
y5
M-36
M-18
a4
0.000e+0
5.000e+2
1.000e+3
1.500e+3
2.000e+3
2.500e+3
500
1000
1500
