## Supplementary Information 5 for "Analysis of coagulation factor IX in bioreactor cell culture medium predicts yield and quality of the purified product"

D64-K.D[+15.995]DINSYEC[+71.037]WC[+71.037]PFGFEGK.N_z_3 (2) 1
D64-K.DD[+15.995]INSYEC[+71.037]WC[+71.037]PFGFEGK.N 2
D64-K.DDINSYEC[+71.037]WC[+71.037]PFGFEGK.N_z_3 3
D64-K.DDINSYEC[+71.037]WC[+71.037]PFGFEGK.N 4
D85-K.NC[+71.037]ELD[+15.995]VTC[+71.037]NIK.N 5
D85-K.NC[+71.037]ELDVTC[+71.037]NIK.N_z_3 6
D85-K.NC[+71.037]ELDVTC[+71.037]NIK.N 7

D104-K.NSAD[+15.995]NKVVC[+71.037]SC[+71.037]TEGYR.L_z_3 8
D104-K.NSADNKVVC[+71.037]SC[+71.037]TEGYR.L_z_3 9
D104-K.NSADNKVVC[+71.037]SC[+71.037]TEGYR.L 10
D203-K.VD[+15.995]AFC[+71.037]GGSIVNEK.W 11
D203-K.VDAFC[+71.037]GGSIVNEK.W 12
E7E8E15-K.LE[+43.990]E[+43.990]FVQGNLER.E 13
E7E8E15-K.LEE[+43.990]FVQGNLER.E 14

E7E8E15-K.LEEFVQGNLER.E 15
E7E8E15-K.RYNSGKLE[+43.990]EFVQGNLER.E 16
E7E8E15-K.RYNSGKLEE[+43.990]FVQGNLE[+43.990]R.E_z_4 17
E7E8E15-K.RYNSGKLEE[+43.990]FVQGNLE[+43.990]R.E 18
E7E8E15-K.RYNSGKLEEFVQGNLE[+43.990]R.E 19
E7E8E15-K.RYNSGKLEEFVQGNLER.E_z_4 20
E7E8E15-K.RYNSGKLEEFVQGNLER.E 21

E7E8E15-R.YNSGKLE[+43.990]EFVQGNLER.E 22
E7E8E15-R.YNSGKLEE[+43.990]FVQGNLE[+43.990]R.E 23
E7E8E15-R.YNSGKLEEFVQGNLER.E 24
E26E27E30E33E36-K.C[+71.037]SFEEARE[+43.990]VFENTER.T 25
E26E27E30E33E36-K.C[+71.037]SFEEAREVFENTER.T 26
N258_R.IIPHHNYNAAINK.Y_z_3 27
N258-R.IIPHHN[+2204.772]YNAAINK.Y 28

N258-R.IIPHHNYNAAINK.Y_z_2 ____ 29
S141-R.VSVSQTS[+656.228]KLTR.A_z_2 ____ 30
S141-R.VSVSQTS[+656.228]KLTR.A ____ 31
S141-R.VSVSQTS[+947.323]KLTR.A ____ 32
S141-R.VSVSQTS[+963.318]KLTR.A ____ 33
S141-R.VSVSQTSKLTR.A_z_3 ____ 34
S141-R.VSVSQTSKLTR.A ____ 35

T39E40-R.T[+656.228]TEFWK.Q 36
T39E40-R.TT[+947.323]EFWK.Q 37
T39E40-R.TTE[+43.990]FWK.Q 38
T39E40-R.TTEFWK.Q 39
Y45S53S61-K.QYVDGDQC[+71.037]ES[+426.137]NPC[+71.037]LNGGS[+802.286]C[+71.037]K 40

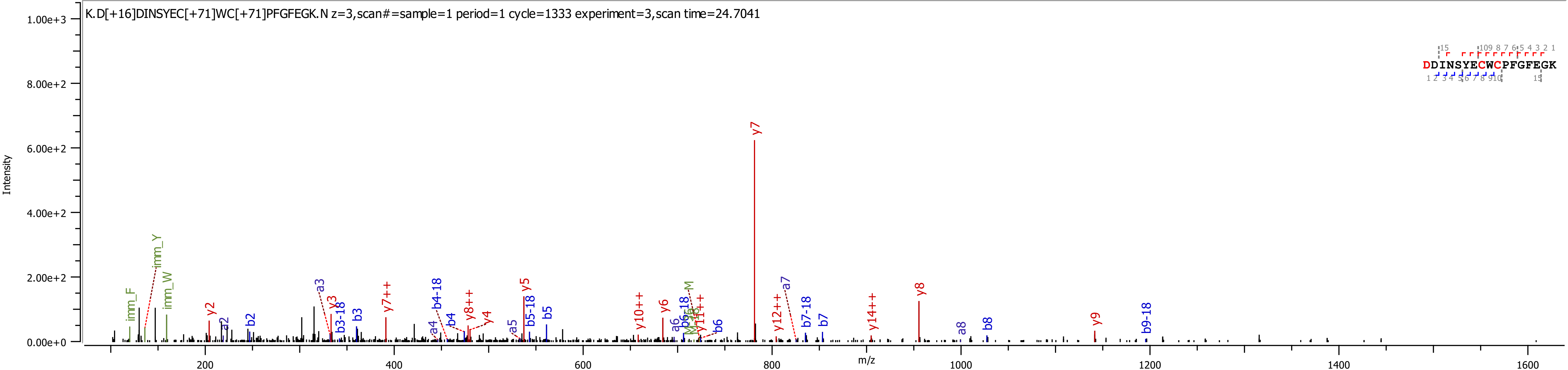

K.D[+16]DINSYEC[+71]WC[+71]PFGFEGK.N z=3,scan#=sample=1 period=1 cycle=1333 experiment=3,scan time=24.7041
DDINSYECWCPFGFEGK
Intensity
m/z
1.00e+3
8.00e+2
6.00e+2
4.00e+2
2.00e+2
0.00e+0
200
400
600
800
1000
1200
1400
1600
imm_F
imm_Y
imm_W
y2
a2
b2
a3
y3
b3-18
b3
y7++
a4
b4-18
b4
y8++
y4
a5
y5
b5-18
b5
y10++
y6
a6
b6-18
y11++
b6
y7
y12++
a7
b7-18
b7
y14++
y8
a8
b8
y9
b9-18

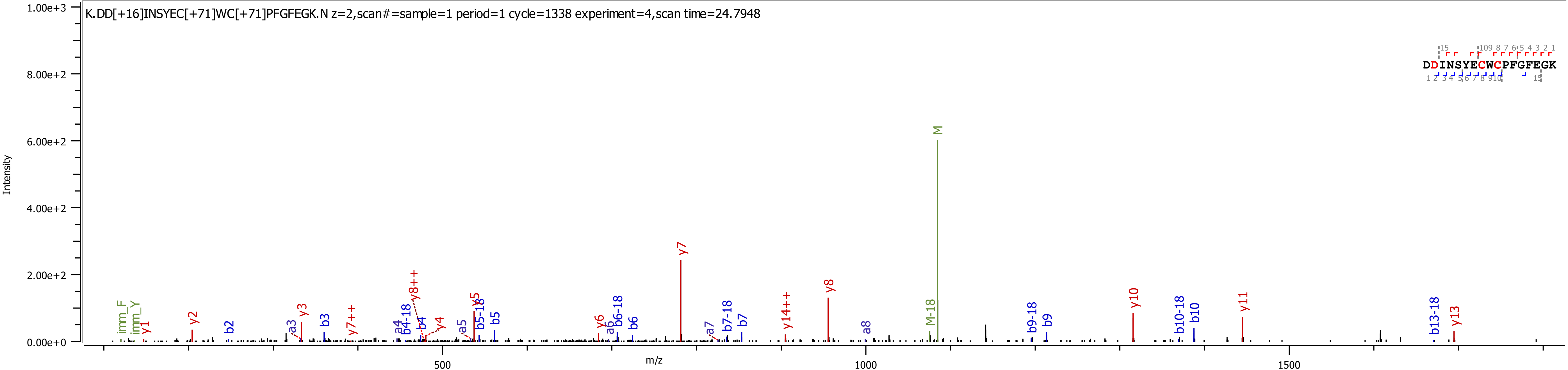

K.DD[+16]INSYEC[+71]WC[+71]PFGFEGK.N z=2,scan#=sample=1 period=1 cycle=1338 experiment=4,scan time=24.7948
DDINSYECWCPFGFEGK
Intensity
m/z
1.00e+3
8.00e+2
6.00e+2
4.00e+2
2.00e+2
0.00e+0
500
1000
1500
M
y7
y8
y10
y11
y13
M-18
b13-18
b10-18
b10
b9-18
b9
a8
y14++
b7-18
b7
a7
y6
a6
b6-18
b6
y5
b5-18
b5
a5
y4
b4
a4
b4-18
y8++
y7++
b3
y3
a3
b2
y2
y1
imm_F
imm_Y

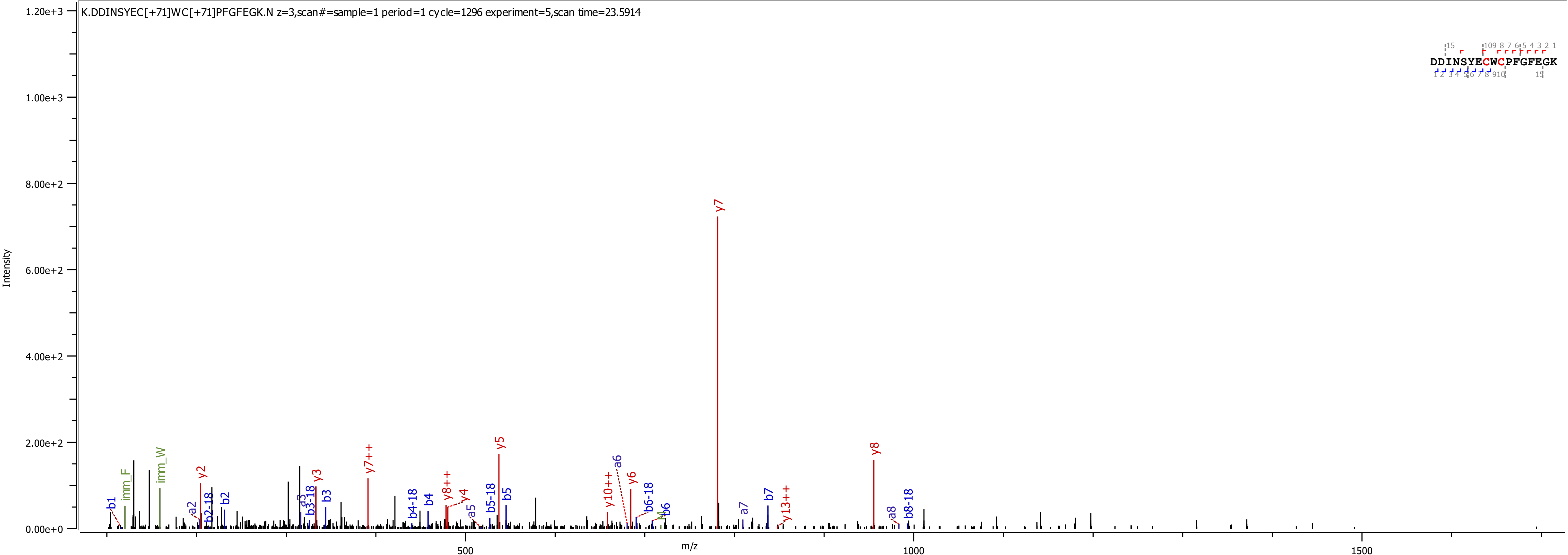
K.DDINSYEC[+71]WC[+71]PFGFEGK.N z=3,scan#=sample=1 period=1 cycle=1296 experiment=5,scan time=23.5914
DDINSYECWCPFGFEGK
Intensity
m/z
1.20e+3
1.00e+3
8.00e+2
6.00e+2
4.00e+2
2.00e+2
0.00e+0
500
1000
1500
y7
y8
y5
y7++
y2
y3
y6
y10++
y8++
y4
y13++
b1
b2
b3
b4
b5
b6
b7
a2
a3
a5
a6
a7
a8
b2-18
b3-18
b4-18
b5-18
b6-18
b8-18
imm_F
imm_W

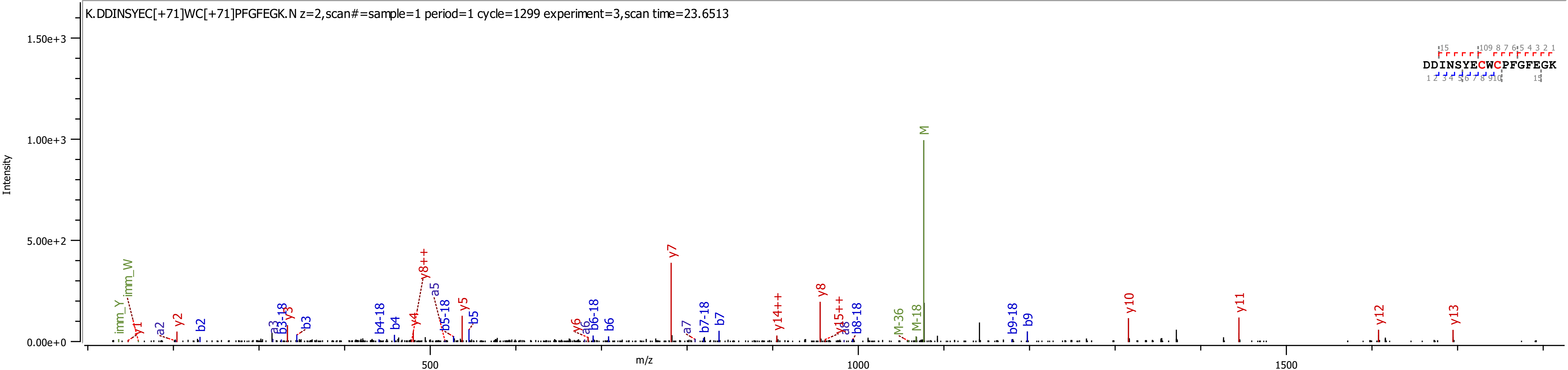

K.DDINSYEC[+71]WC[+71]PFGFEGK.N z=2,scan#=sample=1 period=1 cycle=1299 experiment=3,scan time=23.6513
Intensity
m/z
1.50e+3
1.00e+3
5.00e+2
0.00e+0
500
1000
1500
DDINSYECWCPFGFEGK
M
y7
y8++
y8
y10
y11
y12
y13
y14++
y15++
y5
y6
y4
y3
y2
y1
imm_Y
imm_W
a2
b2
a3
b3-18
b3
b4-18
b4
a5
b5-18
b5
a6
b6-18
b6
a7
b7-18
b7
a8
b8-18
M-36
M-18
b9-18
b9

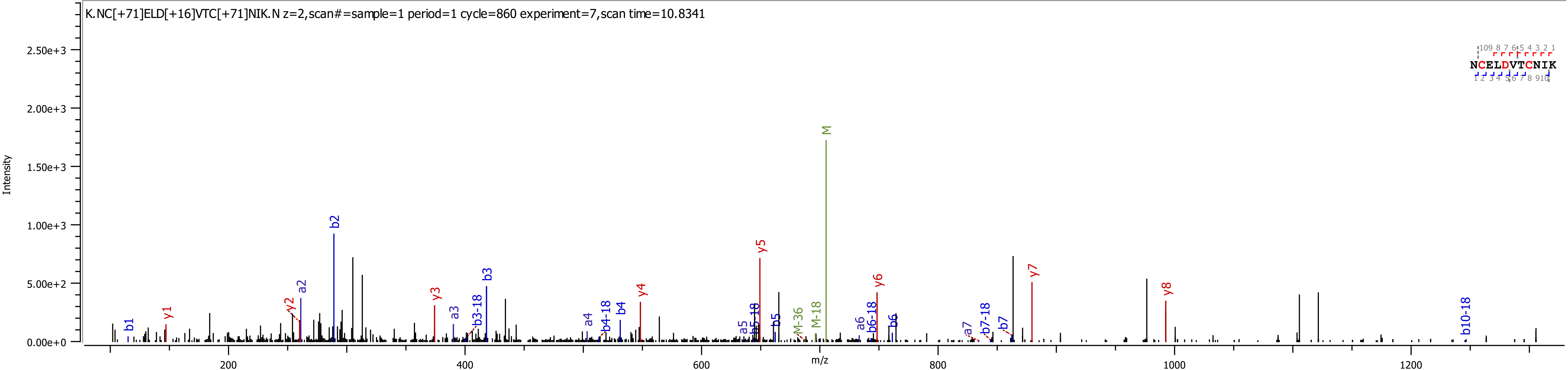
K.NC[+71]ELD[+16]VTC[+71]NIK.N z=2,scan#=sample=1 period=1 cycle=860 experiment=7,scan time=10.8341
Intensity
m/z
NCELDVTCNIK
2.50e+3
2.00e+3
1.50e+3
1.00e+3
5.00e+2
0.00e+0
200
400
600
800
1000
1200
b1
y1
y2
a2
b2
y3
a3
b3-18
b3
a4
b4-18
b4
y4
a5
b5-18
b5
y5
M-36
M-18
M
a6
b6-18
b6
y6
a7
b7-18
b7
y7
y8
b10-18

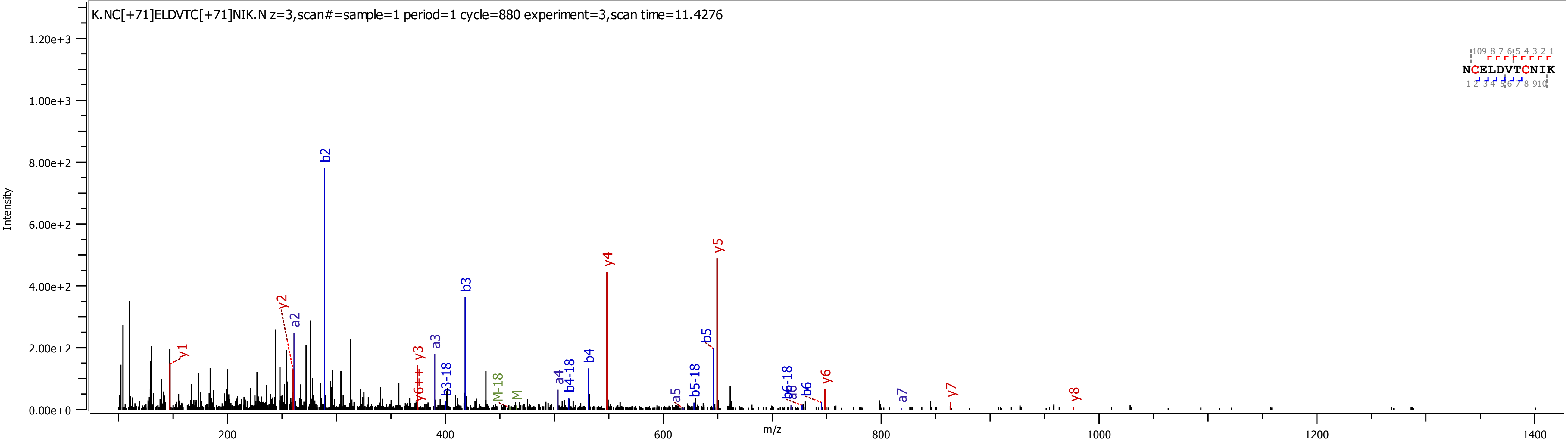

K.NC[+71]ELDVTC[+71]NIK.N z=3,scan#=sample=1 period=1 cycle=880 experiment=3,scan time=11.4276
Intensity
m/z
NCELDVTCNIK
b2
b3
y4
y5
y2
a2
y1
y3
a3
b3-18
y6++
M-18
M
a4
b4-18
b4
a5
b5-18
b5
b6-18
a6
b6
y6
a7
y7
y8

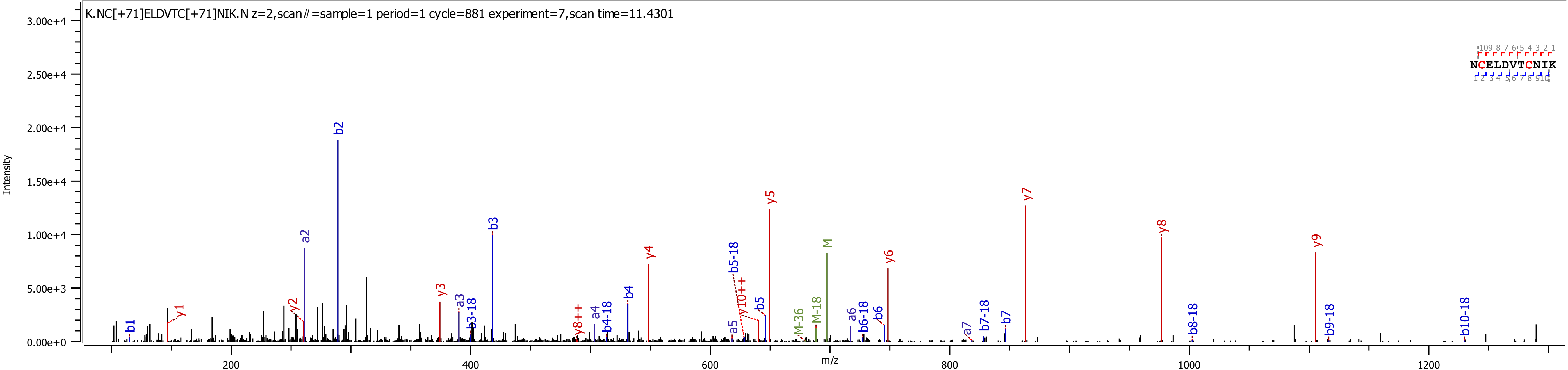
K.NC[+71]ELDVTC[+71]NIK.N z=2,scan#=sample=1 period=1 cycle=881 experiment=7,scan time=11.4301
NCELDVTCNIK
Intensity
m/z
b1
y1
y2
a2
b2
y3
a3
b3-18
b3
y8++
a4
b4-18
b4
y4
a5
b5-18
y10++
b5
y5
M-36
M-18
M
a6
b6-18
b6
y6
a7
b7-18
b7
y7
y8
b8-18
y9
b9-18
b10-18

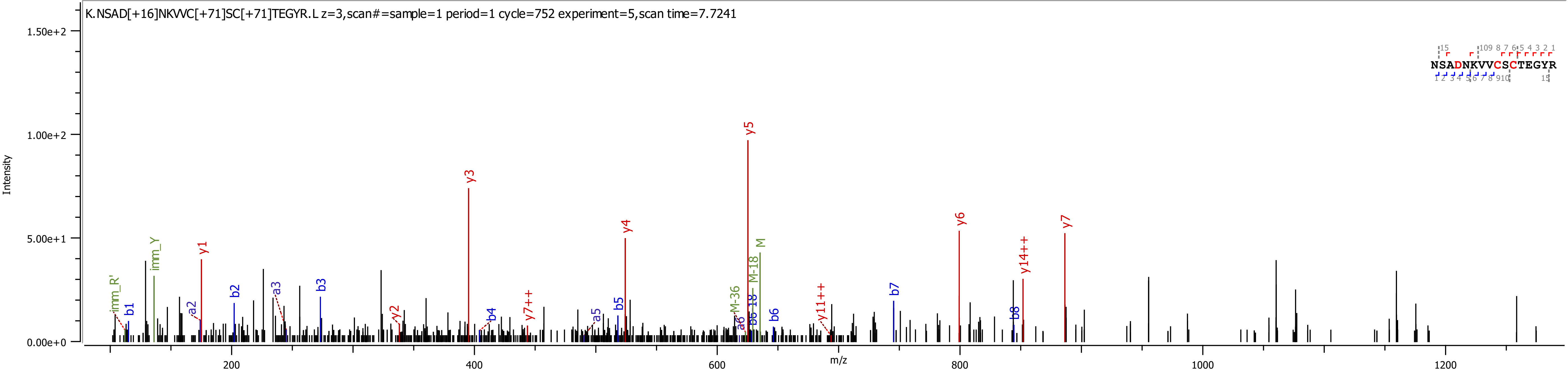
K.NSAD[+16]NKVVC[+71]SC[+71]TEGYR.L z=3,scan#=sample=1 period=1 cycle=752 experiment=5,scan time=7.7241
NSADNKVVCSCTEGYR
Intensity
m/z
1.50e+2
1.00e+2
5.00e+1
0.00e+0
200
400
600
800
1000
1200
imm_R'
b1
imm_Y
a2
y1
b2
a3
b3
y2
y3
b4
y7++
a5
b5
y4
M-36
a6
b6-18
M-18
M
y5
b6
y11++
b7
y6
b8
y14++
y7

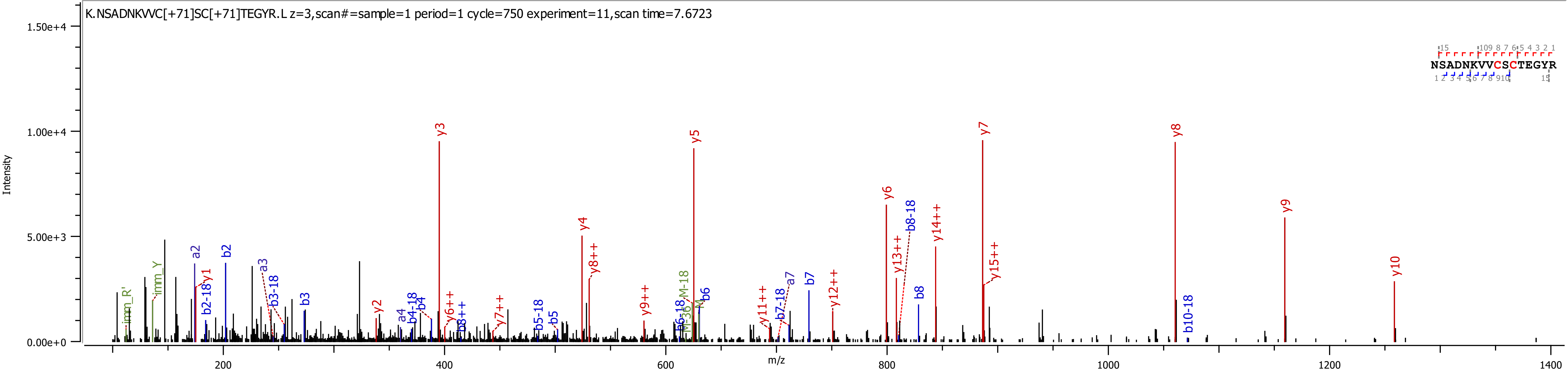

K.NSADNKVVC[+71]SC[+71]TEGYR.L z=3,scan#=sample=1 period=1 cycle=750 experiment=11,scan time=7.6723
NSADNKVVCSCTEGYR
Intensity
m/z
y3
y5
y7
y8
y9
y10
y6
y4
y8++
y14++
y15++
y13++
b8-18
b2
a2
b2-18
y1
imm_Y
imm_R'
a3
b3-18
b3
y2
a4
b4-18
b4
y6++
b8++
y7++
b5-18
b5
y9++
b6-18
M-36
M-18
M
b6
y11++
b7-18
a7
b7
y12++
b8
b10-18

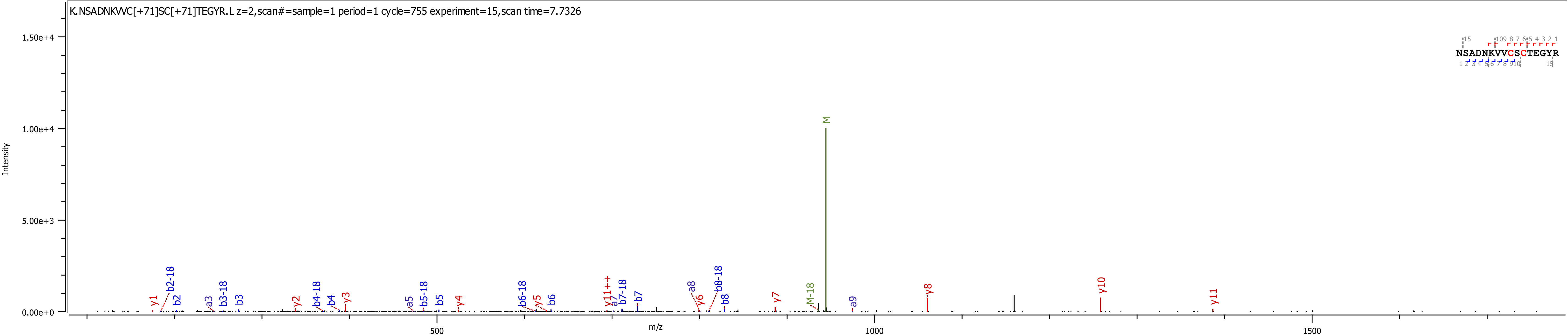
K.NSADNKVVC[+71]SC[+71]TEGYR.L z=2,scan#=sample=1 period=1 cycle=755 experiment=15,scan time=7.7326
Intensity
m/z
NSADNKVVCSCTEGYR
M
M-18
y1
b2-18
b2
a3
b3-18
b3
y2
b4-18
b4
y3
a5
b5-18
b5
y4
b6-18
y5
b6
y11++
a7
b7-18
b7
a8
y6
b8-18
b8
y7
a9
y8
y10
y11
1.50e+4
1.00e+4
5.00e+3
0.00e+0
500
1000
1500

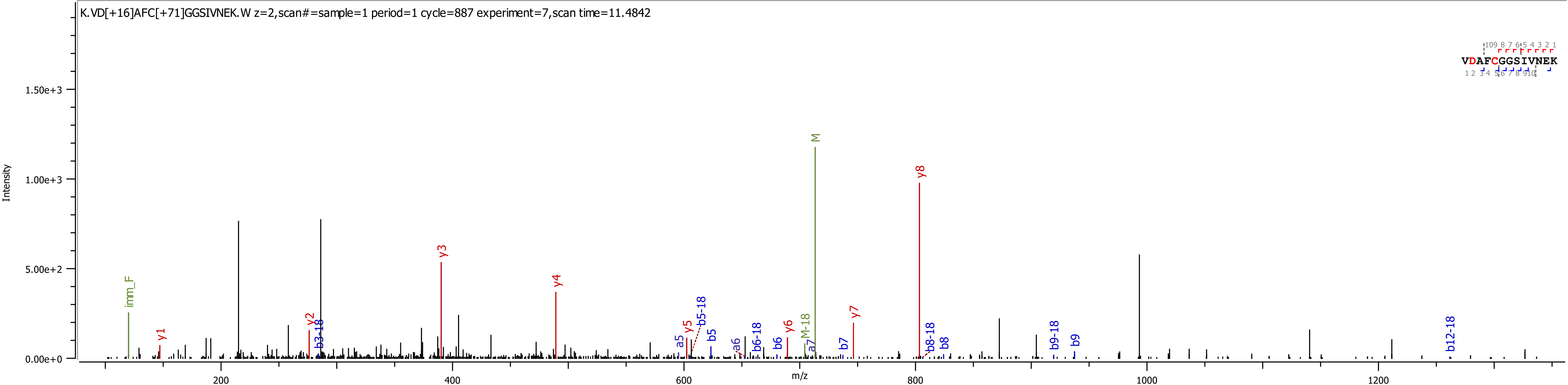
K.VD[+16]AFC[+71]GGSIVNEK.W z=2,scan#=sample=1 period=1 cycle=887 experiment=7,scan time=11.4842
Intensity
m/z
VDAFCGGSIVNEK
imm_F
y1
y2
b3-18
y3
y4
a5
y5
b5-18
b5
a6
b6-18
b6
y6
M-18
a7
M
b7
y7
y8
b8-18
b8
b9-18
b9
b12-18
0.00e+0
5.00e+2
1.00e+3
1.50e+3
200
400
600
800
1000
1200

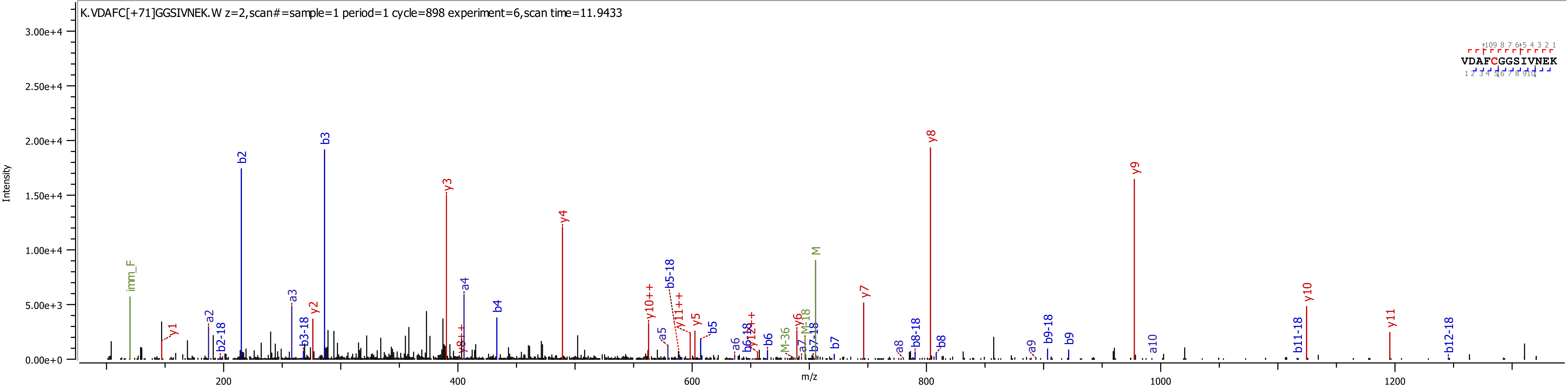
K.VDAFC[+71]GGSIVNEK.W z=2,scan#=sample=1 period=1 cycle=898 experiment=6,scan time=11.9433
VDAFCGGSIVNEK
Intensity
m/z
imm_F
y1
a2
b2-18
b2
a3
b3-18
y2
b3
y3
y8++
a4
b4
y4
y10++
a5
b5-18
y11++
y5
b5
a6
b6-18
y12++
b6
M-36
y6
a7
M-18
b7-18
M
b7
y7
a8
b8-18
y8
b8
a9
b9-18
b9
y9
a10
b11-18
y10
y11
b12-18

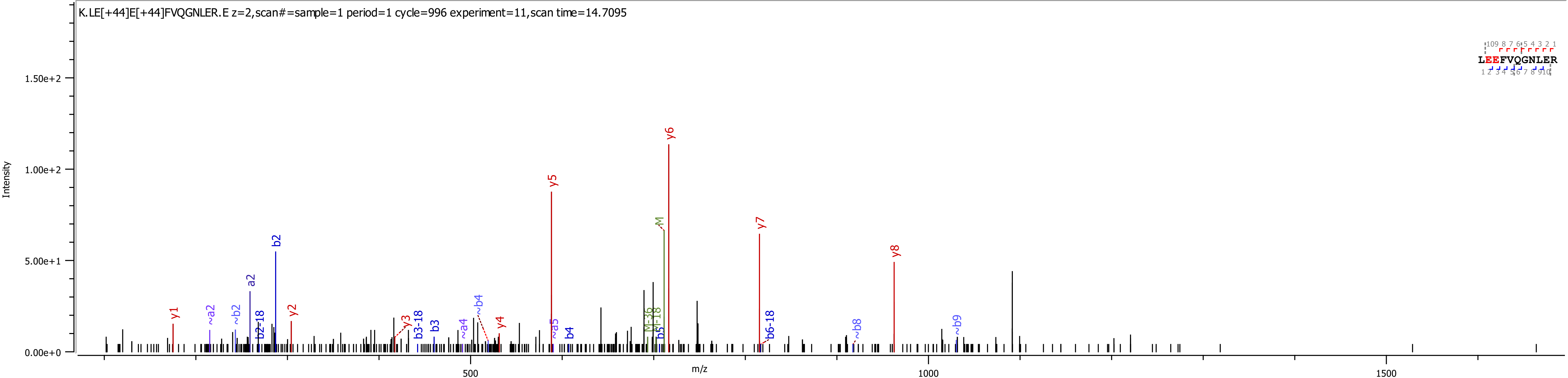
K.LE[+44]E[+44]FVQGNLER.E z=2,scan#=sample=1 period=1 cycle=996 experiment=11,scan time=14.7095
Intensity
m/z
0.00e+0
5.00e+1
1.00e+2
1.50e+2
500
1000
1500
LEEFVQGNLER
y1
~a2
~b2
a2
b2-18
b2
y2
y3
b3-18
b3
~a4
~b4
y4
y5
~a5
b4
M-36
M-18
b5
M
y6
y7
b6-18
~b8
y8
~b9

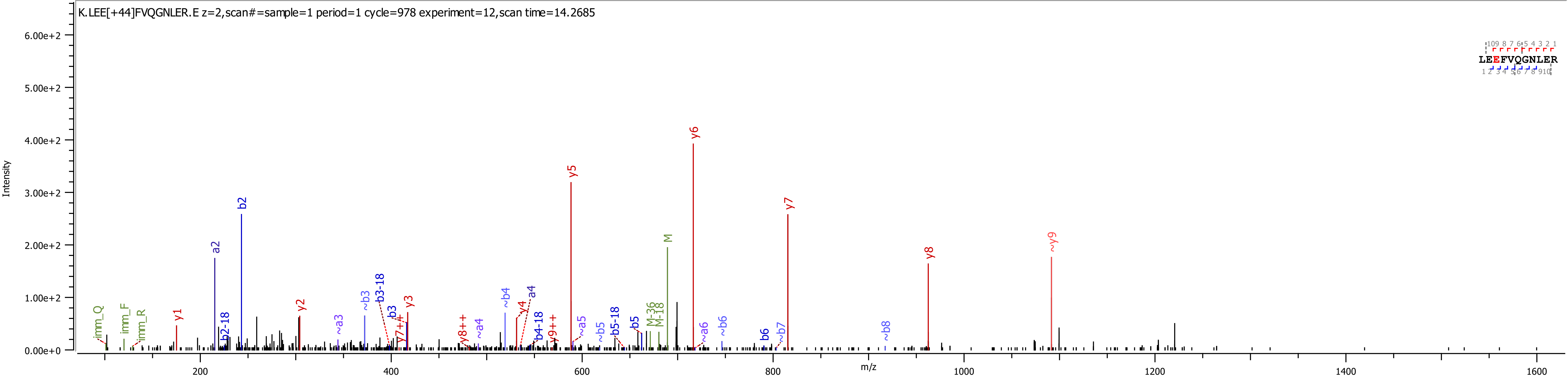
K.LEE[+44]FVQGNLER.E z=2,scan#=sample=1 period=1 cycle=978 experiment=12,scan time=14.2685
LEEFVQGNLER
Intensity
m/z
y6
y5
y7
b2
a2
M
~y9
y8
y1
y2
y3
b3
b3-18
~b3
~a3
y4
a4
~b4
b4-18
y8++
y7++
y9++
~a4
~a5
~b5
b5-18
b5
M-36
M-18
~a6
~b6
b6
~b7
~b8
imm_Q
imm_F
imm_R
b2-18

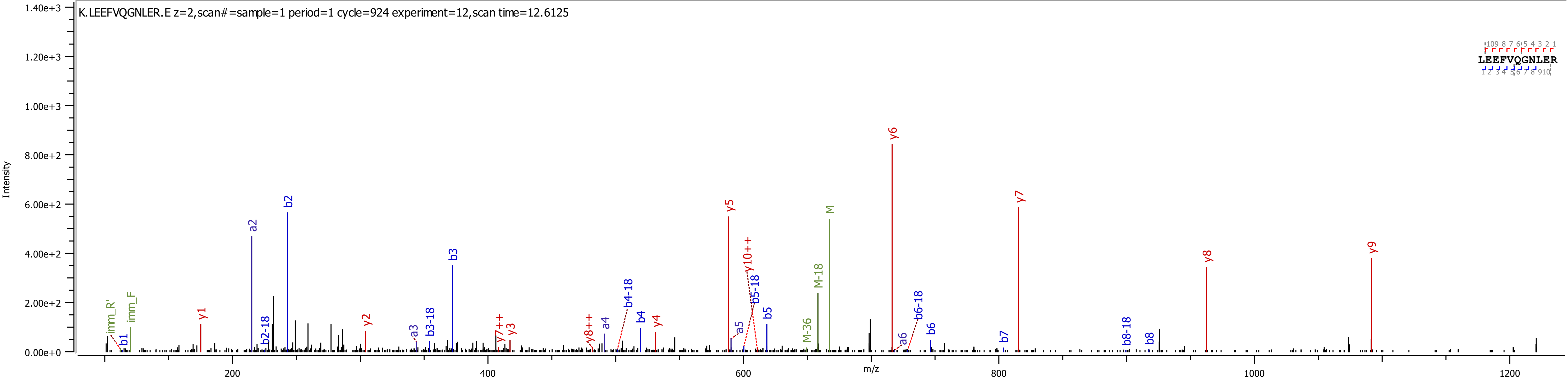
K.LEEFVQGNLER.E z=2,scan#=sample=1 period=1 cycle=924 experiment=12,scan time=12.6125
LEEFVQGNLER
Intensity
m/z
imm_R'
imm_F
b1
y1
a2
b2-18
b2
y2
a3
b3-18
b3
y7++
y3
y8++
a4
b4-18
b4
y4
y5
a5
y10++
b5-18
b5
M-36
M-18
M
y6
a6
b6-18
b6
b7
y7
b8-18
b8
y8
y9

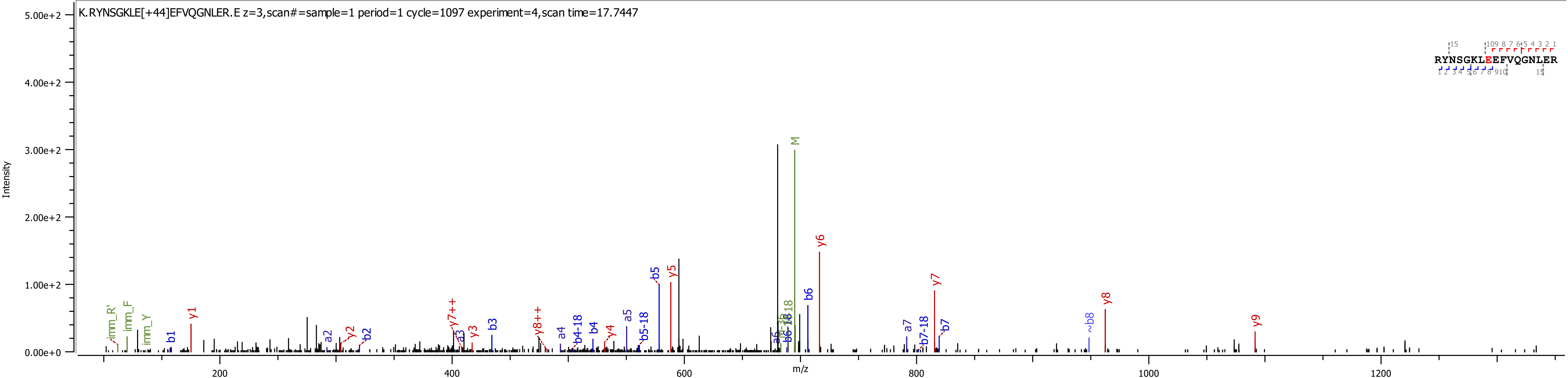

K.RYNSGKLE[+44]EFVQGNLER.E z=3,scan#=sample=1 period=1 cycle=1097 experiment=4,scan time=17.7447
Intensity
m/z
5.00e+2
4.00e+2
3.00e+2
2.00e+2
1.00e+2
0.00e+0
200
400
600
800
1000
1200
RYNSGKLEEFVQGNLER
M
y1
y2
y3
y4
y5
y6
y7
y8
y9
y7++
y8++
b1
b2
b3
b4
b5
b6
b7
~b8
a2
a3
a4
a5
a7
b4-18
b5-18
b7-18
imm_R'
imm_F
imm_Y

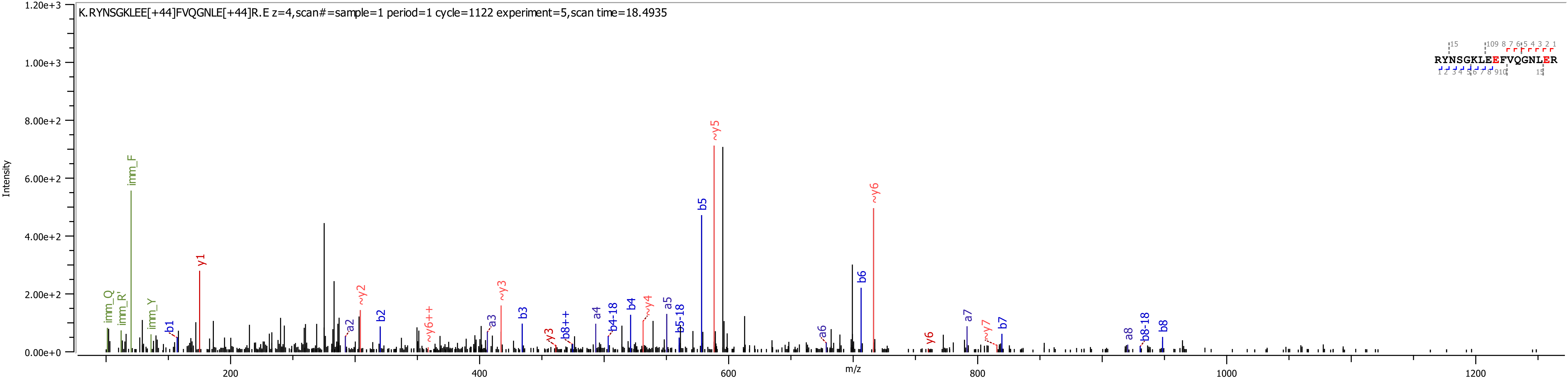
K.RYNSGKLEE[+44]FVQGNLE[+44]R.E z=4,scan#=sample=1 period=1 cycle=1122 experiment=5,scan time=18.4935
Intensity
m/z
1.20e+3
1.00e+3
8.00e+2
6.00e+2
4.00e+2
2.00e+2
0.00e+0
200
400
600
800
1000
1200
RYNSGKLEEFVQGNLER
imm_Q
imm_R'
imm_F
imm_Y
b1
y1
a2
~y2
b2
~y6++
a3
~y3
b3
y3
b8++
a4
b4-18
b4
~y4
a5
b5-18
b5
~y5
a6
b6
~y6
y6
a7
~y7
b7
a8
b8-18
b8

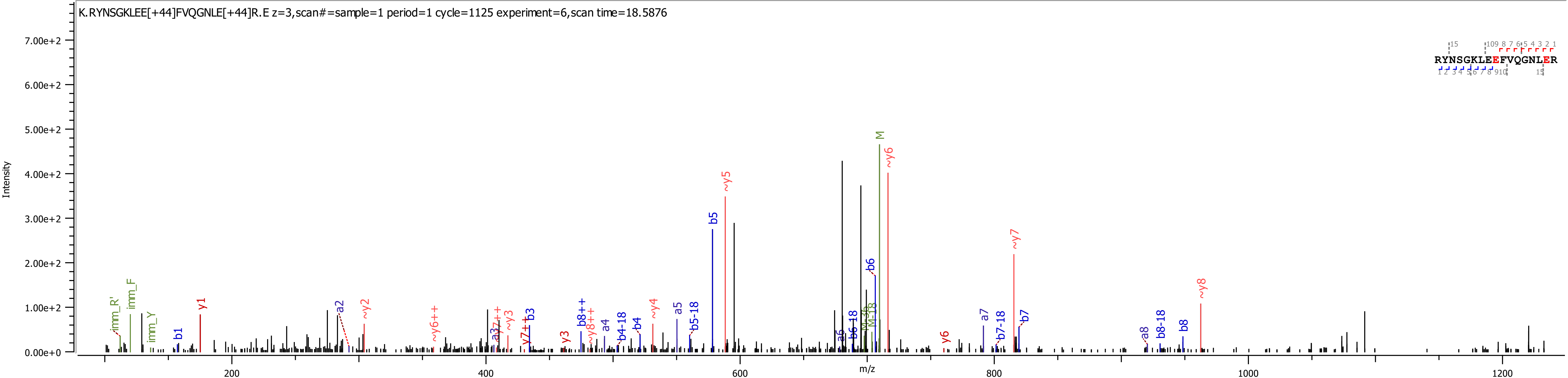
K.RYNSGKLEE[+44]FVQGNLE[+44]R.E z=3,scan#=sample=1 period=1 cycle=1125 experiment=6,scan time=18.5876
Intensity
m/z
RYNSGKLEEFVQGNLER

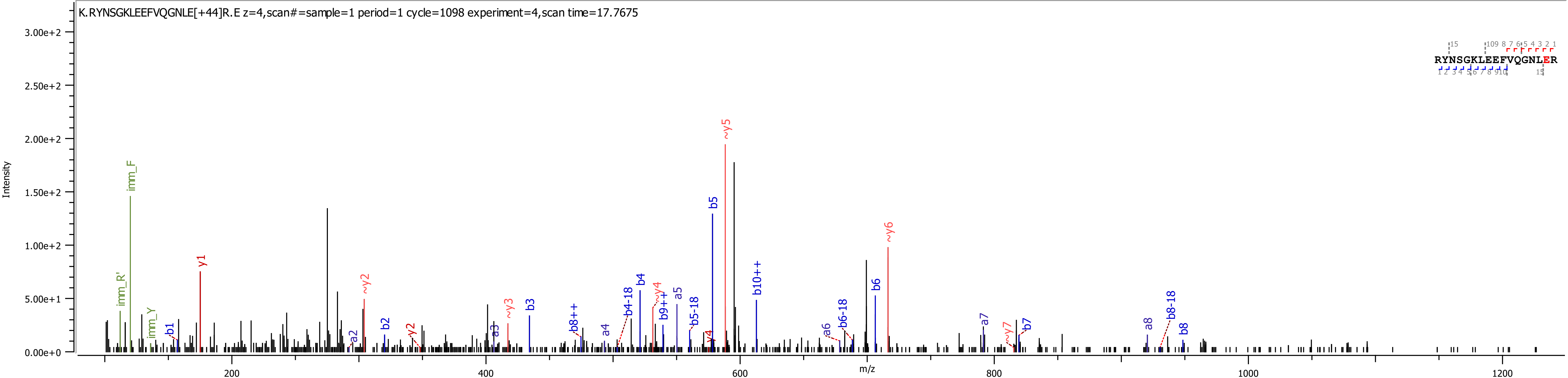
K.RYNSGKLEEFVQGNLE[+44]R.E z=4,scan#=sample=1 period=1 cycle=1098 experiment=4,scan time=17.7675
RYNSGKLEEFVQGNLER
Intensity
m/z

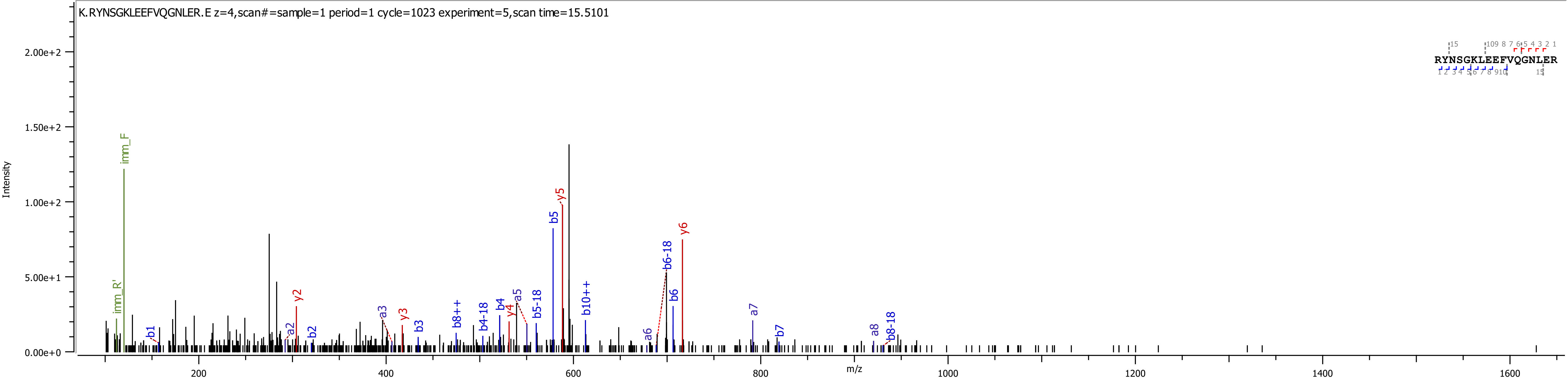

K.RYNSGKLEEFVQGNLER.E z=4,scan#=sample=1 period=1 cycle=1023 experiment=5,scan time=15.5101
RYNSGKLEEFVQGNLER
Intensity
m/z
2.00e+2
1.50e+2
1.00e+2
5.00e+1
0.00e+0
200
400
600
800
1000
1200
1400
1600
imm_F
imm_R'
b1
a2
y2
b2
a3
y3
b3
b8++
b4-18
b4
y4
a5
b5-18
b5
y5
b10++
a6
b6-18
b6
y6
a7
b7
a8
b8-18

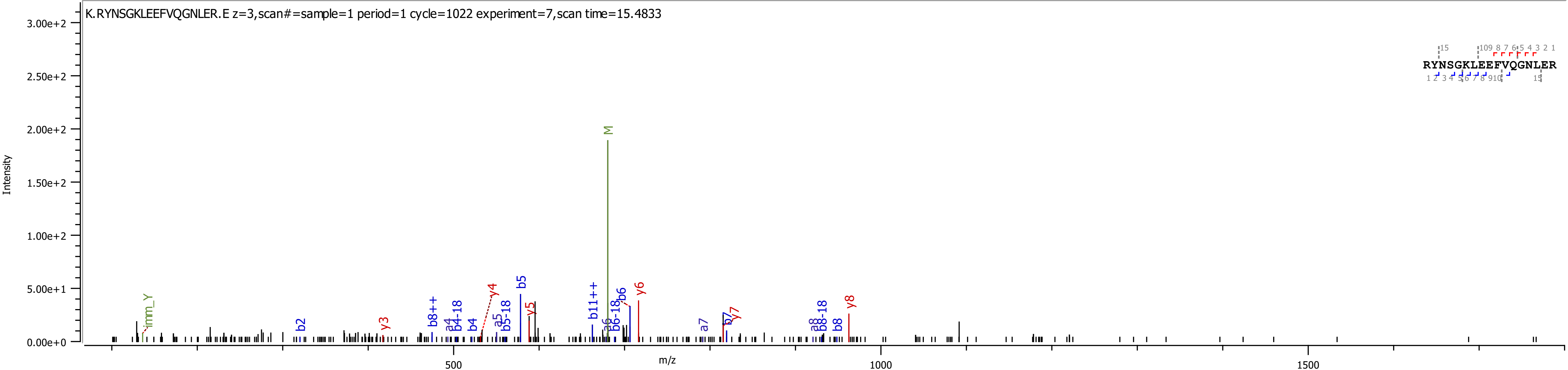

K.RYNSGKLEEFVQGNLER.E z=3,scan#=sample=1 period=1 cycle=1022 experiment=7,scan time=15.4833
Intensity
m/z
RYNSGKLEEFVQGNLER
M
imm_Y
b2
y3
b8++
a4
b4-18
b4
y4
a5
b5-18
b5
y5
b11++
a6
b6-18
b6
y6
a7
b7
y7
a8
b8-18
b8
y8

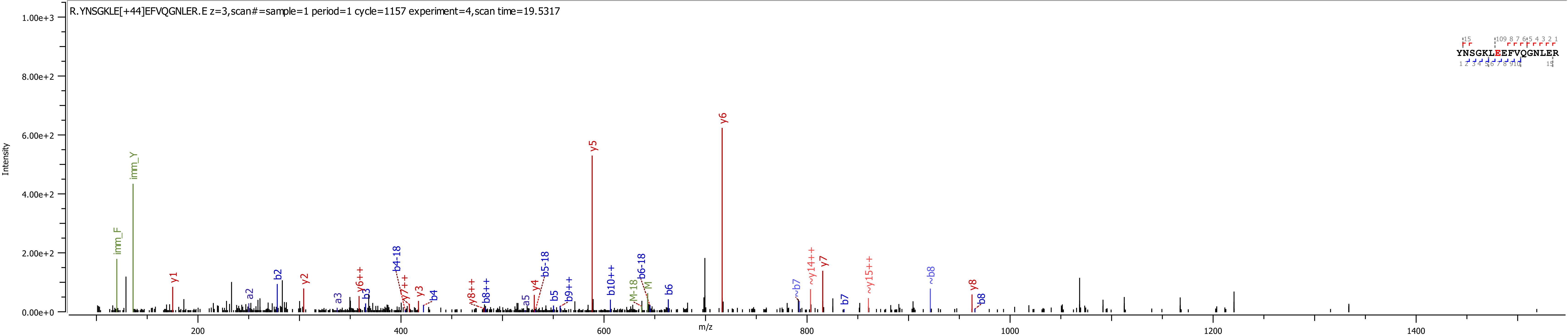
R.YNSGKLE[+44]EFVQGNLER.E z=3,scan#=sample=1 period=1 cycle=1157 experiment=4,scan time=19.5317
Intensity
1.00e+3
8.00e+2
6.00e+2
4.00e+2
2.00e+2
0.00e+0
m/z
200
400
600
800
1000
1200
1400
YNSGKLEEFVQGNLER
imm_Y
imm_F
y1
a2
b2
y2
a3
b3
y6++
b4-18
y7++
y3
b4
y8++
b8++
a5
y4
b5-18
b5
b9++
y5
b10++
M-18
b6-18
M
b6
y6
~b7
~y14++
y7
b7
~y15++
~b8
y8
b8

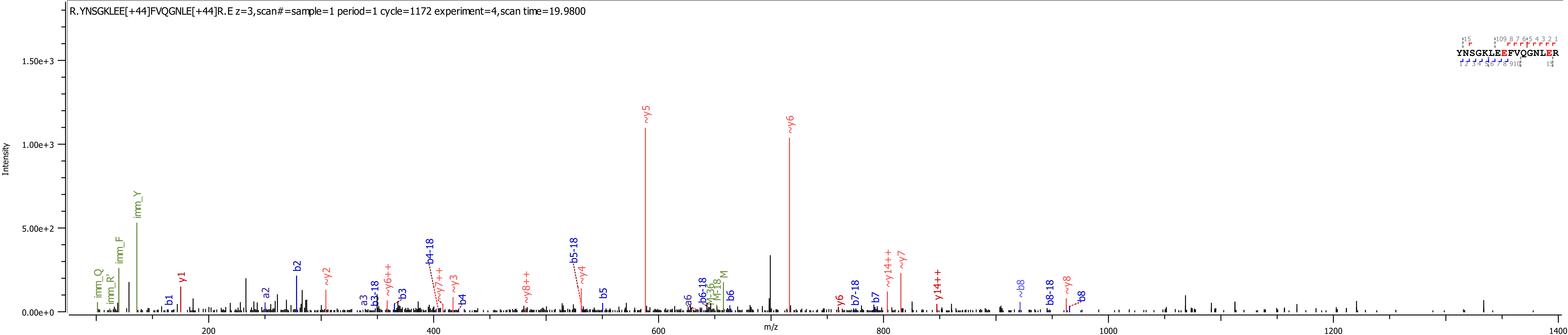
R.YNSGKLEE[+44]FVQGNLE[+44]R.E z=3,scan#=sample=1 period=1 cycle=1172 experiment=4,scan time=19.9800
YNSGKLEEFVQGNLER
Intensity
m/z
0.00e+0
5.00e+2
1.00e+3
1.50e+3
200
400
600
800
1000
1200
1400
imm_Q
imm_R'
imm_F
imm_Y
b1
y1
a2
b2
~y2
a3
b3-18
~y6++
b3
b4-18
~y7++
~y3
b4
~y8++
b5-18
~y4
b5
~y5
a6
b6-18
M-36
M-18
M
b6
~y6
y6
b7-18
b7
~y14++
~y7
y14++
~b8
b8-18
~y8
b8

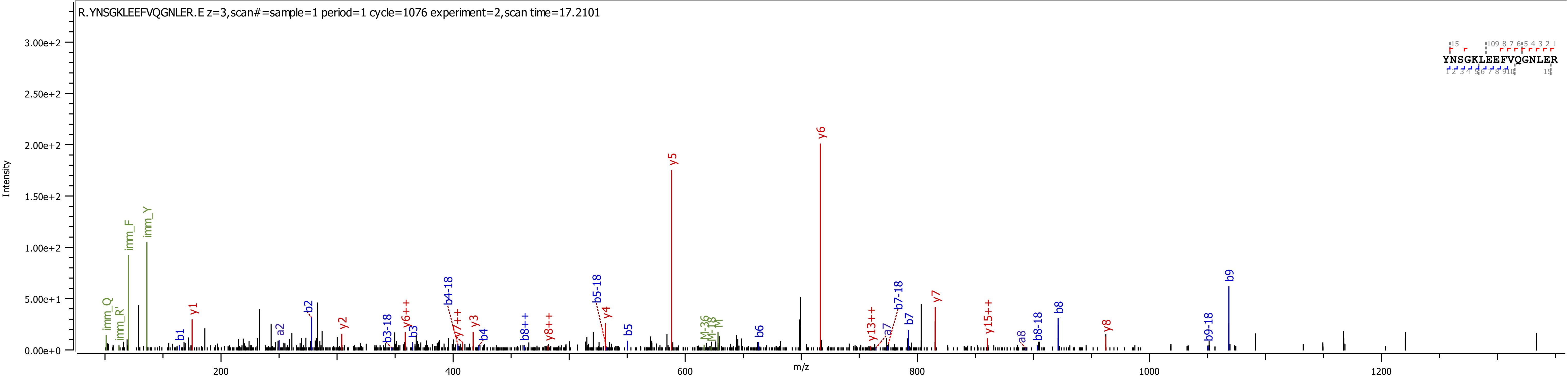
R.YNSGKLEEFVQGNLER.E z=3,scan#=sample=1 period=1 cycle=1076 experiment=2,scan time=17.2101
YNSGKLEEFVQGNLER
Intensity
m/z
3.00e+2
2.50e+2
2.00e+2
1.50e+2
1.00e+2
5.00e+1
0.00e+0
200
400
600
800
1000
1200
imm_Q
imm_R
imm_F
imm_Y
b1
y1
a2
b2
y2
b3-18
y6++
b3
b4-18
y7++
y3
b4
b8++
y8++
b5-18
y4
b5
y5
M-36
M-18
M
b6
y6
y13++
a7
b7-18
b7
y7
y15++
a8
b8-18
b8
y8
b9-18
b9

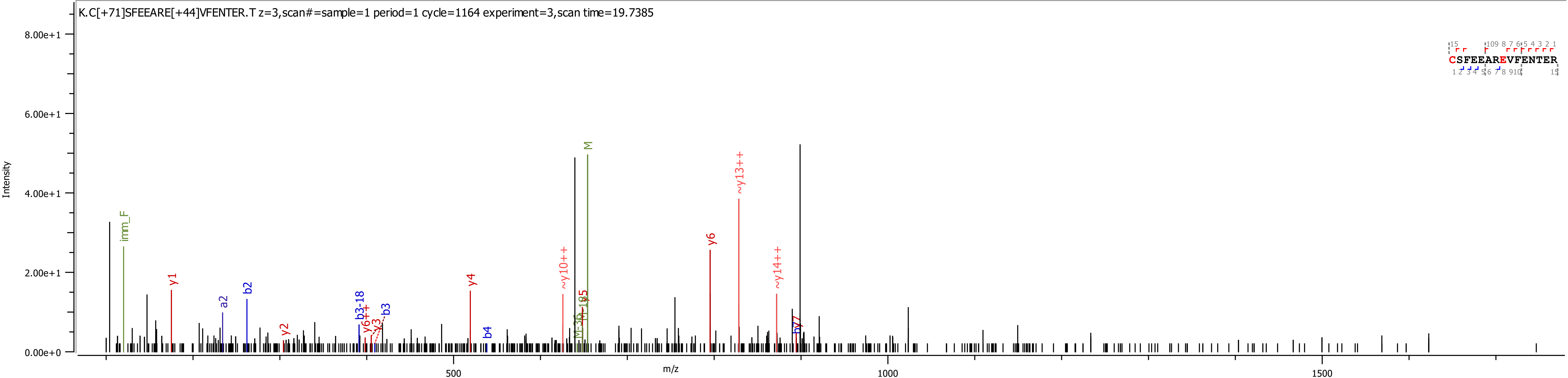

K.C[+71]SFEEARE[+44]VFENTER.T z=3,scan#=sample=1 period=1 cycle=1164 experiment=3,scan time=19.7385
Intensity
8.00e+1
6.00e+1
4.00e+1
2.00e+1
0.00e+0
m/z
500
1000
1500
imm_F
y1
a2
b2
y2
b3-18
y6++
y3
b3
y4
b4
~y10++
M
y5
~y13++
y6
~y14++
b7
CSFEEAREVFENTER

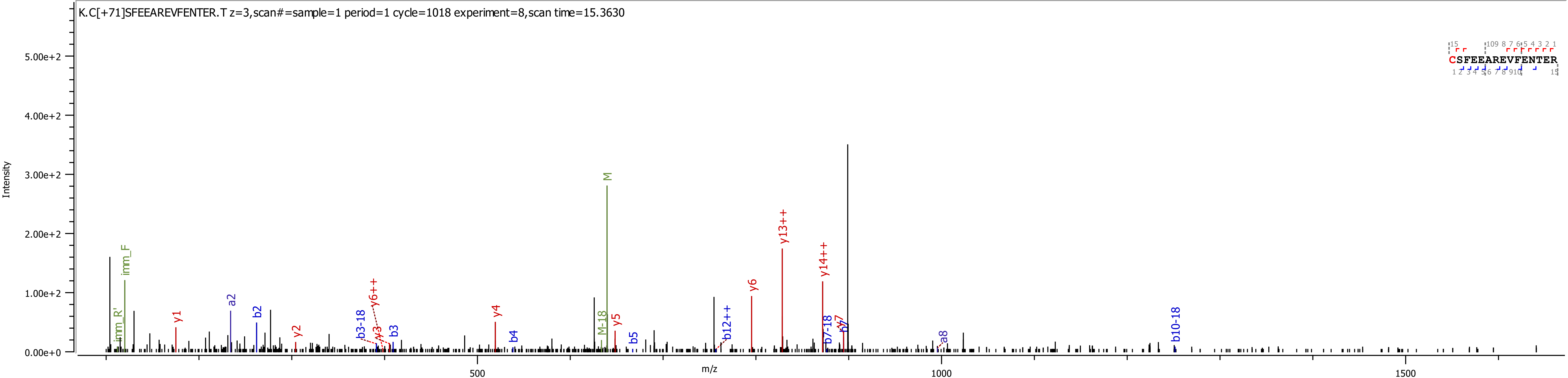
K.C[+71]SFEEAREVFENTER.T z=3,scan#=sample=1 period=1 cycle=1018 experiment=8,scan time=15.3630
CSFEEAREVFENTER
Intensity
m/z
5.00e+2
4.00e+2
3.00e+2
2.00e+2
1.00e+2
0.00e+0
500
1000
1500
imm_R'
imm_F
y1
a2
b2
y2
b3-18
y3
y6++
b3
y4
b4
M-18
M
y5
b5
b12++
y6
y13++
y14++
b7-18
b7
y7
a8
b10-18

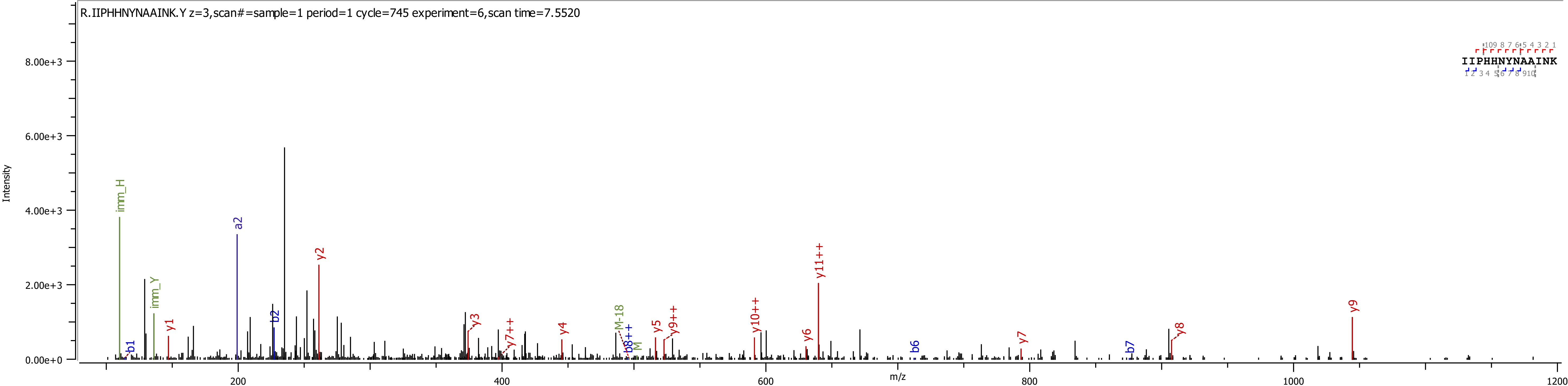
R.IIPHHNYNAAINK.Y z=3,scan#=sample=1 period=1 cycle=745 experiment=6,scan time=7.5520
IIPHHNYNAAINK
Intensity
m/z
8.00e+3
6.00e+3
4.00e+3
2.00e+3
0.00e+0
200
400
600
800
1000
1200
imm_H
imm_Y
a2
b1
b2
b6
b7
b8++
y1
y2
y3
y4
y5
y6
y7
y8
y9
y7++
y9++
y10++
y11++
M-18
M

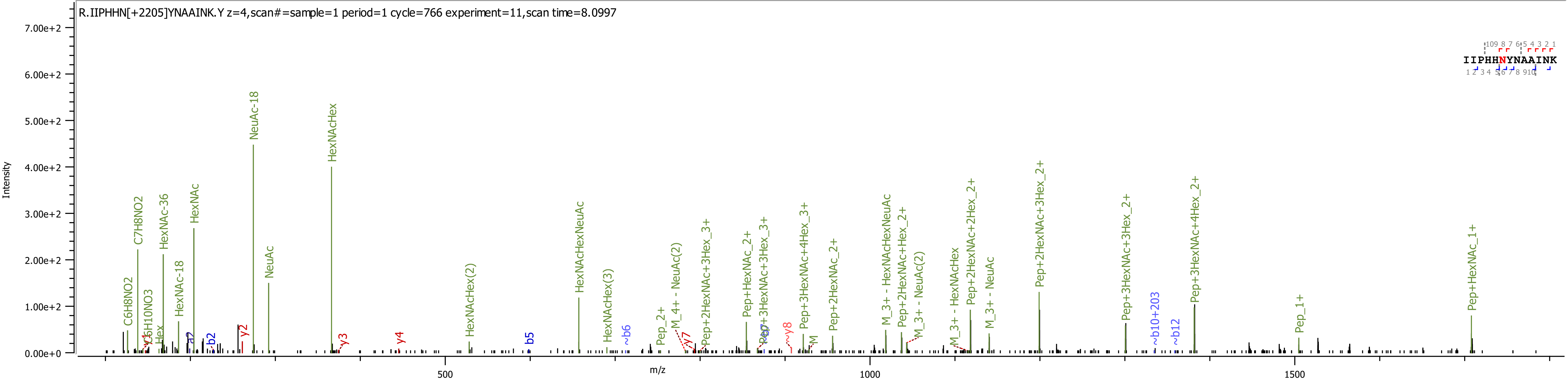

R.IIPHHN[+2205]YNAAINK.Y z=4,scan#=sample=1 period=1 cycle=766 experiment=11,scan time=8.0997
Intensity
m/z
IIPHHNYNAAINK
NeuAc-18
HexNAcHex
C7H8NO2
HexNAc-36
HexNAc
HexNAc-18
C6H8NO2
C6H10NO3
Hex
NeuAc
HexNAcHexNeuAc
HexNAcHex(2)
HexNAcHex(3)
Pep+2HexNAc+3Hex_2+
Pep+3HexNAc+4Hex_2+
Pep+3HexNAc+3Hex_2+
Pep+2HexNAc+2Hex_2+
M_3+ - NeuAc
M_3+ - HexNAcHex
Pep+2HexNAc+Hex_2+
M_3+ - HexNAcHexNeuAc
M_3+ - NeuAc(2)
Pep+2HexNAc_2+
Pep+3HexNAc+4Hex_3+
Pep+3HexNAc+3Hex_3+
Pep+HexNAc_2+
Pep+2HexNAc+3Hex_3+
M_4+ - NeuAc(2)
Pep_2+
Pep_1+
Pep+HexNAc_1+
~b10+203
~b12
~b6
~y8
y7
M
b5
b2
a2
y2
y3
y4
7.00e+2
6.00e+2
5.00e+2
4.00e+2
3.00e+2
2.00e+2
1.00e+2
0.00e+0
500
1000
1500

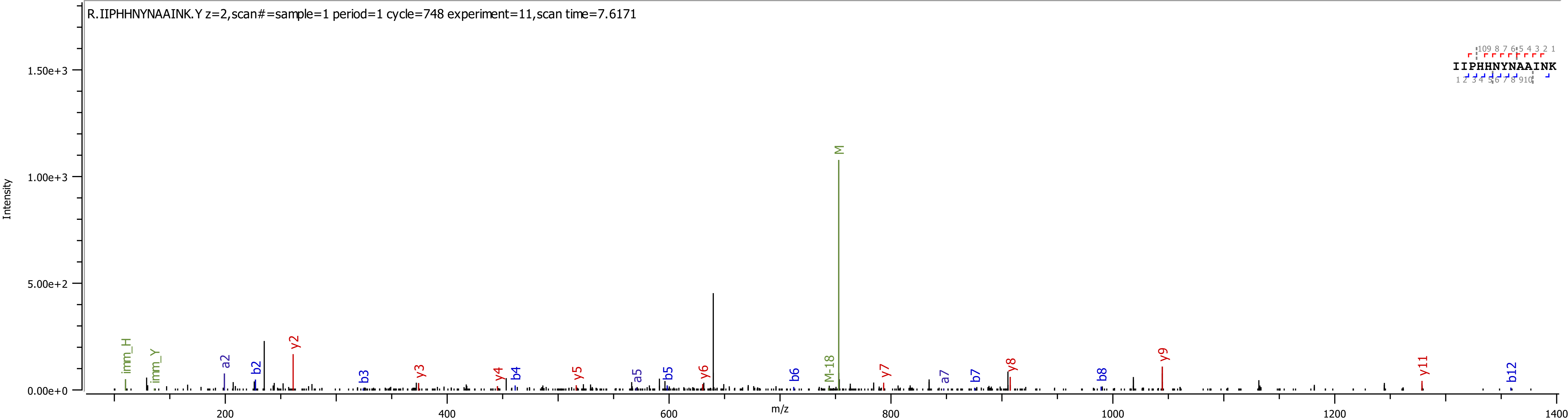
R.IIPHHNYNAAINK.Y z=2,scan#=sample=1 period=1 cycle=748 experiment=11,scan time=7.6171
IIPHHNYNAAINK
Intensity
m/z
M
M-18
imm_H
imm_Y
a2
b2
y2
b3
y3
y4
b4
y5
a5
b5
y6
b6
y7
a7
b7
y8
b8
y9
y11
b12

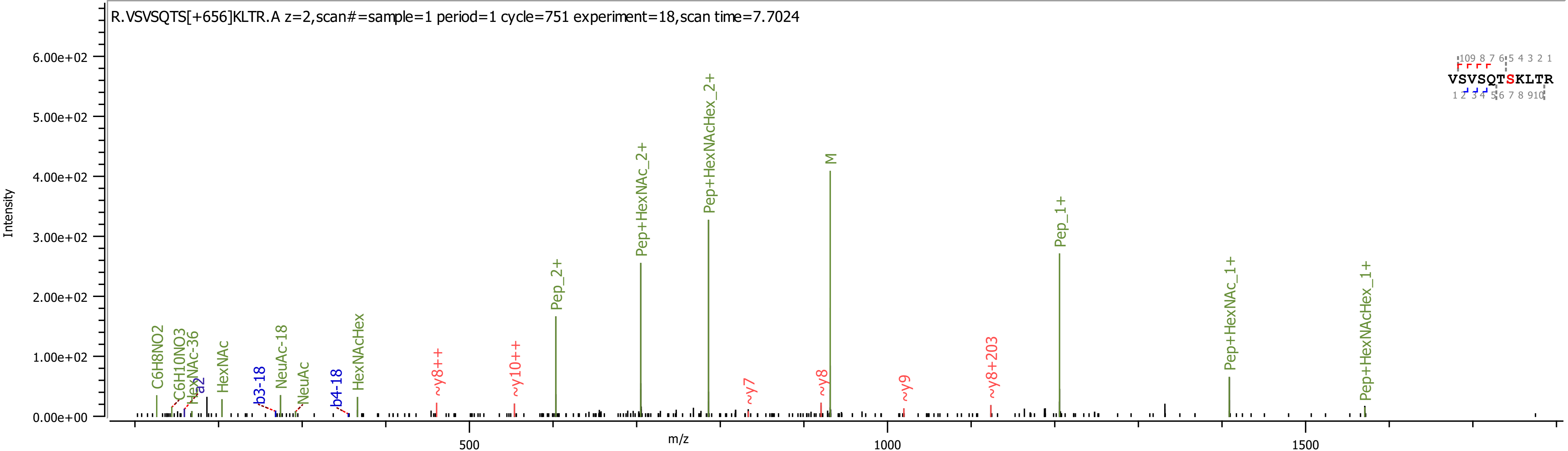
R.VSVSQTS[+656]KLTR.A z=2,scan#=sample=1 period=1 cycle=751 experiment=18,scan time=7.7024
Intensity
m/z
6.00e+02
5.00e+02
4.00e+02
3.00e+02
2.00e+02
1.00e+02
0.00e+00
500
1000
1500
VSVSQTSKLTR
C6H8NO2
C6H10NO3
HexNAc-36
a2
HexNAc
b3-18
NeuAc-18
NeuAc
b4-18
HexNAcHex
~y8++
~y10++
Pep_2+
Pep+HexNAc_2+
Pep+HexNAcHex_2+
~y7
~y8
M
~y9
~y8+203
Pep_1+
Pep+HexNAc_1+
Pep+HexNAcHex_1+

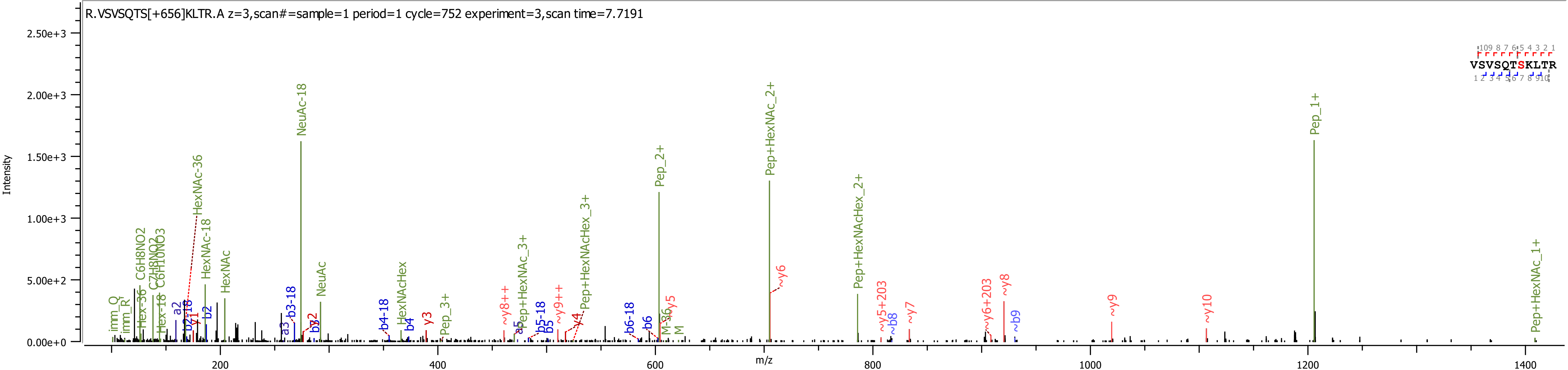
R.VSVSQTS[+656]KLTR.A z=3,scan#=sample=1 period=1 cycle=752 experiment=3,scan time=7.7191
Intensity
m/z
0.00e+0
5.00e+2
1.00e+3
1.50e+3
2.00e+3
2.50e+3
200
400
600
800
1000
1200
1400
VSVSQTSKLTR
NeuAc-18
Pep+HexNAc_2+
Pep_1+
Pep_2+
Pep+HexNAcHex_2+
HexNAc-36
HexNAc-18
HexNAc
NeuAc
HexNAcHex
Pep_3+
Pep+HexNAc_3+
Pep+HexNAcHex_3+
Pep+HexNAc_1+
~y6
~y8
~y9
~y10
~y7
~y5+203
~y6+203
~b8
~b9
~y8++
~y9++
b3-18
b4-18
b5-18
b6-18
b2-18
b2
b3
b4
b5
b6
a2
a3
a5
y1
y2
y3
y4
y5
M-36
M
imm_Q
imm_R
C6H8NO2
C7H8NO2
C6H10NO3
Hex-36
Hex-18

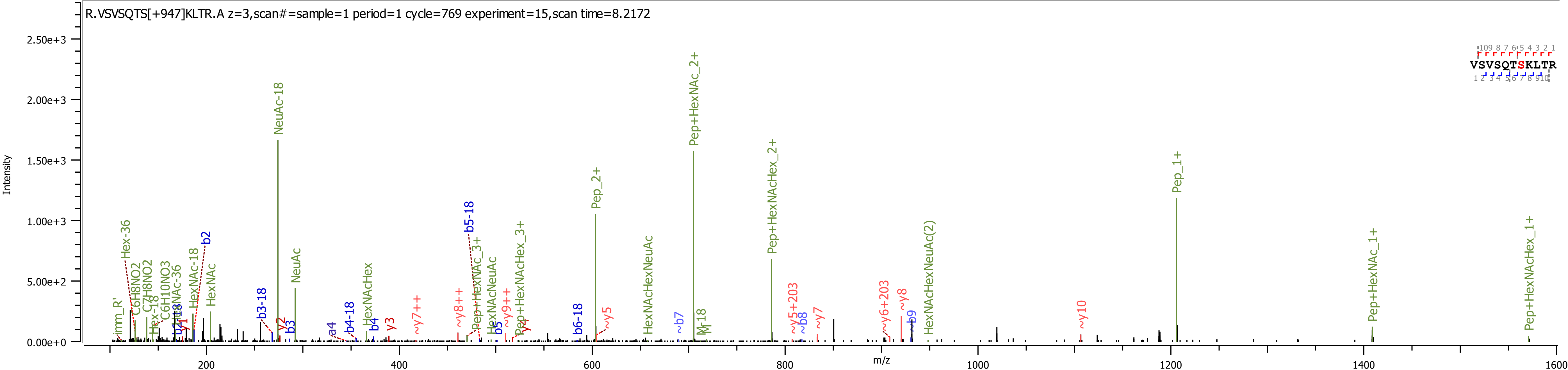
R.VSVSQTS[+947]KLTR.A z=3,scan#=sample=1 period=1 cycle=769 experiment=15,scan time=8.2172
Intensity
m/z
VSVSQTSKLTR
NeuAc-18
Pep+HexNAc_2+
Pep+HexNAcHex_2+
Pep_2+
Pep_1+
Pep+HexNAc_1+
Pep+HexNAcHex_1+
HexNAcHexNeuAc(2)
HexNAcHexNeuAc
Pep+HexNAcHex_3+
Pep+HexNAc_3+
HexNAcNeuAc
HexNAcHex
NeuAc
HexNAc
HexNAc-18
Hex-36
b5-18
b2
b3-18
b4-18
b6-18
~y10
~y8
~y6+203
~y5+203
~y7
~y5
~y9++
~y8++
~y7++
y3
y2
y1
b3
b4
b5
~b7
~b8
b9
a4
M-18
imm_R'

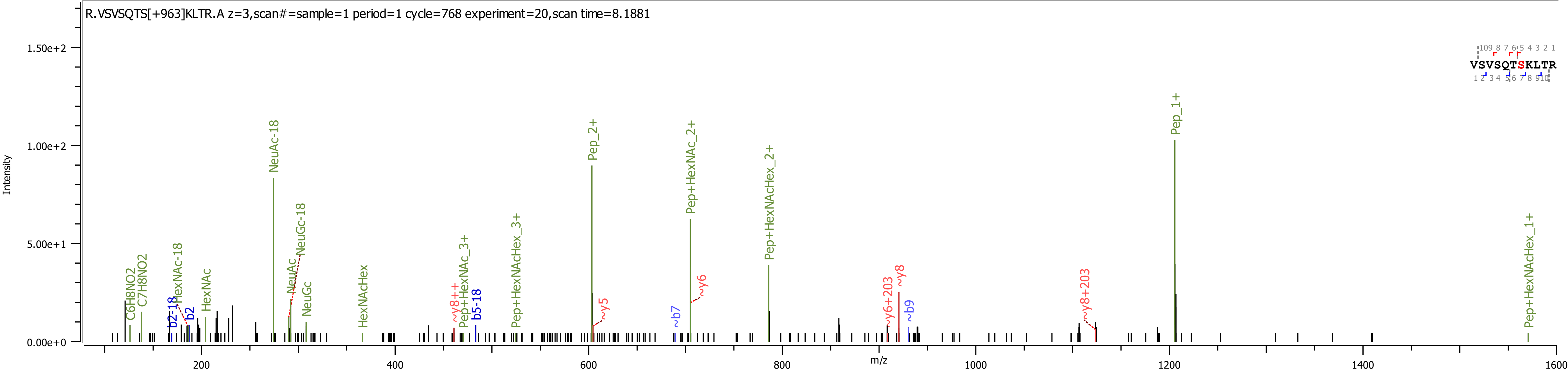

R.VSVSQTS[+963]KLTR.A z=3,scan#=sample=1 period=1 cycle=768 experiment=20,scan time=8.1881
Intensity
m/z
VSVSQTSKLTR
Pep_1+
Pep_2+
NeuAc-18
Pep+HexNAc_2+
NeuGc-18
Pep+HexNAcHex_2+
Pep+HexNAcHex_1+
C6H8NO2
C7H8NO2
HexNAc-18
HexNAc
NeuAc
NeuGc
HexNAcHex
Pep+HexNAc_3+
Pep+HexNAcHex_3+
b2-18
b2
b5-18
~b7
~b9
~y8++
~y5
~y6
~y6+203
~y8
~y8+203

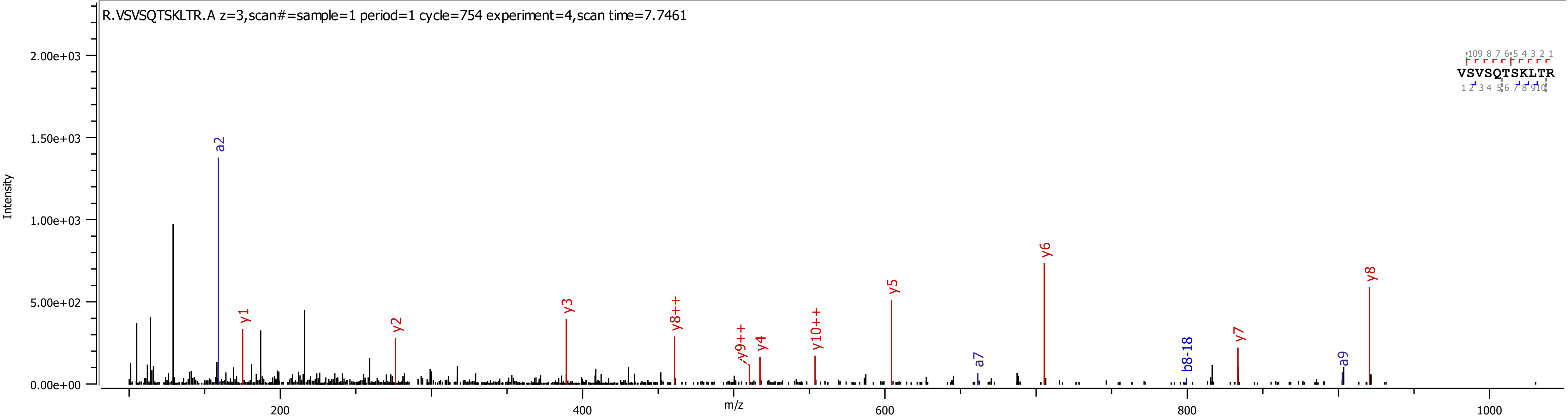

R.VSVSQTSKLTR.A z=3,scan#=sample=1 period=1 cycle=754 experiment=4,scan time=7.7461
Intensity
2.00e+03
1.50e+03
1.00e+03
5.00e+02
0.00e+00
m/z
200
400
600
800
1000
a2
y1
y2
y3
y8++
y9++
y4
y10++
y5
a7
y6
b8-18
y7
a9
y8
VSVSQTSKLTR

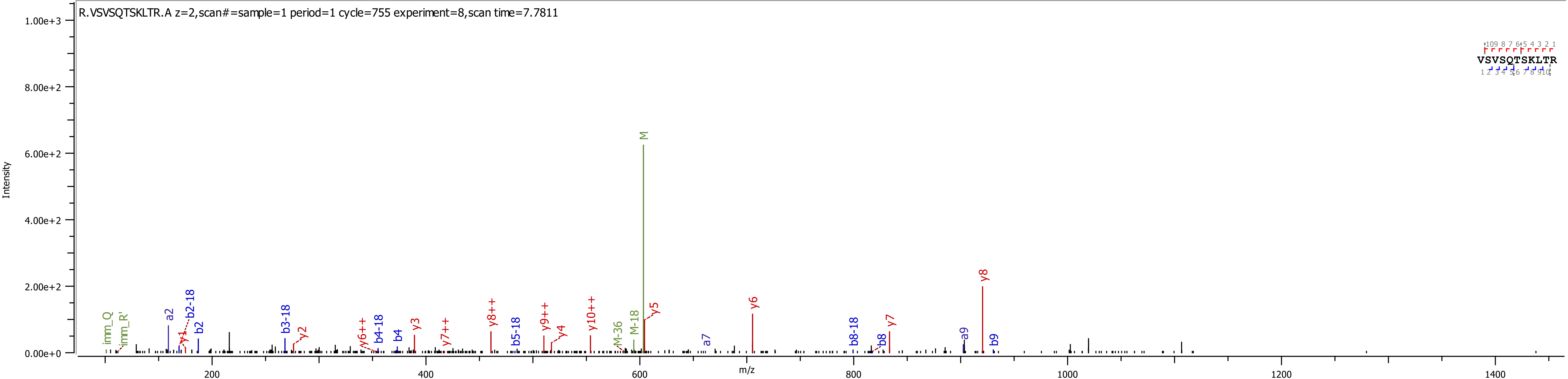
R.VSVSQTSKLTR.A z=2,scan#=sample=1 period=1 cycle=755 experiment=8,scan time=7.7811
VSVSQTSKLTR
Intensity
m/z
M
y8
y6
y7
y5
M-18
M-36
y10++
y9++
y4
y8++
y7++
y6++
b4-18
b4
y3
b5-18
y2
b3-18
a2
b2-18
b2
y1
imm_Q
imm_R'
a7
b8-18
b8
a9
b9

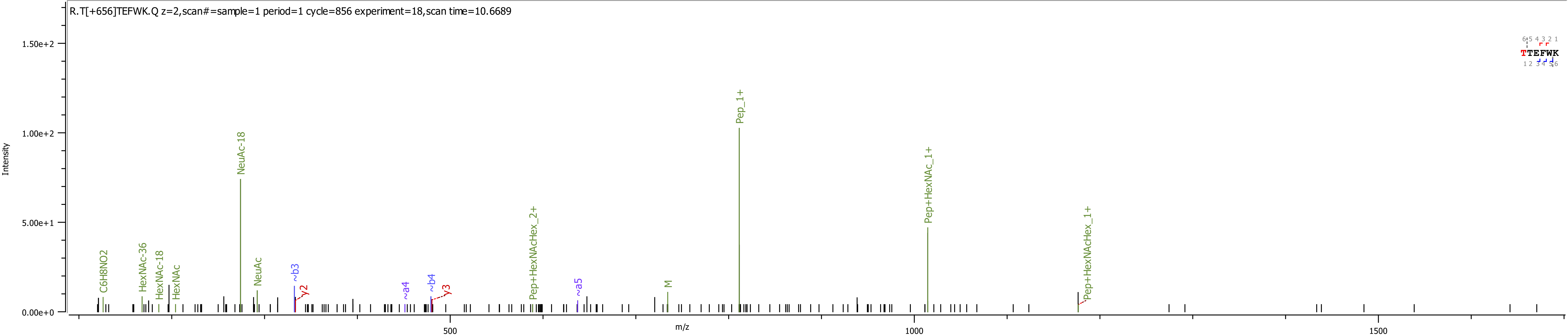
R.T[+656]TEFWK.Q z=2,scan#=sample=1 period=1 cycle=856 experiment=18,scan time=10.6689
Intensity
m/z
0.00e+0
5.00e+1
1.00e+2
1.50e+2
500
1000
1500
C6H8NO2
HexNAc-36
HexNAc-18
HexNAc
NeuAc-18
NeuAc
~b3
y2
~a4
~b4
y3
Pep+HexNAcHex_2+
~a5
M
Pep_1+
Pep+HexNAc_1+
Pep+HexNAcHex_1+
TTEFWK

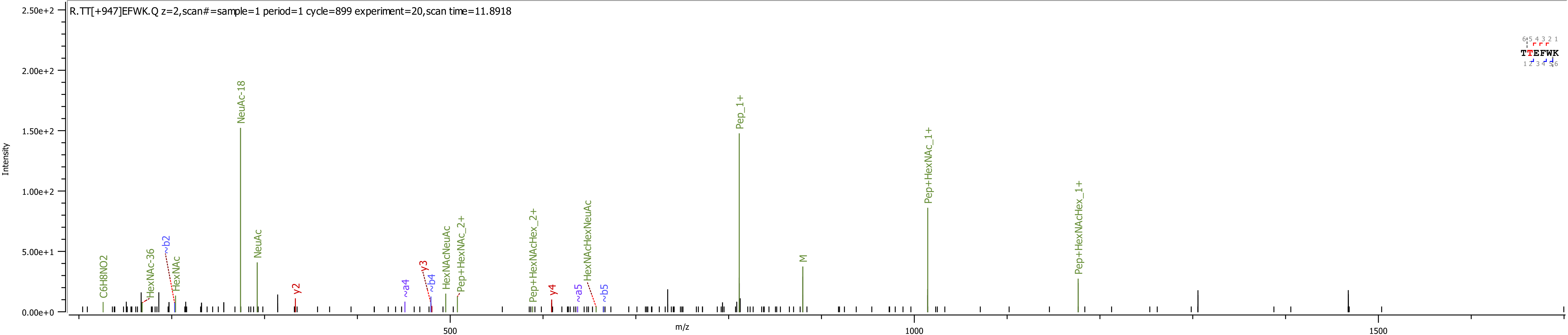
R.TT[+947]EFWK.Q z=2,scan#=sample=1 period=1 cycle=899 experiment=20,scan time=11.8918
Intensity
m/z
TTEFWK
C6H8NO2
HexNAc-36
~b2
HexNAc
NeuAc-18
NeuAc
y2
~a4
y3
~b4
HexNAcNeuAc
Pep+HexNAc_2+
Pep+HexNAcHex_2+
y4
~a5
HexNAcHexNeuAc
~b5
Pep_1+
M
Pep+HexNAc_1+
Pep+HexNAcHex_1+

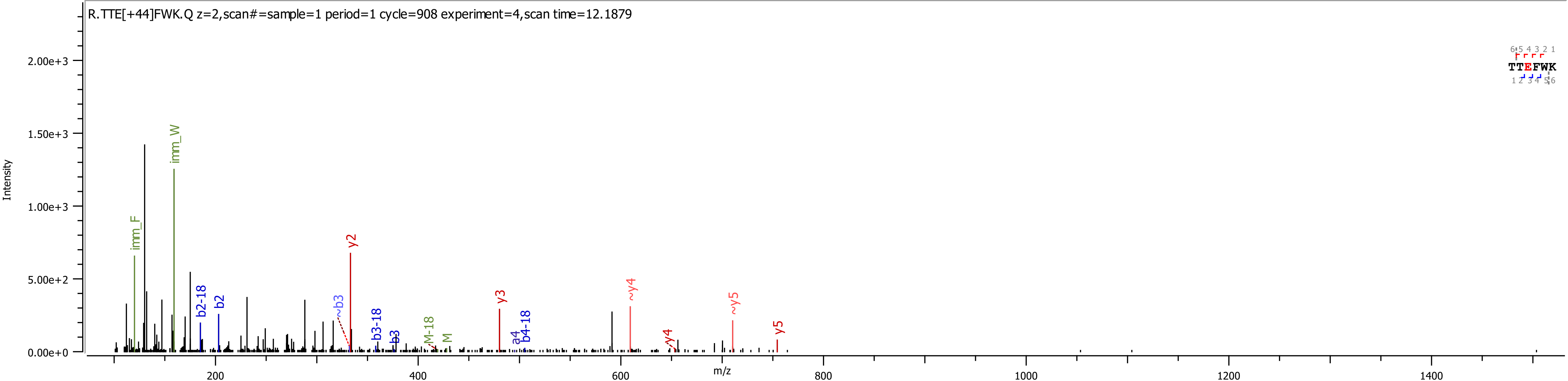
R.TTE[+44]FWK.Q z=2,scan#=sample=1 period=1 cycle=908 experiment=4,scan time=12.1879
Intensity
m/z
imm_F
imm_W
b2-18
b2
~b3
y2
b3-18
b3
M-18
M
y3
a4
b4-18
~y4
y4
~y5
y5
TTEFWK
0.00e+0
5.00e+2
1.00e+3
1.50e+3
2.00e+3
200
400
600
800
1000
1200
1400

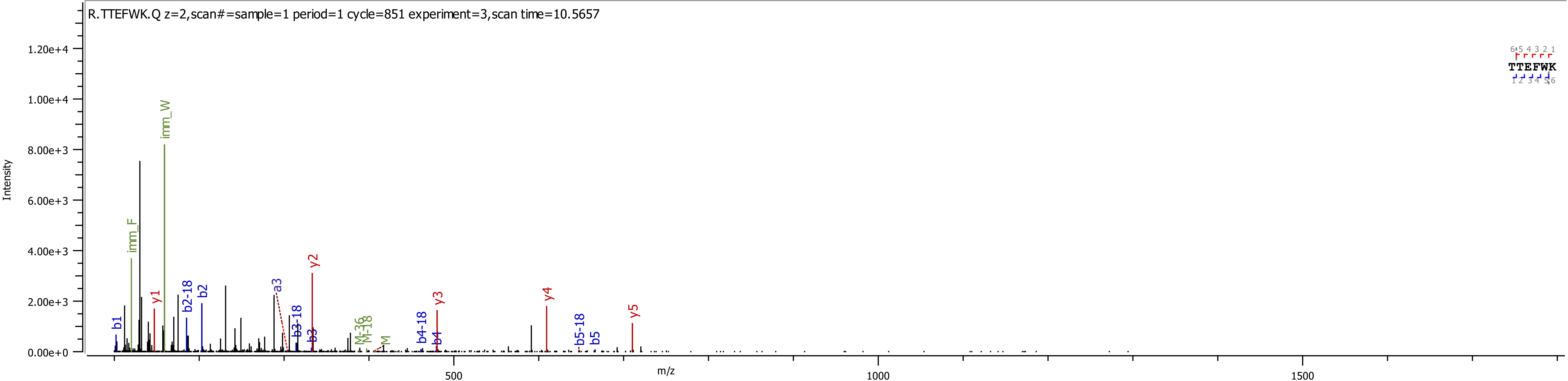
R.TTEFWK.Q z=2,scan#=sample=1 period=1 cycle=851 experiment=3,scan time=10.5657
Intensity
1.20e+4
1.00e+4
8.00e+3
6.00e+3
4.00e+3
2.00e+3
0.00e+0
m/z
500
1000
1500
imm_W
imm_F
b1
y1
b2-18
b2
a3
b3-18
b3
y2
M-36
M-18
M
b4-18
b4
y3
y4
b5-18
b5
y5
TTEFWK

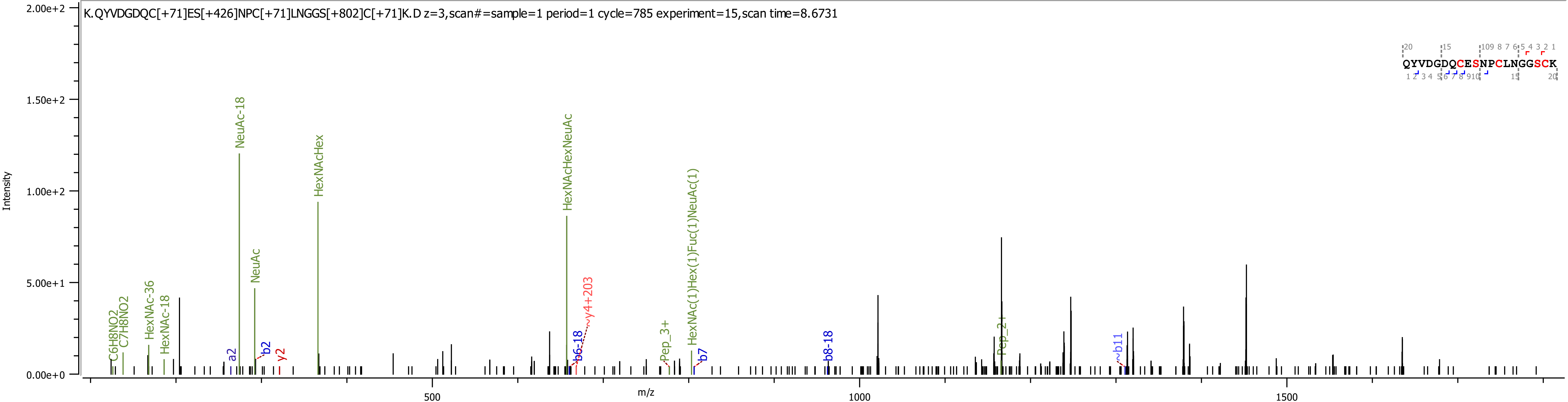
K.QYVDGDQC[+71]ES[+426]NPC[+71]LNGGS[+802]C[+71]K.D z=3,scan#=sample=1 period=1 cycle=785 experiment=15,scan time=8.6731
QYVDGDQCESNPCLNGGSCK
Intensity
m/z
2.00e+2
1.50e+2
1.00e+2
5.00e+1
0.00e+0
500
1000
1500
C6H8NO2
C7H8NO2
HexNAc-36
HexNAc-18
a2
NeuAc-18
NeuAc
b2
y2
HexNAcHex
HexNAcHexNeuAc
b6-18
y4+203
Pep_3+
HexNAc(1)Hex(1)Fuc(1)NeuAc(1)
b7
b8-18
Pep_2+
b11
