## Supplementary Information 6 for "Analysis of coagulation factor IX in bioreactor cell culture medium predicts yield and quality of the purified product"

K.[+14.016]LE[+43.990]E[+14.061]FVQGNLE[+14.016]R.E 1

K.[+14.016]LE[+43.990]E[+43.990]FVQGNLE[+14.016]R.E 2

K.[+14.016]LE[+43.990]E[+43.990]FVQGNLE[+43.990]R.E 3

K.[+14.016]LE[+43.990]E[+43.990]FVQGNLE[+58.005]R.E 4

K.[+14.016]LE[+43.990]E[+43.990]FVQGNLE[+72.021]R.E 5

K.[+14.016]LE[+43.990]E[+58.005]FVQGN[+0.984]LER.E 6
K.[+14.016]LE[+43.990]E[+58.005]FVQGNLE[+14.016]R.E 7
K.[+14.016]LE[+43.990]E[+58.005]FVQGNLE[+43.990]R.E 8
K.[+14.016]LE[+43.990]E[+58.005]FVQGNLE[+58.005R.E 9
K.[+14.016]LE[+43.990]E[+58.005]FVQGNLE[+72.021]R.E 10

K.[+14.016]LE[+43.990]E[+58.005]FVQGNLER.E 11

K.[+14.016]LE[+43.990]E[+72.021]FVQGNLE[+72.021]R.E 12

K.[+14.016]LE[+58.005]E[+43.990]FVQGNLE[+14.016]R.E 13

K.[+14.016]LE[+58.005]E[+43.990]FVQGNLE[+72.021]R.E 14

K.[+14.016]LE[+58.005]E[+58.005]FVQGNLE[+14.016]R.E 15

K.[+14.016]LE[+58.005]E[+58.005]FVQGNLE[+72.021]R.E 16
K.[+14.016]LE[+58.005]E[+72.021]FVQGNLE[+14.016]R.E 17
K.[+14.016]LE[+58.005]E[+72.021]FVQGNLE[+72.021]R.E 18
K.[+14.016]LE[+58.005]E[+72.021]FVQGNLER.E 19
K.[+14.016]LE[+72.021]E[+72.021]FVQGNLE[+58.005]R.E 20

K.[+14.016]LEE[+14.016]FVQGNLE[+14.016]R.E 21

K.[+14.016]LEE[+43.990]FVQGNLER.E 22

K.[+14.016]LEE[+58.005]FVQGN[+0.984]LE[+14.016]R.E 23

K.[+14.016]LEE[+58.005]FVQGNLE[+43.990]R.E 24

K.C[+71.037]SFE[+43.990]E[+14.061]AR.E 25

K.C[+71.037]SFE[+43.990]E[+58.005]AR.E 26
K.C[+71.037]SFE[+43.990]E[+58.005]AR[+28.031].E 27
K.C[+71.037]SFE[+43.990]E[+72.021]AR.E 28
K.C[+71.037]SFE[+58.005]E[+14.061]AR.E 29
K.C[+71.037]SFE[+58.005]E[+58.005]AR.E 30

K.C[+71.037]SFE[+58.005]E[+72.021]AR.E 31
K.C[+71.037]SFE[+72.021]E[+43.990]AR.E 32
K.C[+71.037]SFE[+72.021]E[+58.005]AR.E 33
K.C[+71.037]SFEE[+72.021]AR.E 34
K.LE[+14.016]E[+43.990]FVQGNLE[+14.016]R.E 35

K.LE[+14.016]E[+58.005]FVQGNLE[+14.016]R.E 36
K.LE[+14.016]E[+58.005]FVQGNLER.E 37
K.LE[+14.016]EFVQGNLE[+14.016]R.E 38
K.LE[+43.990]E[+43.990]FVQGNLE[+58.005]R.E 39
K.LE[+43.990]E[+58.005]FVQGN[+14.016]LE[+43.990]R.E 40

K.LE[+43.990]E[+58.005]FVQGNLE[+58.005]R.E 41

K.LE[+43.990]E[+72.021]FVQGNLE[+14.016]R.E 42

K.LE[+58.005]E[+14.016]FVQGN[+14.016]LE[+43.990]R.E 43

K.LE[+58.005]E[+14.016]FVQGNLE[+14.016]R.E 44

K.LE[+58.005]E[+14.016]FVQGNLE[+43.990]R.E 45

K.LE[+58.005]E[+14.016]FVQGNLER.E 46

K.LE[+58.005]E[+43.990]FVQGN[+14.016]LE[+43.990]R.E 47

K.LE[+58.005]E[+43.990]FVQGN[+14.061]LE[+43.990]R.E 48

K.LE[+58.005]E[+43.990]FVQGN[+14.061]LE[+58.005]R.E 49

K.LE[+58.005]E[+43.990]FVQGNLE[+14.016]R.E 50

K.LE[+58.005]E[+43.990]FVQGNLE[+43.990]R.E 51
K.LE[+58.005]E[+43.990]FVQGNLE[+43.990]R[+14.016].E 52
K.LE[+58.005]E[+43.990]FVQGNLE[+72.021]R.E 53
K.LE[+58.005]E[+43.990]FVQGNLER.E 54
K.LE[+58.005]E[+58.005]FVQ[+0.984]GNLE[+14.016]R.E 55

K.LE[+58.005]E[+58.005]FVQ[+0.984]GNLE[+58.005]R.E 56
K.LE[+58.005]E[+58.005]FVQ[+0.984]GNLE[+72.021]R.E 57
K.LE[+58.005]E[+58.005]FVQGN[+0.984]LER.E 58
K.LE[+58.005]E[+58.005]FVQGN[+14.061]LE[+43.990]R.E 59
K.LE[+58.005]E[+58.005]FVQGN[+14.061]LE[+58.005]R.E 60

K.LE[+58.005]E[+58.005]FVQGNLE[+14.016]R.E 61
K.LE[+58.005]E[+58.005]FVQGNLE[+43.990]R.E 62
K.LE[+58.005]E[+58.005]FVQGNLE[+43.990]R[+14.016].E 63
K.LE[+58.005]E[+58.005]FVQGNLE[+58.005]R.E 64
K.LE[+58.005]E[+58.005]FVQGNLE[+72.021]R.E 65

K.LE[+58.005]E[+58.005]FVQGNLER.E 66
K.LE[+58.005]E[+72.021]FVQGN[+0.984]LE[+14.016]R.E 67
K.LE[+58.005]E[+72.021]FVQGN[+14.061]LE[+43.990]R.E 68
K.LE[+58.005]E[+72.021]FVQGNLE[+14.016]R.E 69
K.LE[+58.005]E[+72.021]FVQGNLE[+43.990]R[+14.061].E 70

K.LE[+58.005]E[+72.021]FVQGNLE[+58.005]R.E 71

K.LE[+58.005]E[+72.021]FVQGNLE[+72.021]R.E 72

K.LE[+58.005]EFVQGN[+14.016]LE[+43.990]R.E 73

K.LE[+72.021]E[+43.990]FVQ[+0.984]GNLE[+72.021]R.E 74

K.LE[+72.021]E[+43.990]FVQGN[+14.061]LE[+43.990]R.E 75

K.LE[+72.021]E[+43.990]FVQGNLE[+14.061]R.E 76
K.LE[+72.021]E[+43.990]FVQGNLE[+43.990]R[+14.016].E 77
K.LE[+72.021]E[+43.990]FVQGNLE[+58.005]R.E 78
K.LE[+72.021]E[+43.990]FVQGNLE[+72.021]R.E 79
K.LE[+72.021]E[+43.990]FVQGNLER.E 80

K.LE[+72.021]E[+58.005]FVQGN[+14.016]LE[+43.990]R.E 81
K.LE[+72.021]E[+58.005]FVQGNLE[+14.061]R.E 82
K.LE[+72.021]E[+58.005]FVQGNLE[+43.990]R[+14.016].E 83
K.LE[+72.021]E[+58.005]FVQGNLE[+58.005]R.E 84
K.LE[+72.021]E[+58.005]FVQGNLE[+72.021]R.E 85

K.LE[+72.021]E[+72.021]FVQGN[+14.016]LE[+43.990]R.E 86

K.LE[+72.021]E[+72.021]FVQGNLE[+14.061]R.E 87

K.LE[+72.021]E[+72.021]FVQGNLE[+43.990]R[+14.016].E 88

K.LE[+72.021]E[+72.021]FVQGNLE[+58.005]R.E 89

K.LE[+72.021]E[+72.021]FVQGNLE[+72.021]R.E 90

K.LEE[+14.016]FVQGNLER.E 91
K.LEE[+43.990]FVQGNLE[+14.016]R.E 92
K.LEE[+58.005]FVQGNLE[+14.016]R.E 93
K.LEE[+58.005]FVQGNLER.E 94
K.LEE[+72.021]FVQGNLER.E 95

R.[+14.016]T[+656.228]TEFWK.Q 96
R.[+14.016]T[+947.323]TEFWK.Q 97
R.[+14.016]T[+947.323]TEFWK[+14.016].Q 98
R.[+14.016]TTE[+43.990]FWK.Q 99
R.[+14.016]TTEFWK.Q 100

R.[+14016]T[+656.228]TEFWK[+14.016].Q 101

R.E[-18.010]VFE[+14.016]N[+14.016]TE[+43.990]R.T 102

R.E[-18.010]VFE[+14.016]NTE[+14.016]R.T 103

R.E[-18.010]VFE[+43.990]N[+14.016]TE[+43.990]R.T 104

R.E[-18.010]VFE[+43.990]N[+14.016]TE[+58.005]R.T 105

R.E[-18.010]VFE[+43.990]NTE[+43.990]R[+14.061].T 106
R.E[-18.010]VFE[+43.990]NTE[+58.005]R.T 107
R.E[-18.010]VFE[+58.005]N[+14.016]TE[+43.990]R.T 108
R.E[-18.010]VFE[+58.005]NTE[+14.016]R.T 109
R.E[-18.010]VFE[+58.005]NTE[+43.990]R[+14.016].T 110

R.E[-18.010]VFE[+72.021]N[+14.061]TE[+43.990]R.T 111
R.E[-18.010]VFE[+72.021]NTE[+43.990]R.T 112
R.E[-18.010]VFE[+72.021]NTER.T 113
R.E[-18.010]VFENTE[+14.016]R.T 114
R.E[-18.011]VFE[+58.005]NTE[+43.990]R.T 115

R.E[-18.011]VFE[+72.021]NTE[+43.990]R[+14.016].T 116
R.E[+43.990]VFE[+58.005]NTE[+14.016]R.T 117
R.TT[+656.228]E[+14.016]FWK.Q 118
R.TT[+656.228]EFWK.Q 119
R.TT[+947.323]E[+14.016]FWK.Q 120

R.TTE[+14.016]FWK.Q 121
R.TTE[+43.990]FWK.Q 122
R.TTE[+58.005]FWK.Q 123
R.TTE[+72.021]FWK.Q 124
R.TTEFWK.Q 125

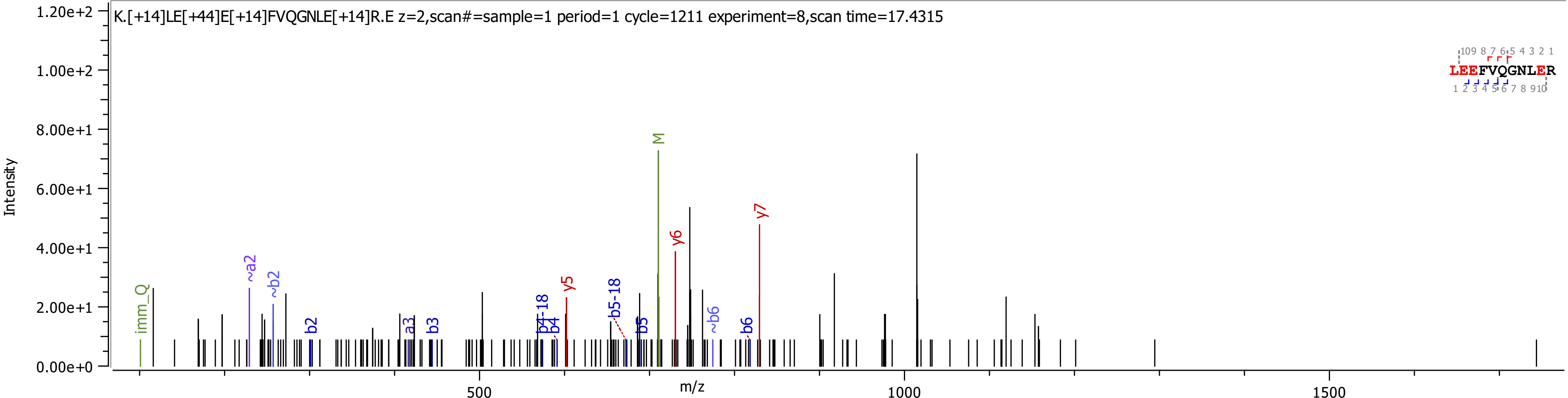

K.[+14]LE[+44]E[+14]FVQGNLE[+14]R.E z=2,scan#=sample=1 period=1 cycle=1211 experiment=8,scan time=17.4315
Intensity
m/z
LEEFVQGNLER
imm_Q
~a2
~b2
b2
a3
b3
b4-18
b4
y5
b5-18
b5
M
y6
~b6
b6
y7

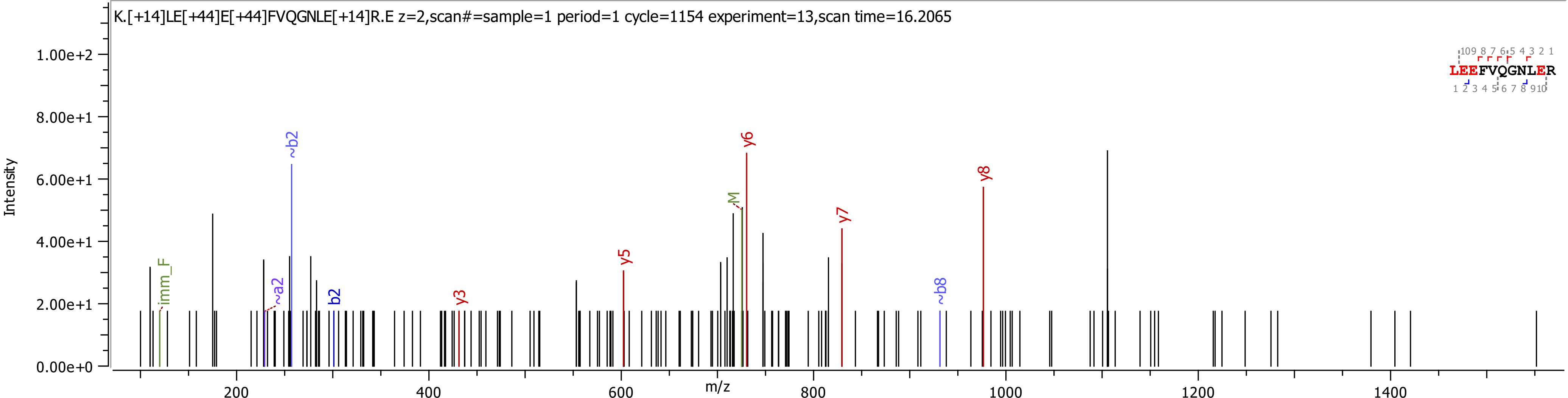
K.[+14]LE[+44]E[+44]FVQGNLE[+14]R.E z=2,scan#=sample=1 period=1 cycle=1154 experiment=13,scan time=16.2065
Intensity
m/z
1.00e+2
8.00e+1
6.00e+1
4.00e+1
2.00e+1
0.00e+0
200
400
600
800
1000
1200
1400
imm_F
~a2
~b2
b2
y3
y5
M
y6
y7
~b8
y8
LEEFVQGNLER

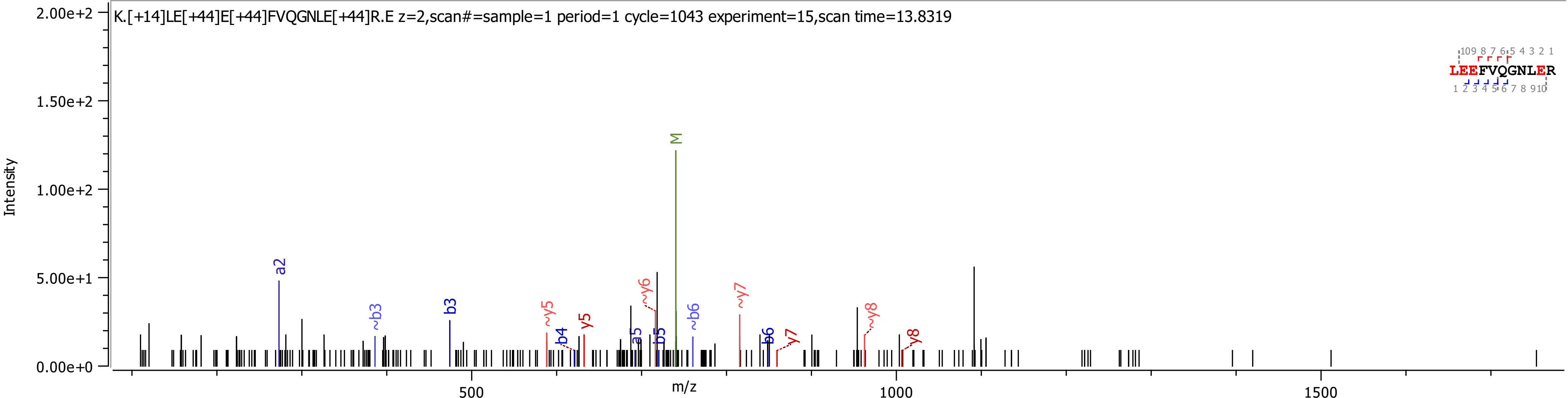
K.[+14]LE[+44]E[+44]FVQGNLE[+44]R.E z=2,scan#=sample=1 period=1 cycle=1043 experiment=15,scan time=13.8319
Intensity
m/z
2.00e+2
1.50e+2
1.00e+2
5.00e+1
0.00e+0
500
1000
1500
a2
~b3
b3
~y5
b4
y5
a5
b5
~y6
M
~b6
~y7
b6
y7
~y8
y8
LEEFVQGNLER

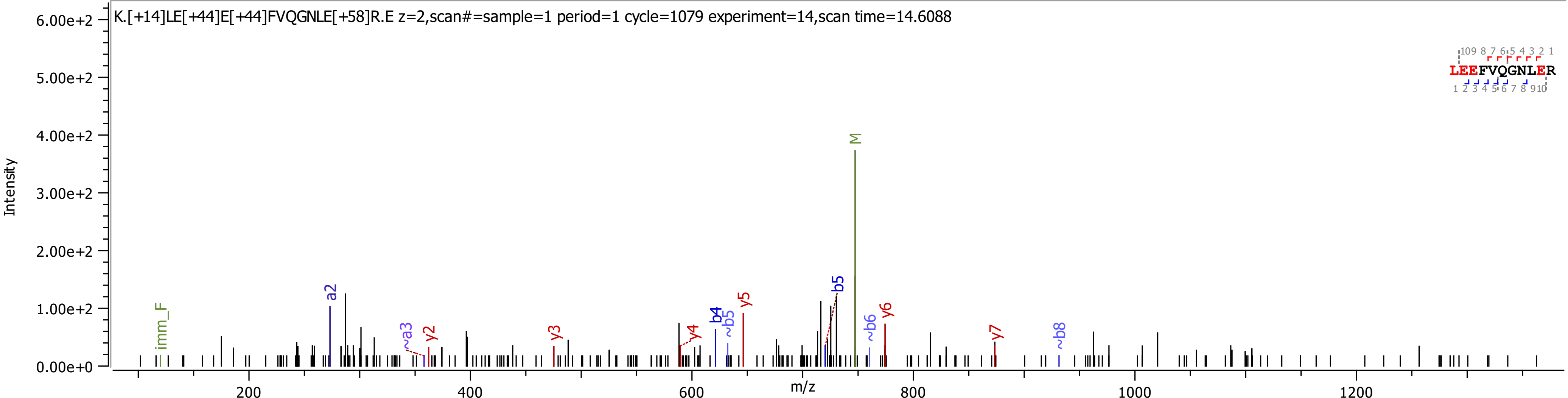
K.[+14]LE[+44]E[+44]FVQGNLE[+58]R.E z=2,scan#=sample=1 period=1 cycle=1079 experiment=14,scan time=14.6088
Intensity
6.00e+2
5.00e+2
4.00e+2
3.00e+2
2.00e+2
1.00e+2
0.00e+0
m/z
200
400
600
800
1000
1200
imm_F
a2
~a3
y2
y3
y4
b4
~b5
y5
b5
M
~b6
y6
y7
~b8
LEEFVQGNLER
109 8 7 6 5 4 3 2 1
1 2 3 4 5 6 7 8 910

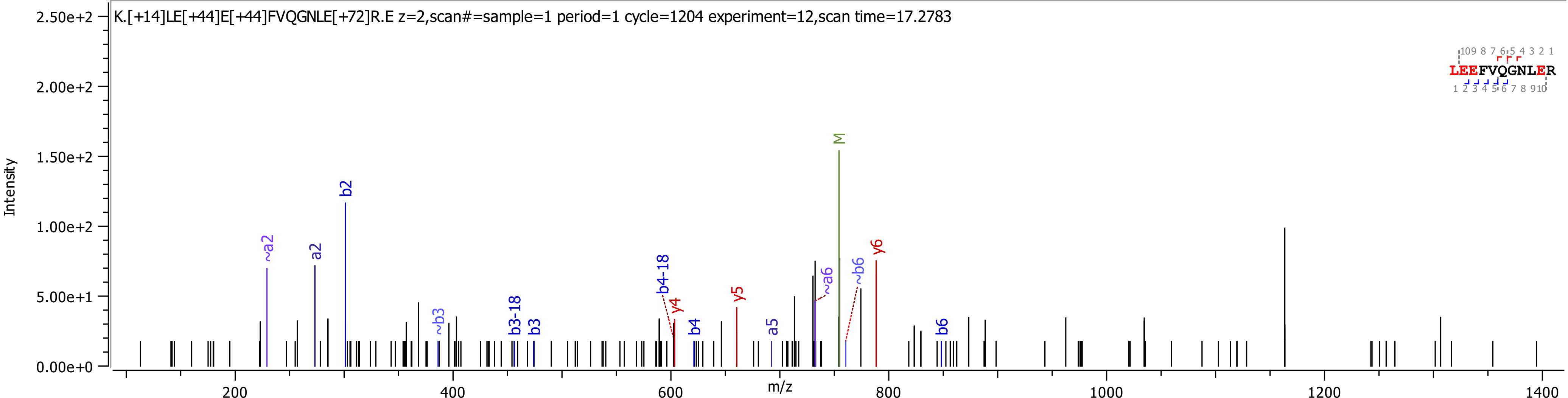
K.[+14]LE[+44]E[+44]FVQGNLE[+72]R.E z=2,scan#=sample=1 period=1 cycle=1204 experiment=12,scan time=17.2783
Intensity
m/z
LEEFVQGNLER
~a2
a2
b2
~b3
b3-18
b3
b4-18
y4
b4
y5
a5
~a6
M
~b6
y6
b6

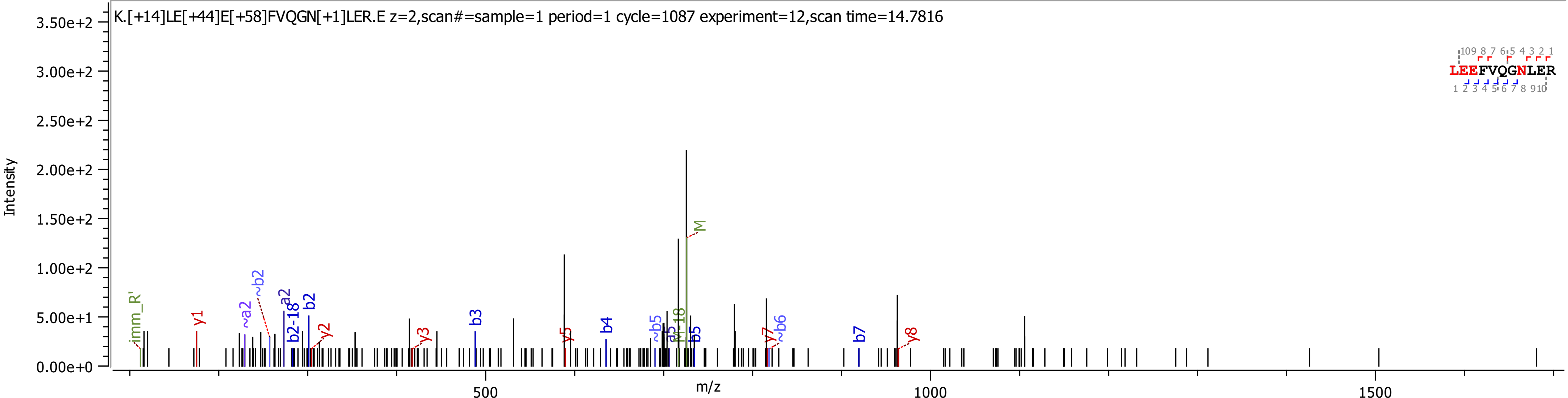
K.[+14]LE[+44]E[+58]FVQGN[+1]LER.E z=2,scan#=sample=1 period=1 cycle=1087 experiment=12,scan time=14.7816
Intensity
3.50e+2
3.00e+2
2.50e+2
2.00e+2
1.50e+2
1.00e+2
5.00e+1
0.00e+0
m/z
500
1000
1500
LEEFVQGNLER
imm_R'
y1
~a2
~b2
a2
b2-18
b2
y2
y3
b3
y5
b4
~b5
b5
M
y7
~b6
b7
y8

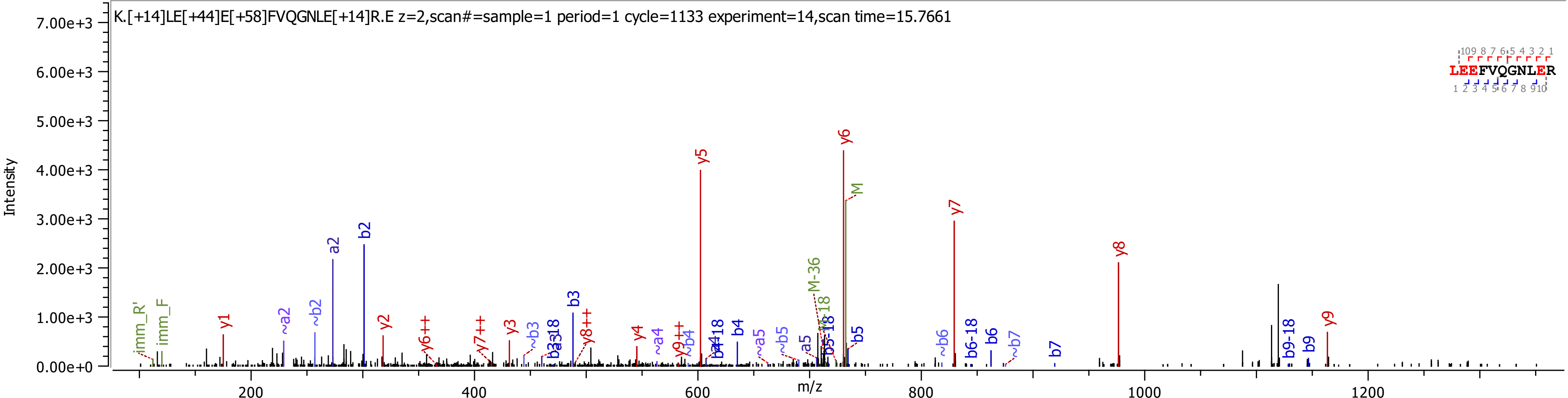

K.[+14]LE[+44]E[+58]FVQGNLE[+14]R.E z=2,scan#=sample=1 period=1 cycle=1133 experiment=14,scan time=15.7661
Intensity
m/z
LEEFVQGNLER
y1
y2
y3
y4
y5
y6
y7
y8
y9
a2
b2
b3
b4
b5
b6
b7
b9
M
M-36

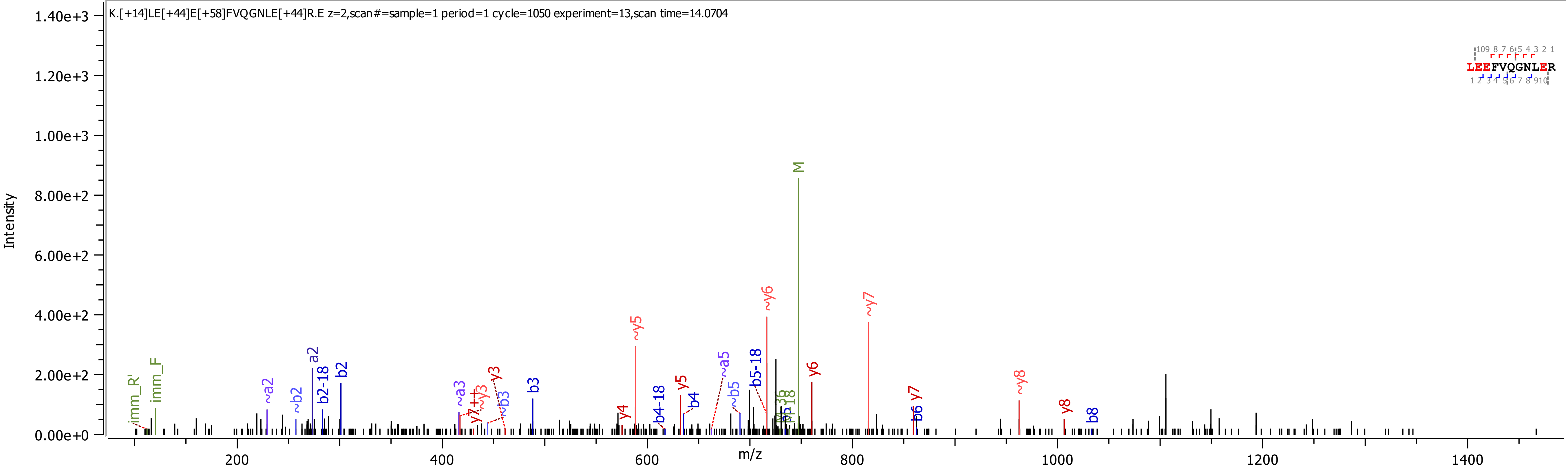
K.[+14]LE[+44]E[+58]FVQGNLE[+44]R.E z=2,scan#=sample=1 period=1 cycle=1050 experiment=13,scan time=14.0704
Intensity
m/z
LEEFVQGNLER
M
~y6
~y7
~y5
y6
y5
y4
~y8
y8
y7
b6
b8
b3
b2
a2
b2-18
b4-18
b4
b5-18
~b5
~a5
~a2
~b2
~a3
~y3
y3
~b3
y7++
imm_R'
imm_F

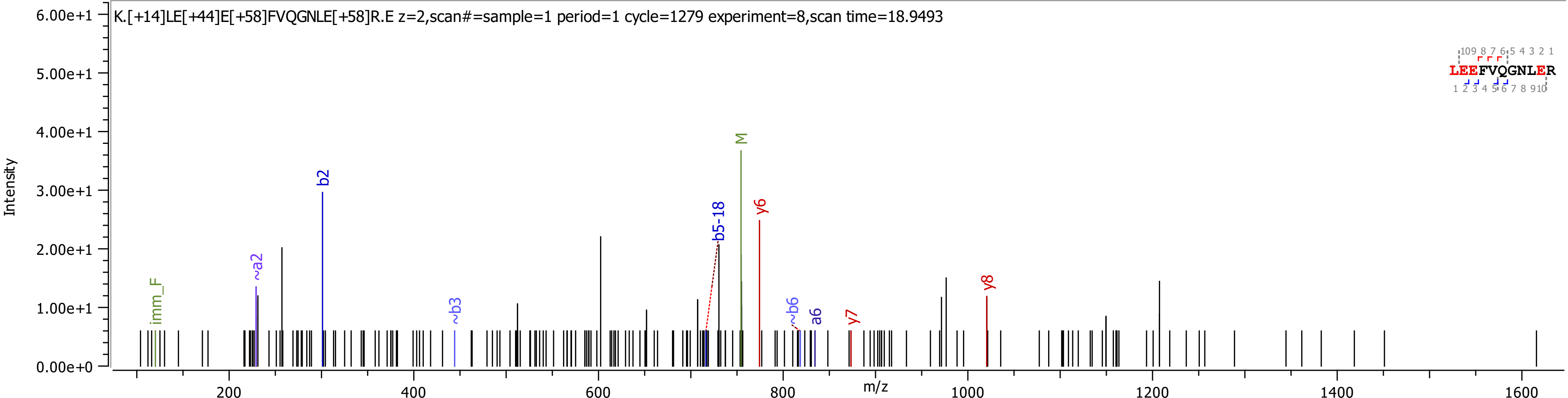

K.[+14]LE[+44]E[+58]FVQGNLE[+58]R.E z=2,scan#=sample=1 period=1 cycle=1279 experiment=8,scan time=18.9493
Intensity
6.00e+1
5.00e+1
4.00e+1
3.00e+1
2.00e+1
1.00e+1
0.00e+0
m/z
200
400
600
800
1000
1200
1400
1600
imm_F
~a2
b2
~b3
b5-18
M
y6
~b6
a6
y7
y8
LEEFVQGNLER

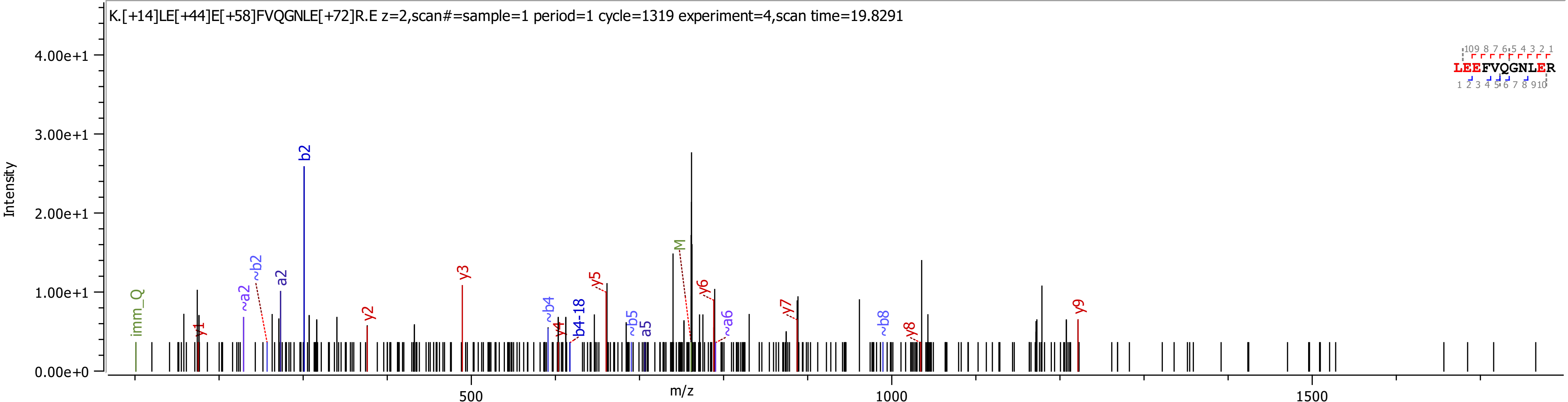
K.[+14]LE[+44]E[+58]FVQGNLE[+72]R.E z=2,scan#=sample=1 period=1 cycle=1319 experiment=4,scan time=19.8291
Intensity
4.00e+1
3.00e+1
2.00e+1
1.00e+1
0.00e+0
m/z
500
1000
1500
imm_Q
y1
~a2
~b2
a2
b2
y2
y3
~b4
y4
b4-18
y5
~b5
a5
M
y6
~a6
y7
~b8
y8
y9
LEEFVQGNLER

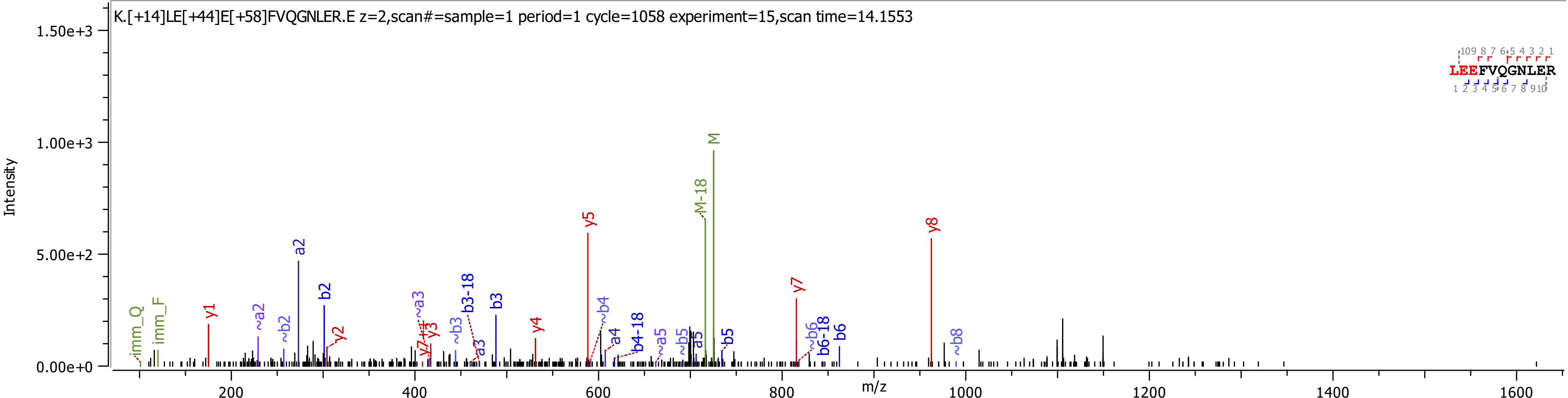

K.[+14]LE[+44]E[+58]FVQGNLER.E z=2,scan#=sample=1 period=1 cycle=1058 experiment=15,scan time=14.1553
Intensity
m/z
LEEFVQGNLER
imm_Q
imm_F
y1
~a2
~b2
a2
b2
y2
y7++
y3
~a3
~b3
a3
b3-18
b3
y4
y5
~b4
a4
b4-18
~a5
~b5
a5
b5
M-18
M
y7
~b6
b6-18
b6
y8
~b8

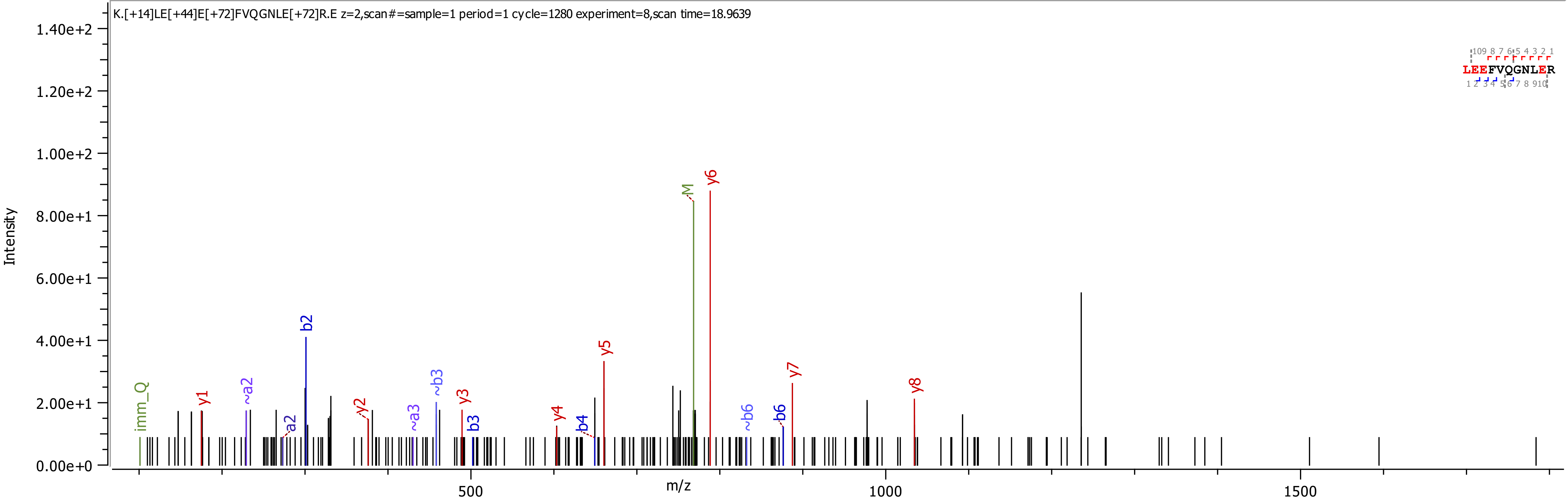
K.[+14]LE[+44]E[+72]FVQGNLE[+72]R.E z=2,scan#=sample=1 period=1 cycle=1280 experiment=8,scan time=18.9639
Intensity
1.40e+2
1.20e+2
1.00e+2
8.00e+1
6.00e+1
4.00e+1
2.00e+1
0.00e+0
m/z
500
1000
1500
imm_Q
y1
~a2
a2
b2
y2
~a3
~b3
y3
b3
y4
b4
y5
M
y6
~b6
b6
y7
y8
LEEFVQGNLER

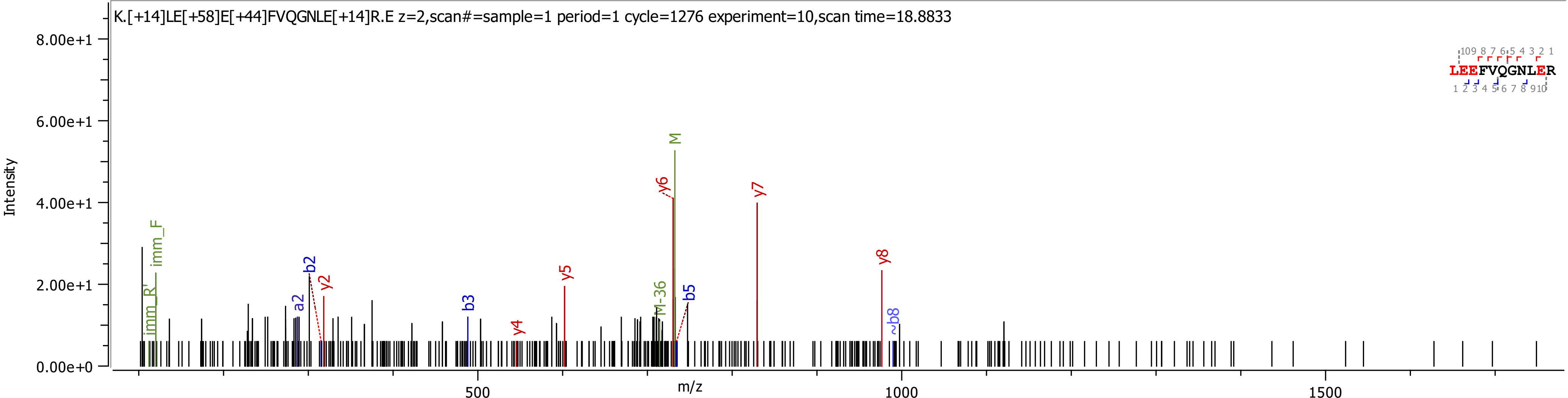
K.[+14]LE[+58]E[+44]FVQGNLE[+14]R.E z=2,scan#=sample=1 period=1 cycle=1276 experiment=10,scan time=18.8833
Intensity
m/z
8.00e+1
6.00e+1
4.00e+1
2.00e+1
0.00e+0
500
1000
1500
imm_R'
imm_F
a2
b2
y2
b3
y4
y5
M-36
y6
M
b5
y7
y8
~b8
LEEFVQGNLER

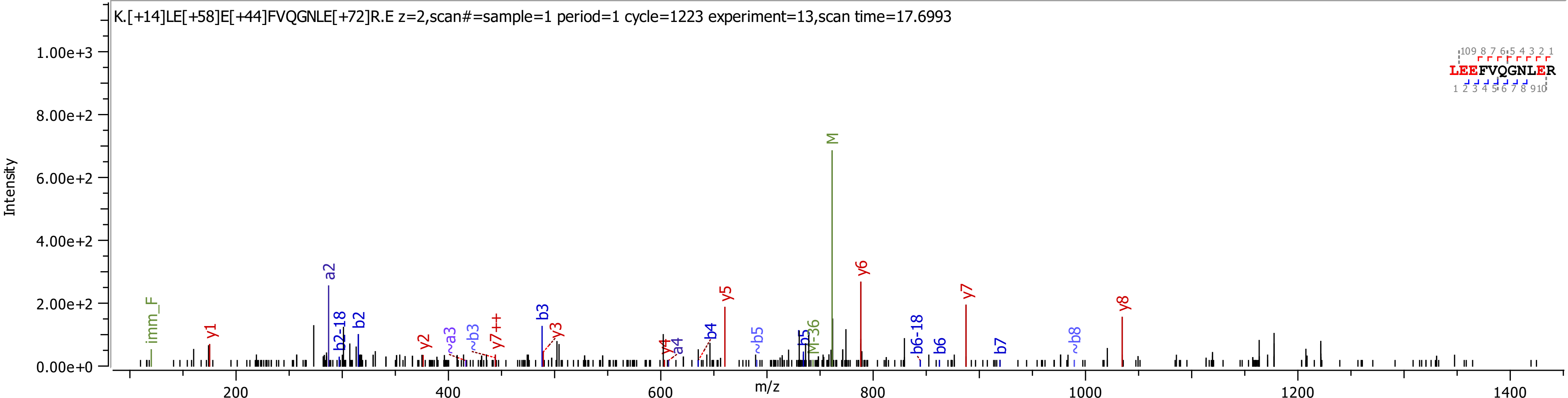

K.[+14]LE[+58]E[+44]FVQGNLE[+72]R.E z=2,scan#=sample=1 period=1 cycle=1223 experiment=13,scan time=17.6993
Intensity
m/z
1.00e+3
8.00e+2
6.00e+2
4.00e+2
2.00e+2
0.00e+0
200
400
600
800
1000
1200
1400
imm_F
y1
a2
b2-18
b2
y2
~a3
~b3
y7++
b3
y3
y4
a4
b4
y5
~b5
b5
M-36
M
y6
b6-18
b6
y7
b7
~b8
y8
LEEFVQGNLER

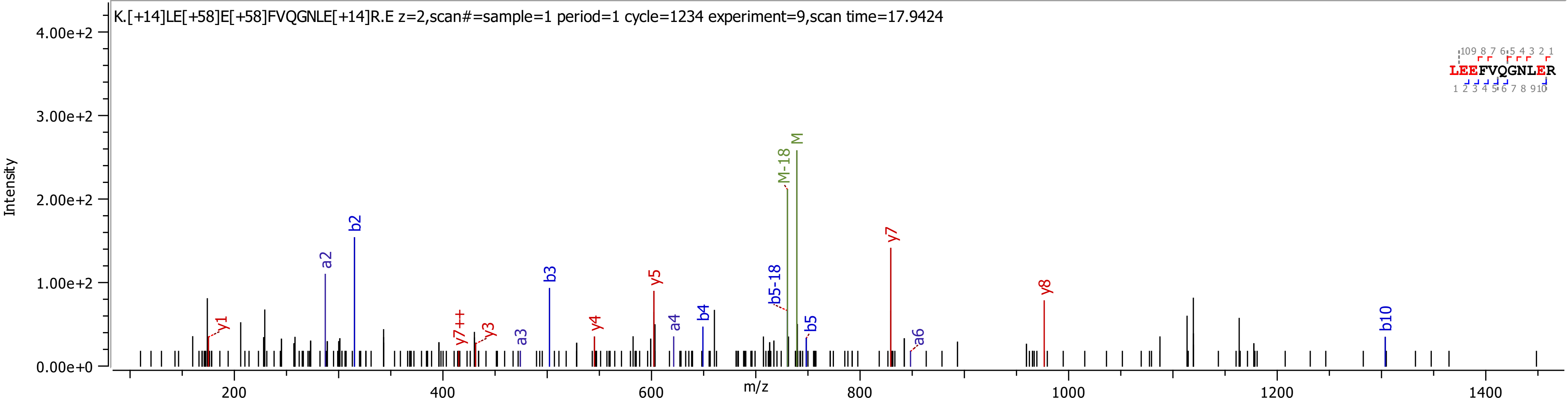
K.[+14]LE[+58]E[+58]FVQGNLE[+14]R.E z=2,scan#=sample=1 period=1 cycle=1234 experiment=9,scan time=17.9424
Intensity
m/z
LEEFVQGNLER
y1
a2
b2
y7++
y3
a3
b3
y4
y5
a4
b4
b5-18
M-18
M
b5
y7
a6
y8
b10

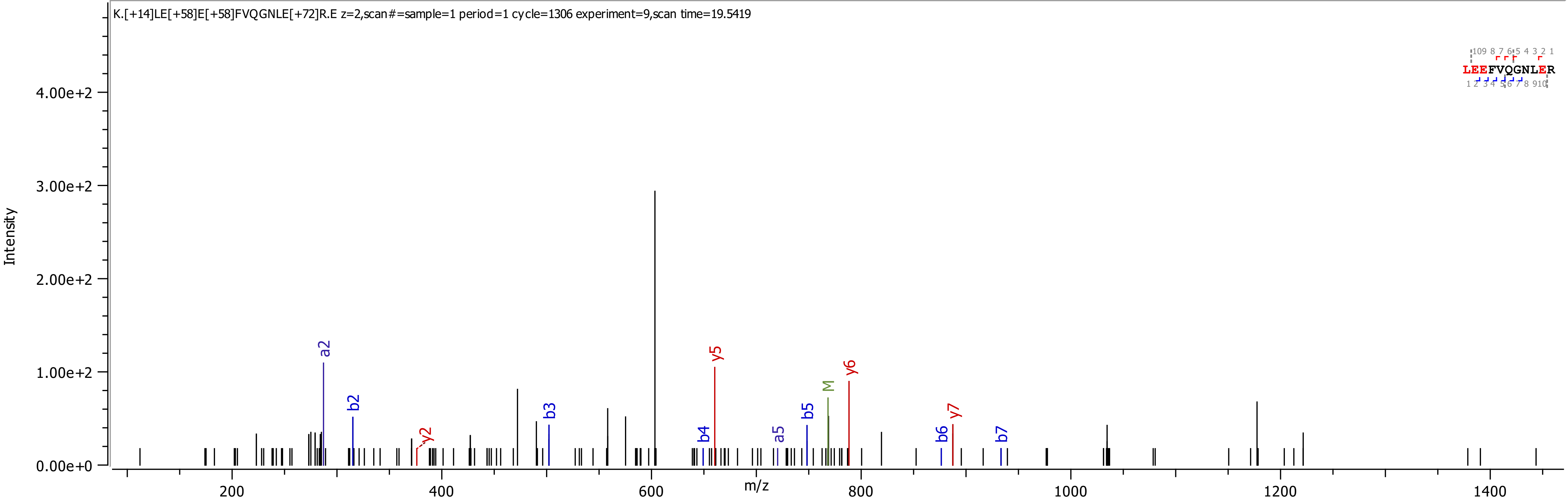

K.[+14]LE[+58]E[+58]FVQGNLE[+72]R.E z=2,scan#=sample=1 period=1 cycle=1306 experiment=9,scan time=19.5419
LEEFVQGNLER
Intensity
m/z
4.00e+2
3.00e+2
2.00e+2
1.00e+2
0.00e+0
200
400
600
800
1000
1200
1400
a2
b2
y2
b3
b4
y5
a5
b5
M
y6
b6
y7
b7

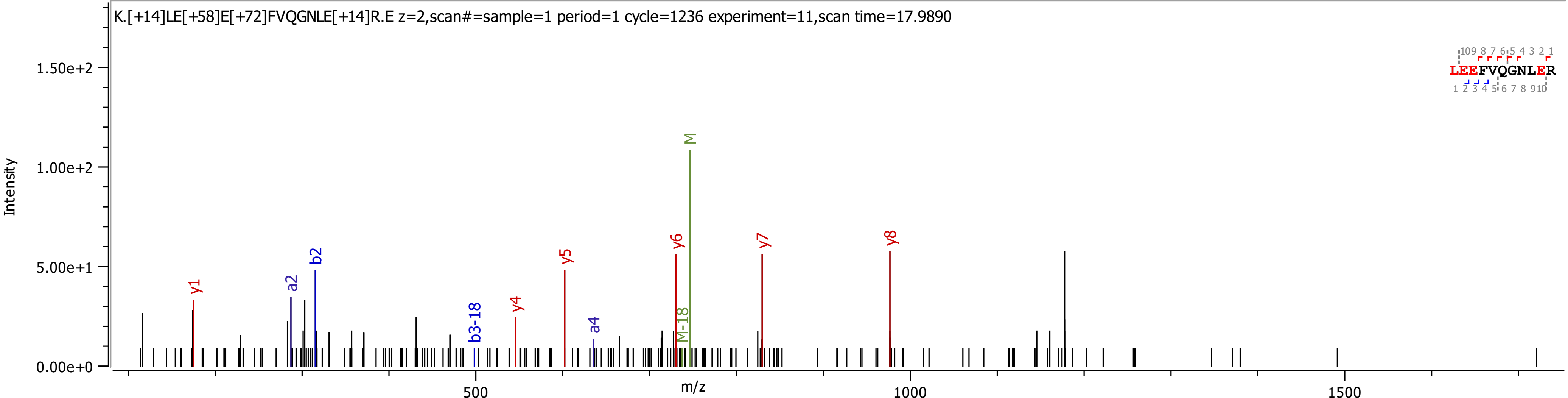
K.[+14]LE[+58]E[+72]FVQGNLE[+14]R.E z=2,scan#=sample=1 period=1 cycle=1236 experiment=11,scan time=17.9890
Intensity
m/z
0.00e+0
5.00e+1
1.00e+2
1.50e+2
500
1000
1500
y1
a2
b2
b3-18
y4
y5
a4
y6
M-18
M
y7
y8
LEEFVQGNLER

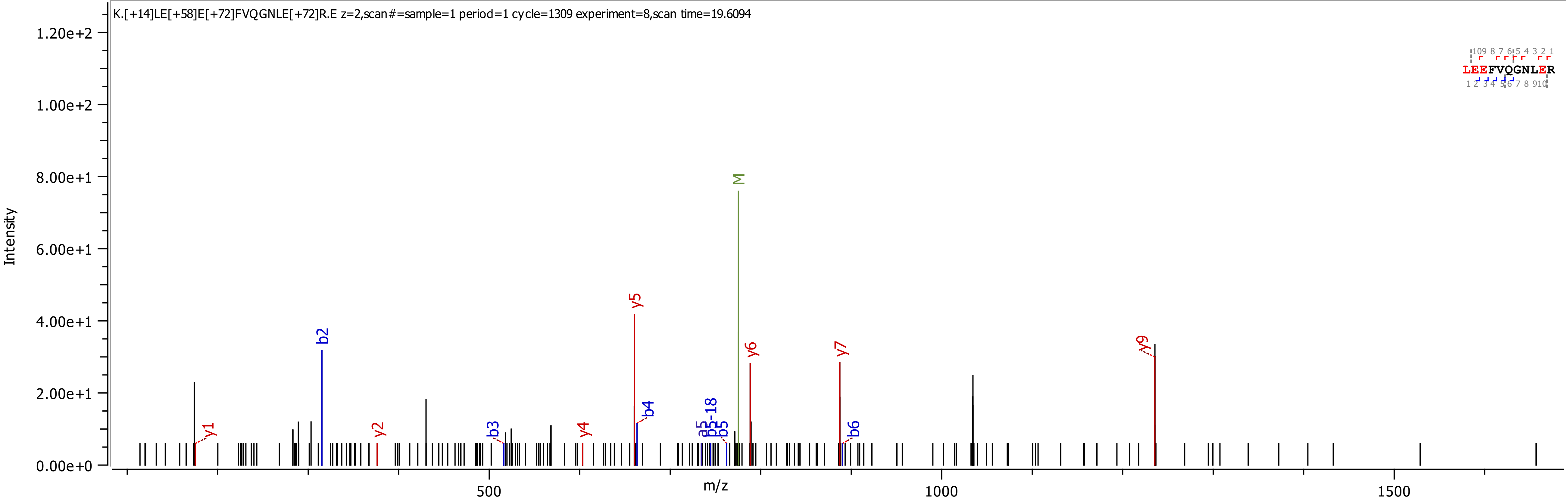
K.[+14]LE[+58]E[+72]FVQGNLE[+72]R.E z=2,scan#=sample=1 period=1 cycle=1309 experiment=8,scan time=19.6094
Intensity
m/z
LEEFVQGNLER
y1
b2
y2
b3
y4
y5
b4
a5
b5-18
b5
M
y6
y7
b6
y9
0.00e+0
2.00e+1
4.00e+1
6.00e+1
8.00e+1
1.00e+2
1.20e+2
500
1000
1500

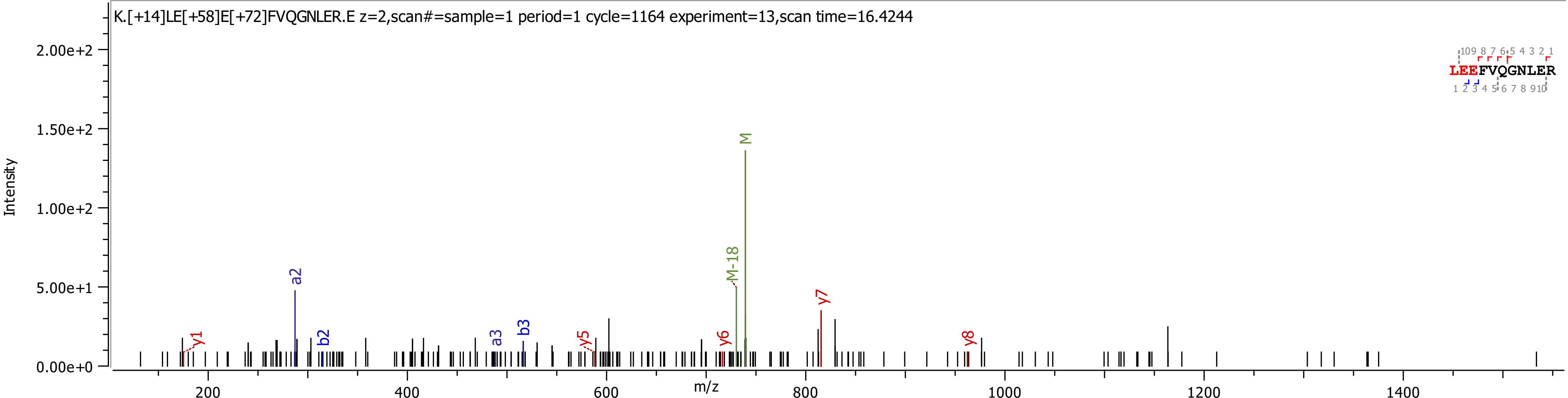

K.[+14]LE[+58]E[+72]FVQGNLER.E z=2,scan#=sample=1 period=1 cycle=1164 experiment=13,scan time=16.4244
Intensity
m/z
2.00e+2
1.50e+2
1.00e+2
5.00e+1
0.00e+0
200
400
600
800
1000
1200
1400
y1
a2
b2
a3
b3
y5
y6
M-18
M
y7
y8
LEEFVQGNLER

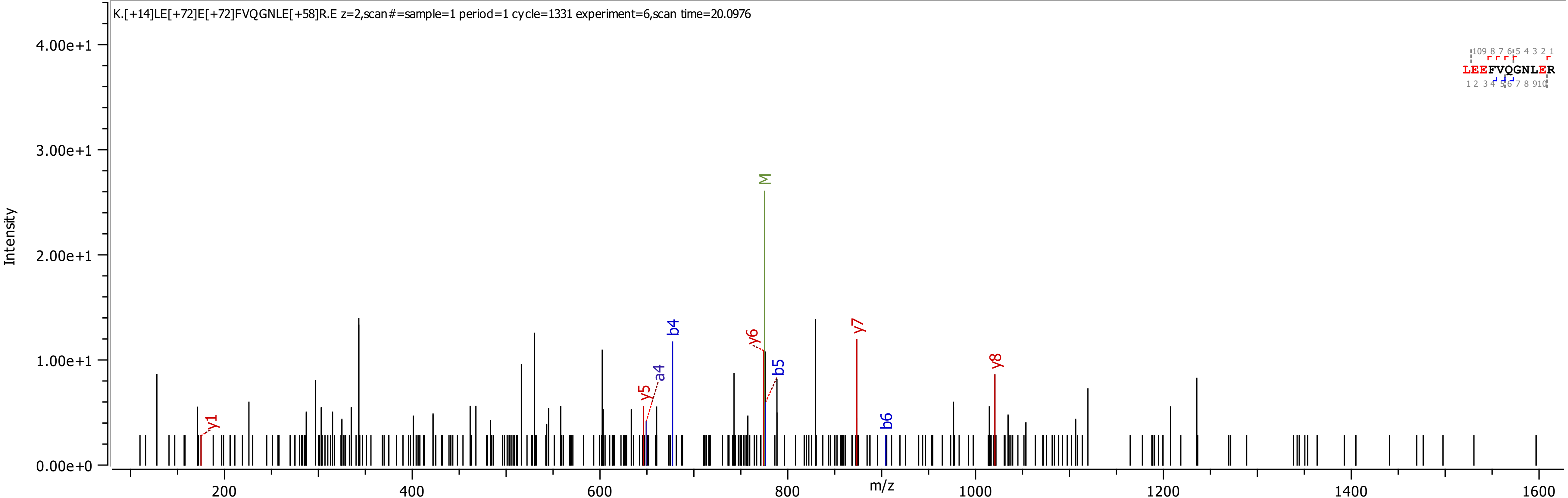

K.[+14]LE[+72]E[+72]FVQGNLE[+58]R.E z=2,scan#=sample=1 period=1 cycle=1331 experiment=6,scan time=20.0976
Intensity
4.00e+1
3.00e+1
2.00e+1
1.00e+1
0.00e+0
m/z
200
400
600
800
1000
1200
1400
1600
M
y1
y5
a4
b4
y6
b5
y7
b6
y8
LEEFVQGNLER

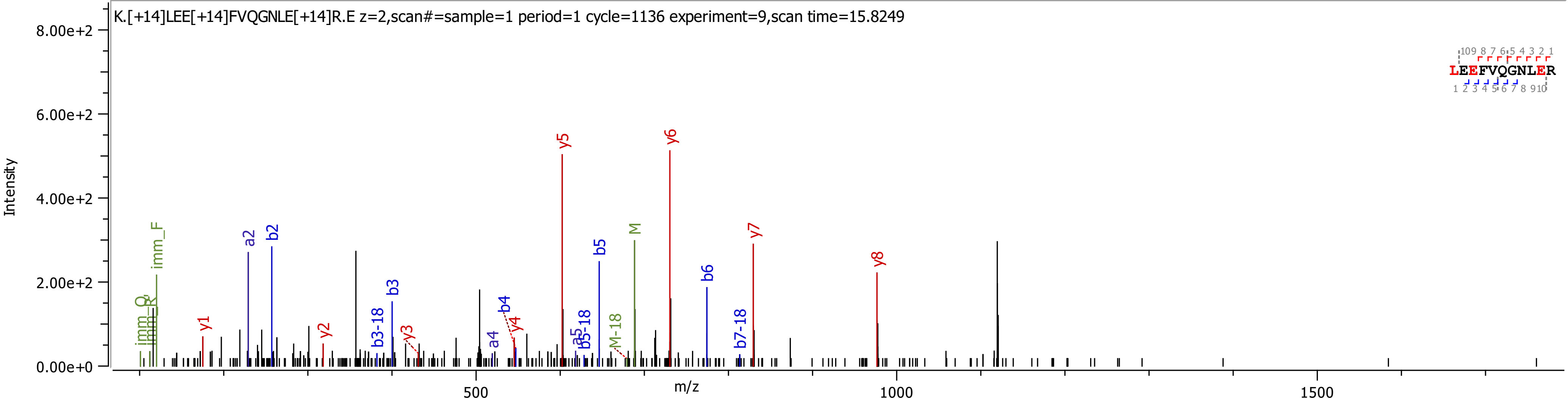
K.[+14]LEE[+14]FVQGNLE[+14]R.E z=2,scan#=sample=1 period=1 cycle=1136 experiment=9,scan time=15.8249
Intensity
8.00e+2
6.00e+2
4.00e+2
2.00e+2
0.00e+0
m/z
500
1000
1500
y1 y2 y3 y4 y5 y6 y7 y8
a2 b2 b3-18 b3 a4 b4 a5 b5-18 b5 b6 b7-18
imm_Q imm_R imm_F M-18 M
LEEFVQGNLER

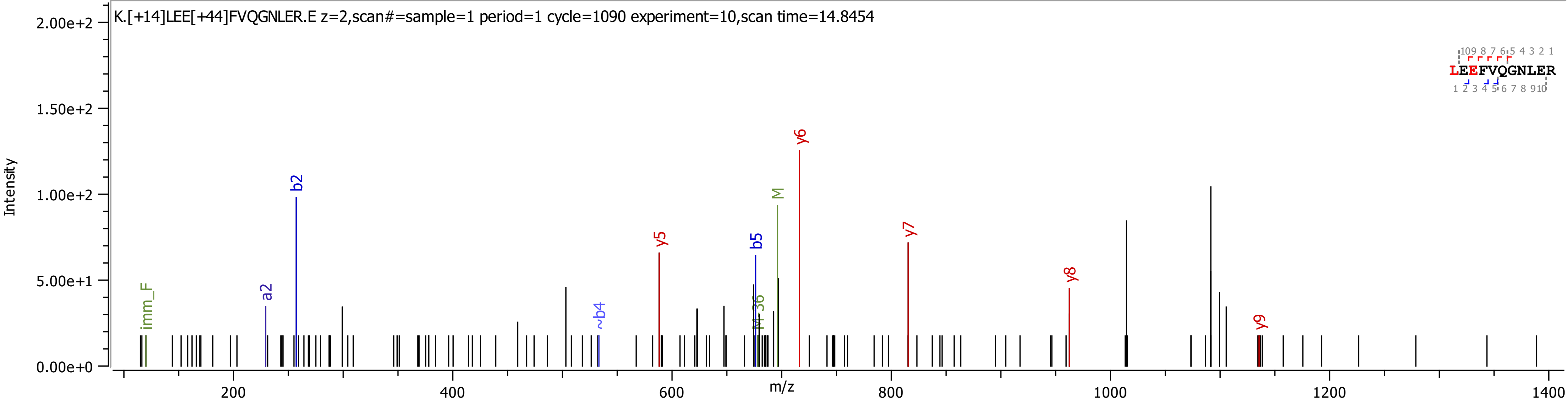
K.[+14]LEE[+44]FVQGNLER.E z=2,scan#=sample=1 period=1 cycle=1090 experiment=10,scan time=14.8454
Intensity
m/z
imm_F
a2
b2
~b4
y5
M-36
b5
M
y6
y7
y8
y9
LEEFVQGNLER
0.00e+0
5.00e+1
1.00e+2
1.50e+2
2.00e+2
200
400
600
800
1000
1200
1400

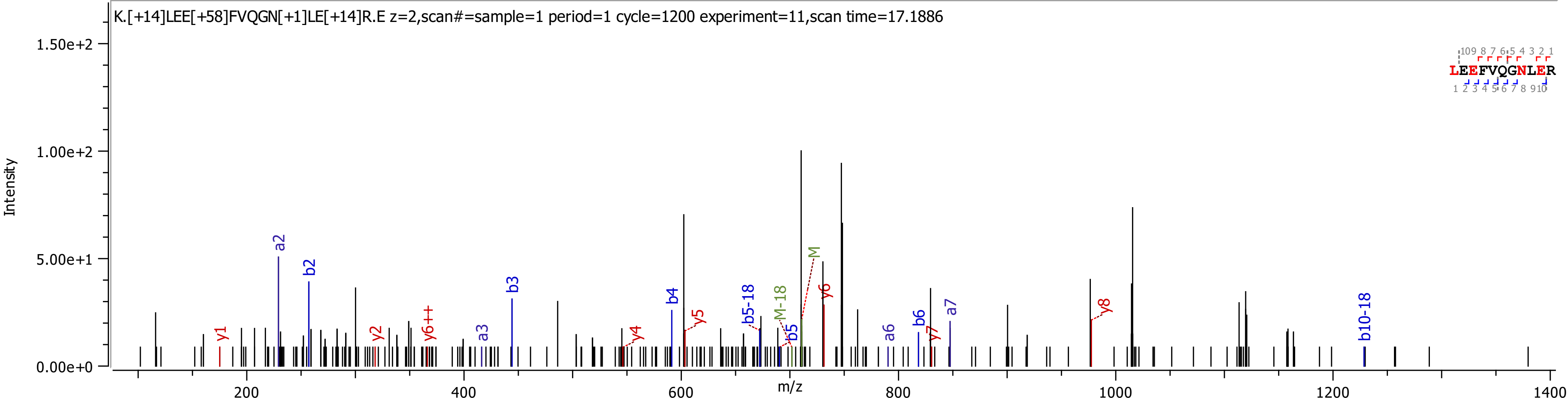
K.[+14]LEE[+58]FVQGN[+1]LE[+14]R.E z=2,scan#=sample=1 period=1 cycle=1200 experiment=11,scan time=17.1886
Intensity
1.50e+2
1.00e+2
5.00e+1
0.00e+0
m/z
200
400
600
800
1000
1200
1400
y1
a2
b2
y2
y6++
a3
b3
y4
b4
y5
b5-18
M-18
b5
M
y6
a6
b6
y7
a7
y8
b10-18
LEEFVQGNLER

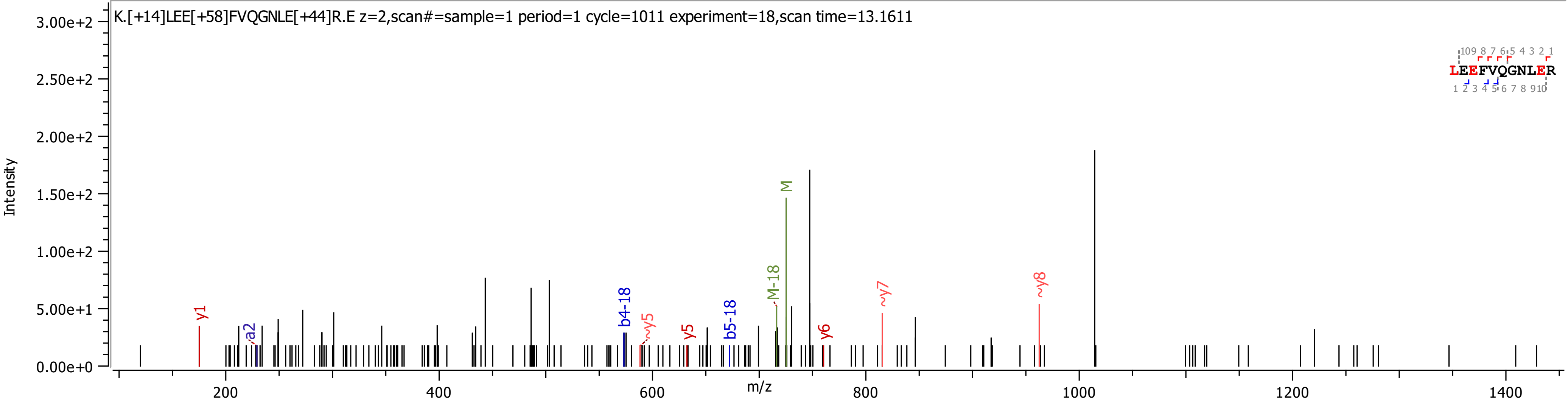
K.[+14]LEE[+58]FVQGNLE[+44]R.E z=2,scan#=sample=1 period=1 cycle=1011 experiment=18,scan time=13.1611
Intensity
3.00e+2
2.50e+2
2.00e+2
1.50e+2
1.00e+2
5.00e+1
0.00e+0
m/z
200
400
600
800
1000
1200
1400
y1
a2
b4-18
~y5
y5
b5-18
M-18
M
y6
~y7
~y8
LEEFVQGNLER

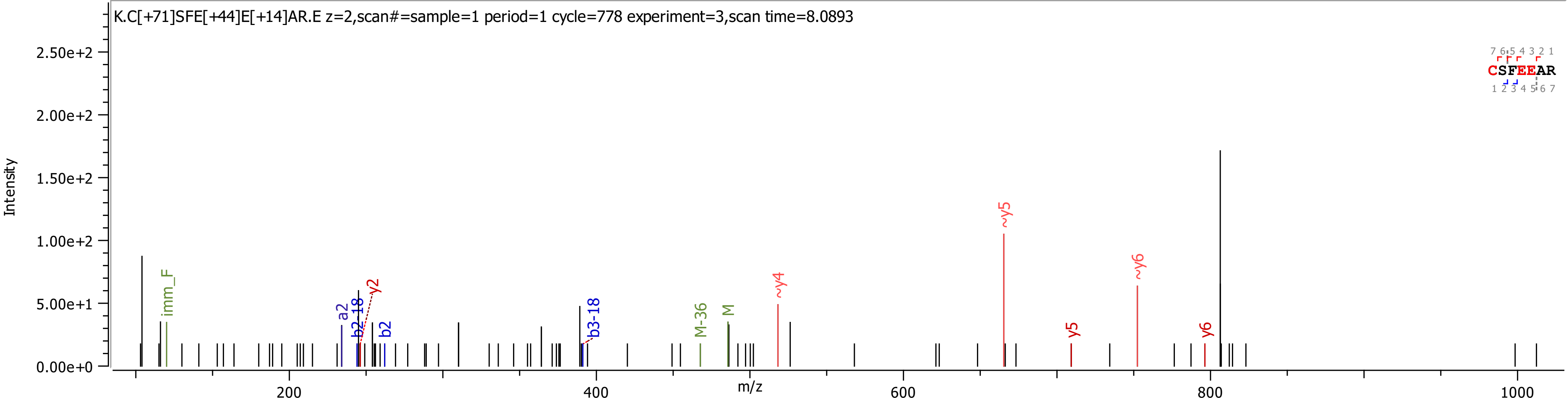
K.C[+71]SFE[+44]E[+14]AR.E z=2,scan#=sample=1 period=1 cycle=778 experiment=3,scan time=8.0893
Intensity
2.50e+2
2.00e+2
1.50e+2
1.00e+2
5.00e+1
0.00e+0
m/z
200
400
600
800
1000
imm_F
a2
b2-18
y2
b2
b3-18
M-36
M
~y4
~y5
y5
~y6
y6
7 6 5 4 3 2 1
CSFEEAR
1 2 3 4 5 6 7

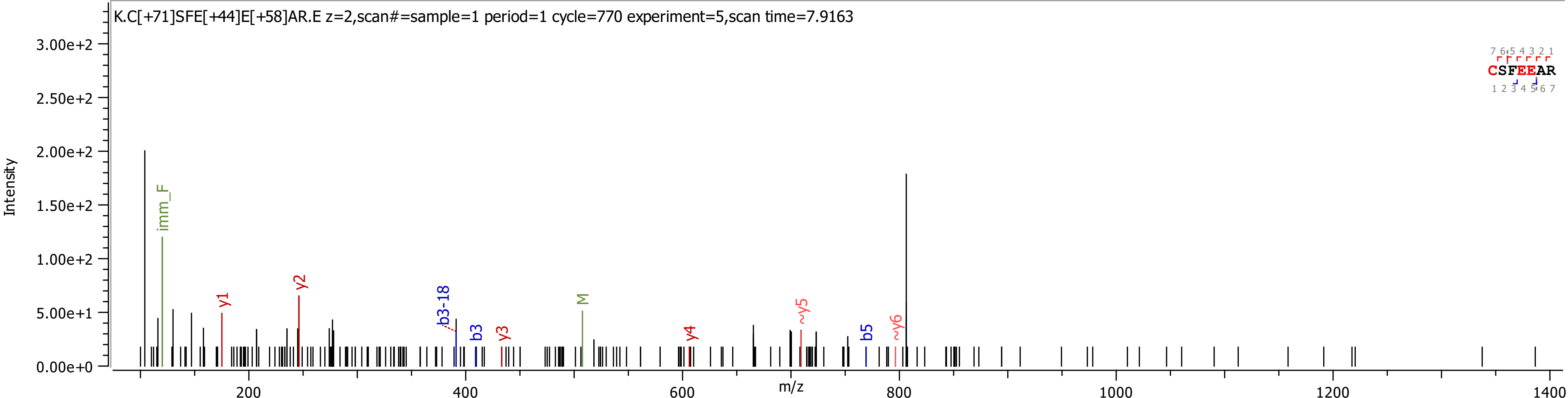
K.C[+71]SFE[+44]E[+58]AR.E z=2,scan#=sample=1 period=1 cycle=770 experiment=5,scan time=7.9163
Intensity
m/z
3.00e+2
2.50e+2
2.00e+2
1.50e+2
1.00e+2
5.00e+1
0.00e+0
200
400
600
800
1000
1200
1400
imm_F
y1
y2
b3-18
b3
y3
M
y4
~y5
b5
~y6
CSFEEAR
7 6 5 4 3 2 1
1 2 3 4 5 6 7

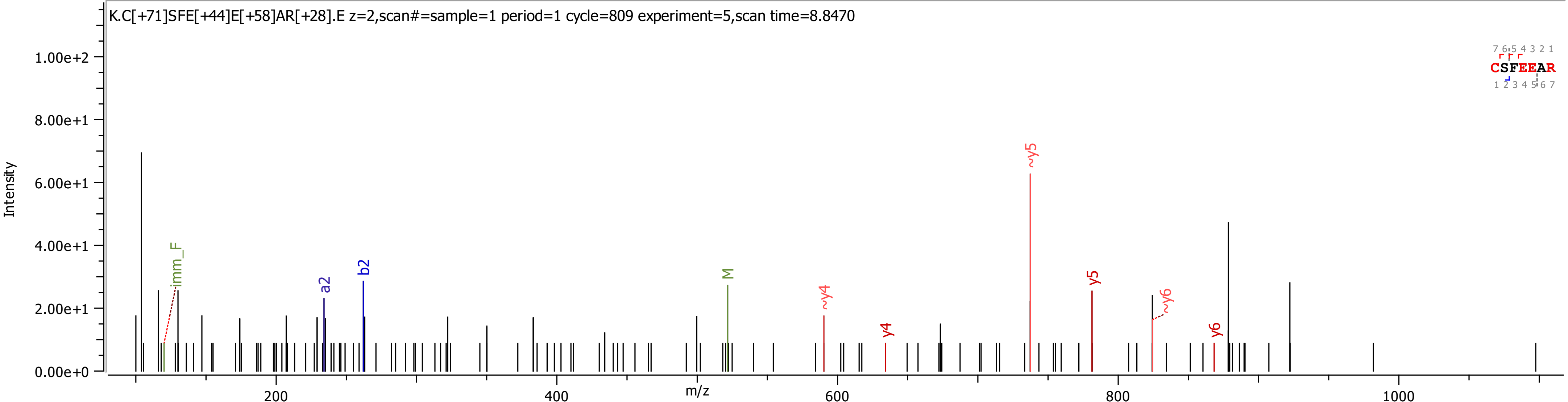

K.C[+71]SFE[+44]E[+58]AR[+28].E z=2,scan#=sample=1 period=1 cycle=809 experiment=5,scan time=8.8470
CSFEEAR
Intensity
m/z
imm_F
a2
b2
M
~y4
y4
~y5
y5
~y6
y6

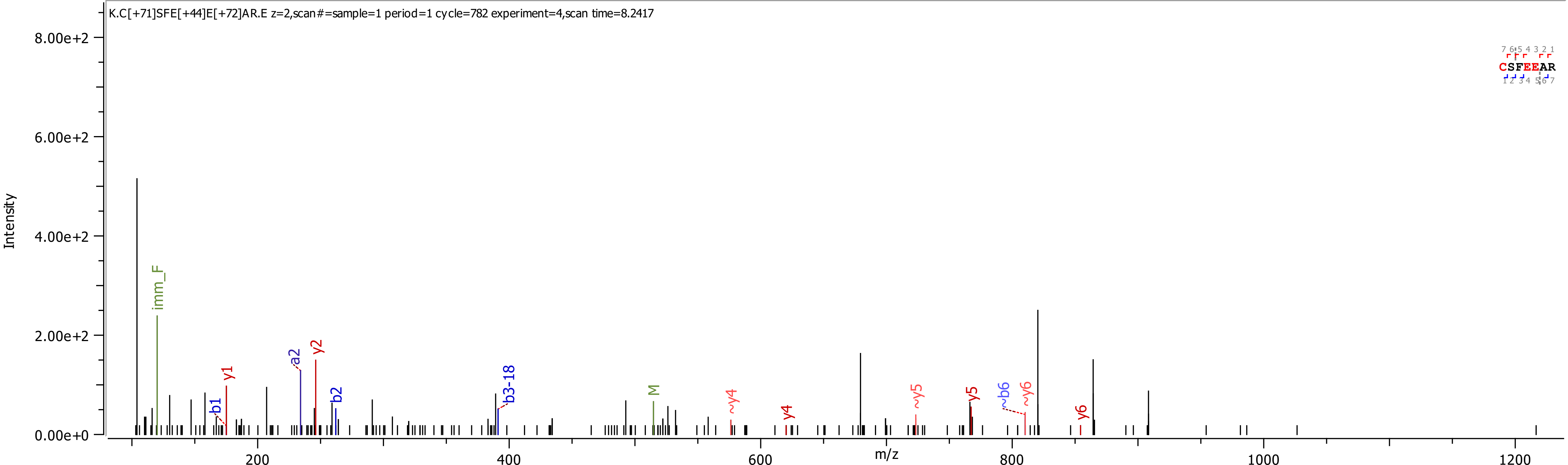
K.C[+71]SFE[+44]E[+72]AR.E z=2,scan#=sample=1 period=1 cycle=782 experiment=4,scan time=8.2417
Intensity
8.00e+2
6.00e+2
4.00e+2
2.00e+2
0.00e+0
m/z
200
400
600
800
1000
1200
CSFEEAR
imm_F
b1
y1
a2
y2
b2
b3-18
M
~y4
y4
~y5
y5
~b6
~y6
y6

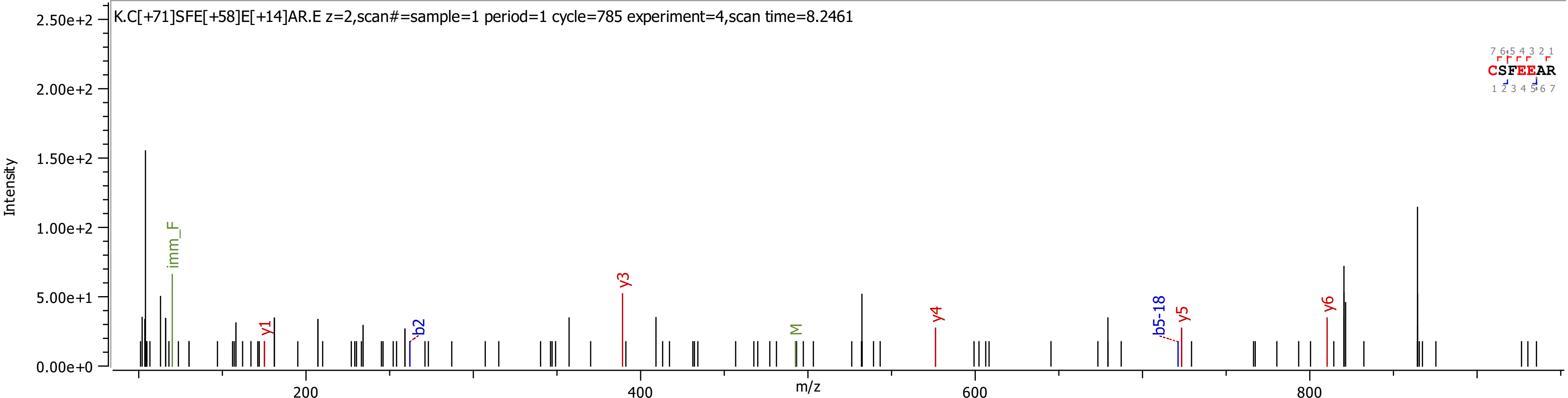

K.C[+71]SFE[+58]E[+14]AR.E z=2,scan#=sample=1 period=1 cycle=785 experiment=4,scan time=8.2461
Intensity
2.50e+2
2.00e+2
1.50e+2
1.00e+2
5.00e+1
0.00e+0
imm_F
y1
b2
y3
M
y4
b5-18
y5
y6
CSFEEAR
7 6 5 4 3 2 1
1 2 3 4 5 6 7
200
400
600
800
m/z

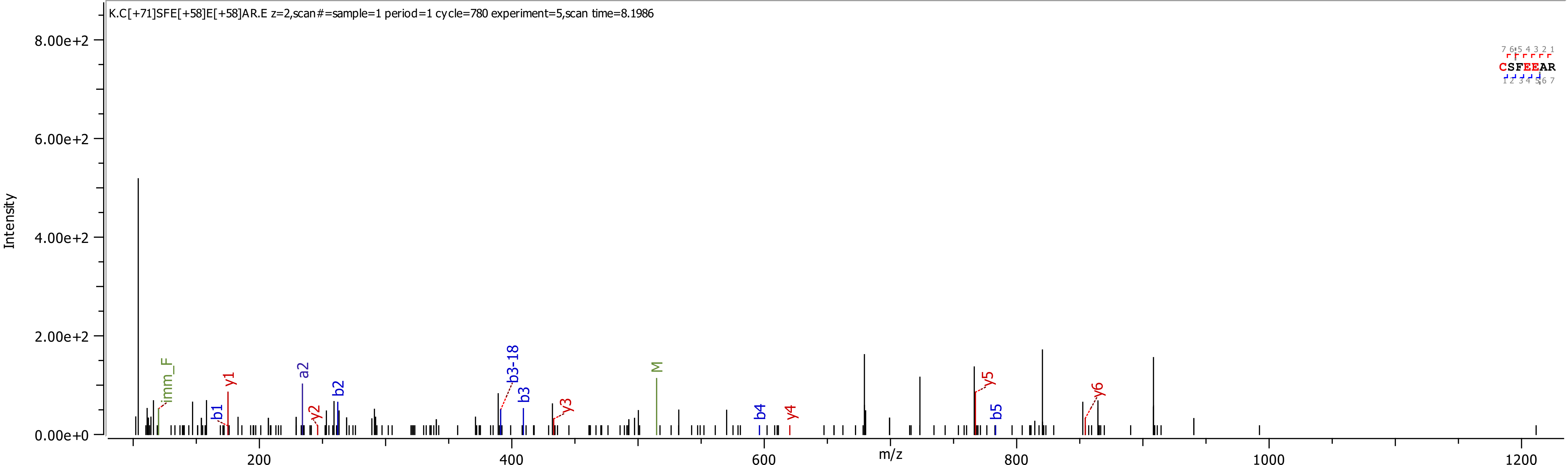
K.C[+71]SFE[+58]E[+58]AR.E z=2,scan#=sample=1 period=1 cycle=780 experiment=5,scan time=8.1986
Intensity
8.00e+2
6.00e+2
4.00e+2
2.00e+2
0.00e+0
m/z
200
400
600
800
1000
1200
imm_F
b1
y1
a2
y2
b2
b3-18
b3
y3
M
b4
y4
y5
b5
y6
CSFEEAR

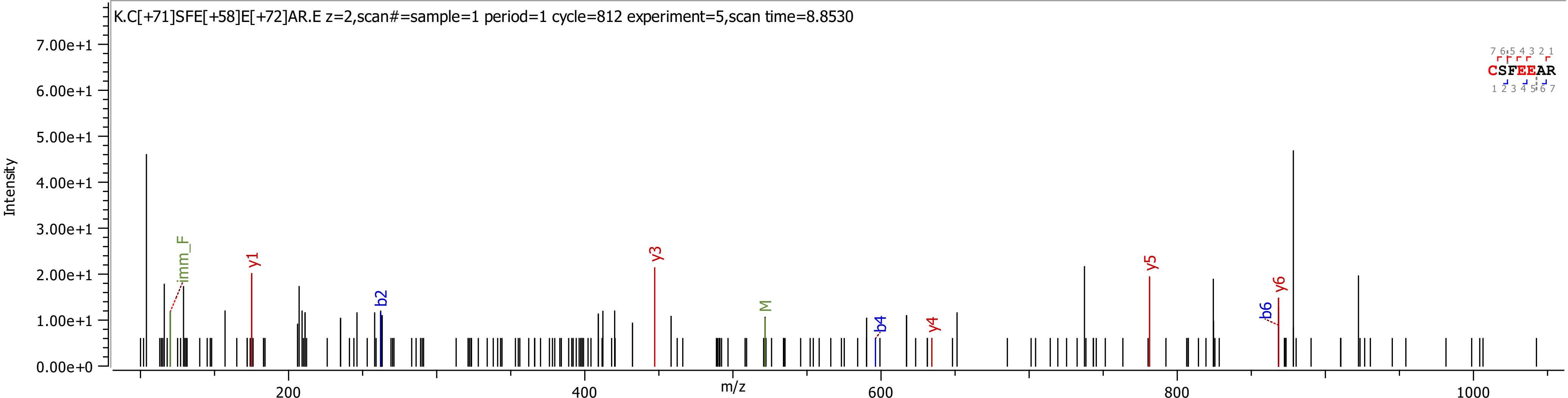
K.C[+71]SFE[+58]E[+72]AR.E z=2,scan#=sample=1 period=1 cycle=812 experiment=5,scan time=8.8530
Intensity
7.00e+1
6.00e+1
5.00e+1
4.00e+1
3.00e+1
2.00e+1
1.00e+1
0.00e+0
m/z
200
400
600
800
1000
imm_F
y1
b2
y3
M
b4
y4
y5
b6
y6
CSFEEAR

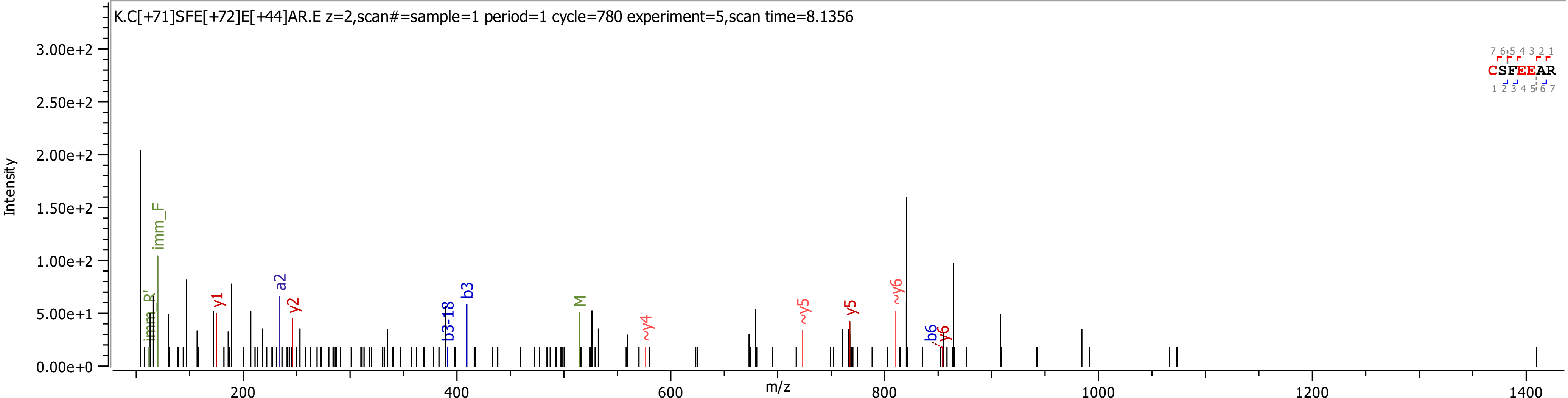

K.C[+71]SFE[+72]E[+44]AR.E z=2,scan#=sample=1 period=1 cycle=780 experiment=5,scan time=8.1356
Intensity
m/z
CSFEEAR
imm_R'
imm_F
y1
a2
y2
b3-18
b3
M
~y4
~y5
y5
~y6
b6
y6

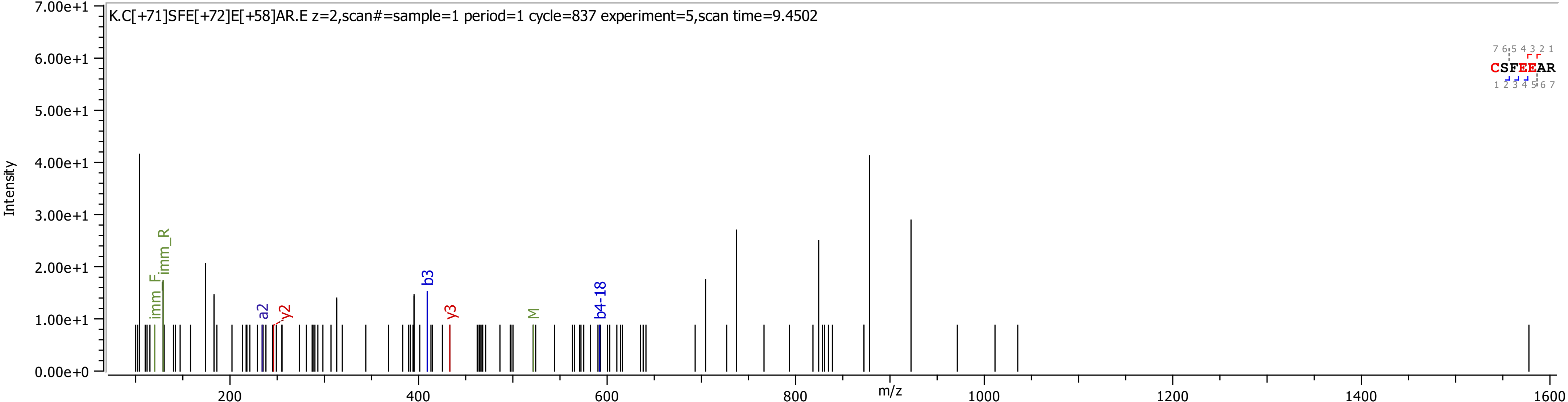
K.C[+71]SFE[+72]E[+58]AR.E z=2,scan#=sample=1 period=1 cycle=837 experiment=5,scan time=9.4502
Intensity
m/z
imm_F
imm_R
a2
y2
b3
y3
M
b4-18
CSFEEAR
7.00e+1
6.00e+1
5.00e+1
4.00e+1
3.00e+1
2.00e+1
1.00e+1
0.00e+0
200
400
600
800
1000
1200
1400
1600

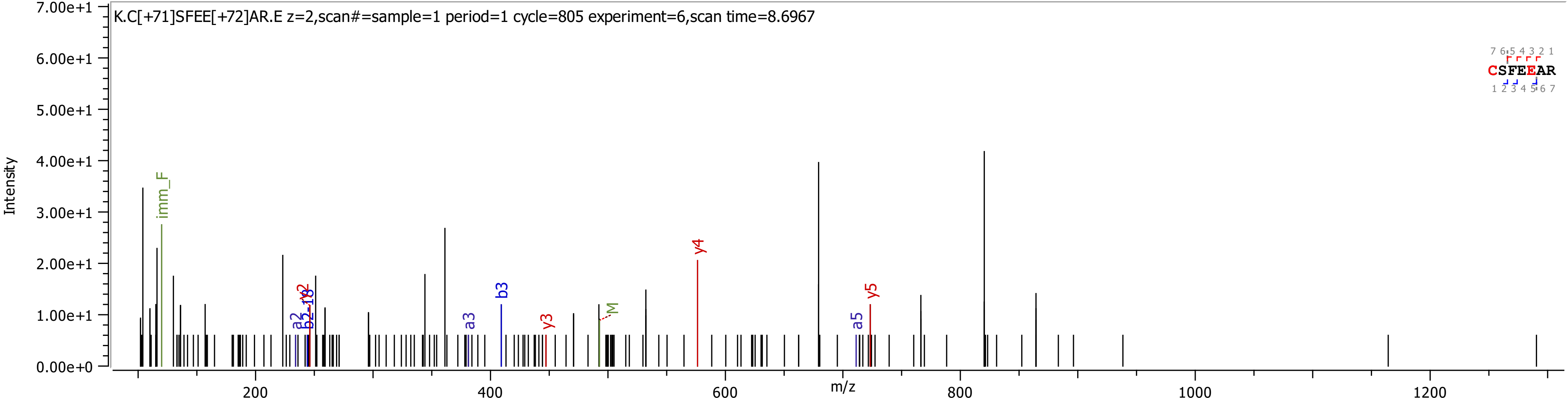

K.C[+71]SFEE[+72]AR.E z=2,scan#=sample=1 period=1 cycle=805 experiment=6,scan time=8.6967
Intensity
7.00e+1
6.00e+1
5.00e+1
4.00e+1
3.00e+1
2.00e+1
1.00e+1
0.00e+0
m/z
200
400
600
800
1000
1200
imm_F
a2
b2
y2
a3
b3
y3
M
y4
a5
y5
CSFEEAR

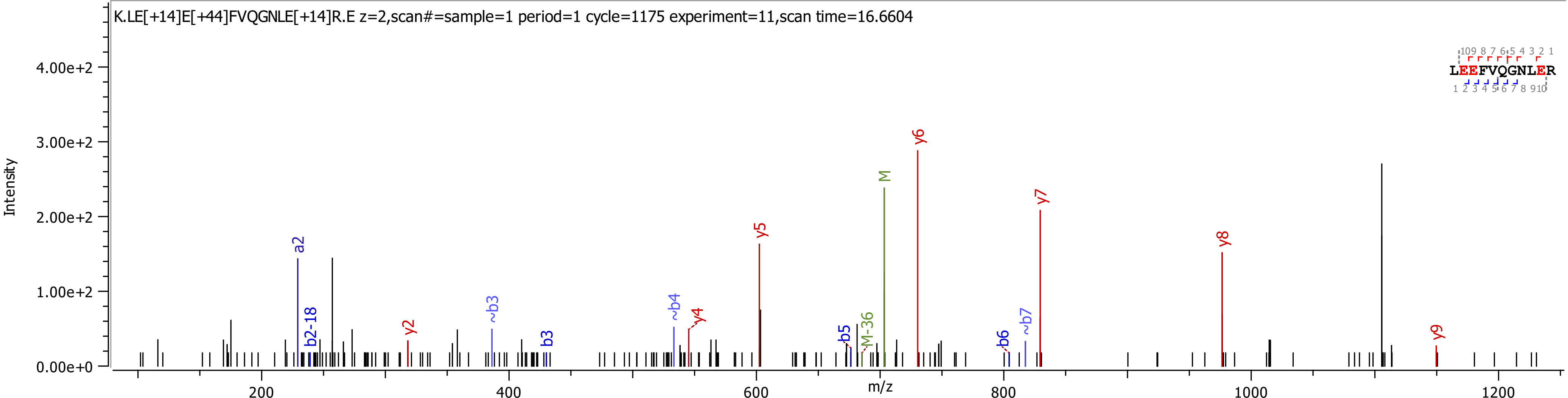

K.LE[+14]E[+44]FVQGNLE[+14]R.E z=2,scan#=sample=1 period=1 cycle=1175 experiment=11,scan time=16.6604
Intensity
4.00e+2
3.00e+2
2.00e+2
1.00e+2
0.00e+0
200
400
600
800
1000
1200
m/z
a2
b2-18
y2
~b3
b3
~b4
y4
y5
b5
M-36
M
y6
b6
~b7
y7
y8
y9
LEEFVQGNLER

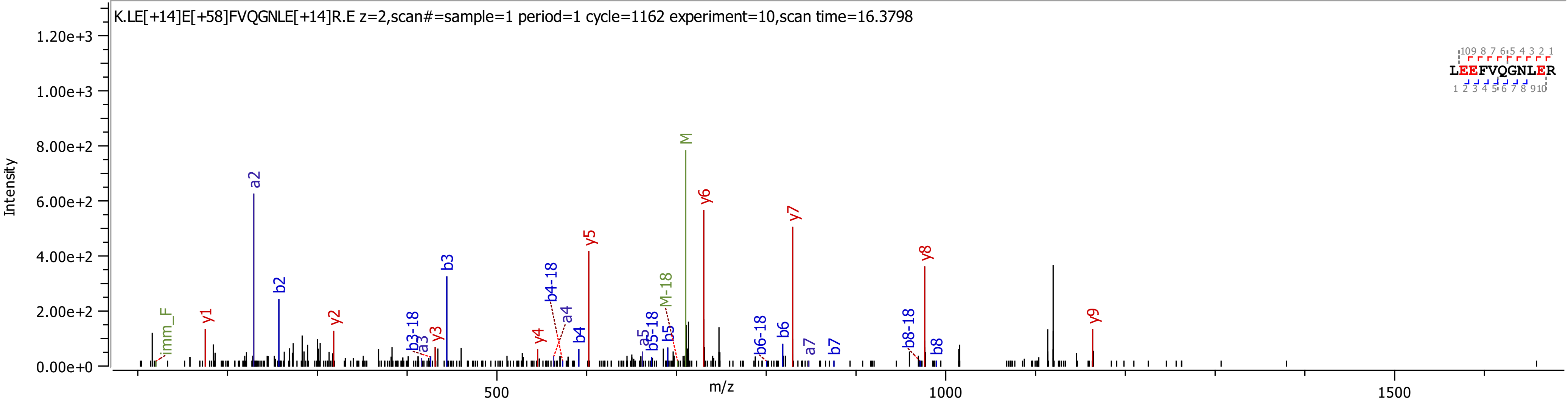

K.LE[+14]E[+58]FVQGNLE[+14]R.E z=2,scan#=sample=1 period=1 cycle=1162 experiment=10,scan time=16.3798
Intensity
m/z
0.00e+0
2.00e+2
4.00e+2
6.00e+2
8.00e+2
1.00e+3
1.20e+3
500
1000
1500
LEEFVQGNLER
imm_F
y1
a2
b2
y2
b3-18
a3
y3
b3
y4
b4-18
a4
b4
y5
a5
b5-18
b5
M-18
M
y6
b6-18
b6
y7
a7
b7
b8-18
y8
b8
y9

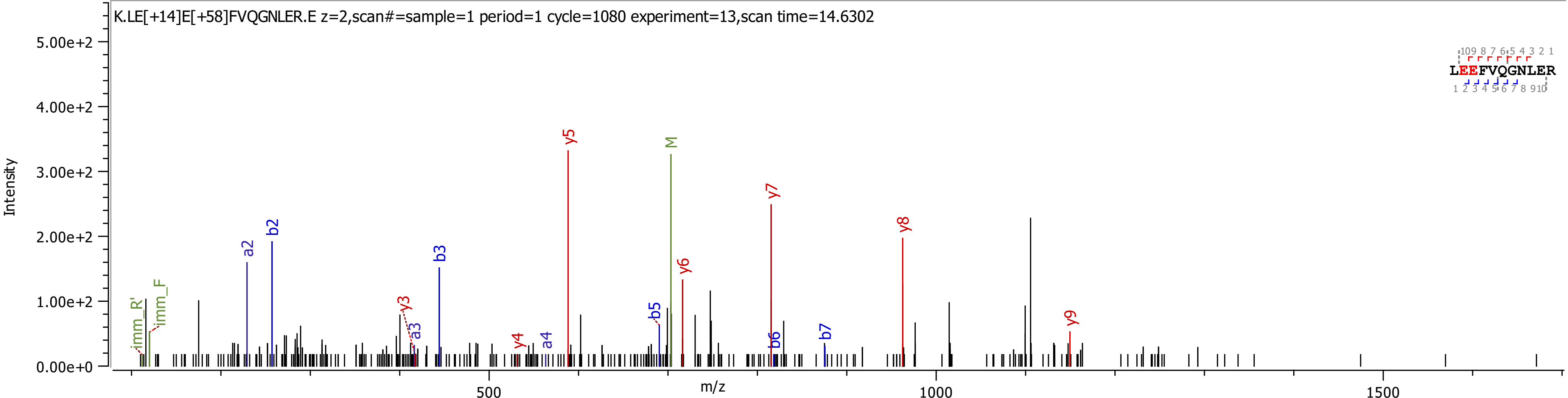

K.LE[+14]E[+58]FVQGNLER.E z=2,scan#=sample=1 period=1 cycle=1080 experiment=13,scan time=14.6302
Intensity
m/z
5.00e+2
4.00e+2
3.00e+2
2.00e+2
1.00e+2
0.00e+0
500
1000
1500
LEEFVQGNLER
imm_R'
imm_F
a2
b2
b3
y3
a3
y4
a4
y5
M
b5
y6
y7
b6
b7
y8
y9

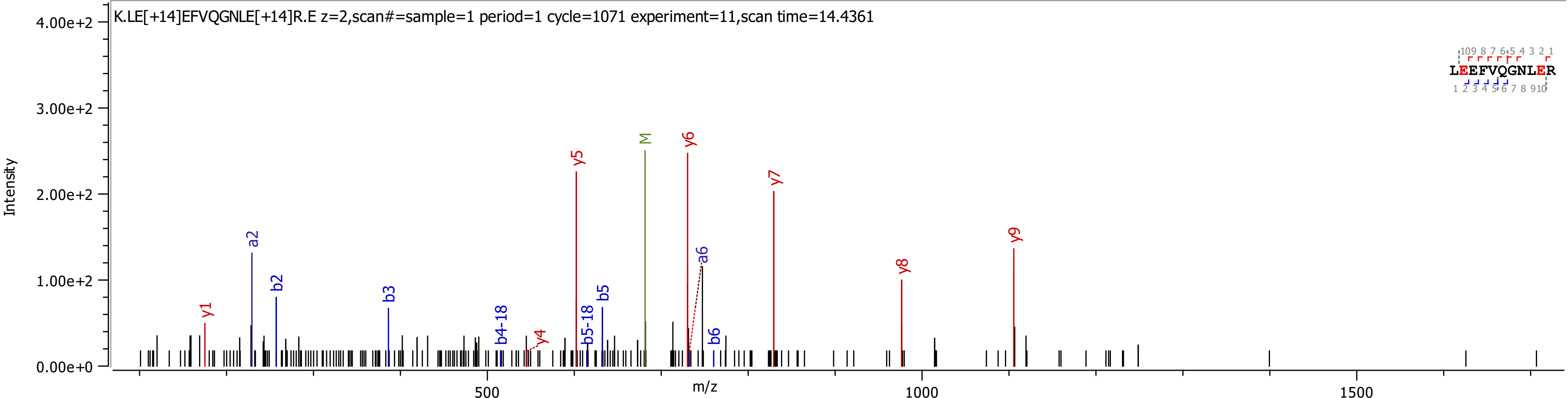
K.LE[+14]EFVQGNLE[+14]R.E z=2,scan#=sample=1 period=1 cycle=1071 experiment=11,scan time=14.4361
Intensity
4.00e+2
3.00e+2
2.00e+2
1.00e+2
0.00e+0
m/z
500
1000
1500
y1
a2
b2
b3
b4-18
y4
y5
b5-18
b5
M
y6
a6
b6
y7
y8
y9
LEEFVQGNLER

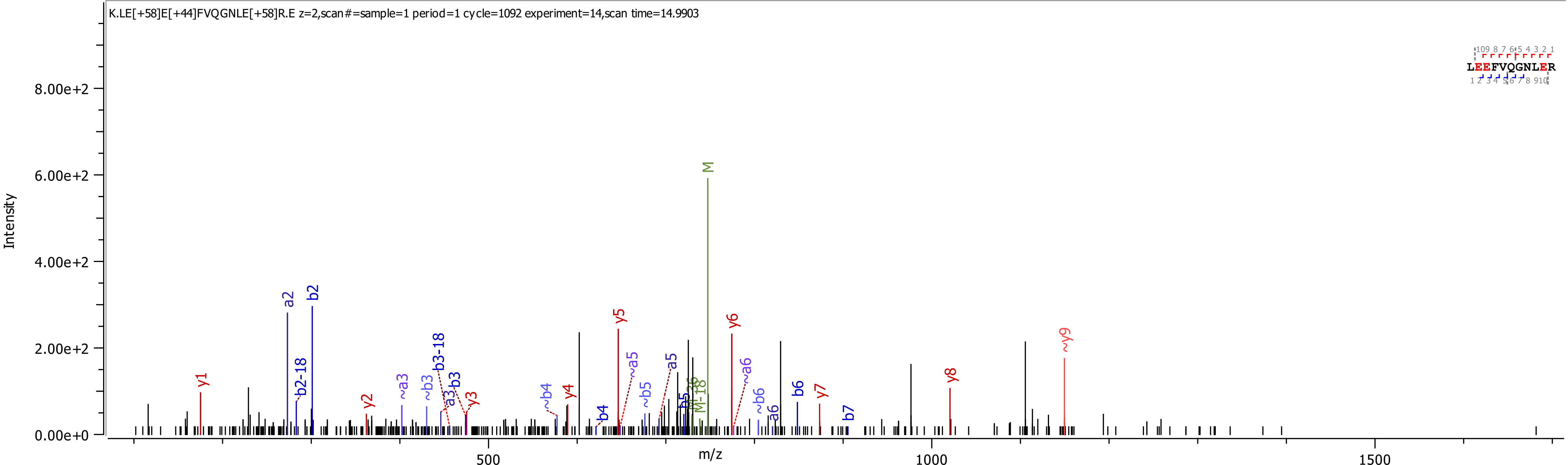

K.LE[+58]E[+44]FVQGNLE[+58]R.E z=2,scan#=sample=1 period=1 cycle=1092 experiment=14,scan time=14.9903
Intensity
8.00e+2
6.00e+2
4.00e+2
2.00e+2
0.00e+0
m/z
500
1000
1500
LEEFVQGNLER
M
a2
b2
b2-18
y1
y2
~a3
~b3
b3-18
a3
b3
y3
~b4
y4
b4
y5
~a5
~b5
a5
b5
M-36
M-18
y6
~a6
~b6
a6
b6
y7
b7
y8
~y9

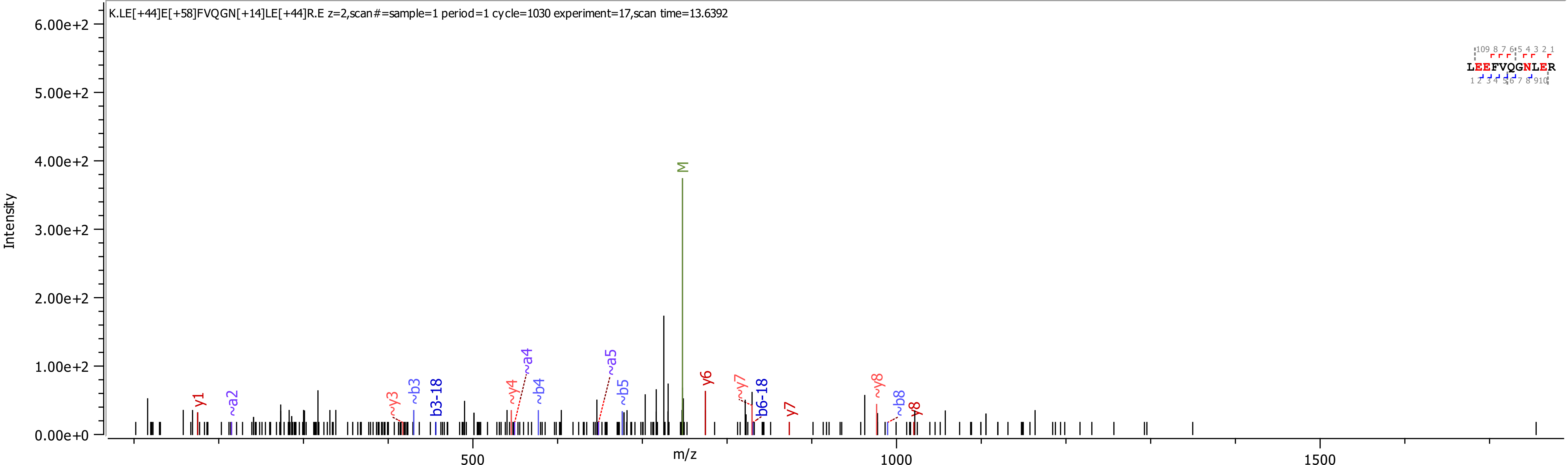
K.LE[+44]E[+58]FVQGN[+14]LE[+44]R.E z=2,scan#=sample=1 period=1 cycle=1030 experiment=17,scan time=13.6392
Intensity
m/z
M
y1
~a2
~y3
~b3
b3-18
~y4
~a4
~b4
~a5
~b5
y6
~y7
b6-18
y7
~y8
~b8
y8
LEEFVQGNLER

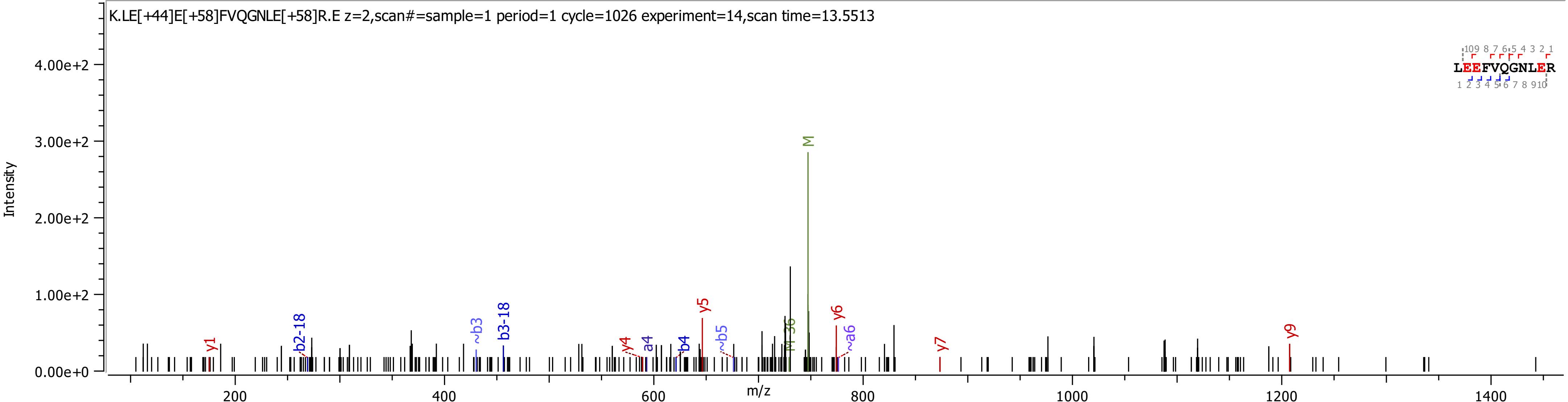
K.LE[+44]E[+58]FVQGNLE[+58]R.E z=2,scan#=sample=1 period=1 cycle=1026 experiment=14,scan time=13.5513
LEEFVQGNLER
Intensity
m/z
4.00e+2
3.00e+2
2.00e+2
1.00e+2
0.00e+0
200
400
600
800
1000
1200
1400
y1
b2-18
~b3
b3-18
y4
a4
b4
y5
~b5
M-36
M
y6
~a6
y7
y9

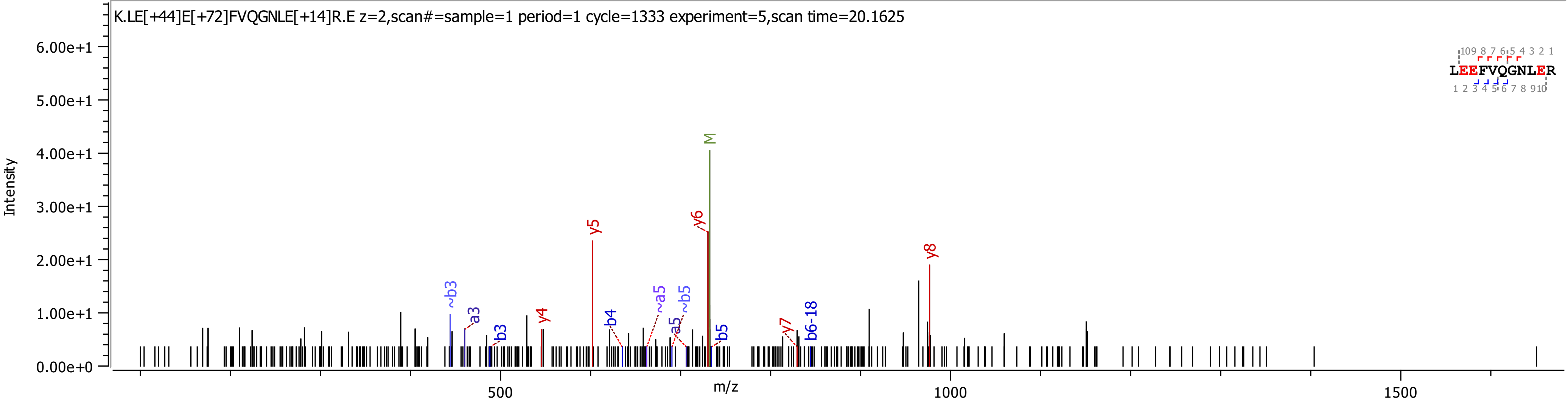

K.LE[+44]E[+72]FVQGNLE[+14]R.E z=2,scan#=sample=1 period=1 cycle=1333 experiment=5,scan time=20.1625
Intensity
6.00e+1
5.00e+1
4.00e+1
3.00e+1
2.00e+1
1.00e+1
0.00e+0
m/z
500
1000
1500
M
y5
y6
y8
y4
y7
~b3
a3
b3
b4
~a5
a5
~b5
b5
b6-18
LEEFVQGNLER

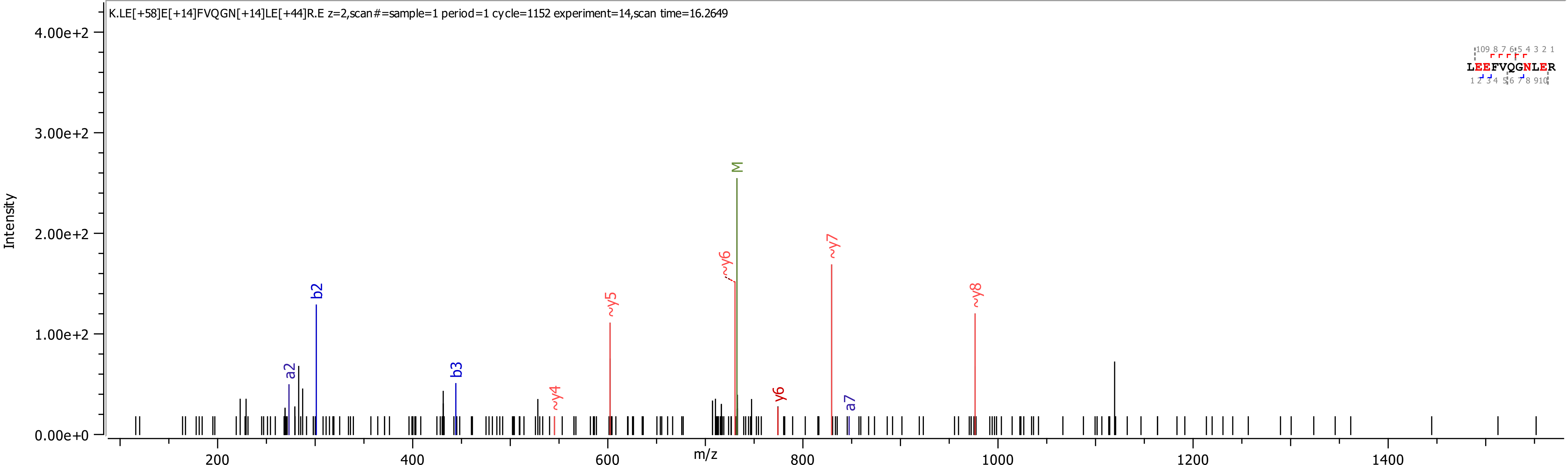

K.LE[+58]E[+14]FVQGN[+14]LE[+44]R.E z=2,scan#=sample=1 period=1 cycle=1152 experiment=14,scan time=16.2649
LEEFVQGNLER
Intensity
m/z
4.00e+2
3.00e+2
2.00e+2
1.00e+2
0.00e+0
200
400
600
800
1000
1200
1400
a2
b2
b3
~y4
~y5
~y6
M
y6
~y7
a7
~y8

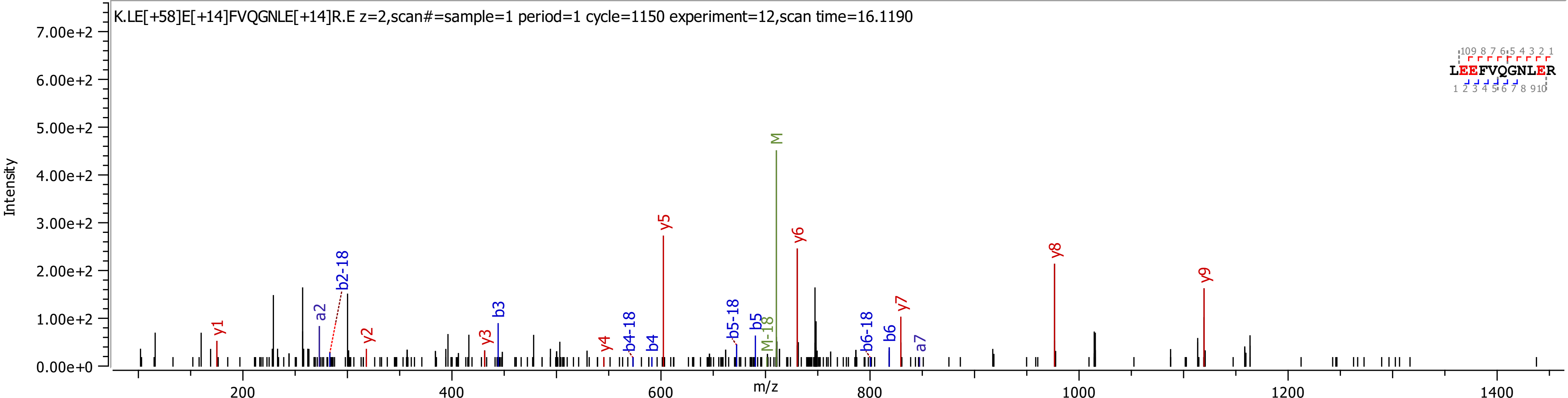

K.LE[+58]E[+14]FVQGNLE[+14]R.E z=2,scan#=sample=1 period=1 cycle=1150 experiment=12,scan time=16.1190
Intensity
7.00e+2
6.00e+2
5.00e+2
4.00e+2
3.00e+2
2.00e+2
1.00e+2
0.00e+0
m/z
200
400
600
800
1000
1200
1400
y1
a2
b2-18
y2
y3
b3
y4
b4-18
b4
y5
b5-18
b5
M-18
M
y6
b6-18
b6
y7
a7
y8
y9
LEEFVQGNLER

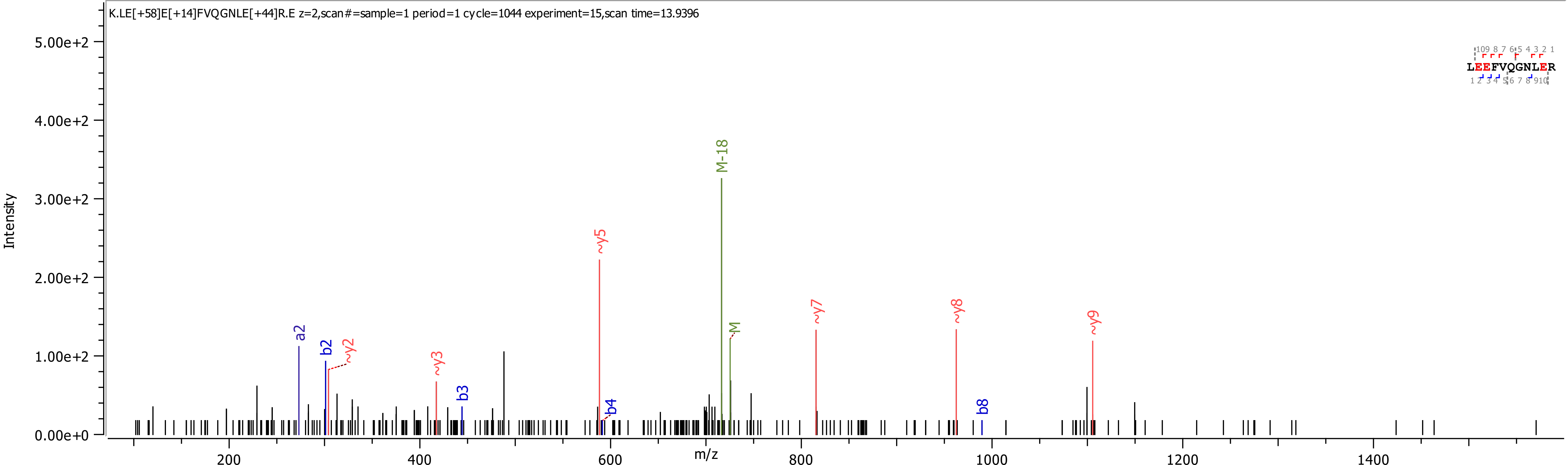

K.LE[+58]E[+14]FVQGNLE[+44]R.E z=2,scan#=sample=1 period=1 cycle=1044 experiment=15,scan time=13.9396
Intensity
m/z
LEEFVQGNLER
a2
b2
~y2
~y3
b3
~y5
b4
M-18
M
~y7
~y8
b8
~y9

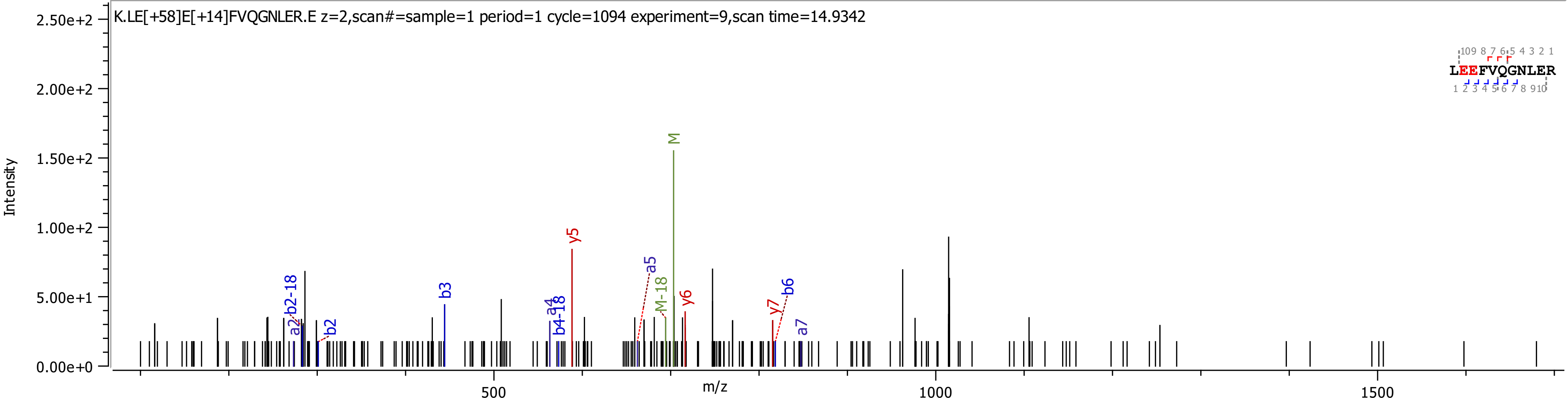
K.LE[+58]E[+14]FVQGNLER.E z=2,scan#=sample=1 period=1 cycle=1094 experiment=9,scan time=14.9342
Intensity
m/z
2.50e+2
2.00e+2
1.50e+2
1.00e+2
5.00e+1
0.00e+0
500
1000
1500
LEEFVQGNLER
M
y5
a5
M-18
y6
y7
b6
a7
b3
a4
b4-18
b2-18
a2
b2

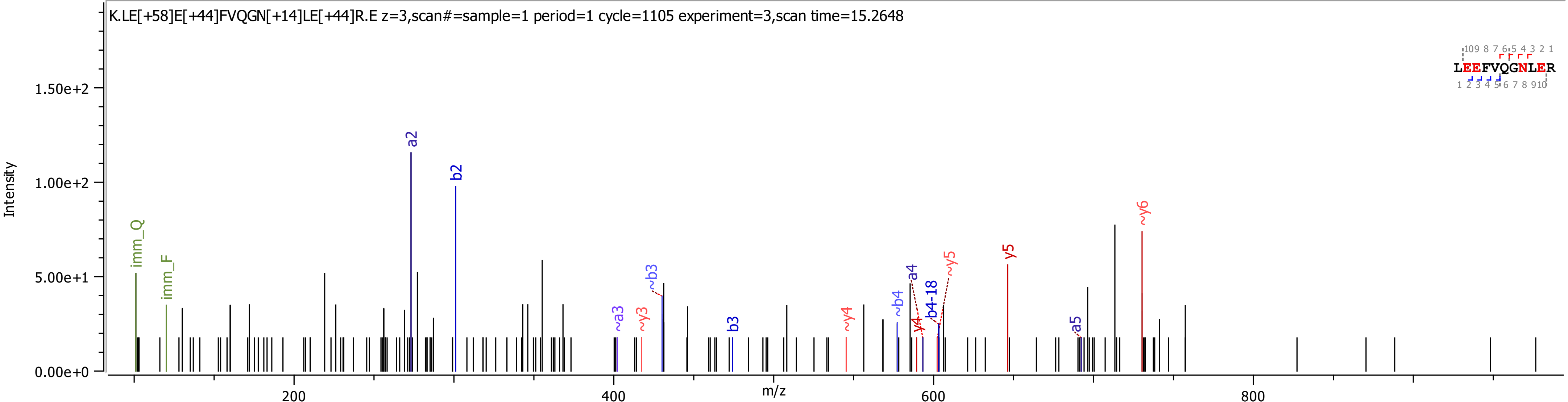

K.LE[+58]E[+44]FVQGN[+14]LE[+44]R.E z=3,scan#=sample=1 period=1 cycle=1105 experiment=3,scan time=15.2648
Intensity
m/z
1.50e+2
1.00e+2
5.00e+1
0.00e+0
200
400
600
800
imm_Q
imm_F
a2
b2
~a3
~y3
~b3
b3
~y4
~b4
a4
y4
b4-18
~y5
y5
a5
~y6
LEEFVQGNLER

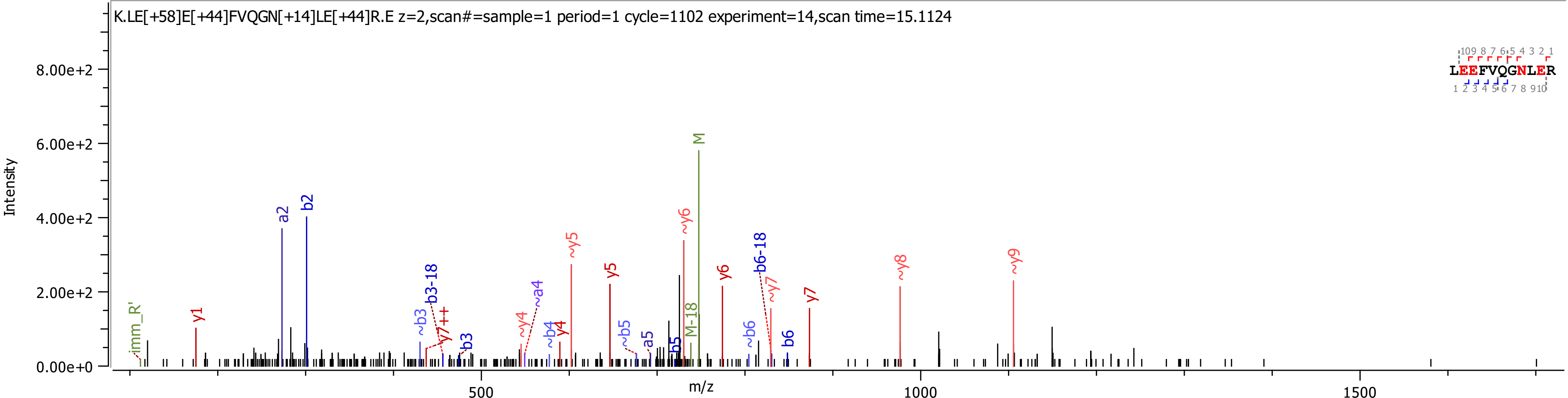

K.LE[+58]E[+44]FVQGN[+14]LE[+44]R.E z=2,scan#=sample=1 period=1 cycle=1102 experiment=14,scan time=15.1124
Intensity
8.00e+2
6.00e+2
4.00e+2
2.00e+2
0.00e+0
m/z
500
1000
1500
imm_R'
y1
a2
b2
~b3
b3-18
y7++
b3
~y4
~a4
~b4
y4
~y5
y5
~b5
a5
b5
M-18
~y6
M
y6
~b6
b6-18
~y7
b6
y7
~y8
~y9
LEEFVQGNLER

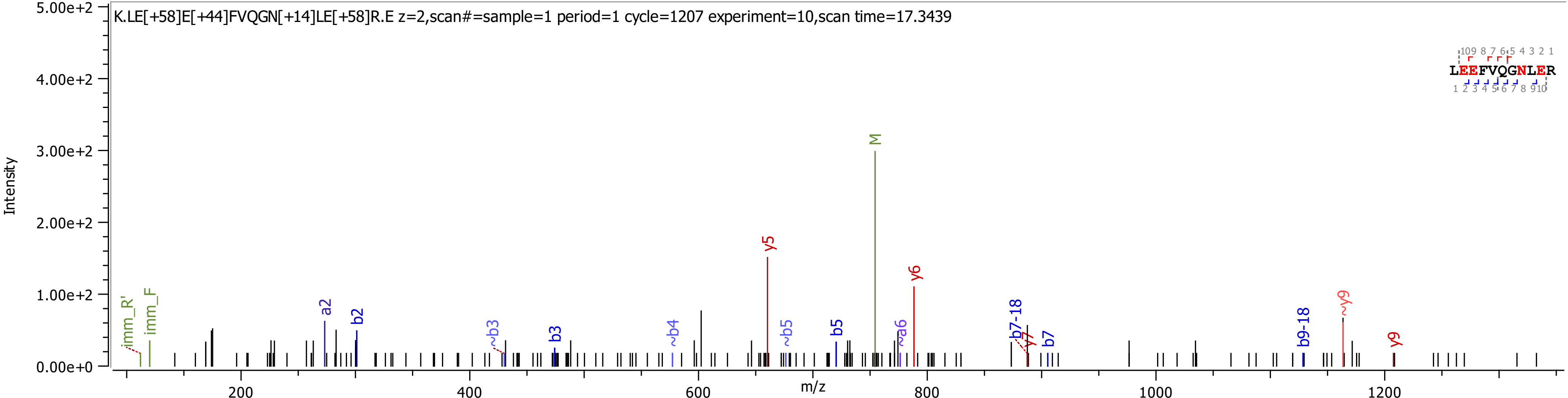

K.LE[+58]E[+44]FVQGN[+14]LE[+58]R.E z=2,scan#=sample=1 period=1 cycle=1207 experiment=10,scan time=17.3439
LEEFVQGNLER
Intensity
m/z
5.00e+2
4.00e+2
3.00e+2
2.00e+2
1.00e+2
0.00e+0
200
400
600
800
1000
1200
imm_R'
imm_F
a2
b2
~b3
b3
~b4
y5
~b5
b5
M
~a6
y6
b7-18
y7
b7
b9-18
~y9
y9

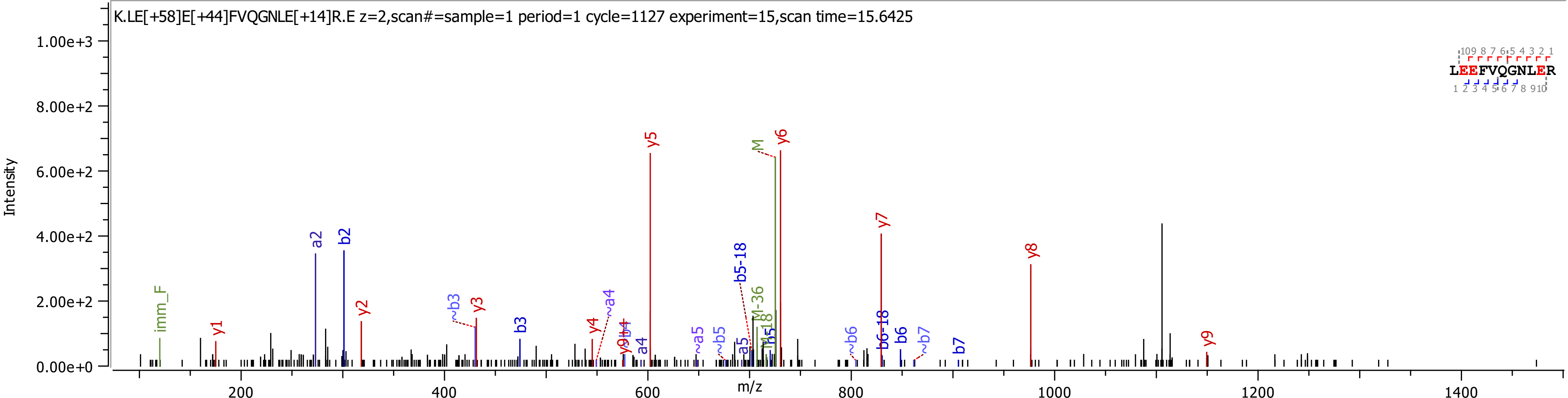

K.LE[+58]E[+44]FVQGNLE[+14]R.E z=2,scan#=sample=1 period=1 cycle=1127 experiment=15,scan time=15.6425
Intensity
m/z
1.00e+3
8.00e+2
6.00e+2
4.00e+2
2.00e+2
0.00e+0
200
400
600
800
1000
1200
1400
LEEFVQGNLER
imm_F
y1
a2
b2
y2
~b3
y3
b3
y4
~a4
y9
b4
a4
y5
~a5
~b5
a5
b5-18
M-36
M-18
b5
M
y6
~b6
b6-18
b6
~b7
b7
y7
y8

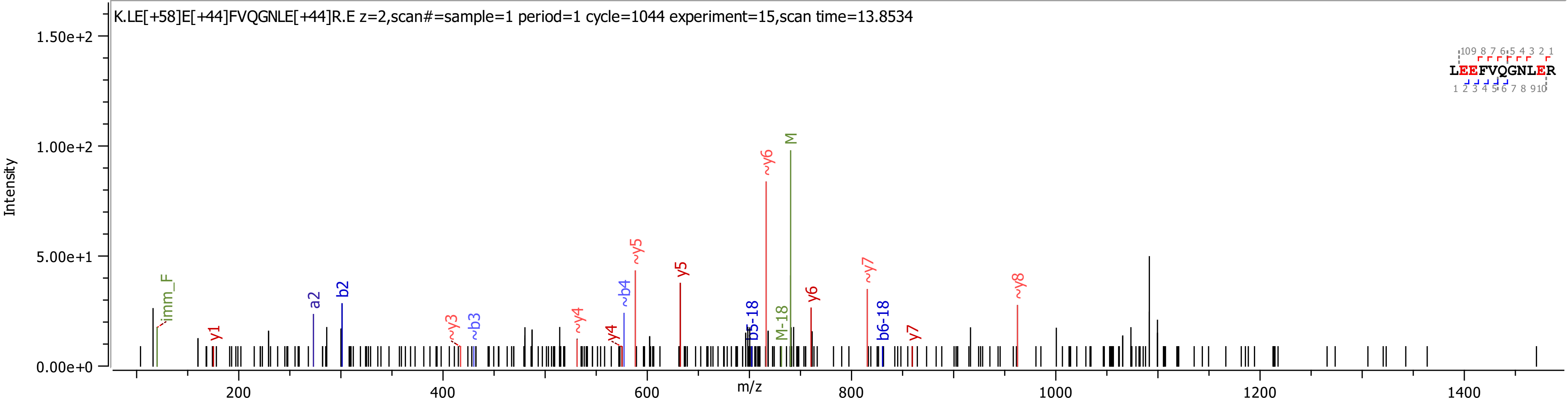

K.LE[+58]E[+44]FVQGNLE[+44]R.E z=2,scan#=sample=1 period=1 cycle=1044 experiment=15,scan time=13.8534
Intensity
m/z
1.50e+2
1.00e+2
5.00e+1
0.00e+0
200
400
600
800
1000
1200
1400
LEEFVQGNLER
imm_F
y1
a2
b2
~y3
~b3
~y4
y4
~b4
~y5
y5
b5-18
~y6
M-18
M
y6
~y7
b6-18
y7
~y8

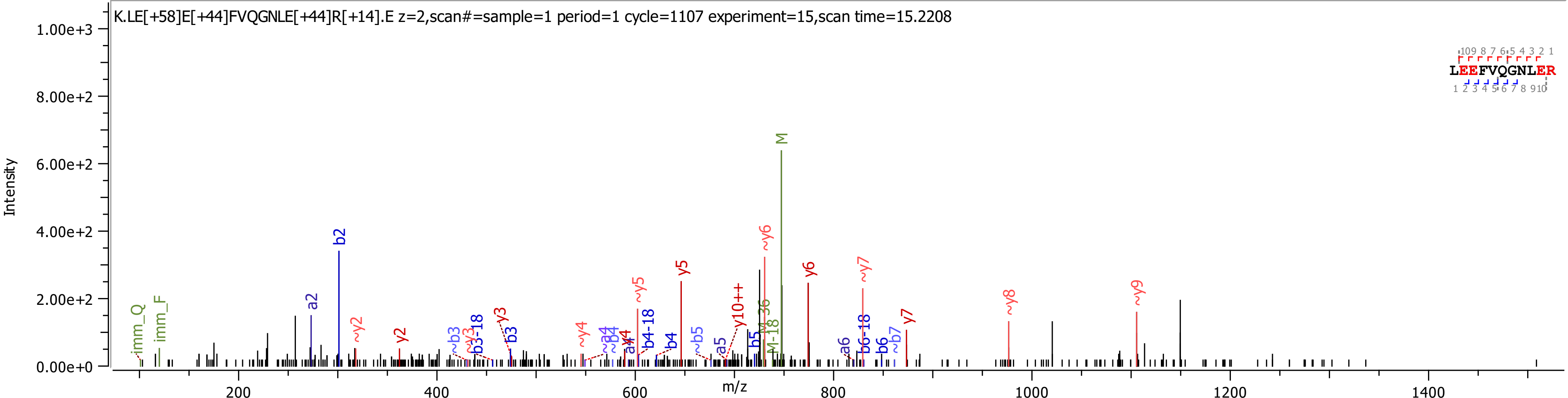
K.LE[+58]E[+44]FVQGNLE[+44]R[+14].E z=2,scan#=sample=1 period=1 cycle=1107 experiment=15,scan time=15.2208
Intensity
1.00e+3
8.00e+2
6.00e+2
4.00e+2
2.00e+2
0.00e+0
m/z
200
400
600
800
1000
1200
1400
LEEFVQGNLER
imm_Q
imm_F
a2
b2
~y2
y2
~b3
~y3
b3-18
y3
b3
~y4
~a4
~b4
a4
y4
~y5
b4-18
b4
y5
~b5
a5
y10++
b5
M-36
M-18
~y6
M
y6
a6
b6-18
~y7
b6
~b7
y7
~y8
~y9

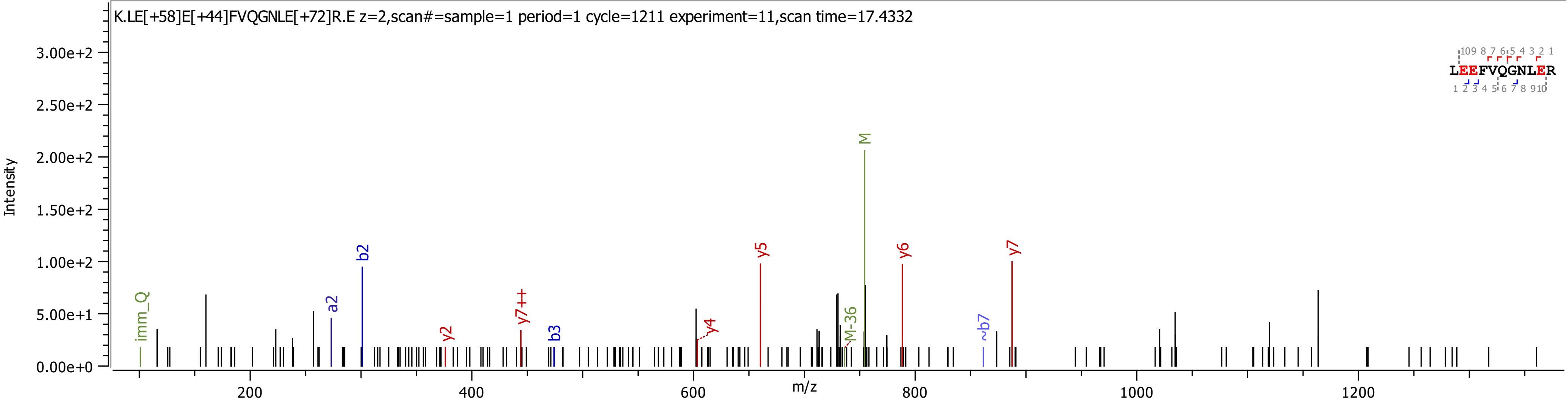

K.LE[+58]E[+44]FVQGNLE[+72]R.E z=2,scan#=sample=1 period=1 cycle=1211 experiment=11,scan time=17.4332
Intensity
m/z
imm_Q
a2
b2
y2
y7++
b3
y4
y5
M-36
M
y6
~b7
y7
LEEFVQGNLER

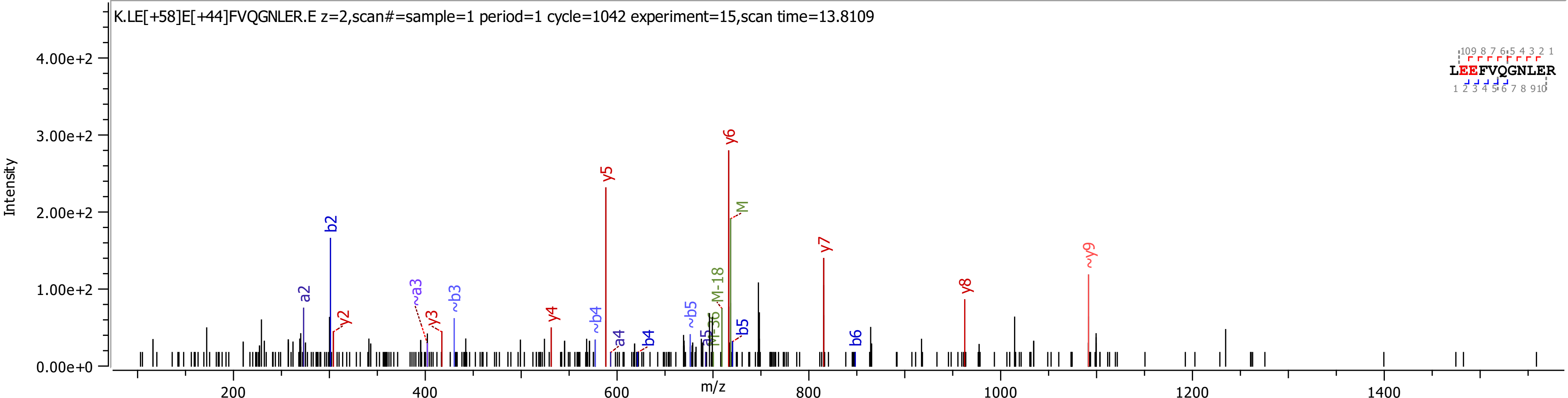
K.LE[+58]E[+44]FVQGNLER.E z=2,scan#=sample=1 period=1 cycle=1042 experiment=15,scan time=13.8109
LEEFVQGNLER
Intensity
m/z
4.00e+2
3.00e+2
2.00e+2
1.00e+2
0.00e+0
200
400
600
800
1000
1200
1400
a2
b2
y2
~a3
y3
~b3
y4
y5
~b4
a4
b4
~b5
M-18
y6
M
b5
y7
b6
y8
~y9

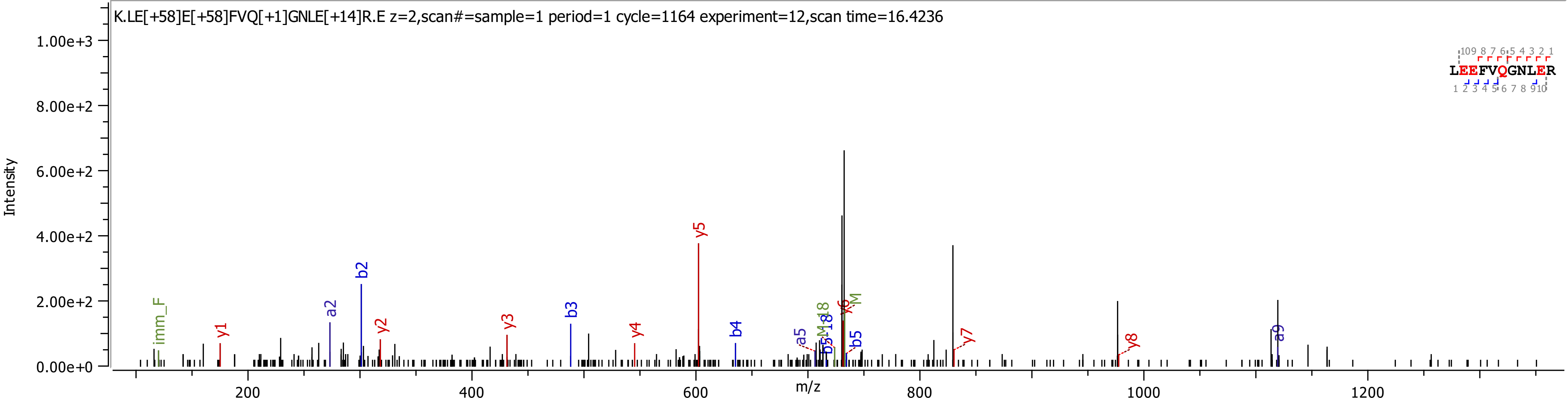

K.LE[+58]E[+58]FVQ[+1]GNLE[+14]R.E z=2,scan#=sample=1 period=1 cycle=1164 experiment=12,scan time=16.4236
Intensity
1.00e+3
8.00e+2
6.00e+2
4.00e+2
2.00e+2
0.00e+0
m/z
200
400
600
800
1000
1200
imm_F
y1
a2
b2
y2
y3
b3
y4
y5
b4
a5
b5
y6
b5
y7
y8
a9
LEEFVQGNLER

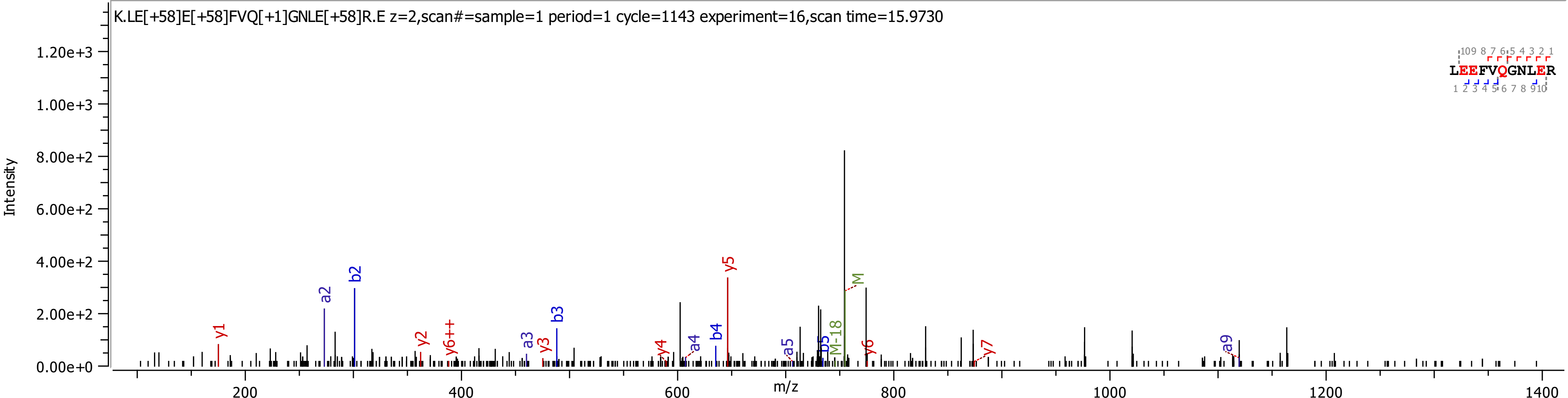
K.LE[+58]E[+58]FVQ[+1]GNLE[+58]R.E z=2,scan#=sample=1 period=1 cycle=1143 experiment=16,scan time=15.9730
LEEFVQGNLER
Intensity
m/z
y1
a2
b2
y2
y6++
a3
y3
b3
y4
a4
b4
y5
a5
b5
M-18
M
y6
y7
a9

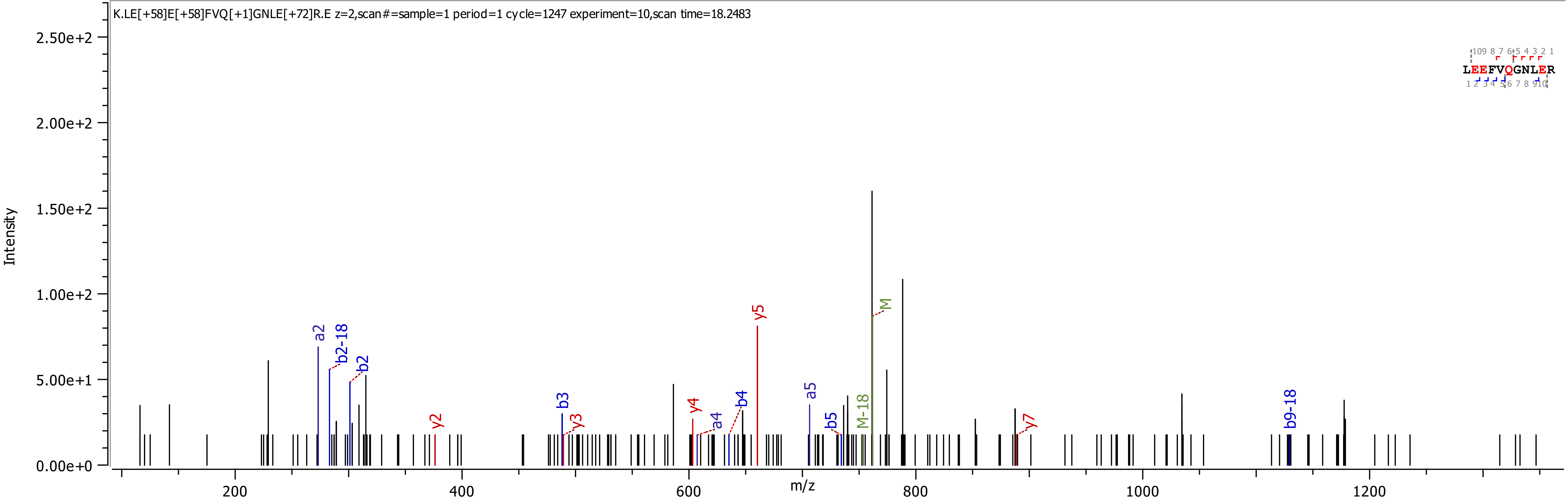

K.LE[+58]E[+58]FVQ[+1]GNLE[+72]R.E z=2,scan#=sample=1 period=1 cycle=1247 experiment=10,scan time=18.2483
Intensity
m/z
2.50e+2
2.00e+2
1.50e+2
1.00e+2
5.00e+1
0.00e+0
200
400
600
800
1000
1200
LEEFVQGNLER
a2
b2-18
b2
y2
b3
y3
y4
a4
b4
y5
a5
b5
M-18
M
y7
b9-18

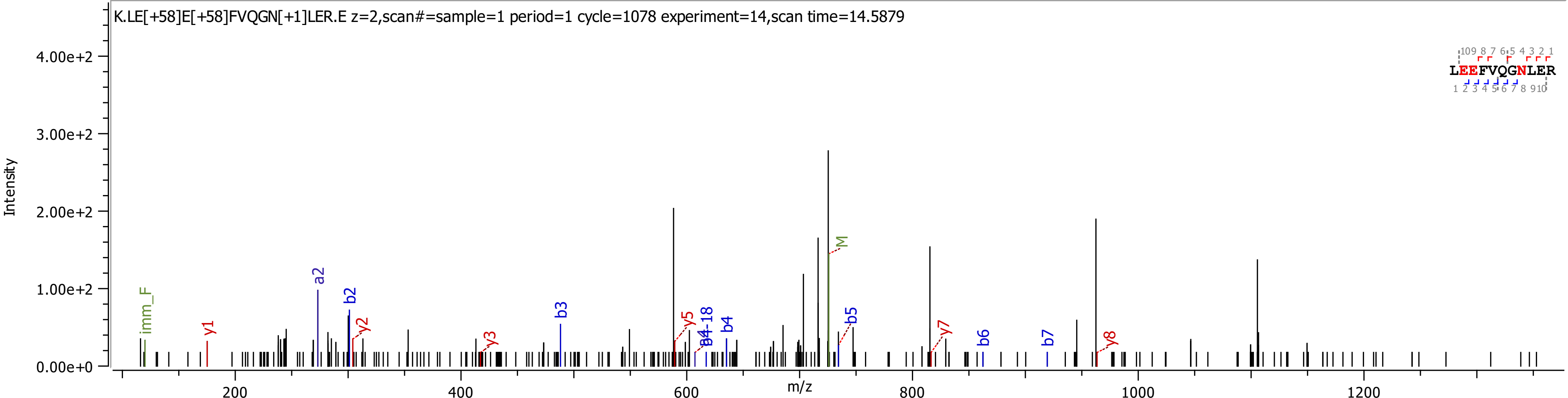
K.LE[+58]E[+58]FVQGN[+1]LER.E z=2,scan#=sample=1 period=1 cycle=1078 experiment=14,scan time=14.5879
Intensity
4.00e+2
3.00e+2
2.00e+2
1.00e+2
0.00e+0
m/z
200
400
600
800
1000
1200
imm_F
y1
a2
b2
y2
y3
b3
y5
b4-18
b4
M
b5
y7
b6
b7
y8
LEEFVQGNLER

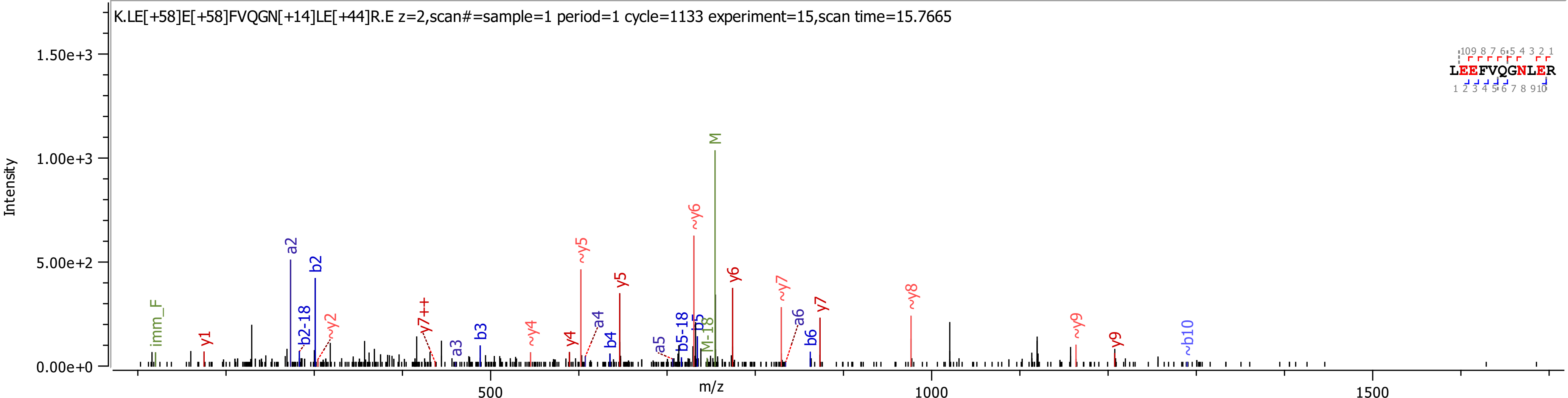
K.LE[+58]E[+58]FVQGN[+14]LE[+44]R.E z=2,scan#=sample=1 period=1 cycle=1133 experiment=15,scan time=15.7665
Intensity
m/z
LEEFVQGNLER

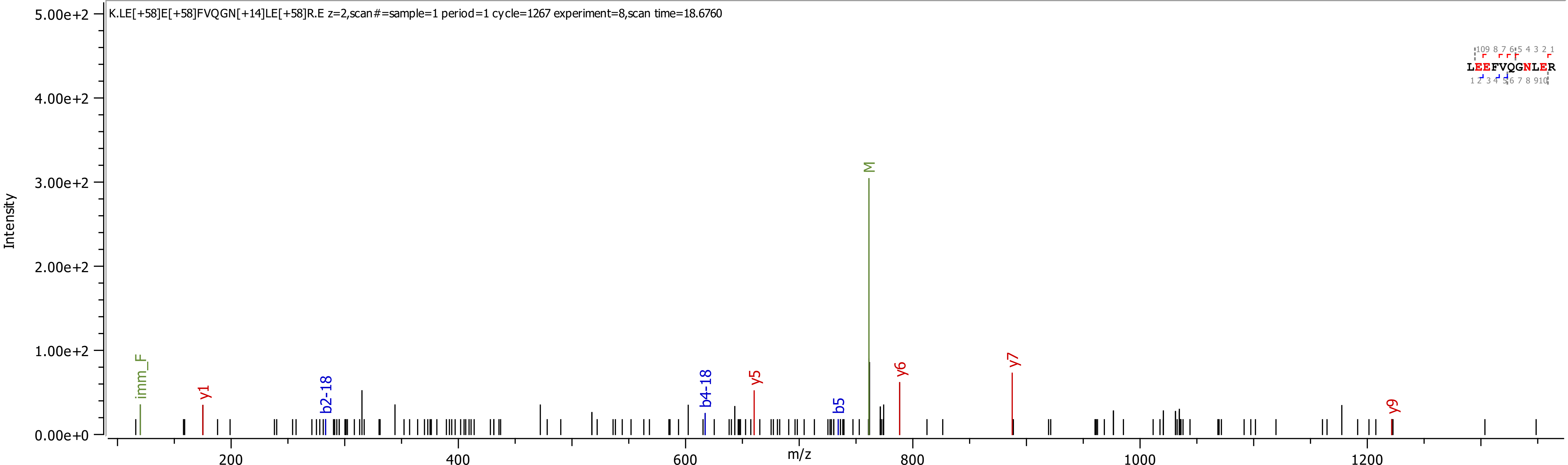
K.LE[+58]E[+58]FVQGN[+14]LE[+58]R.E z=2,scan#=sample=1 period=1 cycle=1267 experiment=8,scan time=18.6760
Intensity
5.00e+2
4.00e+2
3.00e+2
2.00e+2
1.00e+2
0.00e+0
m/z
200
400
600
800
1000
1200
imm_F
y1
b2-18
b4-18
y5
b5
M
y6
y7
y9
LEEFVQGNLER

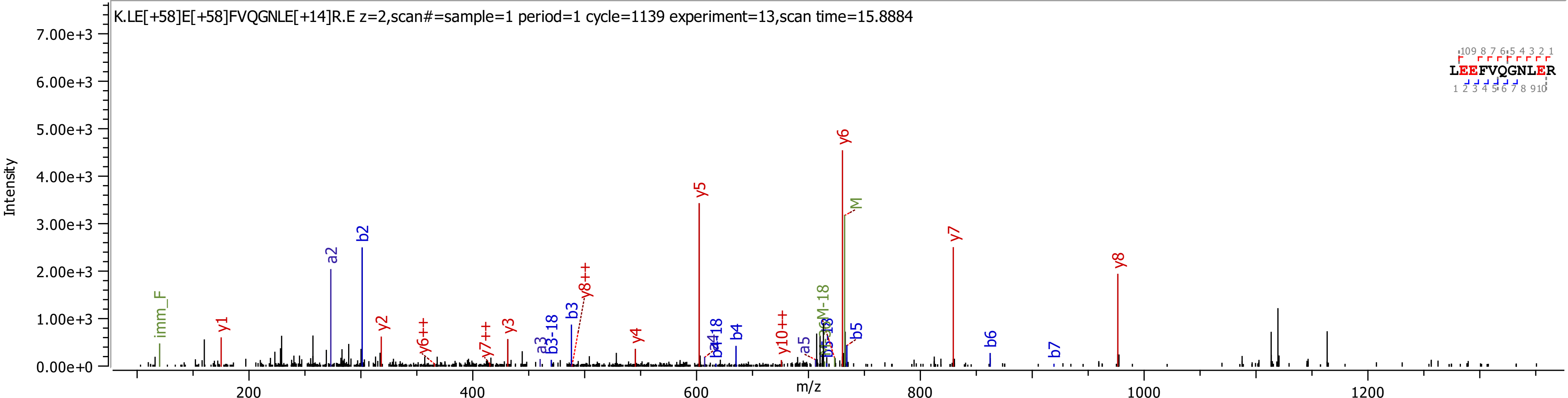

K.LE[+58]E[+58]FVQGNLE[+14]R.E z=2,scan#=sample=1 period=1 cycle=1139 experiment=13,scan time=15.8884
Intensity
m/z
7.00e+3
6.00e+3
5.00e+3
4.00e+3
3.00e+3
2.00e+3
1.00e+3
0.00e+0
200
400
600
800
1000
1200
LEEFVQGNLER
imm_F
y1
a2
b2
y2
y6++
y7++
y3
a3
b3-18
b3
y8++
y4
y5
b4
y10++
a5
y6
M
y7
y8
b5
b6
b7

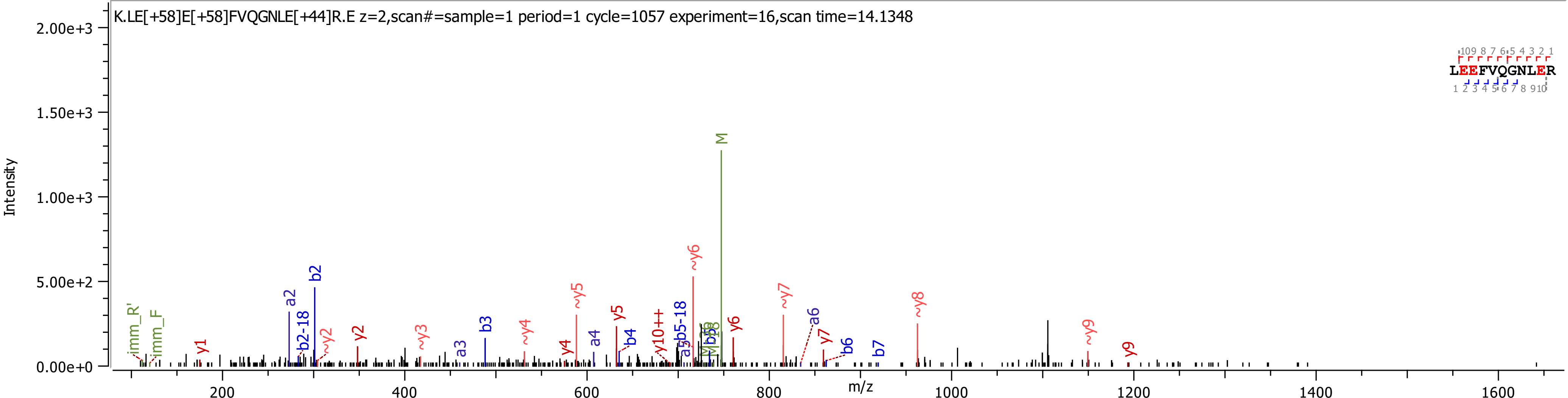
K.LE[+58]E[+58]FVQGNLE[+44]R.E z=2,scan#=sample=1 period=1 cycle=1057 experiment=16,scan time=14.1348
LEEFVQGNLER
Intensity
m/z
2.00e+3
1.50e+3
1.00e+3
5.00e+2
0.00e+0
200
400
600
800
1000
1200
1400
1600
M
~y6
b2
a2
b2-18
~y5
y5
y10++
b5-18
y6
~y7
a6
y7
b6
b7
~y8
~y9
y9
imm_R'
imm_F
y1
~y2
y2
~y3
a3
b3
~y4
y4
a4
b4

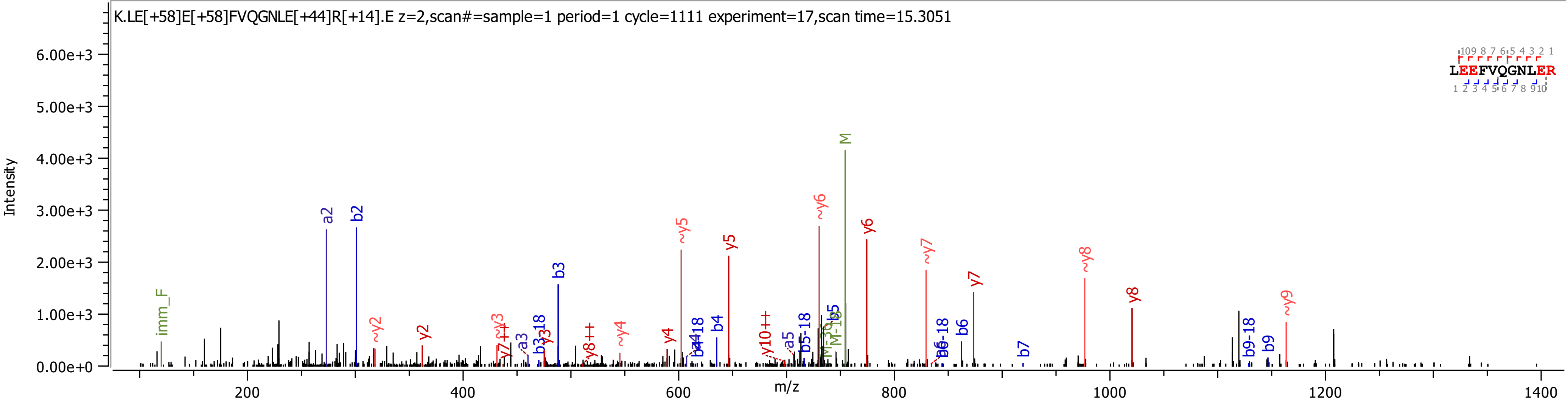
K.LE[+58]E[+58]FVQGNLE[+44]R[+14].E z=2,scan#=sample=1 period=1 cycle=1111 experiment=17,scan time=15.3051
Intensity
m/z
LEEFVQGNLER
M
a2
b2
b3
y5
y6
~y5
~y6
~y7
~y8
~y9
y7
y8
y2
~y2
imm_F
b4
b6
b7
b9
b9-18
y4
~y4
a5
b5-18
b6-18
y10++
y8++
a3
b3-18
y3
~y3
y7++

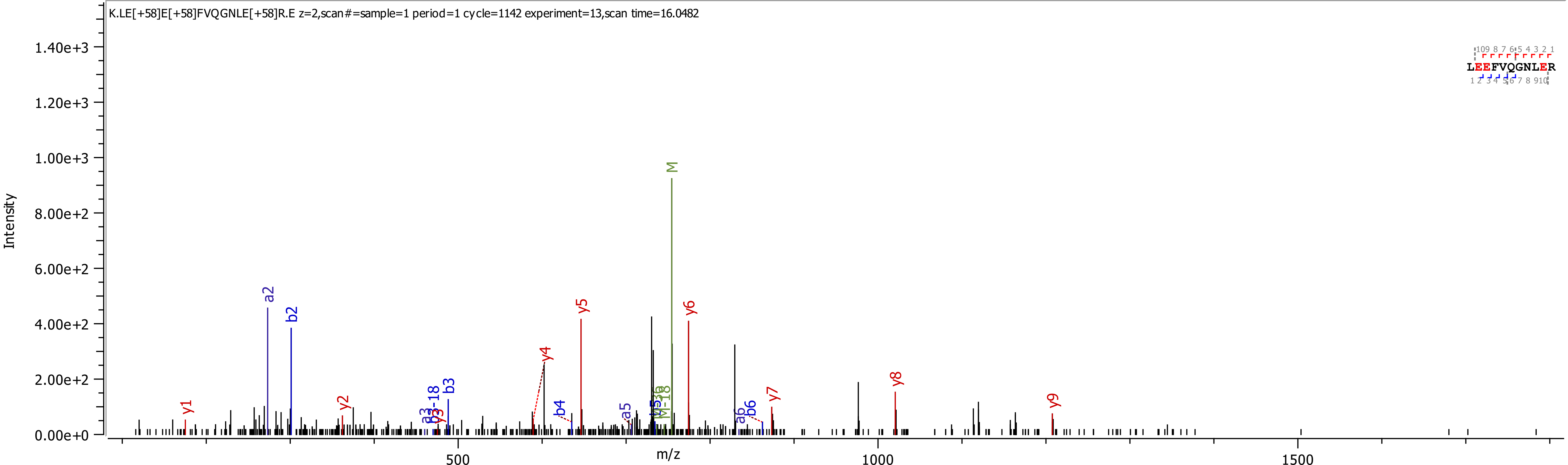
K.LE[+58]E[+58]FVQGNLE[+58]R.E z=2,scan#=sample=1 period=1 cycle=1142 experiment=13,scan time=16.0482
LEEFVQGNLER
Intensity
m/z
1.40e+3
1.20e+3
1.00e+3
8.00e+2
6.00e+2
4.00e+2
2.00e+2
0.00e+0
500
1000
1500
M
a2
b2
y5
y6
y4
y8
y1
y2
b3
y7
y9
b4
a5
b6

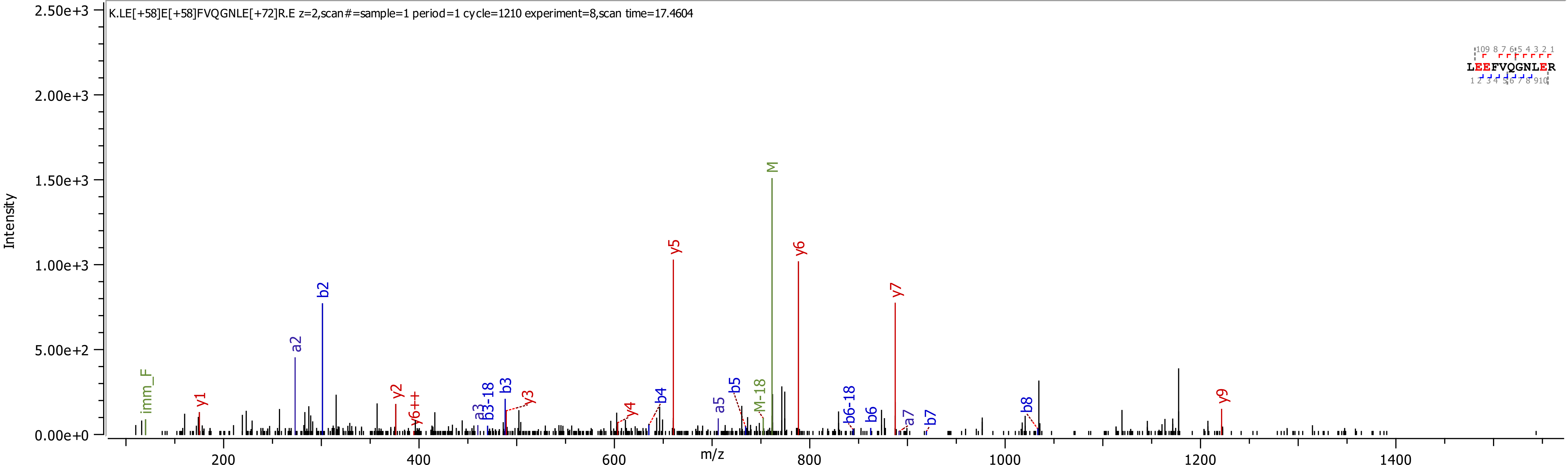
K.LE[+58]E[+58]FVQGNLE[+72]R.E z=2,scan#=sample=1 period=1 cycle=1210 experiment=8,scan time=17.4604
LEEFVQGNLER
Intensity
m/z
2.50e+3
2.00e+3
1.50e+3
1.00e+3
5.00e+2
0.00e+0
200
400
600
800
1000
1200
1400
imm_F
y1
a2
b2
y2
y6++
a3
b3-18
b3
y3
y4
b4
y5
a5
b5
M-18
M
y6
b6-18
b6
y7
a7
b7
b8
y9

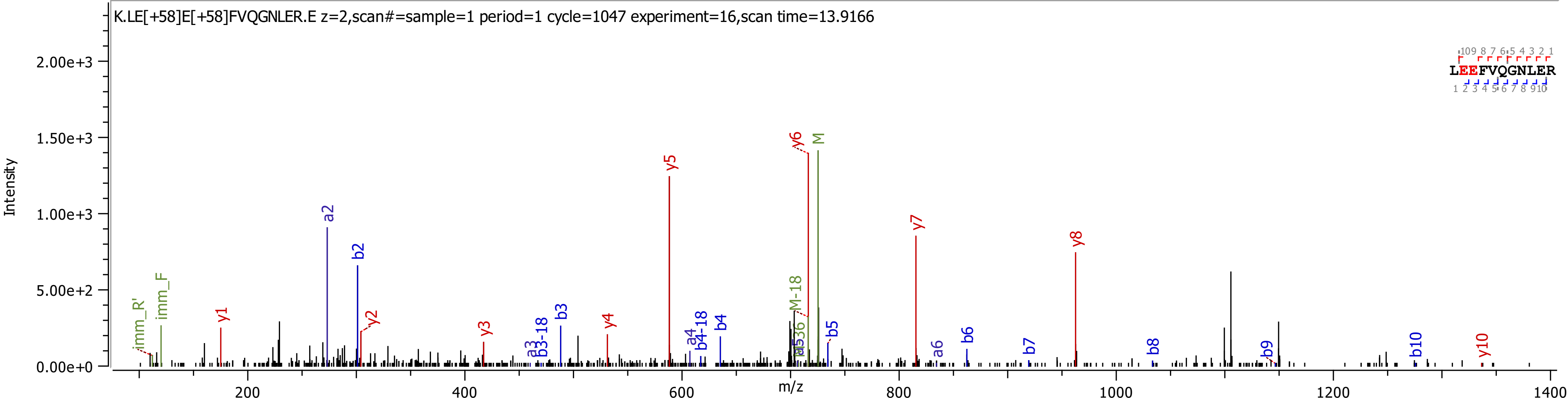

K.LE[+58]E[+58]FVQGNLER.E z=2,scan#=sample=1 period=1 cycle=1047 experiment=16,scan time=13.9166
Intensity
m/z
LEEFVQGNLER
imm_R'
imm_F
y1
a2
b2
y2
y3
a3
b3-18
b3
y4
y5
a4
b4-18
b4
M-18
y6
M
b5
y7
a6
b6
b7
y8
b8
b9
b10
y10

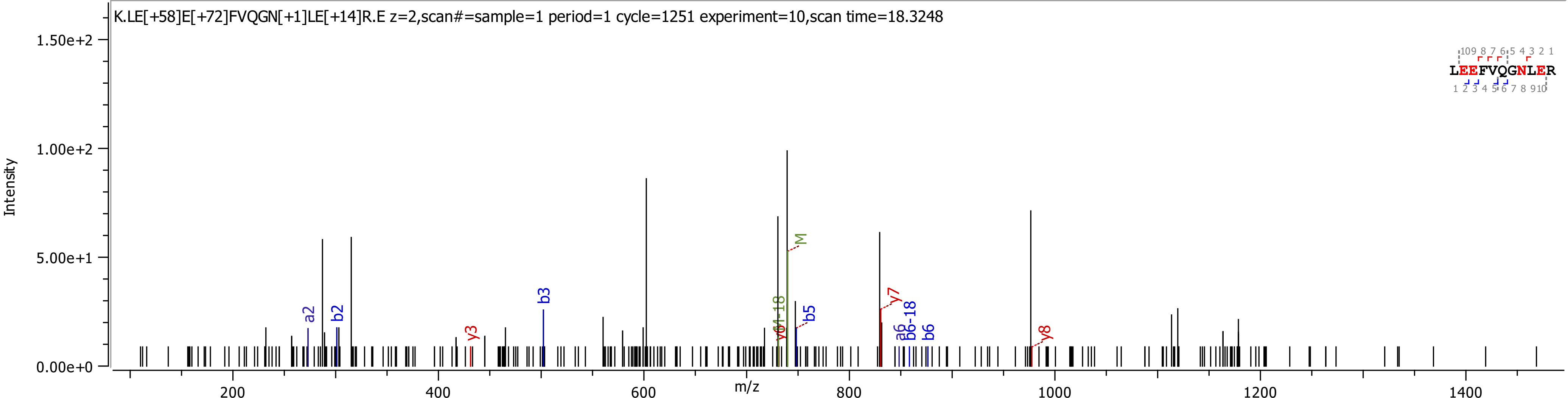
K.LE[+58]E[+72]FVQGN[+1]LE[+14]R.E z=2,scan#=sample=1 period=1 cycle=1251 experiment=10,scan time=18.3248
Intensity
m/z
LEEFVQGNLER
a2
b2
y3
b3
y6
M-18
M
b5
y7
a6
b6-18
b6
y8

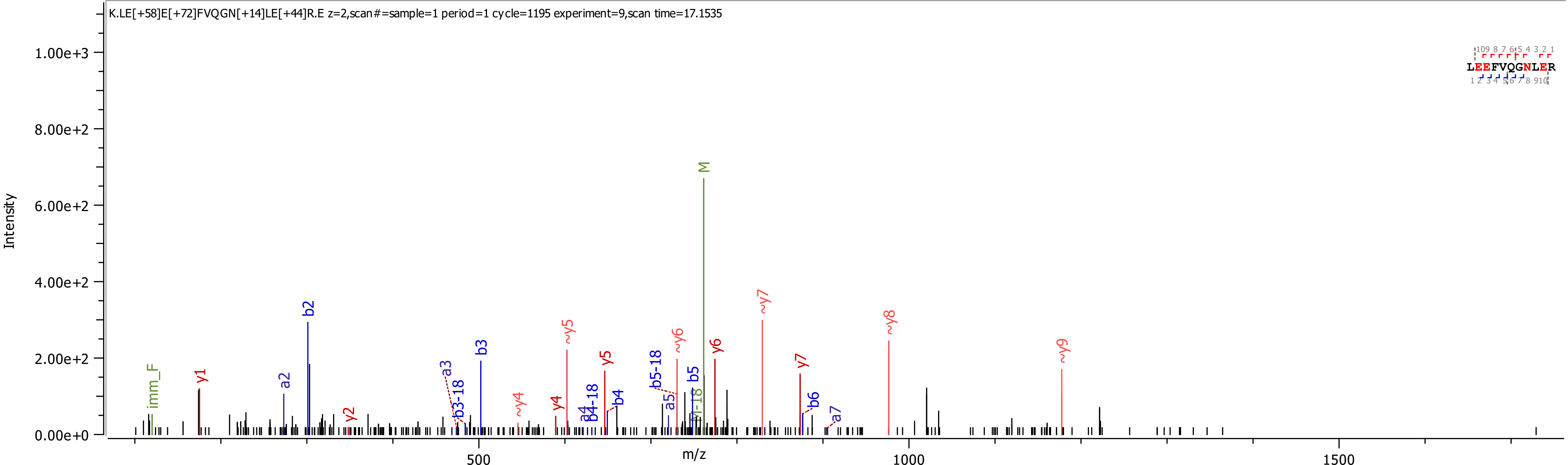
K.LE[+58]E[+72]FVQGN[+14]LE[+44]R.E z=2,scan#=sample=1 period=1 cycle=1195 experiment=9,scan time=17.1535
Intensity
m/z
1.00e+3
8.00e+2
6.00e+2
4.00e+2
2.00e+2
0.00e+0
500
1000
1500
LEEFVQGNLER
imm_F
y1
a2
b2
y2
a3
b3-18
b3
~y4
y4
~y5
a4
b4-18
y5
b4
b5-18
a5
~y6
b5
M-18
M
y6
~y7
y7
b6
a7
~y8
~y9

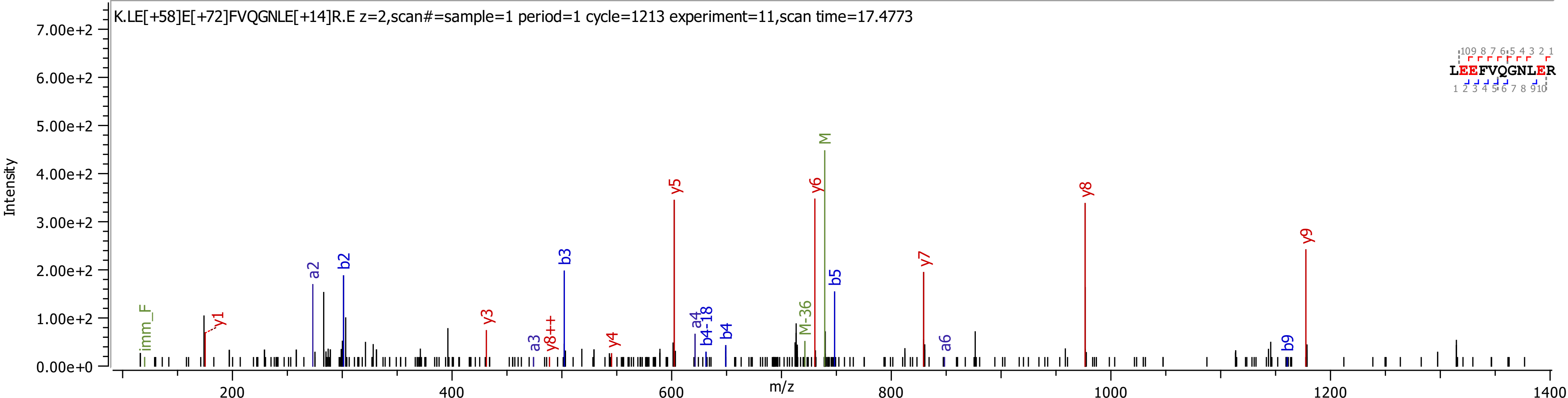
K.LE[+58]E[+72]FVQGNLE[+14]R.E z=2,scan#=sample=1 period=1 cycle=1213 experiment=11,scan time=17.4773
Intensity
m/z
LEEFVQGNLER
imm_F
y1
a2
b2
y3
a3
y8++
b3
y4
y5
a4
b4-18
b4
M-36
y6
M
b5
y7
a6
y8
b9
y9

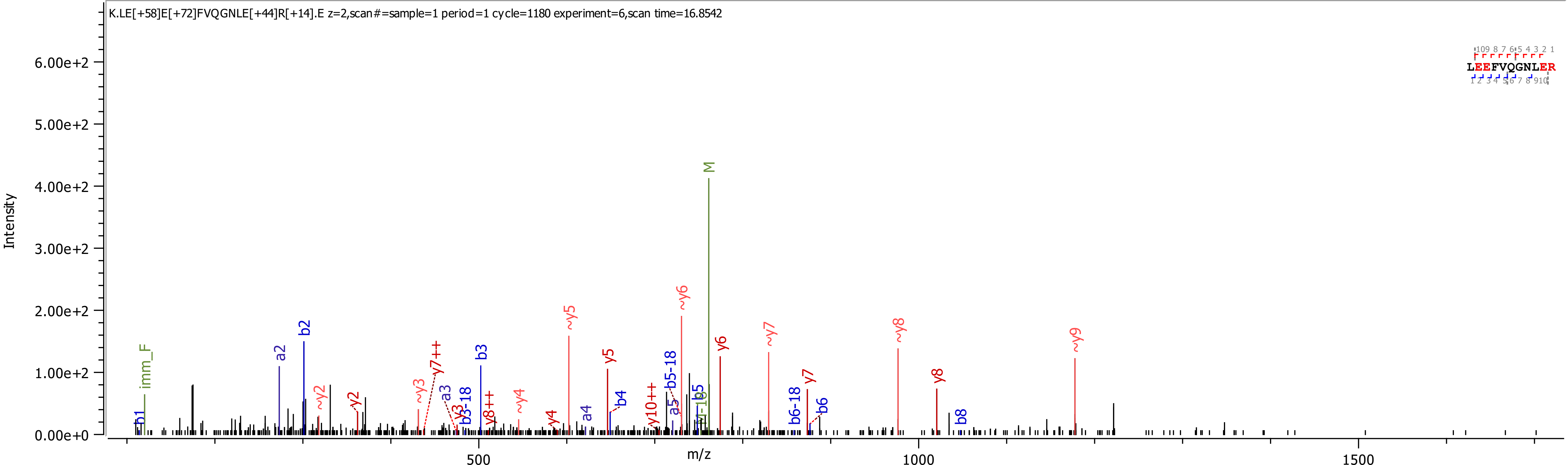
K.LE[+58]E[+72]FVQGNLE[+44]R[+14].E z=2,scan#=sample=1 period=1 cycle=1180 experiment=6,scan time=16.8542
LEEFVQGNLER
Intensity
m/z
0.00e+0
1.00e+2
2.00e+2
3.00e+2
4.00e+2
5.00e+2
6.00e+2
500
1000
1500
M
imm_F
b1
a2
b2
~y2
y2
~y3
y7++
a3
y3
b3-18
b3
y8++
~y4
y4
~y5
a4
y5
b4
y10++
a5
b5-18
~y6
b5
y6
~y7
b6-18
y7
b6
~y8
y8
b8
~y9

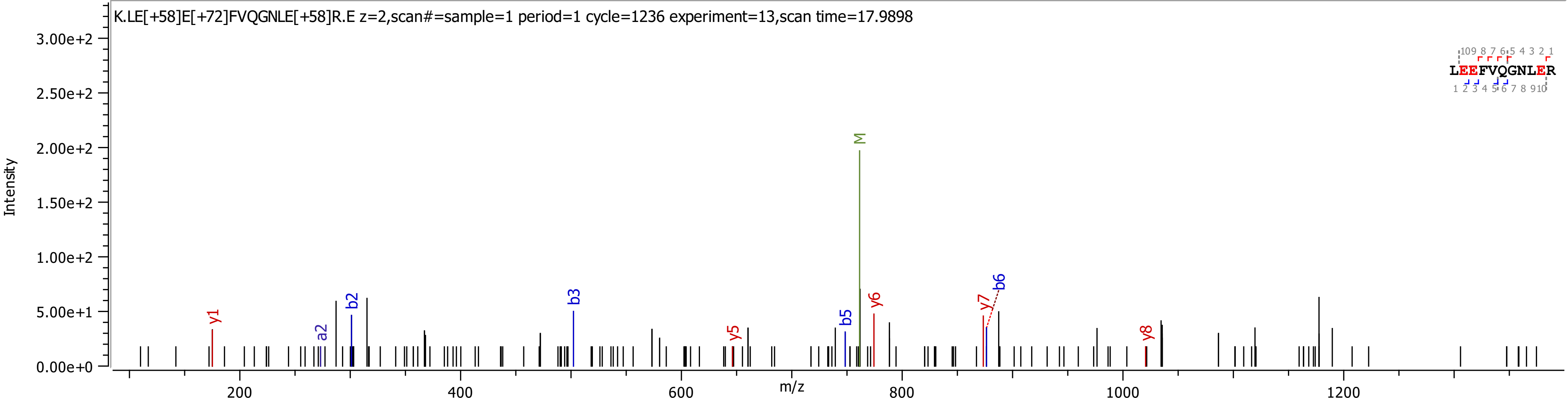
K.LE[+58]E[+72]FVQGNLE[+58]R.E z=2,scan#=sample=1 period=1 cycle=1236 experiment=13,scan time=17.9898
Intensity
m/z
LEEFVQGNLER
y1
a2
b2
b3
y5
b5
M
y6
y7
b6
y8

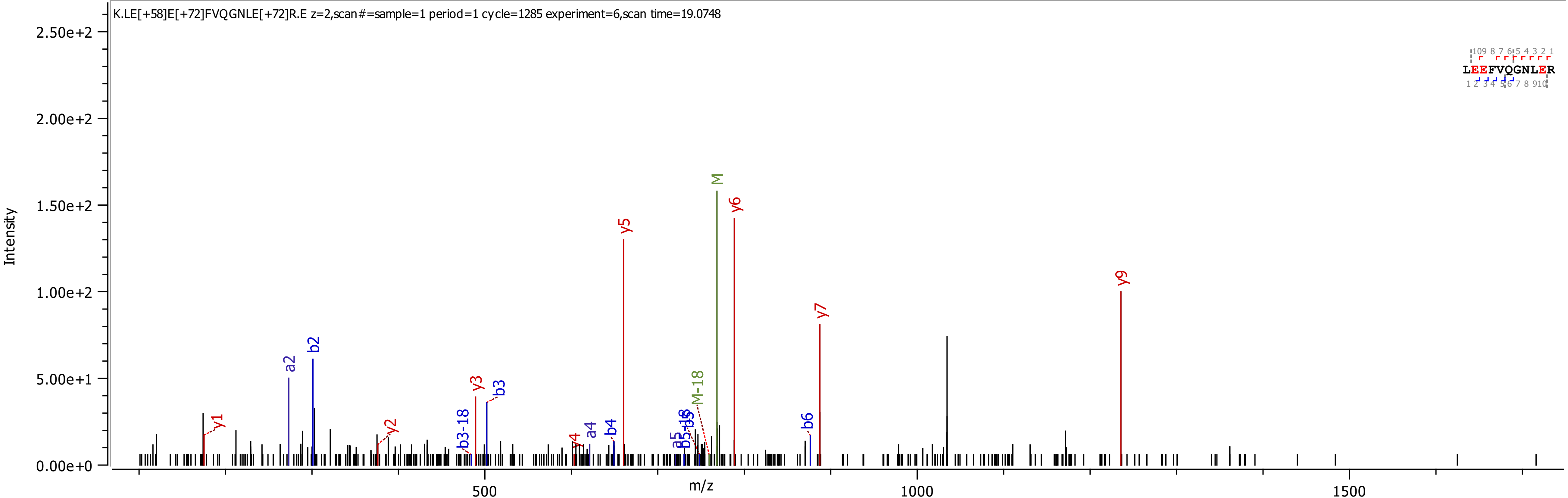

K.LE[+58]E[+72]FVQGNLE[+72]R.E z=2,scan#=sample=1 period=1 cycle=1285 experiment=6,scan time=19.0748
Intensity
2.50e+2
2.00e+2
1.50e+2
1.00e+2
5.00e+1
0.00e+0
m/z
500
1000
1500
LEEFVQGNLER
y1
a2
b2
y2
b3-18
y3
b3
y4
a4
b4
y5
a5
b5-18
b5
M-18
M
y6
b6
y7
y9

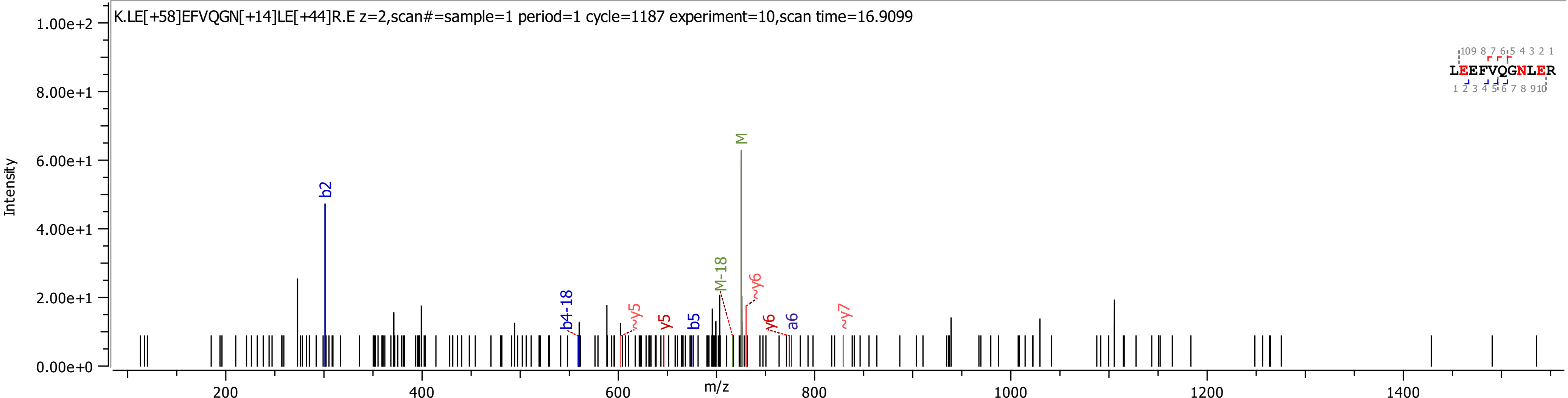
K.LE[+58]EFVQGN[+14]LE[+44]R.E z=2,scan#=sample=1 period=1 cycle=1187 experiment=10,scan time=16.9099
Intensity
m/z
1.00e+2
8.00e+1
6.00e+1
4.00e+1
2.00e+1
0.00e+0
200
400
600
800
1000
1200
1400
b2
b4-18
~y5
y5
b5
M-18
M
~y6
y6
a6
~y7
LEEFVQGNLER

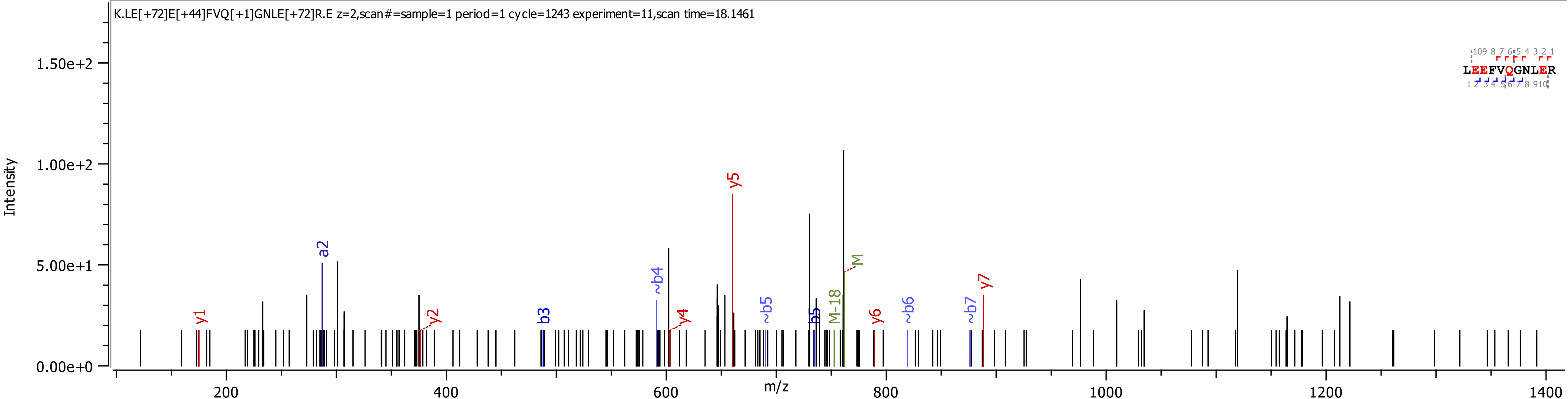

K.LE[+72]E[+44]FVQ[+1]GNLE[+72]R.E z=2,scan#=sample=1 period=1 cycle=1243 experiment=11,scan time=18.1461
Intensity
m/z
y1
a2
y2
b3
~b4
y4
y5
~b5
b5
M-18
M
y6
~b6
~b7
y7
LEEFVQGNLER

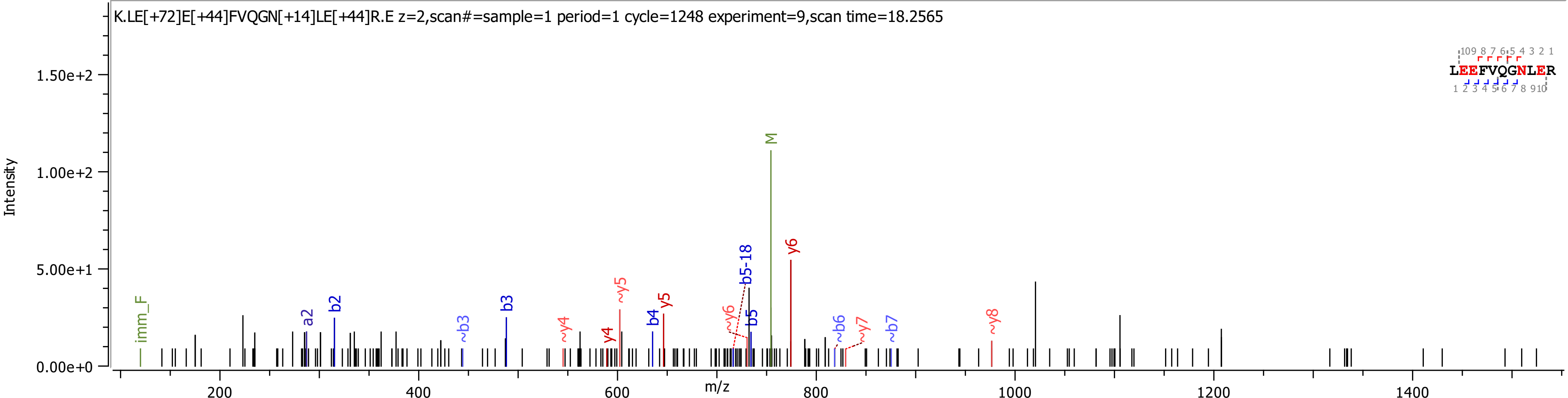
K.LE[+72]E[+44]FVQGN[+14]LE[+44]R.E z=2,scan#=sample=1 period=1 cycle=1248 experiment=9,scan time=18.2565
Intensity
m/z
0.00e+0
5.00e+1
1.00e+2
1.50e+2
200
400
600
800
1000
1200
1400
imm_F
a2
b2
~b3
b3
~y4
y4
~y5
y5
b4
~y6
b5-18
b5
M
y6
~b6
~y7
~b7
~y8
LEEFVQGNLER
10 9 8 7 6 5 4 3 2 1
1 2 3 4 5 6 7 8 9 10

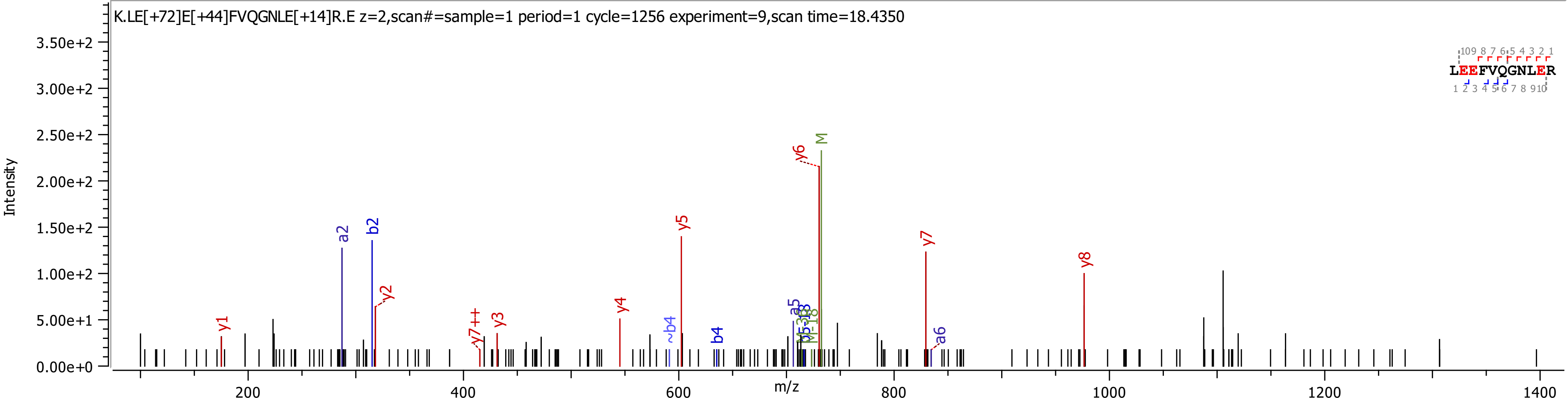

K.LE[+72]E[+44]FVQGNLE[+14]R.E z=2,scan#=sample=1 period=1 cycle=1256 experiment=9,scan time=18.4350
LEEFVQGNLER
Intensity
m/z
0.00e+0
5.00e+1
1.00e+2
1.50e+2
2.00e+2
2.50e+2
3.00e+2
3.50e+2
200
400
600
800
1000
1200
1400
y1
a2
b2
y2
y7++
y3
y4
~b4
y5
b4
a5
y6
M
y7
a6
y8

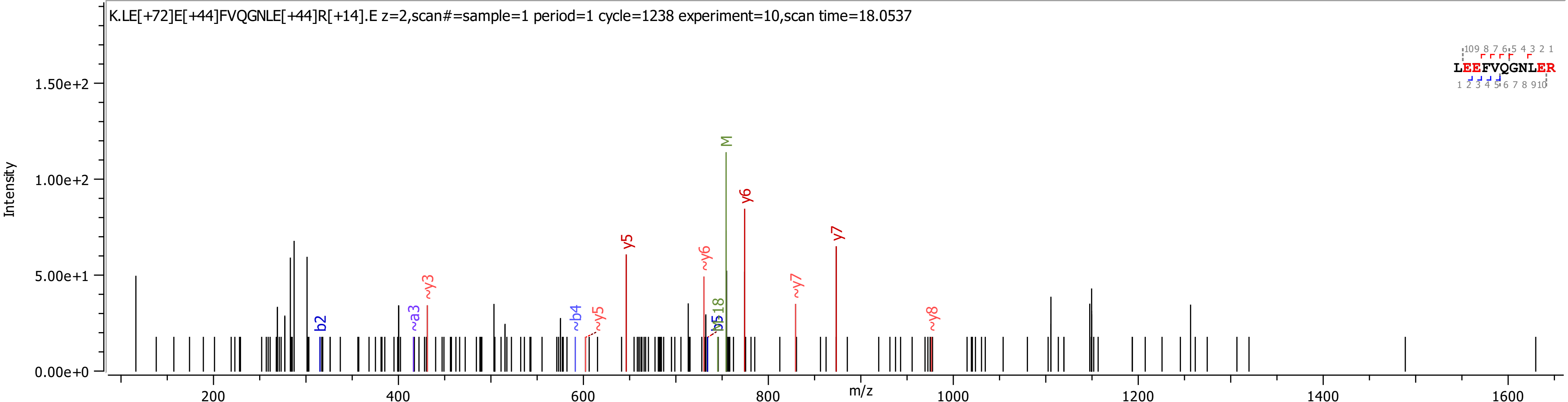
K.LE[+72]E[+44]FVQGNLE[+44]R[+14].E z=2,scan#=sample=1 period=1 cycle=1238 experiment=10,scan time=18.0537
Intensity
0.00e+0
5.00e+1
1.00e+2
1.50e+2
m/z
200
400
600
800
1000
1200
1400
1600
LEEFVQGNLER
b2
~a3
~y3
~b4
~y5
y5
~y6
b5
b518
M
y6
~y7
y7
~y8

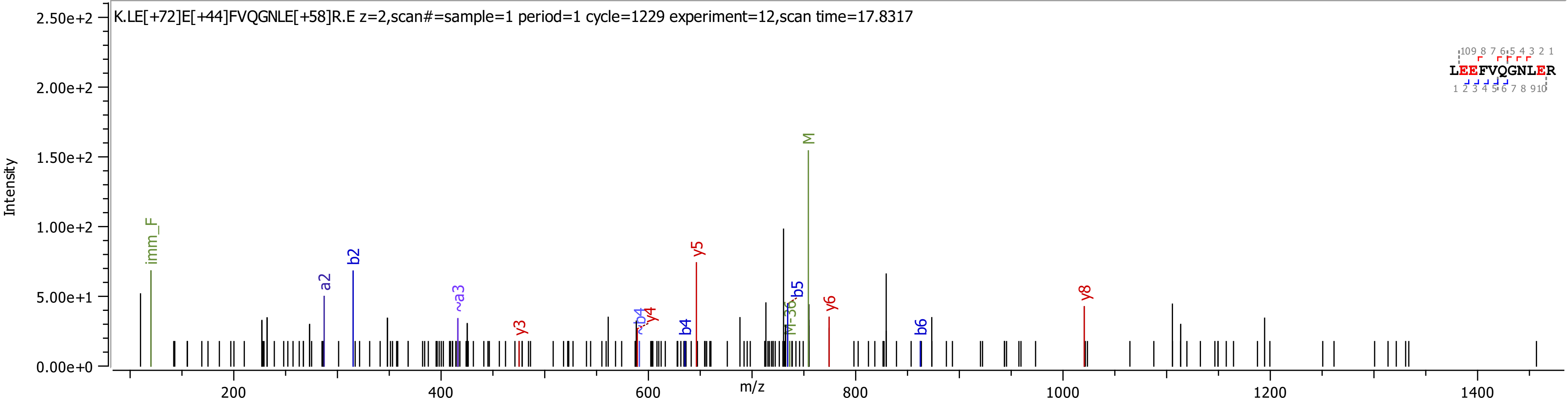
K.LE[+72]E[+44]FVQGNLE[+58]R.E z=2,scan#=sample=1 period=1 cycle=1229 experiment=12,scan time=17.8317
Intensity
m/z
LEEFVQGNLER
imm_F
a2
b2
~a3
y3
~b4
y4
b4
y5
M-36
b5
M
y6
b6
y8
2.50e+2
2.00e+2
1.50e+2
1.00e+2
5.00e+1
0.00e+0
200
400
600
800
1000
1200
1400

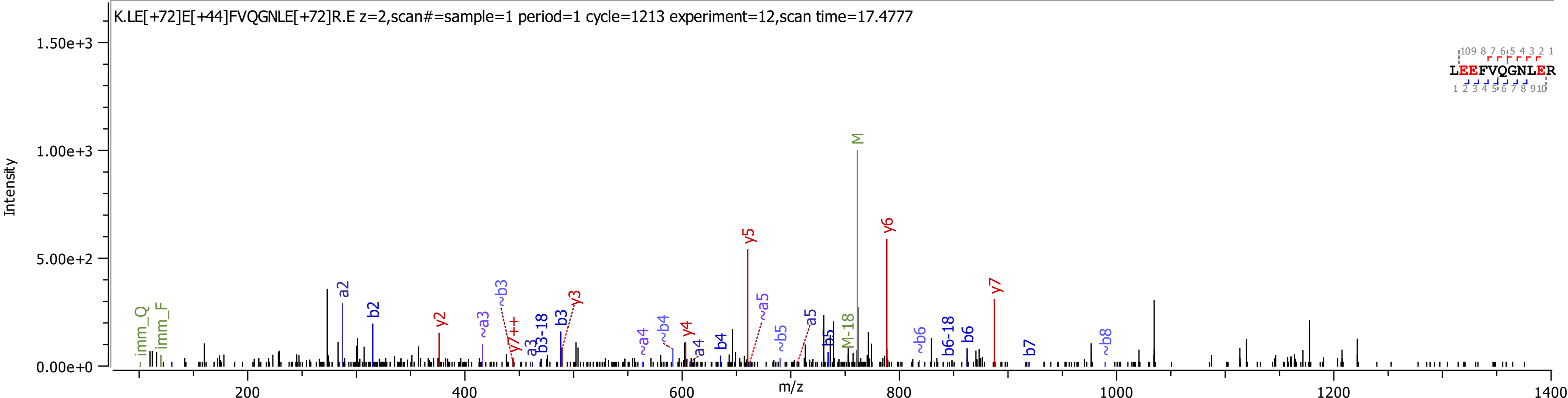
K.LE[+72]E[+44]FVQGNLE[+72]R.E z=2,scan#=sample=1 period=1 cycle=1213 experiment=12,scan time=17.4777
Intensity
m/z
LEEFVQGNLER
1.50e+3
1.00e+3
5.00e+2
0.00e+0
200
400
600
800
1000
1200
1400
imm_Q
imm_F
a2
b2
y2
~a3
~b3
y7++
a3
b3-18
b3
y3
~a4
~b4
y4
a4
b4
y5
~a5
~b5
a5
b5
M-18
M
y6
~b6
b6-18
b6
y7
b7
~b8

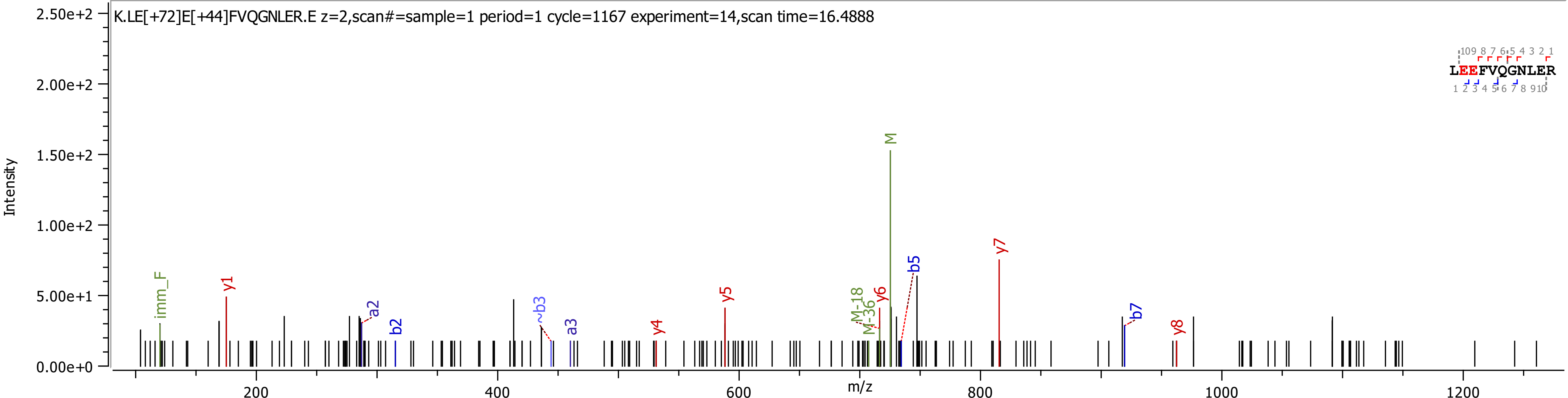
K.LE[+72]E[+44]FVQGNLER.E z=2,scan#=sample=1 period=1 cycle=1167 experiment=14,scan time=16.4888
Intensity
m/z
2.50e+2
2.00e+2
1.50e+2
1.00e+2
5.00e+1
0.00e+0
200
400
600
800
1000
1200
imm_F
y1
a2
b2
~b3
a3
y4
y5
M-18
M-36
y6
M
b5
y7
b7
y8
LEEFVQGNLER

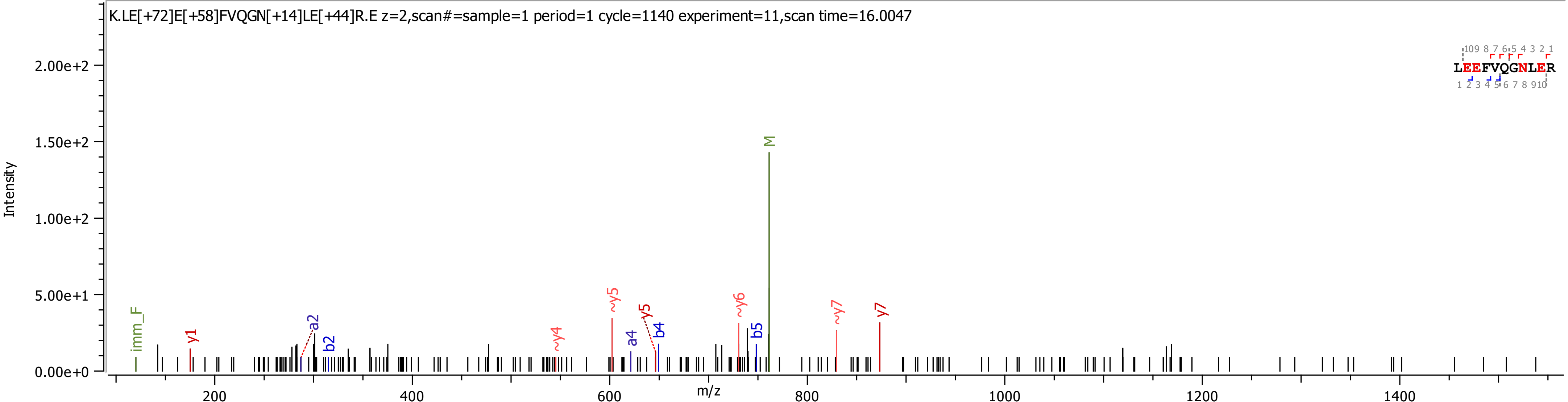

K.LE[+72]E[+58]FVQGN[+14]LE[+44]R.E z=2,scan#=sample=1 period=1 cycle=1140 experiment=11,scan time=16.0047
LEEFVQGNLER
Intensity
m/z
imm_F
y1
a2
b2
~y4
~y5
a4
y5
b4
~y6
b5
M
~y7
y7

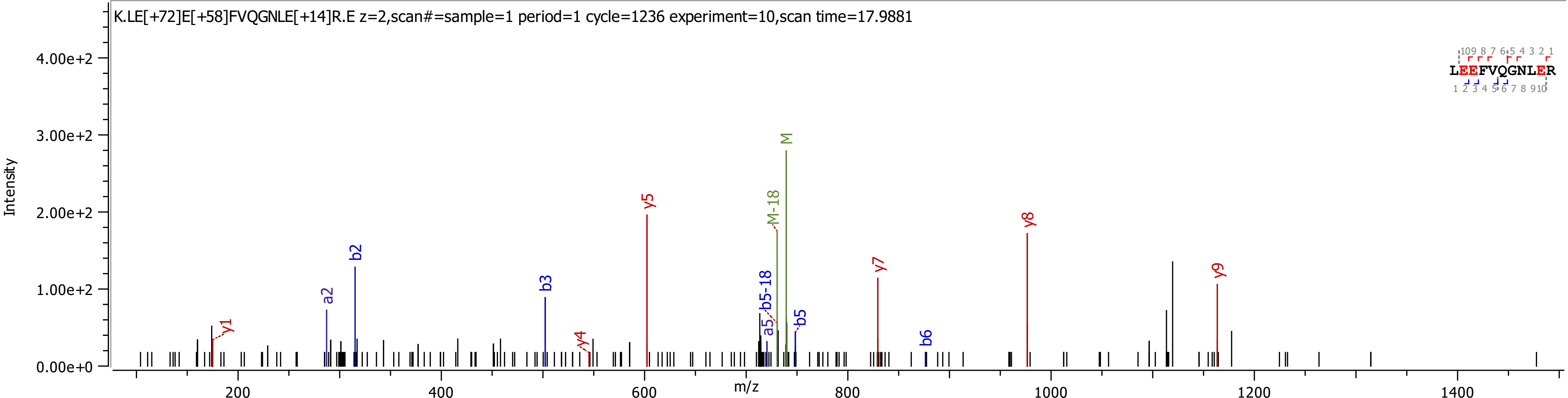

K.LE[+72]E[+58]FVQGNLE[+14]R.E z=2,scan#=sample=1 period=1 cycle=1236 experiment=10,scan time=17.9881
Intensity
m/z
LEEFVQGNLER
y1
a2
b2
b3
y4
y5
a5
b5-18
M-18
M
b5
y7
b6
y8
y9

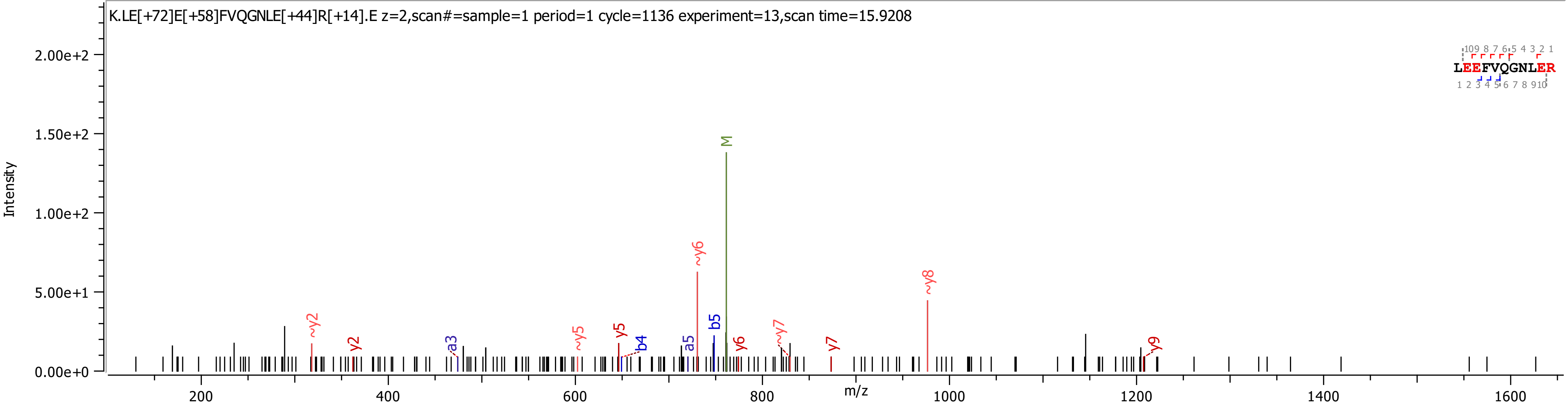
K.LE[+72]E[+58]FVQGNLE[+44]R[+14].E z=2,scan#=sample=1 period=1 cycle=1136 experiment=13,scan time=15.9208
Intensity
m/z
2.00e+2
1.50e+2
1.00e+2
5.00e+1
0.00e+0
200
400
600
800
1000
1200
1400
1600
M
~y6
~y8
~y2
y2
a3
~y5
y5
b4
a5
b5
y6
~y7
y7
y9
LEEFVQGNLER
10 9 8 7 6 5 4 3 2 1
1 2 3 4 5 6 7 8 9 10

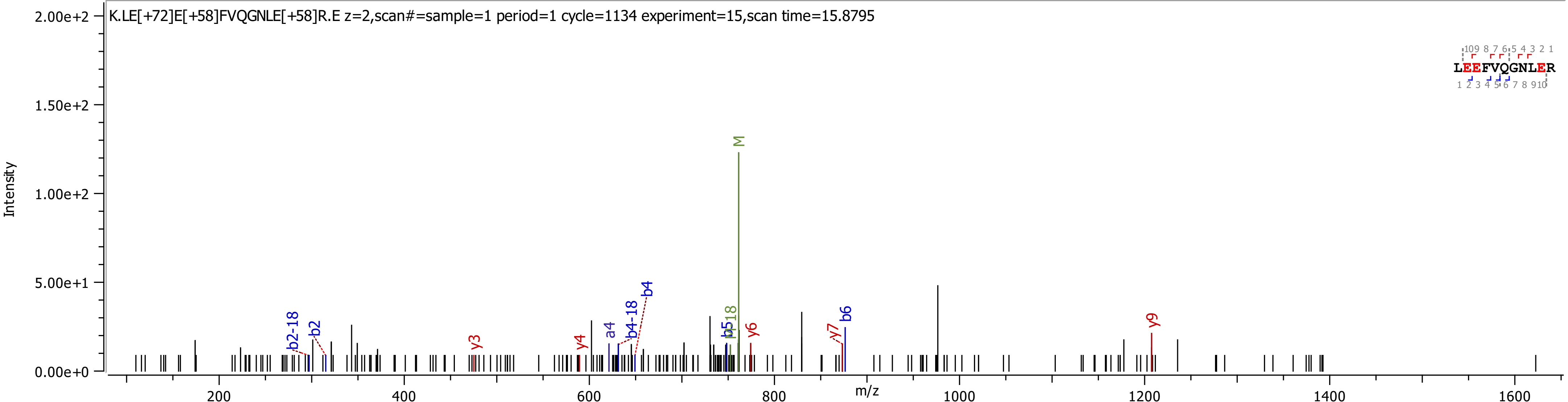

K.LE[+72]E[+58]FVQGNLE[+58]R.E z=2,scan#=sample=1 period=1 cycle=1134 experiment=15,scan time=15.8795
Intensity
m/z
M
b2-18
b2
y3
y4
a4
b4-18
b4
b5
M-18
y6
y7
b6
y9
LEEFVQGNLER

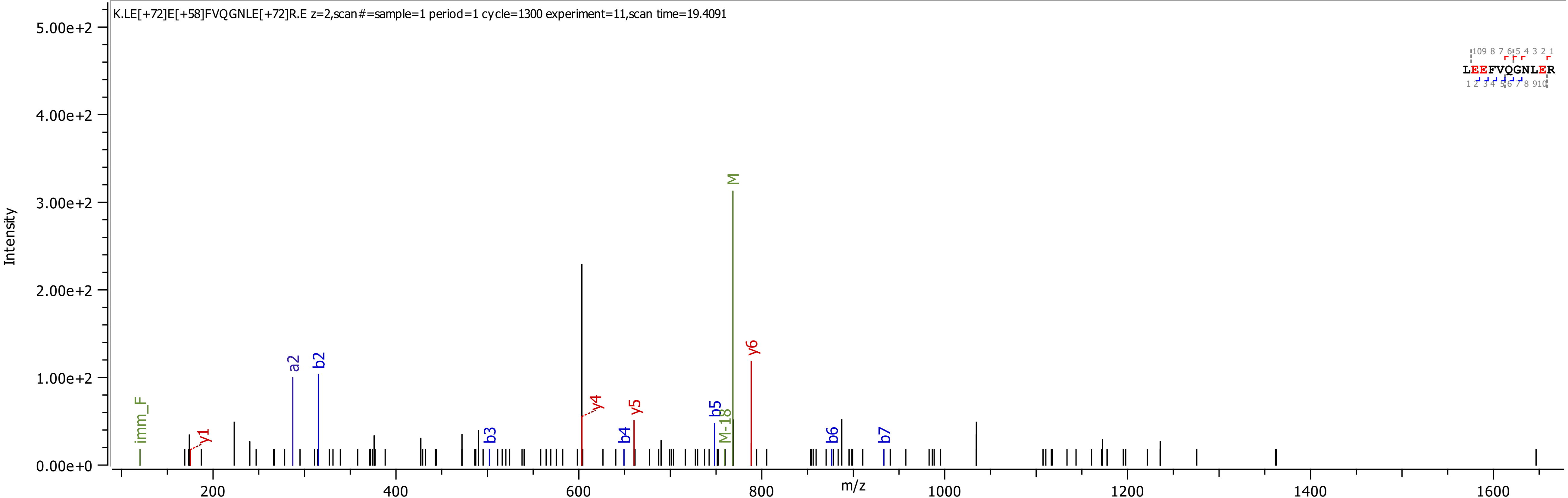
K.LE[+72]E[+58]FVQGNLE[+72]R.E z=2,scan#=sample=1 period=1 cycle=1300 experiment=11,scan time=19.4091
Intensity
m/z
imm_F
y1
a2
b2
b3
y4
b4
y5
b5
M-18
M
y6
b6
b7
LEEFVQGNLER

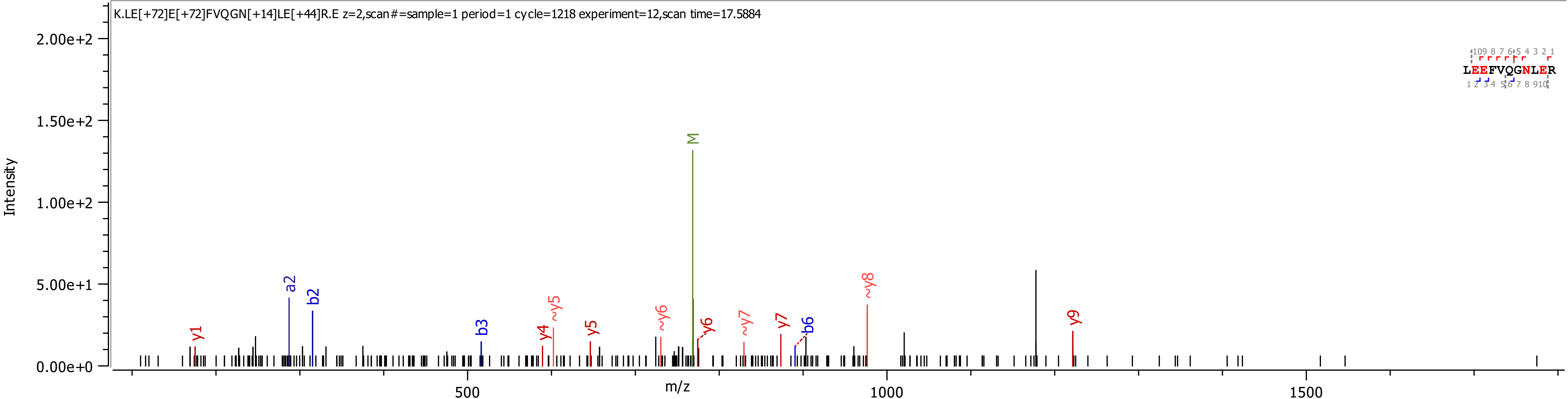
K.LE[+72]E[+72]FVQGN[+14]LE[+44]R.E z=2,scan#=sample=1 period=1 cycle=1218 experiment=12,scan time=17.5884
Intensity
m/z
M
y1
a2
b2
b3
y4
~y5
y5
~y6
y6
~y7
y7
b6
~y8
y9
LEEFVQGNLER

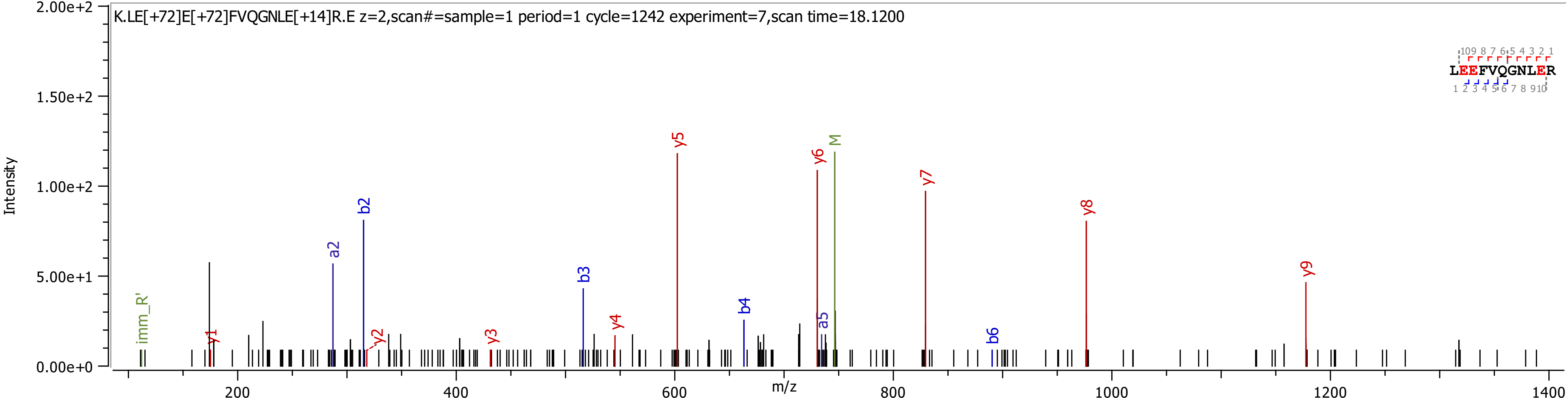
K.LE[+72]E[+72]FVQGNLE[+14]R.E z=2,scan#=sample=1 period=1 cycle=1242 experiment=7,scan time=18.1200
Intensity
m/z
LEEFVQGNLER
imm_R'
y1
a2
b2
y2
y3
b3
y4
y5
b4
y6
a5
M
y7
b6
y8
y9

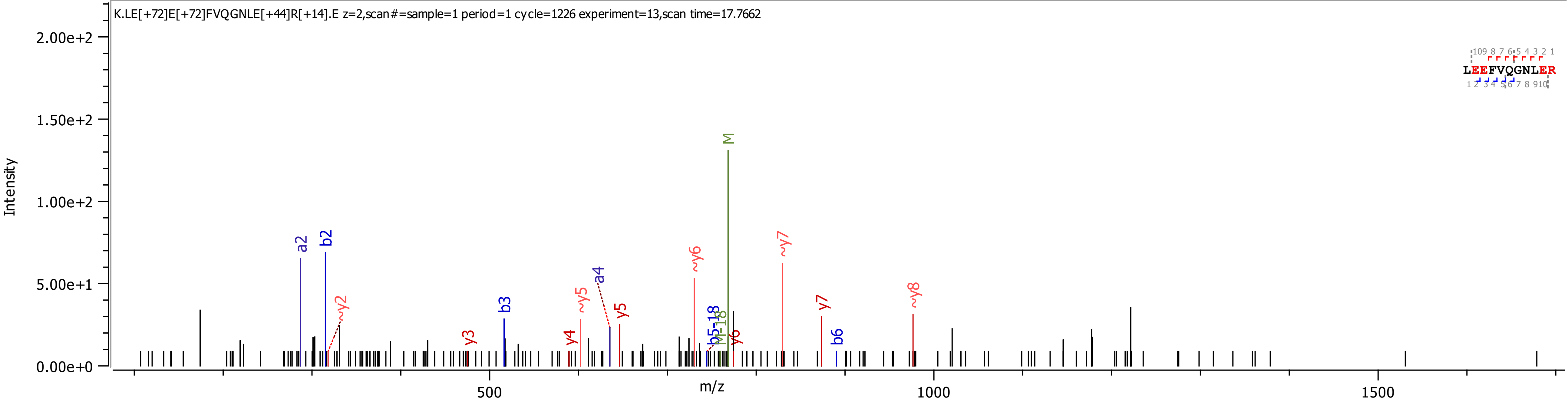
K.LE[+72]E[+72]FVQGNLE[+44]R[+14].E z=2,scan#=sample=1 period=1 cycle=1226 experiment=13,scan time=17.7662
Intensity
2.00e+2
1.50e+2
1.00e+2
5.00e+1
0.00e+0
m/z
500
1000
1500
a2
b2
~y2
y3
b3
y4
~y5
a4
y5
~y6
b5-18
M-18
y6
M
~y7
y7
b6
~y8
LEEFVQGNLER

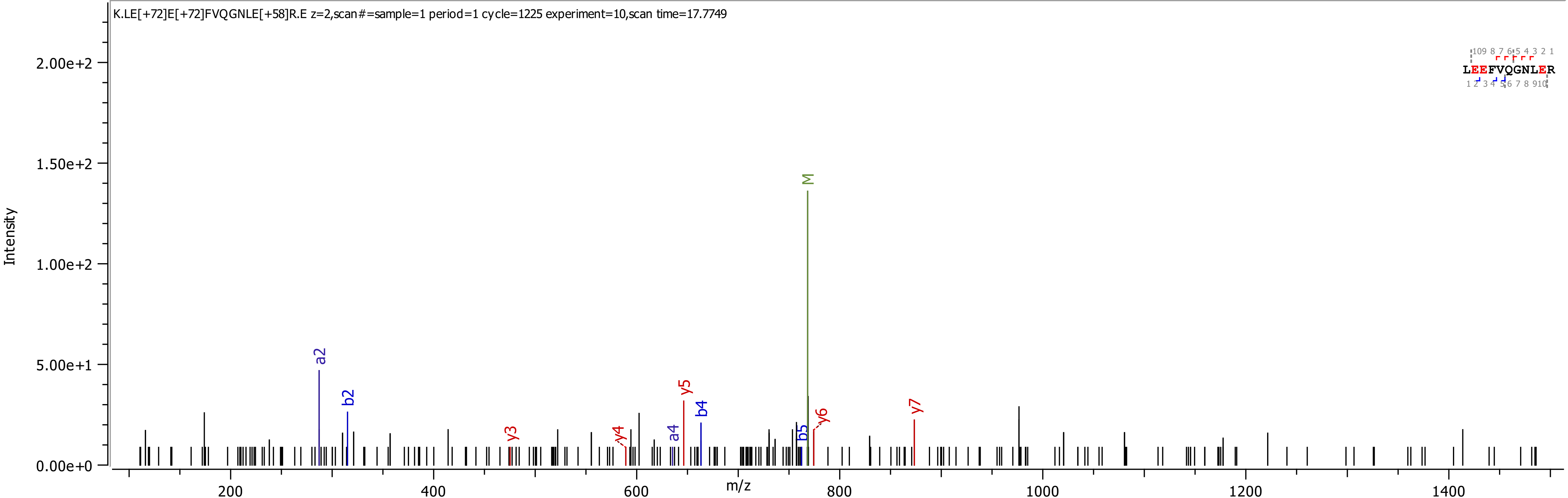

K.LE[+72]E[+72]FVQGNLE[+58]R.E z=2,scan#=sample=1 period=1 cycle=1225 experiment=10,scan time=17.7749
Intensity
m/z
LEEFVQGNLER
M
a2
b2
y3
y4
a4
y5
b4
b5
y6
y7
2.00e+2
1.50e+2
1.00e+2
5.00e+1
0.00e+0
200
400
600
800
1000
1200
1400

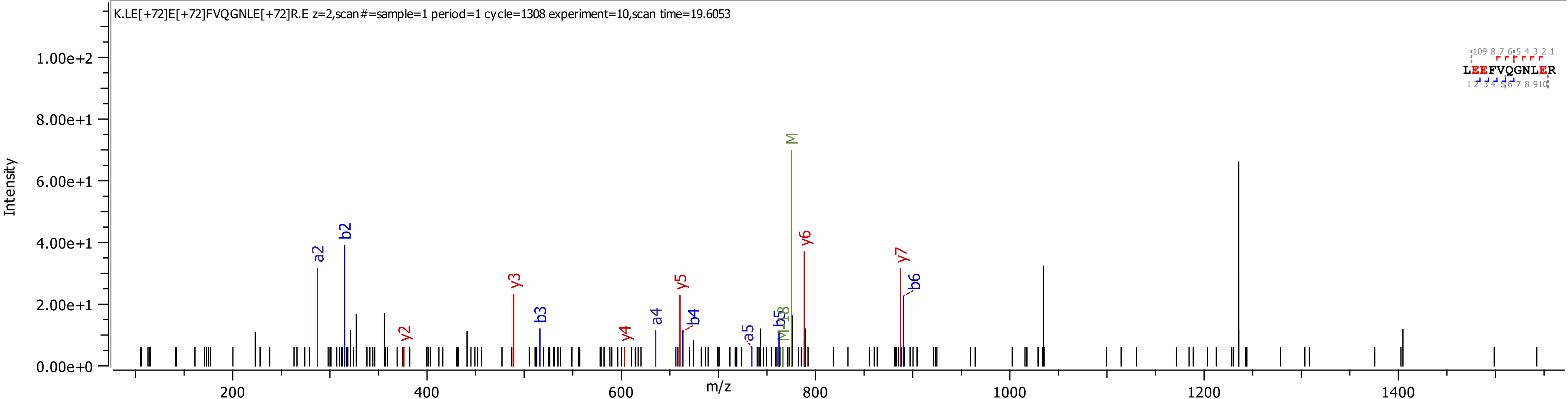

K.LE[+72]E[+72]FVQGNLE[+72]R.E z=2,scan#=sample=1 period=1 cycle=1308 experiment=10,scan time=19.6053
Intensity
m/z
LEEFVQGNLER
a2
b2
y2
y3
b3
y4
a4
y5
b4
a5
b5
M-18
M
y6
y7
b6

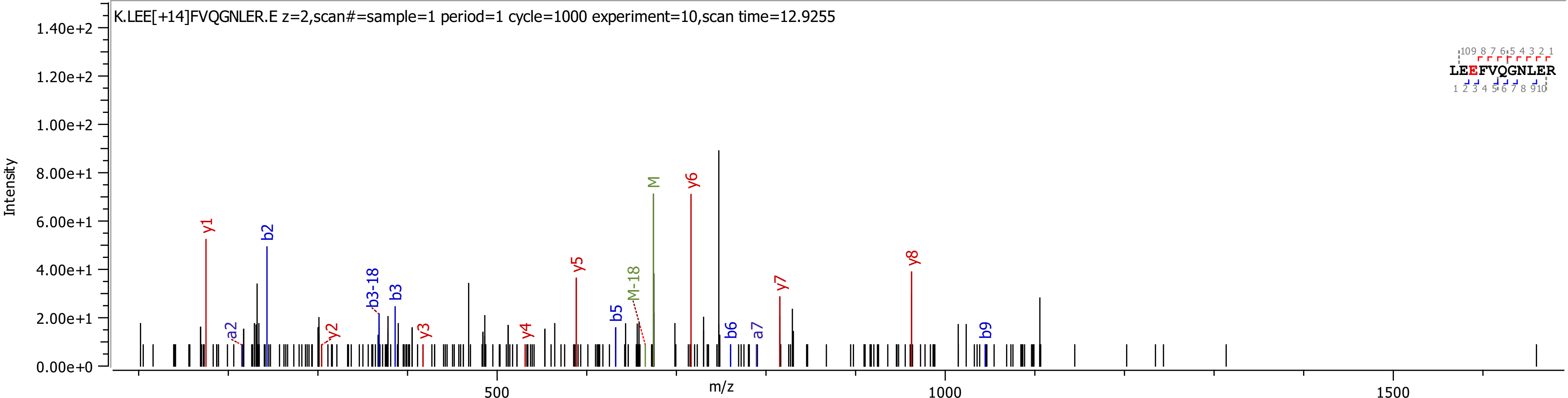

K.LEE[+14]FVQGNLER.E z=2,scan#=sample=1 period=1 cycle=1000 experiment=10,scan time=12.9255
Intensity
m/z
LEEFVQGNLER
y1
b2
a2
y2
b3-18
b3
y3
y4
y5
b5
M-18
M
y6
b6
a7
y7
y8
b9

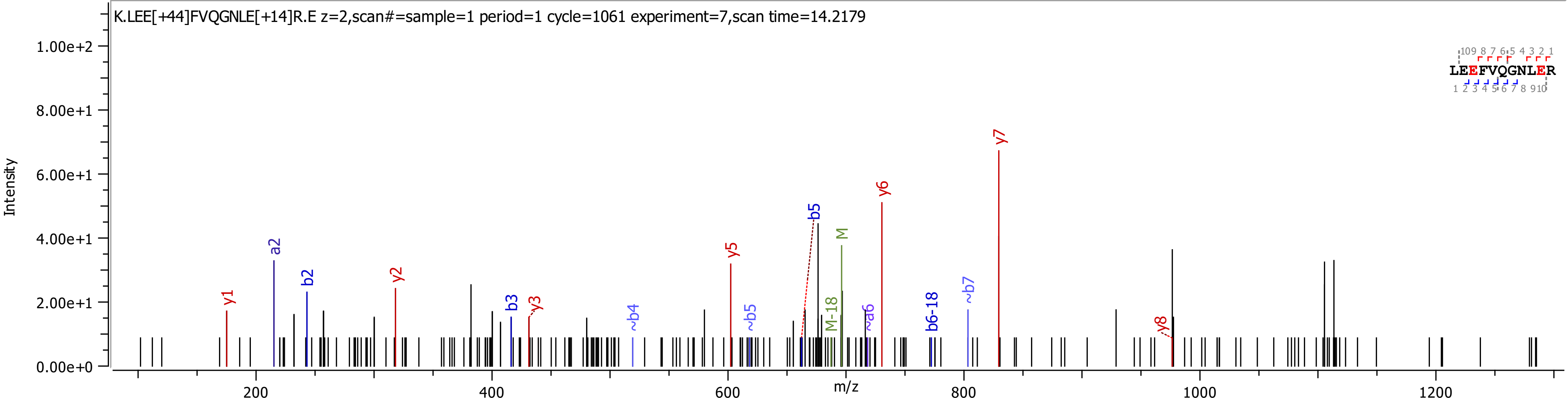
K.LEE[+44]FVQGNLE[+14]R.E z=2,scan#=sample=1 period=1 cycle=1061 experiment=7,scan time=14.2179
Intensity
m/z
y1
a2
b2
y2
b3
y3
~b4
y5
~b5
b5
M-18
M
~a6
y6
b6-18
~b7
y7
y8
LEEFVQGNLER

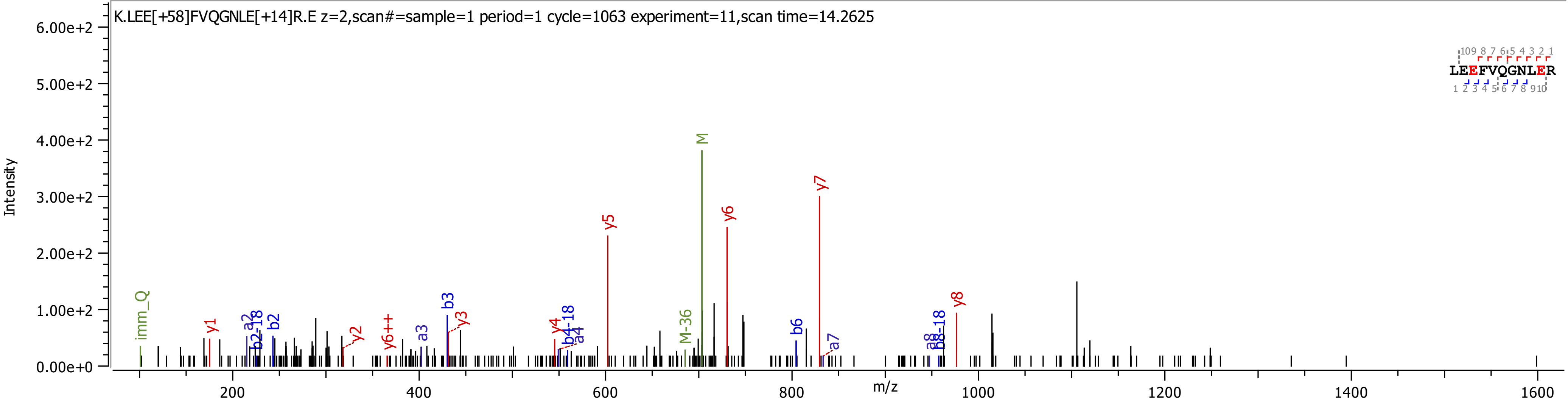

K.LEE[+58]FVQGNLE[+14]R.E z=2,scan#=sample=1 period=1 cycle=1063 experiment=11,scan time=14.2625
LEEFVQGNLER
Intensity
m/z
6.00e+2
5.00e+2
4.00e+2
3.00e+2
2.00e+2
1.00e+2
0.00e+0
200
400
600
800
1000
1200
1400
1600
imm_Q
y1
a2
b2-18
b2
y2
y6++
a3
b3
y3
y4
b4-18
a4
y5
M-36
M
y6
b6
y7
a7
a8
b8-18
y8

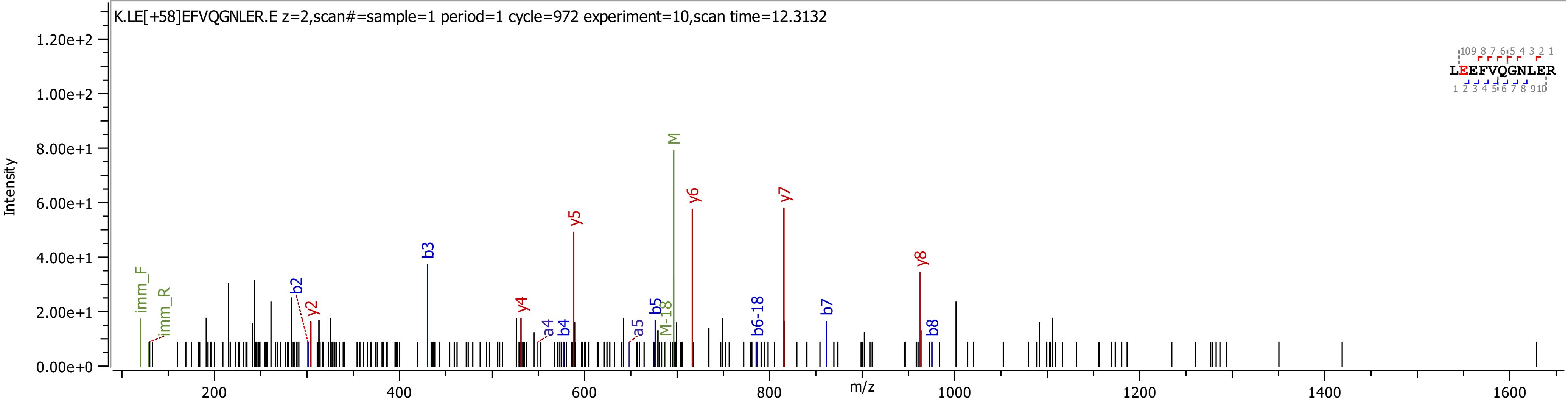
K.LE[+58]EFVQGNLER.E z=2,scan#=sample=1 period=1 cycle=972 experiment=10,scan time=12.3132
Intensity
m/z
LEEFVQGNLER
imm_F
imm_R
b2
y2
b3
y4
a4
b4
y5
a5
b5
M-18
M
y6
b6-18
y7
b7
y8
b8

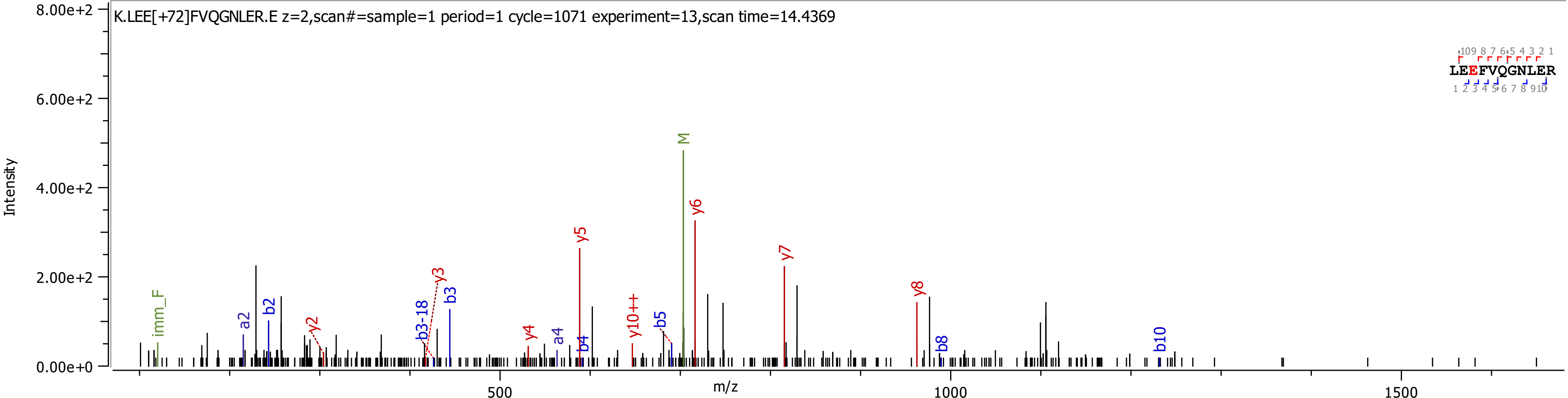
K.LEE[+72]FVQGNLER.E z=2,scan#=sample=1 period=1 cycle=1071 experiment=13,scan time=14.4369
Intensity
m/z
LEEFVQGNLER
imm_F
a2
b2
y2
b3-18
y3
b3
y4
a4
y5
b4
y10++
b5
M
y6
y7
y8
b8
b10
8.00e+2
6.00e+2
4.00e+2
2.00e+2
0.00e+0
500
1000
1500

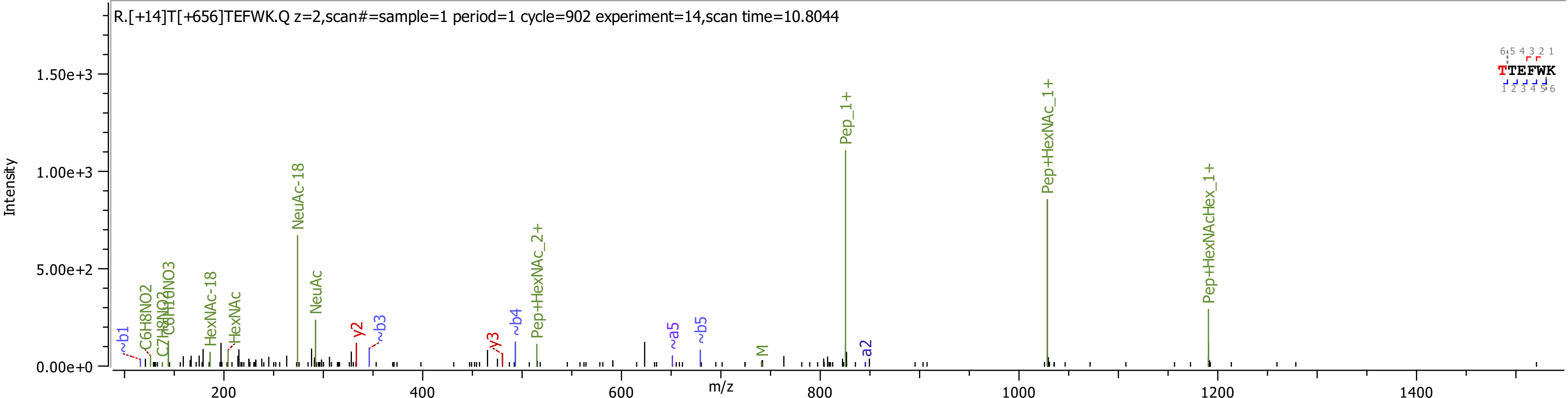
R.[+14]T[+656]TEFWK.Q z=2,scan#=sample=1 period=1 cycle=902 experiment=14,scan time=10.8044
TTEFWK
Intensity
m/z
NeuAc-18
NeuAc
HexNAc-18
HexNAc
C6H8NO2
C7H8NO3
C8H10NO3
~b1
y2
~b3
y3
~b4
Pep+HexNAc_2+
~a5
~b5
M
a2
Pep_1+
Pep+HexNAc_1+
Pep+HexNAcHex_1+

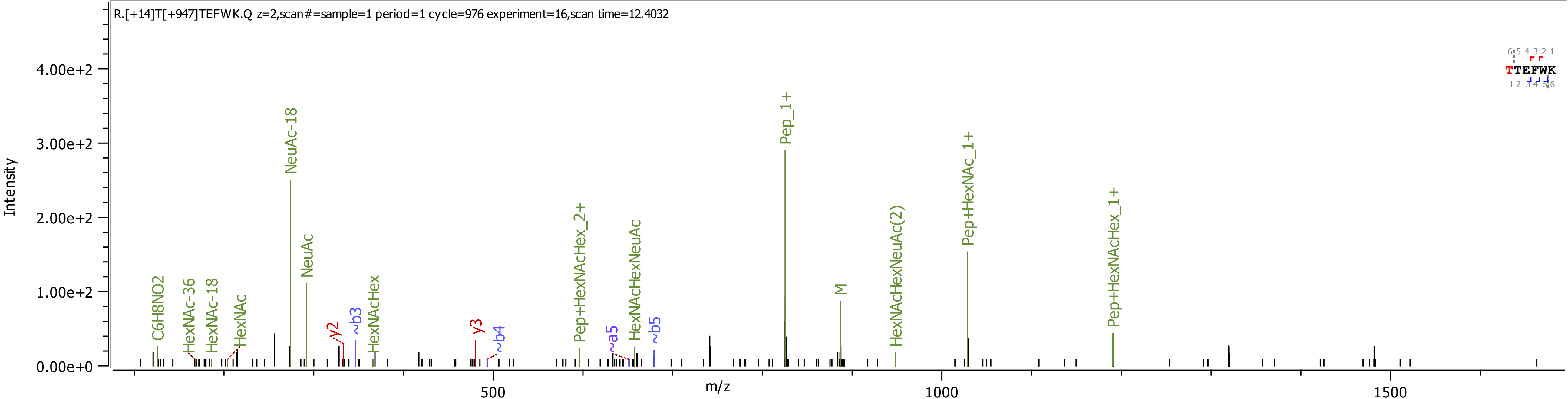

R.[+14]T[+947]TEFWK.Q z=2,scan#=sample=1 period=1 cycle=976 experiment=16,scan time=12.4032
Intensity
4.00e+2
3.00e+2
2.00e+2
1.00e+2
0.00e+0
m/z
500
1000
1500
C6H8NO2
HexNAc-36
HexNAc-18
HexNAc
NeuAc-18
NeuAc
y2
~b3
HexNAcHex
y3
~b4
Pep+HexNAcHex_2+
~a5
HexNAcHexNeuAc
~b5
Pep_1+
M
HexNAcHexNeuAc(2)
Pep+HexNAc_1+
Pep+HexNAcHex_1+
TTEFWK

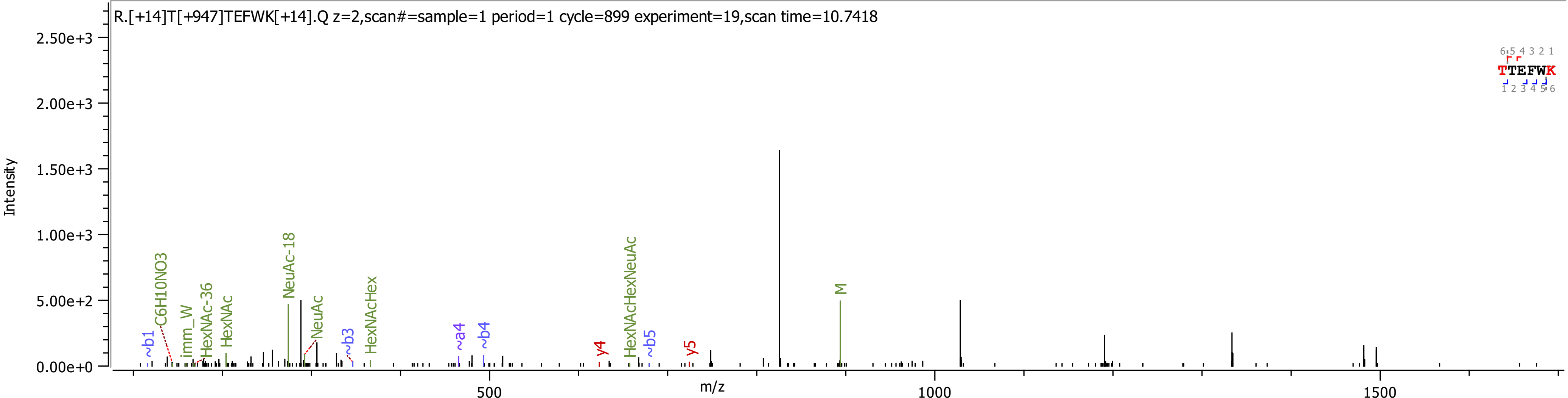

R.[+14]T[+947]TEFWK[+14].Q z=2,scan#=sample=1 period=1 cycle=899 experiment=19,scan time=10.7418
Intensity
m/z
2.50e+3
2.00e+3
1.50e+3
1.00e+3
5.00e+2
0.00e+0
500
1000
1500
~b1
C6H10NO3
imm_W
HexNAc-36
HexNAc
NeuAc-18
NeuAc
~b3
HexNAcHex
~a4
~b4
y4
HexNAcHexNeuAc
~b5
y5
M
TTEFWK

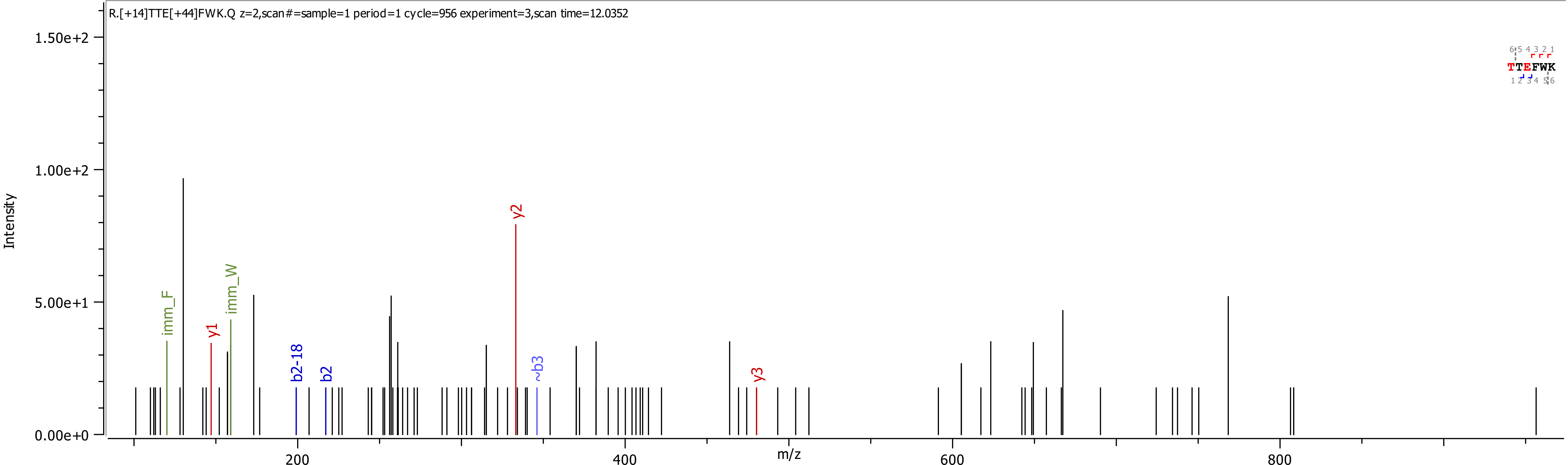
R.[+14]TTE[+44]FWK.Q z=2,scan#=sample=1 period=1 cycle=956 experiment=3,scan time=12.0352
Intensity
1.50e+2
1.00e+2
5.00e+1
0.00e+0
m/z
200
400
600
800
imm_F
y1
imm_W
b2-18
b2
y2
~b3
y3
6 5 4 3 2 1
TTEFWK
1 2 3 4 5 6

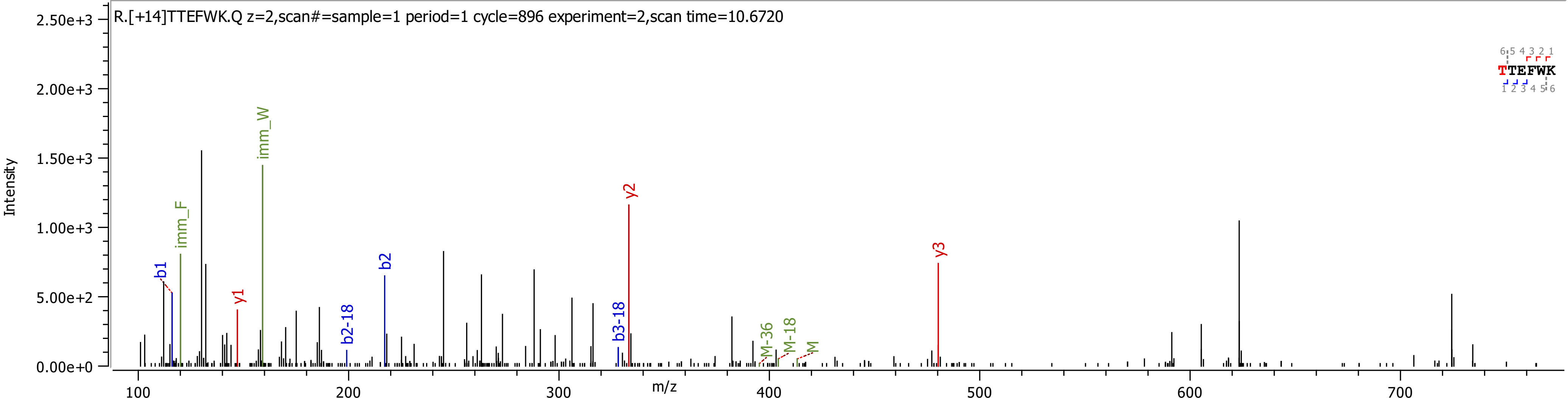

R.[+14]TTEFWK.Q z=2,scan#=sample=1 period=1 cycle=896 experiment=2,scan time=10.6720
Intensity
m/z
b1
imm_F
y1
imm_W
b2-18
b2
b3-18
y2
M-36
M-18
M
y3
TTEFWK

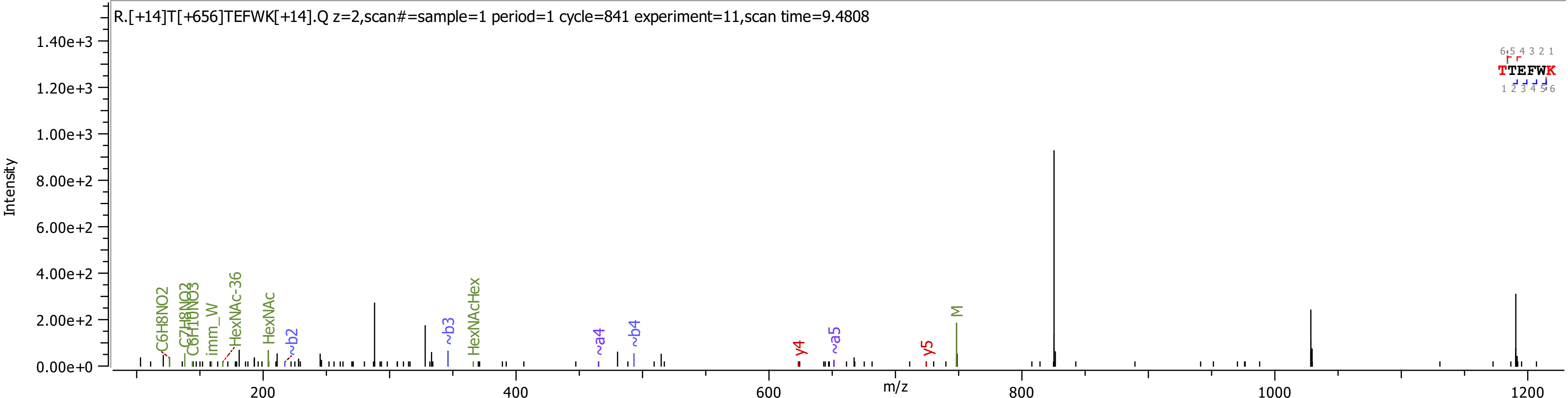

R.[+14]T[+656]TEFWK[+14].Q z=2,scan#=sample=1 period=1 cycle=841 experiment=11,scan time=9.4808
Intensity
1.40e+3
1.20e+3
1.00e+3
8.00e+2
6.00e+2
4.00e+2
2.00e+2
0.00e+0
m/z
200
400
600
800
1000
1200
C6H8NO2
C7H8NO2
C6H10NO3
imm_W
HexNAc-36
HexNAc
~b2
~b3
HexNAcHex
~a4
~b4
y4
~a5
y5
M
TTEFWK

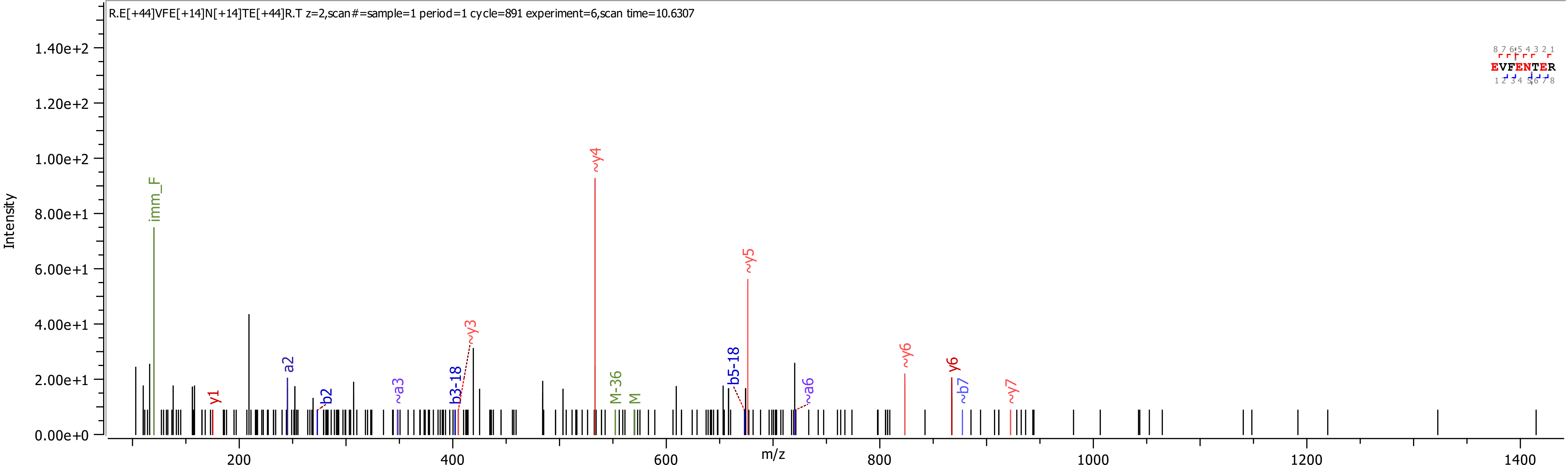
R.E[+44]VFE[+14]N[+14]TE[+44]R.T z=2,scan#=sample=1 period=1 cycle=891 experiment=6,scan time=10.6307
Intensity
m/z
1.40e+2
1.20e+2
1.00e+2
8.00e+1
6.00e+1
4.00e+1
2.00e+1
0.00e+0
200
400
600
800
1000
1200
1400
imm_F
y1
a2
b2
~a3
b3-18
~y3
~y4
M-36
M
b5-18
~y5
~a6
~y6
y6
~b7
~y7
EVFENTER

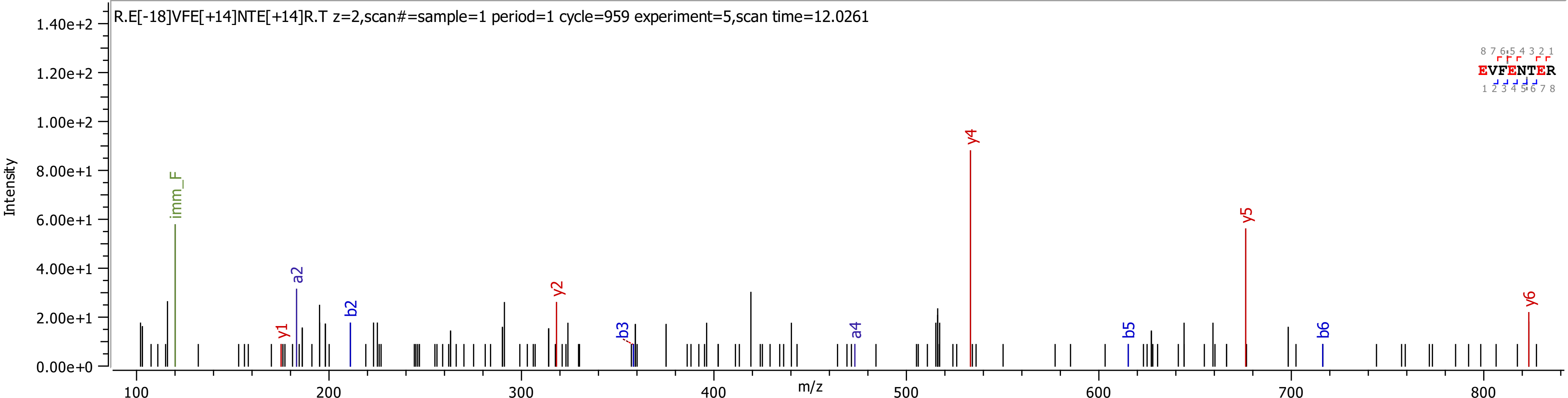
R.E[-18]VFE[+14]NTE[+14]R.T z=2,scan#=sample=1 period=1 cycle=959 experiment=5,scan time=12.0261
Intensity
m/z
imm_F
y1
a2
b2
y2
b3
a4
y4
b5
y5
b6
y6
EVFENTER
1.40e+2
1.20e+2
1.00e+2
8.00e+1
6.00e+1
4.00e+1
2.00e+1
0.00e+0
100
200
300
400
500
600
700
800

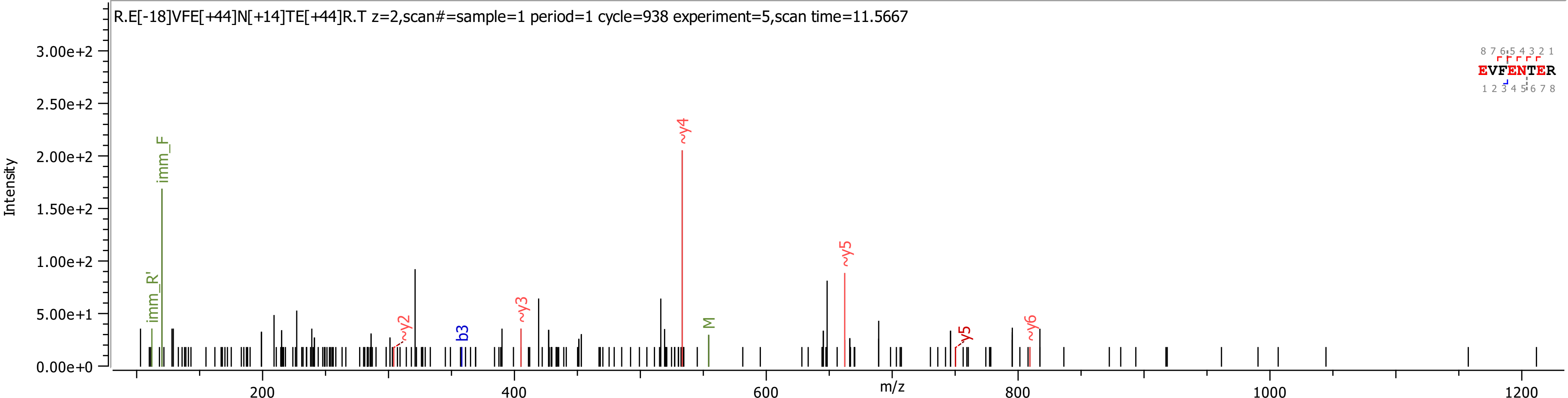
R.E[-18]VFE[+44]N[+14]TE[+44]R.T z=2,scan#=sample=1 period=1 cycle=938 experiment=5,scan time=11.5667
Intensity
m/z
imm_R'
imm_F
~y2
b3
~y3
~y4
M
~y5
y5
~y6
EVFENTER
0.00e+0
5.00e+1
1.00e+2
1.50e+2
2.00e+2
2.50e+2
3.00e+2
200
400
600
800
1000
1200

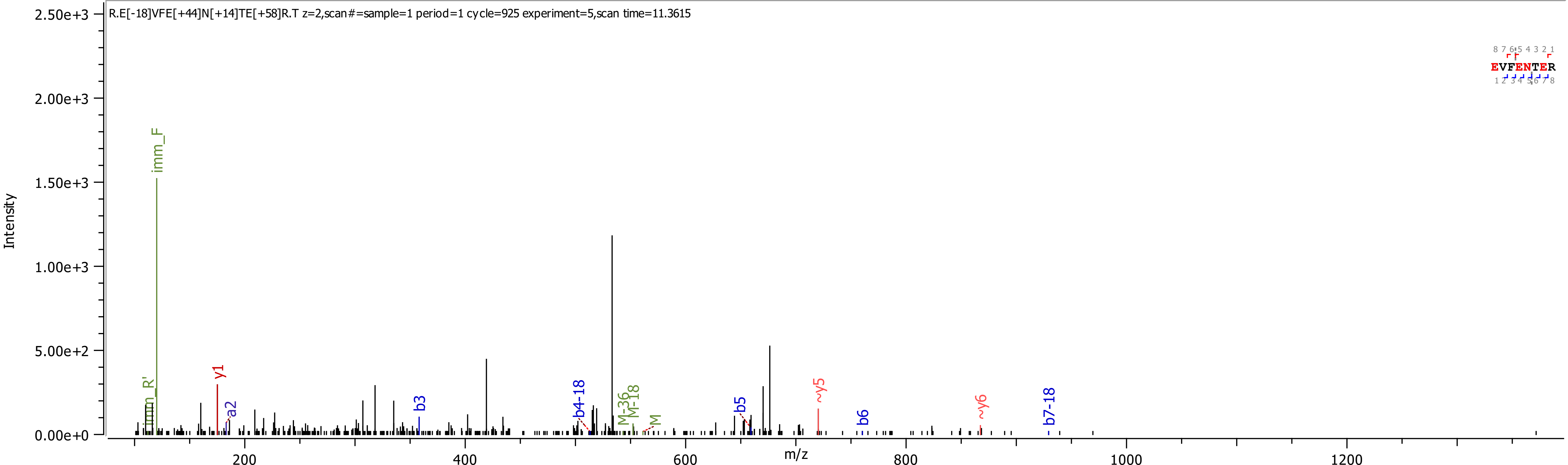
R.E[-18]VFE[+44]N[+14]TE[+58]R.T z=2,scan#=sample=1 period=1 cycle=925 experiment=5,scan time=11.3615
Intensity
2.50e+3
2.00e+3
1.50e+3
1.00e+3
5.00e+2
0.00e+0
m/z
200
400
600
800
1000
1200
imm_F
imm_R'
y1
a2
b3
b4-18
M-36
M-18
M
b5
~y5
b6
~y6
b7-18
EVFENTER
8 7 6 5 4 3 2 1
1 2 3 4 5 6 7 8

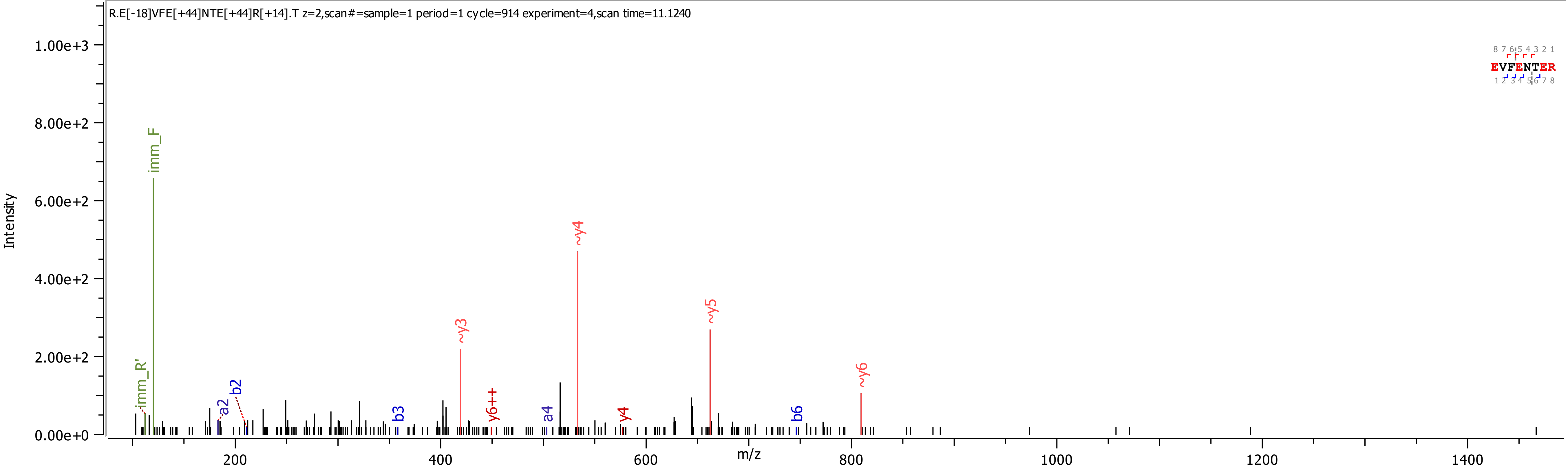

R.E[-18]VFE[+44]NTE[+44]R[+14].T z=2,scan#=sample=1 period=1 cycle=914 experiment=4,scan time=11.1240
EVFENTER
Intensity
m/z
imm_F
imm_R'
a2
b2
b3
~y3
y6++
a4
~y4
y4
~y5
b6
~y6

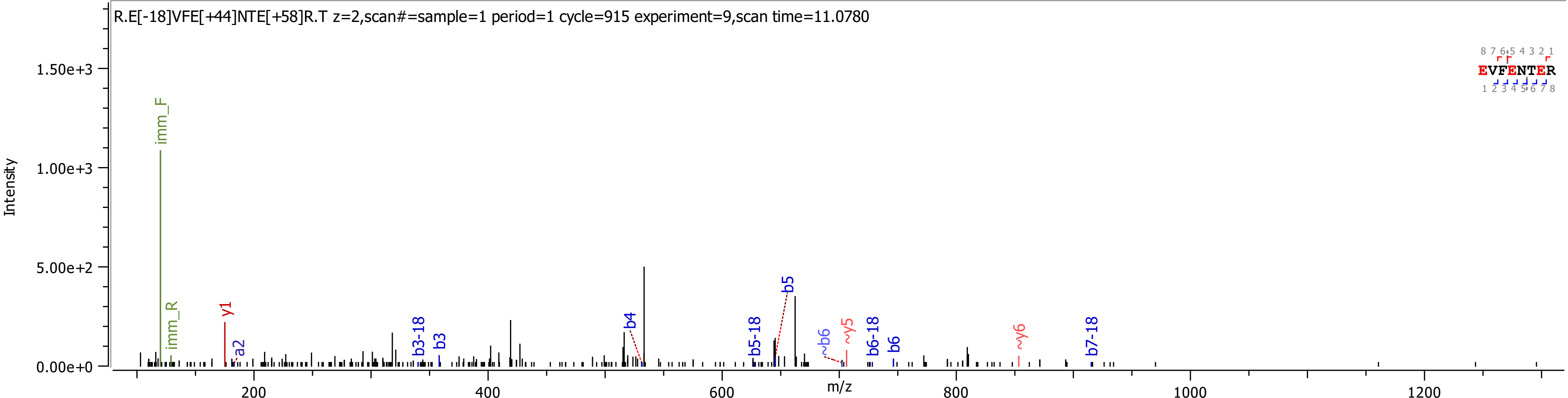
R.E[-18]VFE[+44]NTE[+58]R.T z=2,scan#=sample=1 period=1 cycle=915 experiment=9,scan time=11.0780
Intensity
1.50e+3
1.00e+3
5.00e+2
0.00e+0
m/z
200
400
600
800
1000
1200
imm_F
imm_R
y1
a2
b3-18
b3
b4
b5-18
b5
~b6
~y5
b6-18
b6
~y6
b7-18
8 7 6 5 4 3 2 1
EVFENTER
1 2 3 4 5 6 7 8

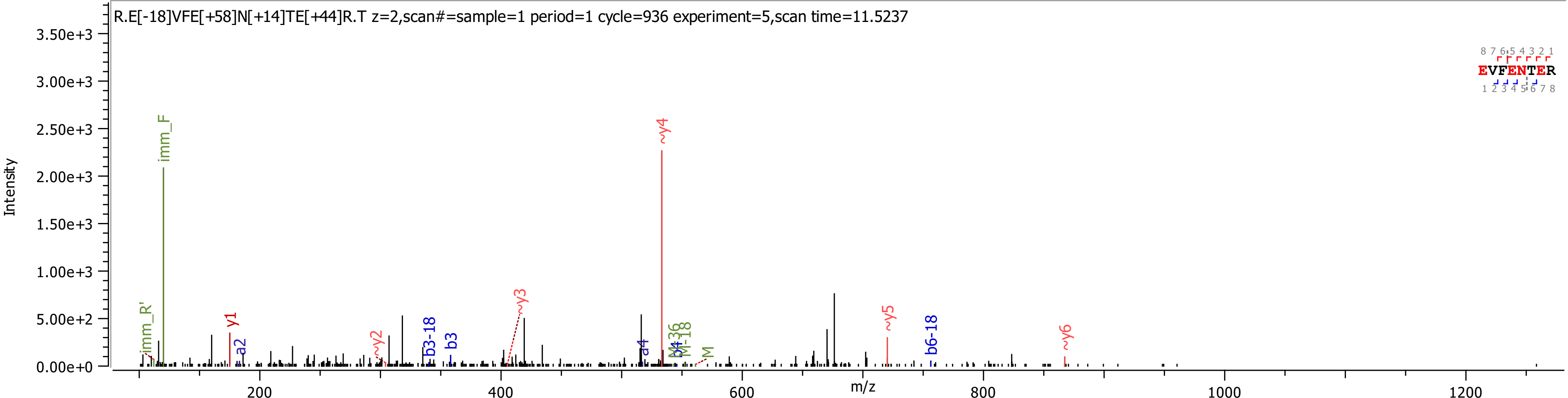

R.E[-18]VFE[+58]N[+14]TE[+44]R.T z=2,scan#=sample=1 period=1 cycle=936 experiment=5,scan time=11.5237
Intensity
m/z
EVFENTER
imm_F
imm_R'
y1
a2
~y2
b3-18
b3
~y3
a4
~y4
b4
M-36
M-18
M
~y5
b6-18
~y6

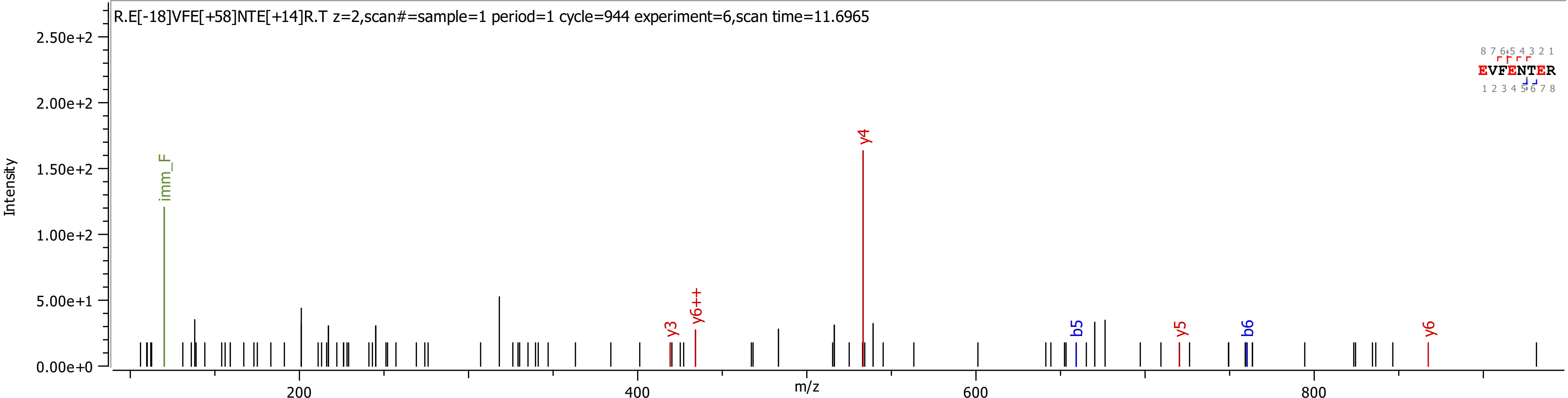

R.E[-18]VFE[+58]NTE[+14]R.T z=2,scan#=sample=1 period=1 cycle=944 experiment=6,scan time=11.6965
Intensity
2.50e+2
2.00e+2
1.50e+2
1.00e+2
5.00e+1
0.00e+0
imm_F
y3
y6++
y4
b5
y5
b6
y6
EVFENTER
8 7 6 5 4 3 2 1
1 2 3 4 5 6 7 8
200
400
600
800
m/z

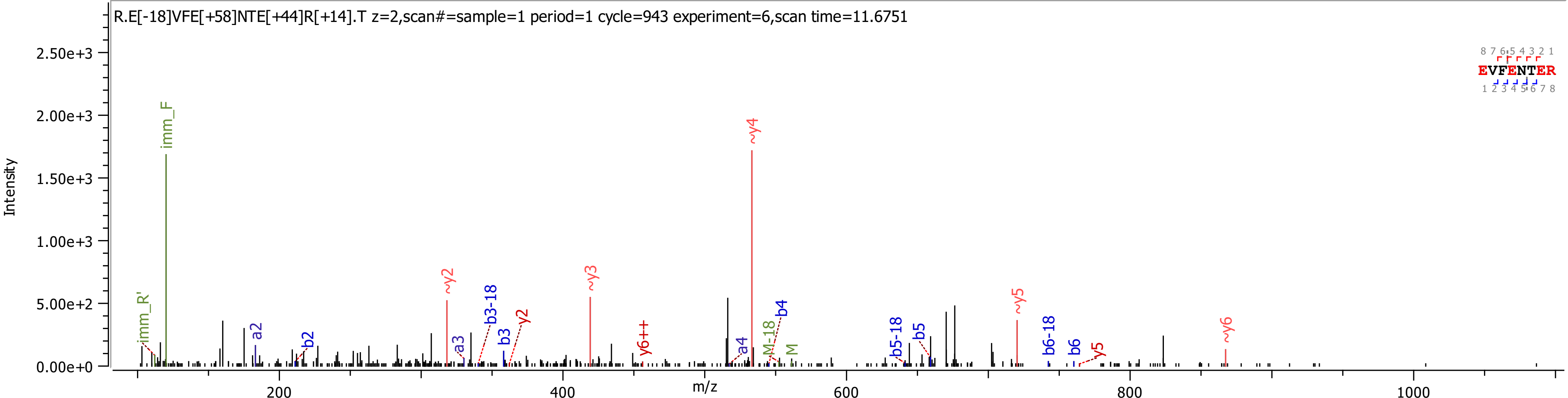

R.E[-18]VFE[+58]NTE[+44]R[+14].T z=2,scan#=sample=1 period=1 cycle=943 experiment=6,scan time=11.6751
Intensity
m/z
EVFENTER
imm_R'
imm_F
a2
b2
~y2
a3
b3-18
b3
y2
~y3
y6++
a4
M-18
b4
~y4
M
b5-18
b5
~y5
b6-18
b6
y5
~y6

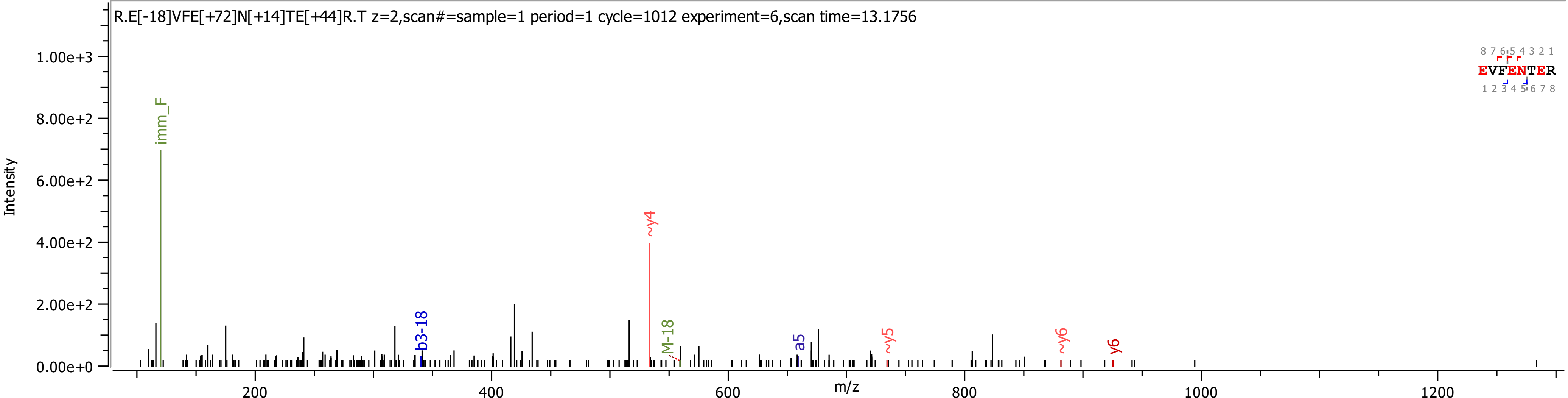
R.E[-18]VFE[+72]N[+14]TE[+44]R.T z=2,scan#=sample=1 period=1 cycle=1012 experiment=6,scan time=13.1756
Intensity
m/z
imm_F
b3-18
~y4
M-18
a5
~y5
~y6
y6
EVFENTER

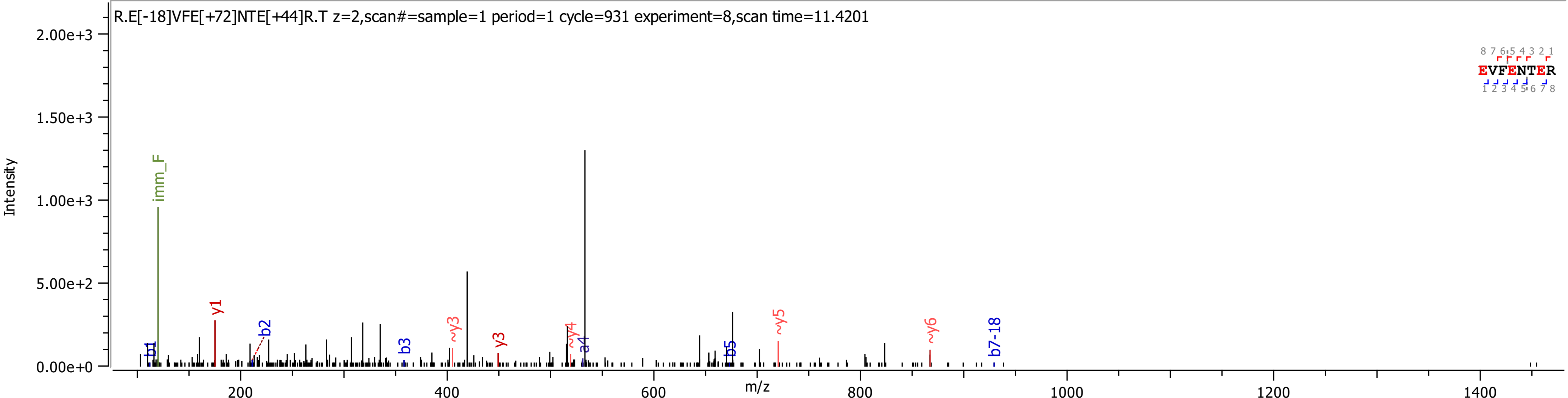

R.E[-18]VFE[+72]NTE[+44]R.T z=2,scan#=sample=1 period=1 cycle=931 experiment=8,scan time=11.4201
Intensity
m/z
imm_F
y1
b1
b2
b3
~y3
y3
~y4
a4
b5
~y5
~y6
b7-18
EVFENTER

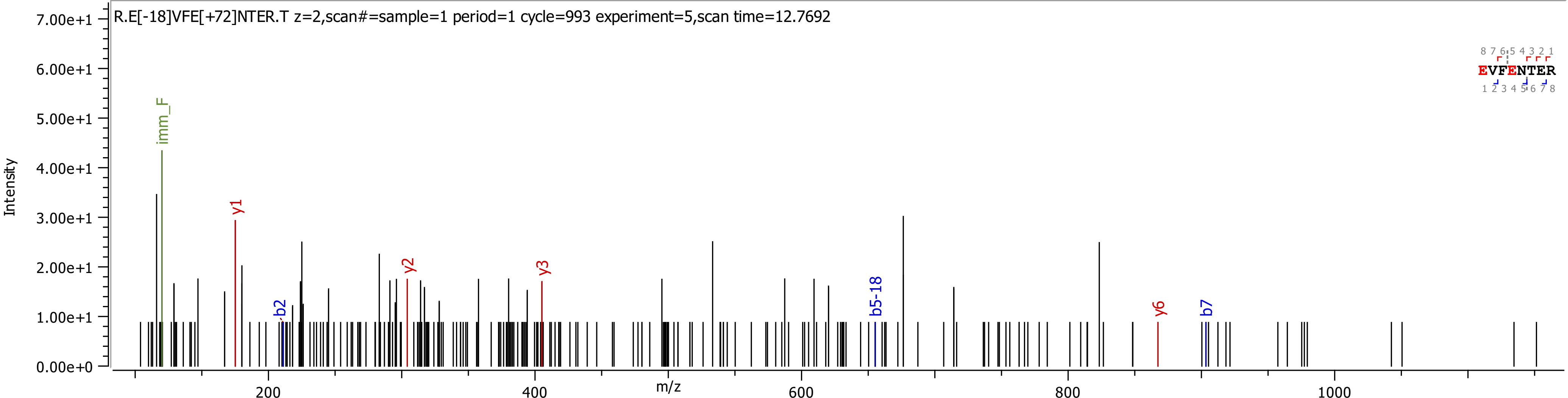

R.E[-18]VFE[+72]NTER.T z=2,scan#=sample=1 period=1 cycle=993 experiment=5,scan time=12.7692
Intensity
m/z
imm_F
y1
y2
y3
y6
b2
b5-18
b7
EVFENTER

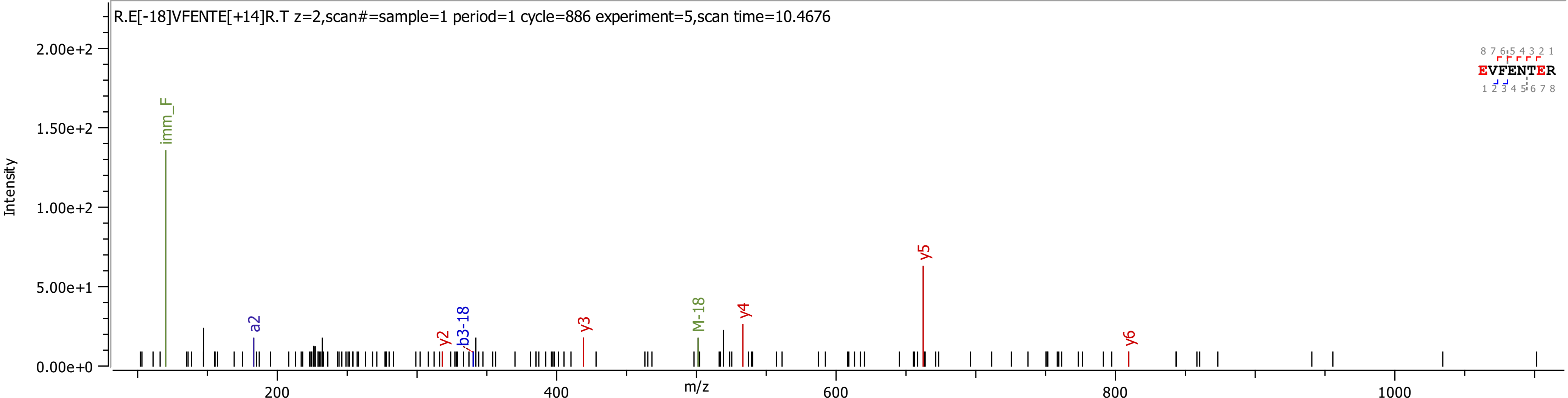
R.E[-18]VFENTE[+14]R.T z=2,scan#=sample=1 period=1 cycle=886 experiment=5,scan time=10.4676
Intensity
m/z
imm_F
a2
y2
b3-18
y3
M-18
y4
y5
y6
EVFENTER
2.00e+2
1.50e+2
1.00e+2
5.00e+1
0.00e+0
200
400
600
800
1000

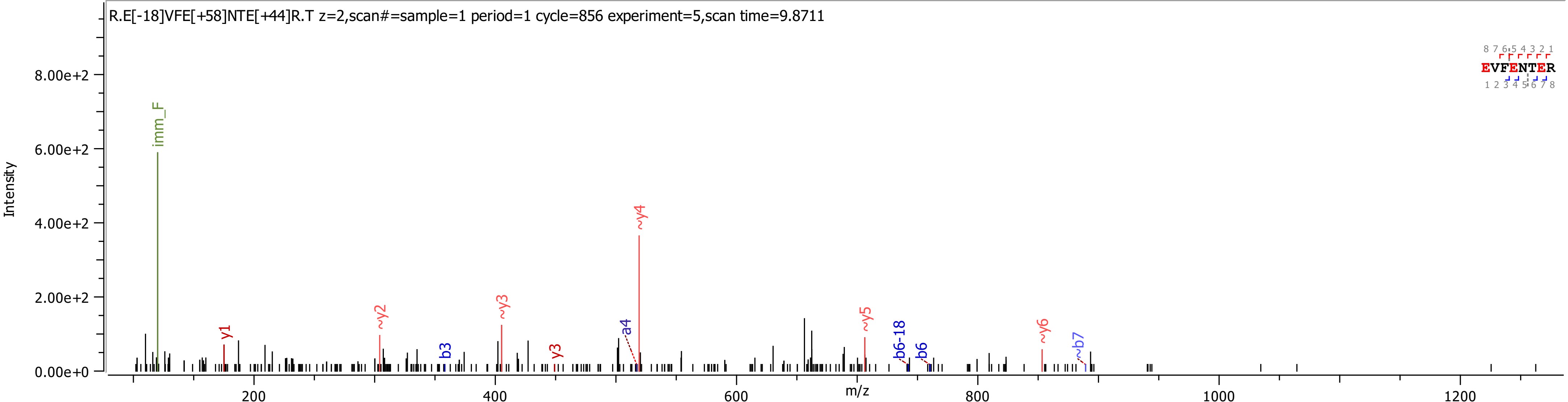
R.E[-18]VFE[+58]NTE[+44]R.T z=2,scan#=sample=1 period=1 cycle=856 experiment=5,scan time=9.8711
Intensity
8.00e+2
6.00e+2
4.00e+2
2.00e+2
0.00e+0
m/z
200
400
600
800
1000
1200
imm_F
y1
~y2
b3
~y3
y3
a4
~y4
~y5
b6-18
b6
~y6
~b7
EVFENTER
8 7 6 5 4 3 2 1
1 2 3 4 5 6 7 8

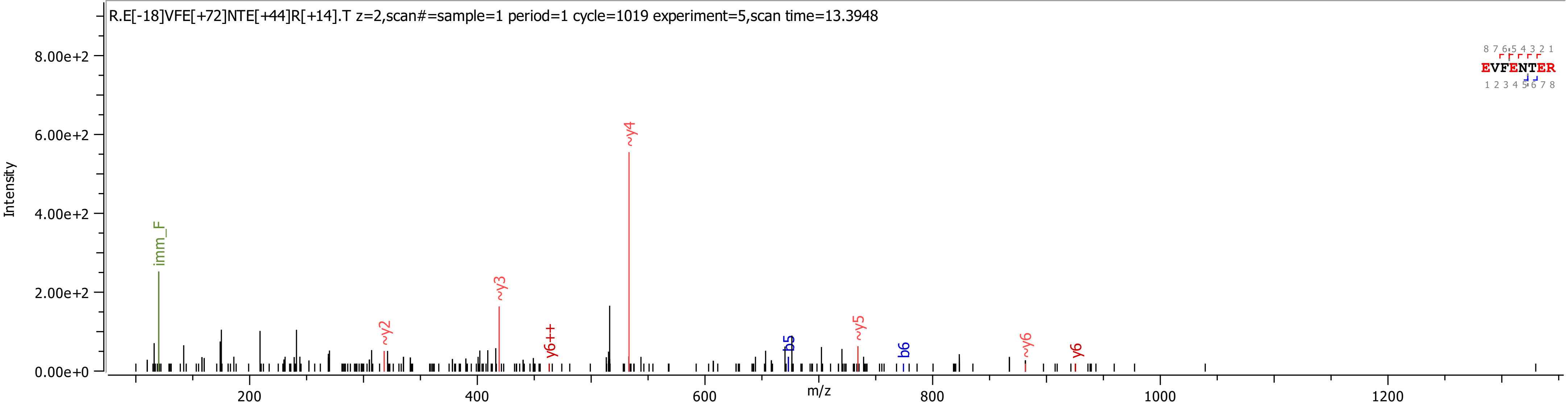

R.E[-18]VFE[+72]NTE[+44]R[+14].T z=2,scan#=sample=1 period=1 cycle=1019 experiment=5,scan time=13.3948
Intensity
m/z
8.00e+2
6.00e+2
4.00e+2
2.00e+2
0.00e+0
200
400
600
800
1000
1200
imm_F
~y2
~y3
y6++
~y4
b5
~y5
b6
~y6
y6
EVFENTER
8 7 6 5 4 3 2 1
1 2 3 4 5 6 7 8

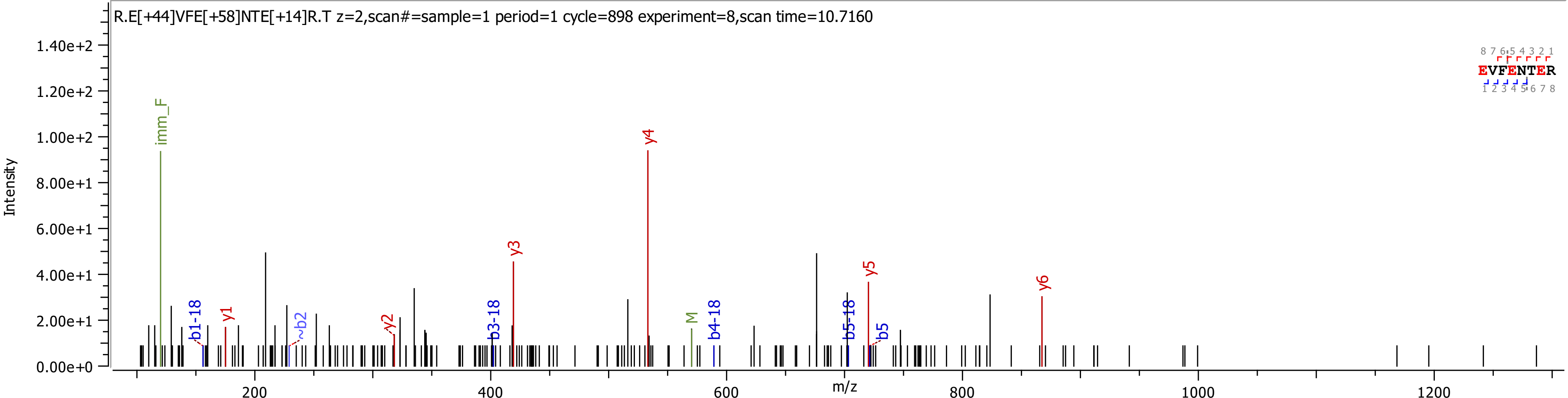
R.E[+44]VFE[+58]NTE[+14]R.T z=2,scan#=sample=1 period=1 cycle=898 experiment=8,scan time=10.7160
Intensity
m/z
1.40e+2
1.20e+2
1.00e+2
8.00e+1
6.00e+1
4.00e+1
2.00e+1
0.00e+0
200
400
600
800
1000
1200
imm_F
b1-18
y1
~b2
y2
b3-18
y3
y4
M
b4-18
b5-18
y5
b5
y6
EVFENTER
8 7 6 5 4 3 2 1
1 2 3 4 5 6 7 8

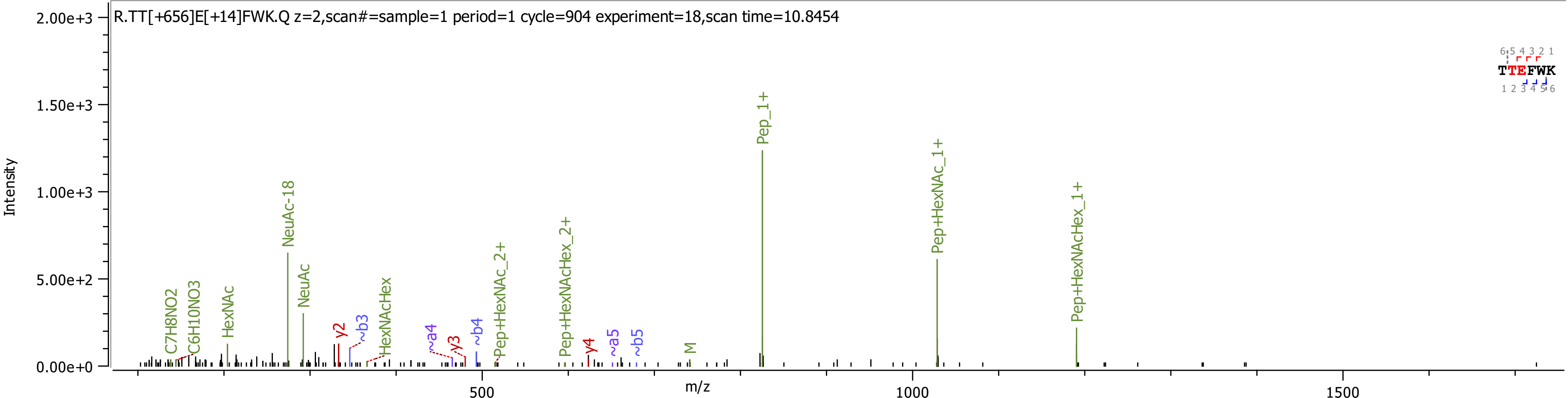

R.TT[+656]E[+14]FWK.Q z=2,scan#=sample=1 period=1 cycle=904 experiment=18,scan time=10.8454
Intensity
m/z
2.00e+3
1.50e+3
1.00e+3
5.00e+2
0.00e+0
500
1000
1500
C7H8NO2
C6H10NO3
HexNAc
NeuAc-18
NeuAc
y2
~b3
HexNAcHex
~a4
y3
~b4
Pep+HexNAc_2+
Pep+HexNAcHex_2+
y4
~a5
~b5
M
Pep_1+
Pep+HexNAc_1+
Pep+HexNAcHex_1+
TTEFWK

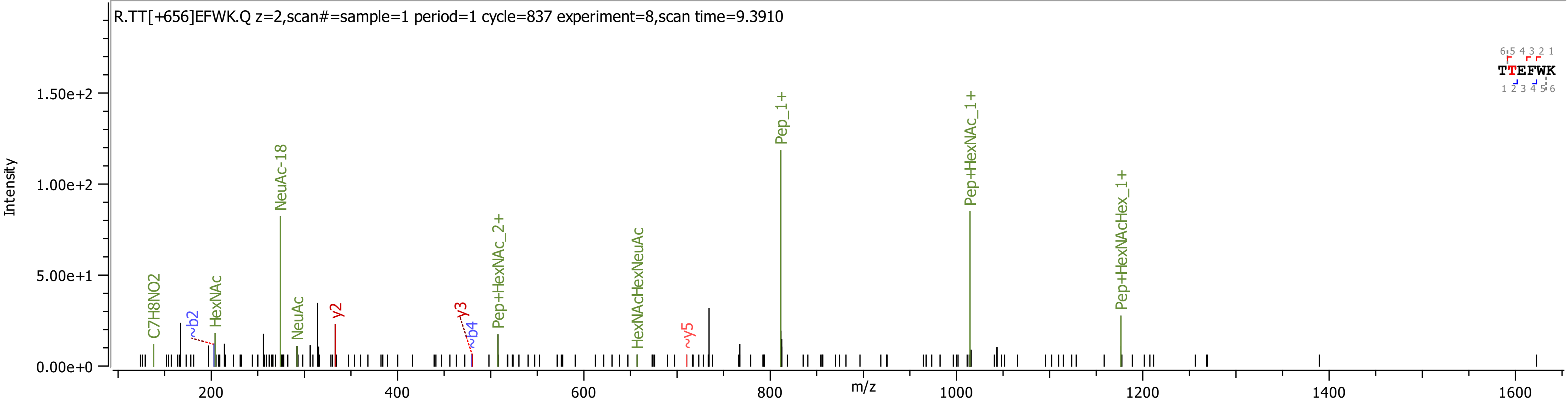

R.TT[+656]EFWK.Q z=2,scan#=sample=1 period=1 cycle=837 experiment=8,scan time=9.3910
Intensity
m/z
C7H8NO2
~b2
HexNAc
NeuAc-18
NeuAc
y2
y3
~b4
Pep+HexNAc_2+
HexNAcHexNeuAc
~y5
Pep_1+
Pep+HexNAc_1+
Pep+HexNAcHex_1+
TTEFWK

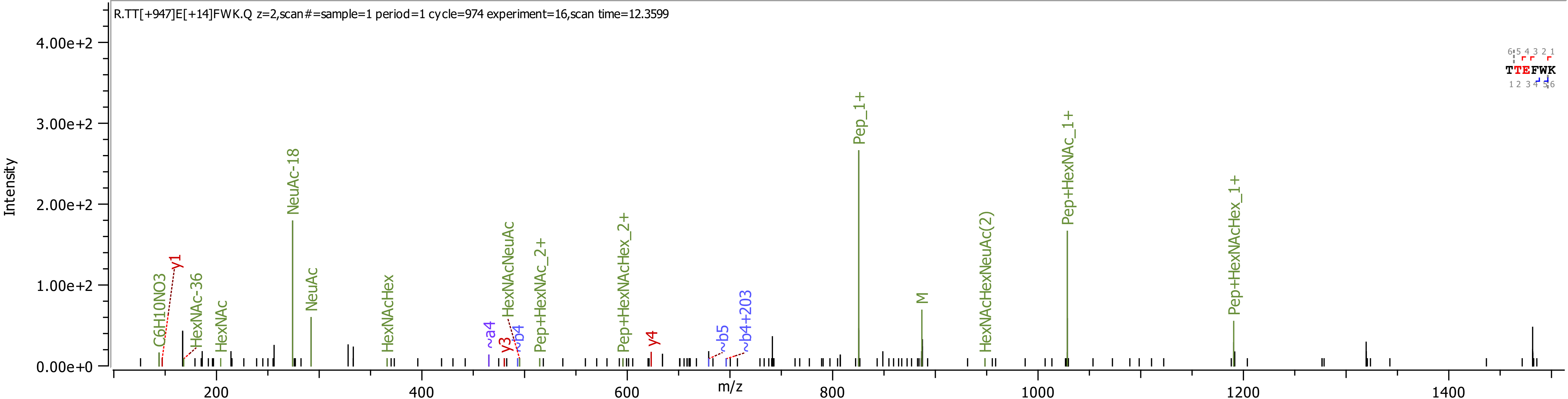
R.TT[+947]E[+14]FWK.Q z=2,scan#=sample=1 period=1 cycle=974 experiment=16,scan time=12.3599
Intensity
4.00e+2
3.00e+2
2.00e+2
1.00e+2
0.00e+0
m/z
200
400
600
800
1000
1200
1400
C6H10NO3
y1
HexNAc-36
HexNAc
NeuAc-18
NeuAc
HexNAcHex
~a4
y3
~b4
HexNAcNeuAc
Pep+HexNAc_2+
Pep+HexNAcHex_2+
y4
~b5
~b4+203
Pep_1+
M
HexNAcHexNeuAc(2)
Pep+HexNAc_1+
Pep+HexNAcHex_1+
TTEFWK

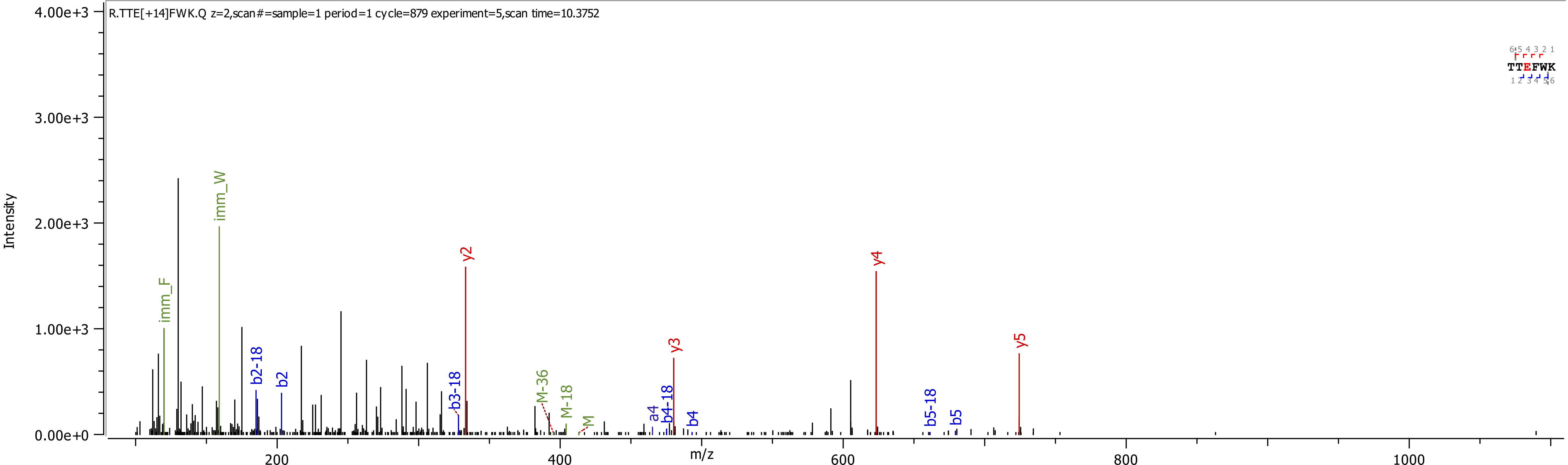

R.TTE[+14]FWK.Q z=2,scan#=sample=1 period=1 cycle=879 experiment=5,scan time=10.3752
Intensity
4.00e+3
3.00e+3
2.00e+3
1.00e+3
0.00e+0
m/z
200
400
600
800
1000
imm_F
imm_W
b2-18
b2
y2
b3-18
M-36
M-18
M
a4
b4-18
y3
b4
y4
b5-18
b5
y5
TTEFWK

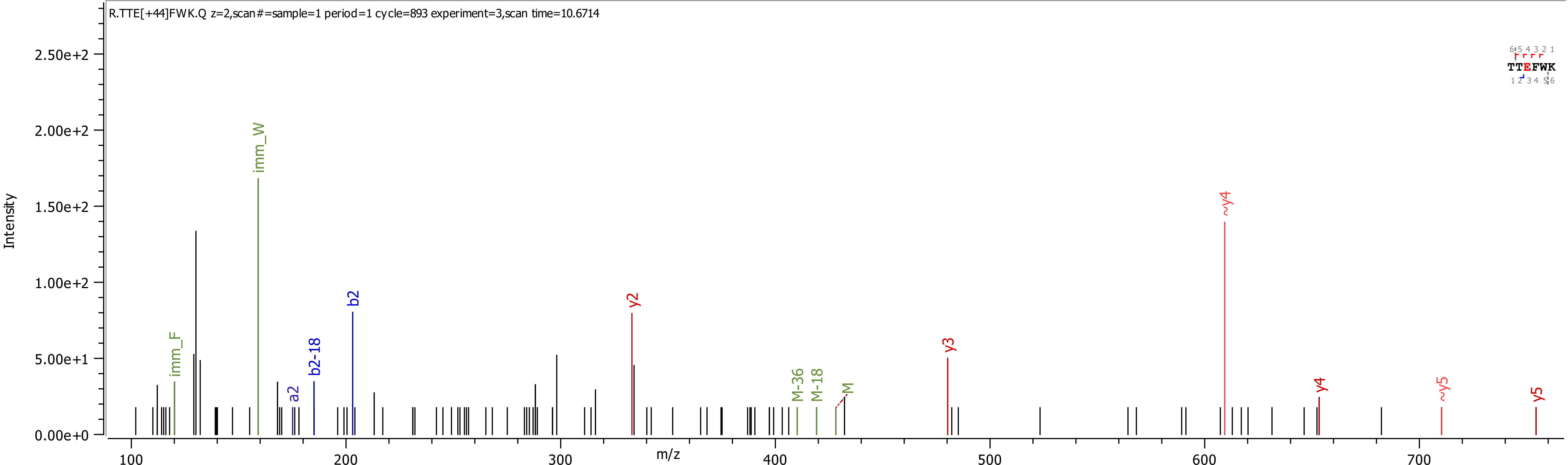
R.TTE[+44]FWK.Q z=2,scan#=sample=1 period=1 cycle=893 experiment=3,scan time=10.6714
TTEFWK
Intensity
m/z
imm_W
imm_F
a2
b2-18
b2
y2
M-36
M-18
M
y3
~y4
y4
~y5
y5

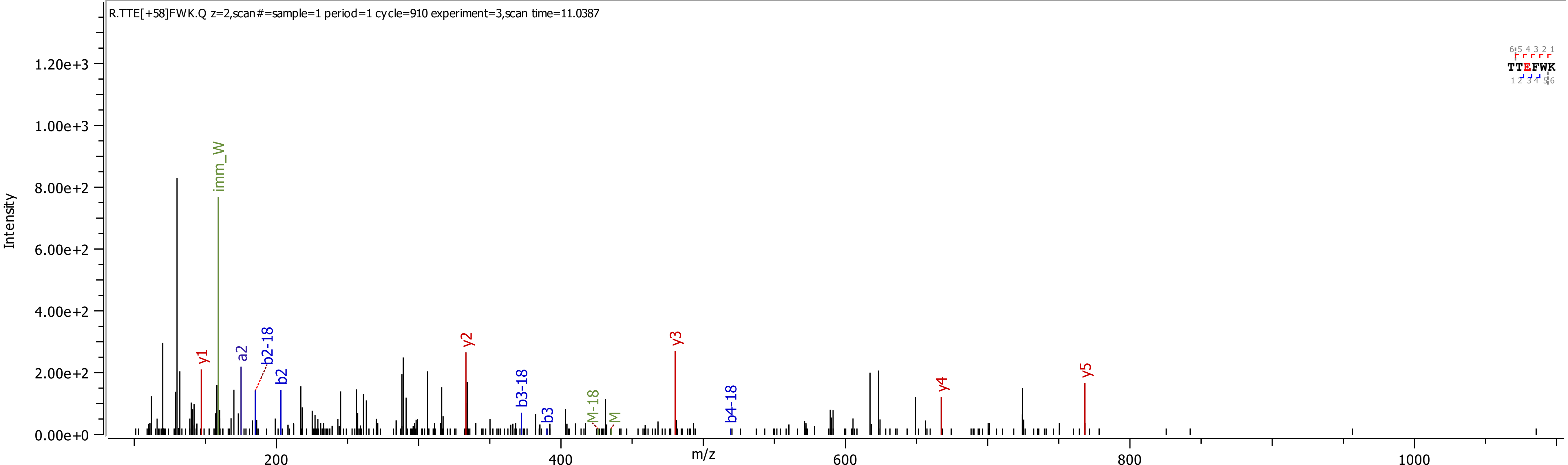
R.TTE[+58]FWK.Q z=2,scan#=sample=1 period=1 cycle=910 experiment=3,scan time=11.0387
Intensity
m/z
TTEFWK
imm_W
y1
a2
b2-18
b2
y2
b3-18
b3
M-18
M
y3
b4-18
y4
y5
1.20e+3
1.00e+3
8.00e+2
6.00e+2
4.00e+2
2.00e+2
0.00e+0
200
400
600
800
1000

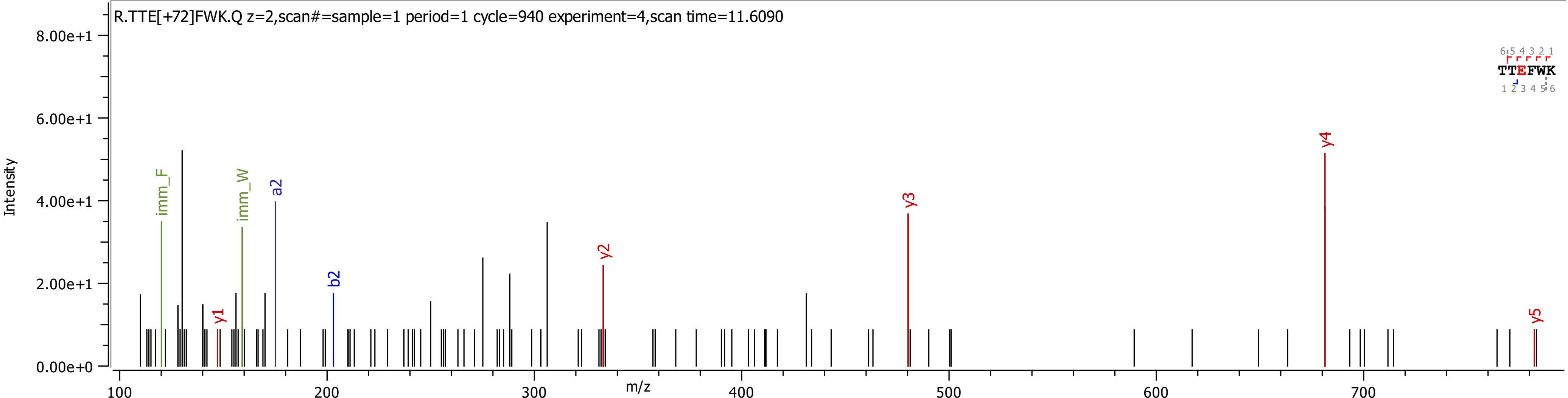
R.TTE[+72]FWK.Q z=2,scan#=sample=1 period=1 cycle=940 experiment=4,scan time=11.6090
Intensity
8.00e+1
6.00e+1
4.00e+1
2.00e+1
0.00e+0
m/z
100
200
300
400
500
600
700
imm_F
imm_W
a2
b2
y1
y2
y3
y4
y5
6 5 4 3 2 1
TTEFWK
1 2 3 4 5 6

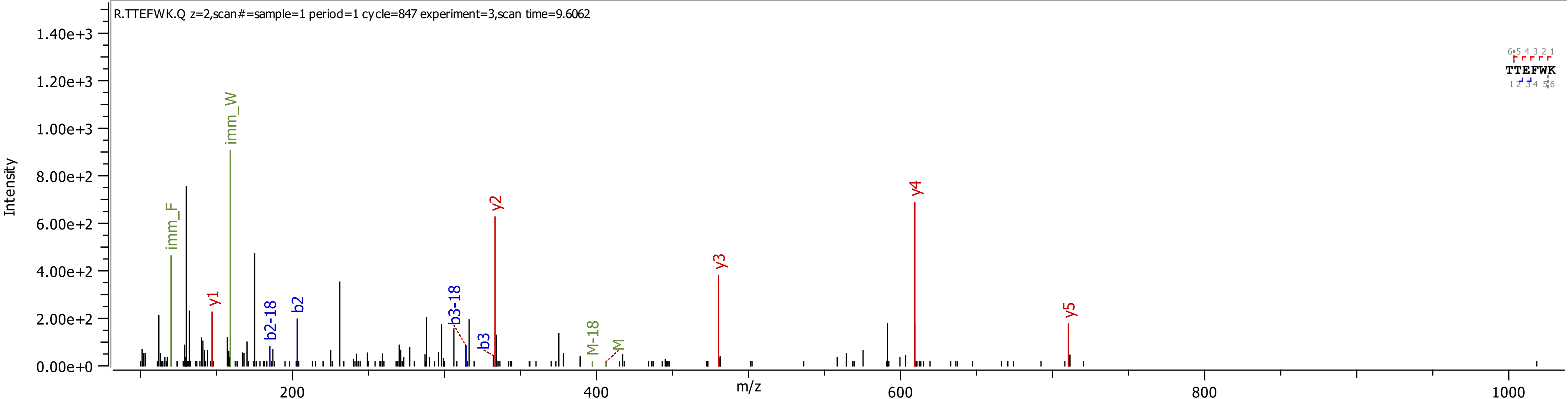

R.TTEFWK.Q z=2,scan#=sample=1 period=1 cycle=847 experiment=3,scan time=9.6062
Intensity
1.40e+3
1.20e+3
1.00e+3
8.00e+2
6.00e+2
4.00e+2
2.00e+2
0.00e+0
imm_F
imm_W
y1
b2-18
b2
b3-18
b3
y2
M-18
M
y3
y4
y5
TTEFWK
m/z
200
400
600
800
1000
